# Facilitating the design of combination therapy in cancer using multipartite network models: Emphasis on acute myeloid leukemia

**DOI:** 10.1101/2021.03.18.436040

**Authors:** Mohieddin Jafari, Mehdi Mirzaie, Jie Bao, Farnaz Barneh, Shuyu Zheng, Johanna Eriksson, Jing Tang

**Author notes:** **Corresponding authors:** Jing Tang, Mohieddin Jafari.

## Abstract

From the drug discovery perspective, combination therapy is recommended in cancer due to efficiency and safety compared to the common cytotoxic and single-targeted monotherapies. However, identifying effective drug combinations is time- and cost-consuming. Here, we offer a novel strategy of predicting potential drug combinations and patient subclasses by constructing multipartite networks using drug response data on patient samples. In the present study, we used Beat AML and GDSC, two comprehensive datasets based on patient-derived and cell line-based samples, to show the potential of multipartite network modeling in cancer combinatorial therapy. We used the median values of cell viability to compare drug potency and reconstruct a weighted bipartite network, which models the interaction of drugs and biological samples. Then, clusters of network communities were identified in two projected networks based on the topological structure of networks. Chemical structures, drug-target networks, protein-protein interactions, and signaling networks were used to corroborate the intra-cluster homogeneity. We further leveraged the community structures within the drug-based multipartite networks to discover effective multi-targeted drug combinations, and the synergy levels which were supported with more evidence using the DrugComb and the ALMANAC databases. Furthermore, we confirmed the potency of selective combinations of drugs against monotherapy *in vitro* experiment using three acute myeloid leukemia (AML) cell lines. Taken together, this study presents an innovative data-driven strategy based on multipartite networks to suggest potential drug combinations to improve treatment of AML.

## Introduction

Studies on cases with advanced cancers has shown that less than 10% of patients have actionable mutations, and improvement of outcomes was not observed in a randomized trial of precision medicine based on genomic profiles (1). The current limitation of genomics-centric personalized medicine falls short on the enormous heterogeneity and lack of actionable and sustainable treatment options. With a few exceptions, patient genomic signatures with clinical pathology do not typically predict drug responses. More precisely, cancer can principally be considered as a signaling disease, not a genetic disease. There is a wealth of data that has validated this hypothesis, including signaling behaviors involved in growth factor and nutrient responses, the process of entering and exiting the cell cycle, ensuring that chromosomes are segregated in an orderly, efficient and accurate manner during mitosis and apoptosis (2, 3). On the other hand, the complexity of crosstalk between signaling pathways necessitates to modulate multiple targets in cancer cells, otherwise lack of complete response, resistance, and relapse will emerge during the course of treatment.

Despite the fact that large amounts of small molecules or drugs have been tested on many cell lines or patient-derived samples, using single drugs as monotherapies to cure cancer might not be a promising strategy, as it is known that the complex interactions of various biological components can lead to drug resistance during the treatment of cancer (4–6). As a matter of fact, monotherapy and the slogan of “one target one drug” is inefficient to cure complex diseases such as cancer (7, 8). Combination therapy or polytherapy with synergistic drugs may achieve a more effective and safer outcome by targeting several targets in the same or separate pathways of the complex system (4). To better identify the synergistic drug combination based on precision medicine, we need ex-vivo drug screening to decipher the functional impact of cancer genomics at the phenotypic level to understand their interactions in the context of biological networks (9, 10). Therefore, understanding network biology may provide a unique opportunity to leverage the rich source of drug response data to offer network-based models for combinatorial therapy. These network models have shown promises for the development of clinical decision support tools to discriminate functional patient subclasses (11, 12). Even though there are networks reconstructed to model biological mechanisms of diseases and predict drug combination synergies based on molecular data (13–16), network models have not been systematically applied to patient data, i.e., drug response data of patient-derived samples to predict patient-customized drug combinations (14). Instead, the ex vivo drug response data are straightforwardly translated into the clinic for patient treatment since these individualized experiments represent the efficiency of some approved drugs on patient-derived primary cultures (17, 18).

In 2018, the Beat AML program reported a cohort of 672 tumor specimens collected from 531 patients, analyzing the ex vivo sensitivity for 122 drugs, as well as the mutational status and the gene expression signatures of the samples (19). Despite the dearth of large patient-related drug response datasets, some large cell line-based datasets, such as GDSC and ALMANAC, can offer a strong source of supporting evidence for predictions. The GDSC (Genomics of Drug Sensitivity in Cancer) database contains the response of 1001 cancer cell lines to 265 anti-cancer drugs, providing a rich source of information to connect genotypes with cellular phenotypes and to identify cancer-specific therapeutic options (20). The largest publicly accessible dataset for cancer combination drugs, i.e., ALMANAC, was recently published by the U.S. National Cancer Institute. This data collection contains more than 5,000 combinations of 104 investigational and licensed drugs, with synergies calculated against 60 cancer cell lines, resulting in more than 290,000 synergy scores (21). Moreover, DrugComb (https://drugcomb.org/), a web-based portal for the storage and study of drug combination screening datasets, offers a comprehensive visualization of drug combination susceptibility and synergy, which can significantly aid understanding of drug interactions at unique dosage levels. Drugcomb now has 751,498 drug combinations and 717,684 single drug screens from 37 trials, which relate to 2040 cell lines and 216 cancer forms (22).

In this study, we developed a network pharmacology approach to predict potential drug combinations for AML based on BeatAML dataset. We proposed a drug combination strategy using multipartite network modeling of ex vivo drug screening data. By ex vivo drug response data, we directly accessed the individual phenotypes of the patients’ cancer cells, and by network modeling, we demonstrated the similarity of drugs and AML patients. Then, we used the community structures within the drug-based multipartite networks to discover effective multi-targeted drug combination regimens. Our predicted combinations of drugs were only suggested on the basis of the phenotypic interactions of the cancer cells or patient samples to the drugs without prior understanding of the genetic origin or molecular understanding of the disease.

## Methods

The entire workflow of the present study is illustrated in Fig. 1. The weighted bipartite network is constructed using BeatAML data set. This data set is a collaborative research program of 11 academic medical centers providing data on AML samples offering genomics, clinical, and drug responses. It includes a cohort study of 672 tumor specimens collected from 531 patients analyzing 122 drug responses. In order to construct a weighted bipartite network, the best response read out of drug potency was defined using information-based measures. Then two unipartite networks obtained using network projection on samples and drugs. In the next step, communities of two projected networks were extracted and intra-cluster homogeneity analysis was performed using the similarity of drugs and patients/ cell members based on available gene expression profiles for patients and protein-protein interaction network and biological pathways. The drug candidates for drug combination were selected at two different communities and a high-throughput drug screening was used to assess their synergetic effects.

**Figure 1:**
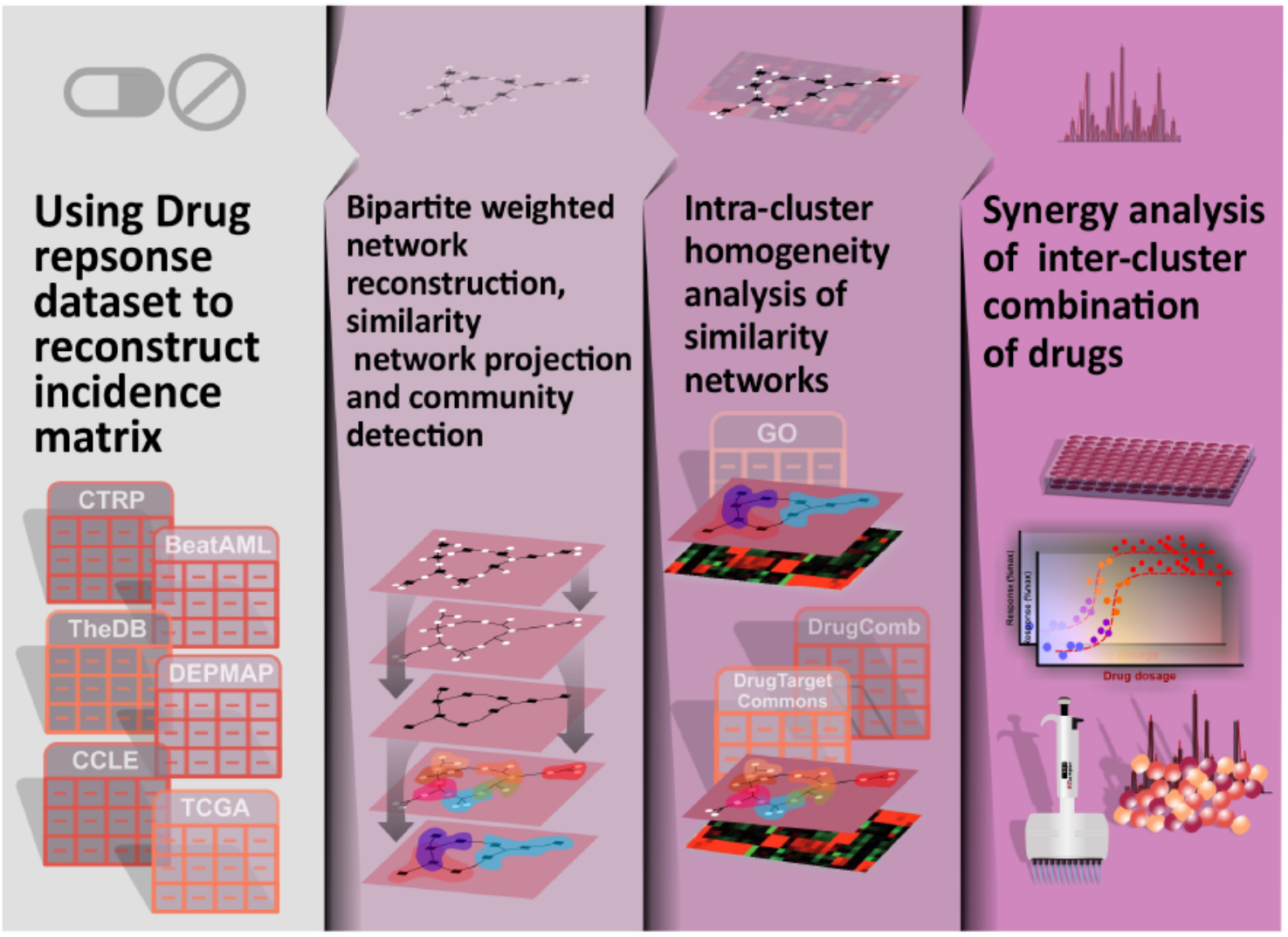
Flowchart of the study. Data collection started from existing drug response databases, then followed by incidence matrix extraction, weighted bipartite network reconstruction, network projection, and community detection. In the next step, the intra-cluster homogeneity analysis was conducted using the similarity of drug and patient/cell members of all clusters according to available gene expression profiles, drug-target interactions, protein-protein interactions, and biological pathways. Finally, a high-throughput drug screening experiment was used to assess the synergistic behavior of the proposed drug combinations.

### Defining the response read-out for drug screening experiments

Pharmacogenomic studies require extensive standardization, to avoid inconsistency of drug responses data for further research and unbiased predictions (23, 24). Therefore, first, we controlled the quality of cell viability data to select the potent compounds. To aim this, we examined the raw datasets in terms of the availability of replicated data and outlier detection, followed by assessment of distribution, pairwise correlation, and homoscedasticity analyses to select the best response read-out or “measure” of drug potency. This analysis was performed using information-based nonparametric measures available in the Minerva package (25) by computing Maximal Information Coefficient (MIC), Maximum Edge Value (MEV), and Maximum Asymmetry Score (MAS). Furthermore, the relative and absolute IC50 (i.e., IC50 measures, which were computed based on the top and bottom plateaus of the curve or based on the blank and the positive control values, respectively), RI value (Relative Inhibition), AUC (Area Under Curve of drug-response fitted line), and the median of cell viability in the drug response experiments were assessed to select the best measurement. The chosen measurement was later used as a weight value for the edges in the weighted bipartite network reconstruction.

### Reconstruction and analysis of the bipartite network model

In our bipartite network model, one group of nodes contained the small molecule drugs and the other group the cancer cell lines or patient samples depends on the dataset. The edges were defined by incidence matrices derived from the min-max normalized values:

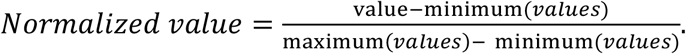

This normalization transforms these values, that are indicative of the potency of small molecules on cancer cell lines or patient samples, into a decimal between 0 and 1. Next, we projected the bipartite network into the two similarity networks, i.e., drug similarity network and sample similarity network. In the network projection, two unipartite graphs are derived from a bipartite graph, which result in deducing the node’s relationships of the same type. In this work, we projected similarity networks, which take into account the edge weights in the bipartite network. Then, we studied the general properties of the networks such as network heterogeneity, centralization, and clustering coefficients. The critical step was community detection within the projected networks in order to discern functionally similar drugs and similar cells or patients in terms of drug response. The modularity index was used to determine the best community detection algorithms, including infomap (26), fast greedy (27), and spinglass (28). In the next step, we explored the network modules to propose a strategy for drug combination design.

### Computational corroboration

Multiple computational methods were applied to validate the predictions of the drug combinations and patient or cell stratification. The validation of the community structures is similar to the general cluster quality assessment method, and we assessed the clustering performance by matching clustering structures to prior knowledge. This validation will be a basis to support the possible drug combination designs. In other words, the combination of distinct drugs in terms of chemical structure, target profile, and implicated biological pathways is most likely more efficient than similar drugs (7). Therefore, we used the drug-target network, protein-protein interactions, and signaling networks to justify the similarity of cluster elements. Thus, Chembl (29), drug target commons (DTC) (30), KEGG (31), and Omnipath database (32) were used to extract prior annotations about the drugs and their targets. To compare the chemical structures of the drugs, Simplified Molecular Input Line Entry System (SMILES) of the drug molecules were retrieved and transformed into extended connectivity fingerprint (ECFP), in order to assess the Dice similarity of the molecules. The Dice similarity is one of the standard metrics for molecular similarity calculations in which *S_A,B_* = 2*c*/(*a* + *b*), where *a* is the number of ON bits in molecule A, *b* is the number of ON bits in molecule B, and *c* is the number of ON bits in both A and B molecules (33). Also, the corresponding gene expression profiles were used to assess similarity within patient or cell line modules in the sample similarity networks. For reads per kilobase per million (RPKM) with negative values and counts per million (CPM), we used Harmonic similarity and Jaccard distance, respectively as follows:

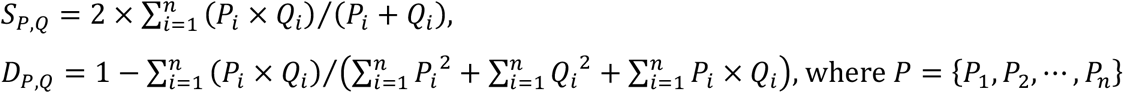

and *Q* = {*Q*_1_, *Q*_2_, … *Q_n_*} denote the vector of gene expression values for patients or cell lines and *n* is the number of genes. In all cases, the similarity or distance scores were compared with the random grouping of small molecules or biological samples to perform statistical testing.

The synergy scores provided by the DrugComb database (34) were used to corroborate synergistic combinations of our network-based predictions, including HSA, Bliss, Loewe, ZIP, CSS and S. Let’s assume that drug 1 at dose x_1_ and drug 2 at dose *x_2_* is used to produce effects of y_1_ and y_2_, and y_c_ is the effect of their combination. Drug’s effect is usually measured as percentage of cell death and a drug combination is classified as synergetic, antagonistic or non-interactive (35). The expected effect denoted by y_e_ represents a non-interactive level and it is quantified based on a reference model. Several mathematical models have been introduced to calculate the expected effect by assuming specific principals. The HSA model (36) considers the expected combination effect as the maximum of single drug effects, i.e. y_e_ = max(y_1_,y_2_). The loewe model (37) assumes that an individual drug produces y_e_ at a higher dose than in the combination. In the Bliss model (38), y_e_ is the effect of the two drugs when they act independently. The ZIP model (35) consider the assumptions of the Loewe and Bliss models by assuming that at reference model two drugs do not potentiate each other. CSS determines the sensitivity of a drug pair and S synergy is based on the difference between the drug combination and the single drug dose response curves (39).

### Cell culture and reagents

AML cell lines MOLM-16, NOMO-1, and OCI-AML3 were a kind gift from Prof. Caroline Heckman (University of Helsinki, Finland). MOLM-16 and NOMO-1 were cultured in RPMI-1640 medium (Gibco/Thermo Fisher Scientific, Waltham, MA, USA) and OCI-AML3 in α-MEM (with nucleosides; Gibco/Thermo Fisher Scientific) supplemented with GlutaMAX (Gibco CTS/Thermo Fisher Scientific), fetal bovine serum (20% for MOLM-16 and OCI-AML3; 10% for NOMO-1) and antibiotics.

### Drug combination testing

The compounds dissolved in DMSO were plated using Beckman Coulter Echo 550 Liquid Handler (Beckman Coulter, Indianapolis, IN, USA) in combinations, seven concentrations for each compound in half-log dilution series with 2.5/7.5/25 nl volumes, covering a 1,000-fold concentration range on black clear-bottom TC treated 384-well plates (Corning #3764, Corning, NY, USA). All doses were randomized across the plate to minimize any plate affects. As positive (total killing) and negative (non-effective) controls 100 μM benzethonium chloride and 0.2% dimethyl sulfoxide (DMSO) were used, respectively.

Cells were plated on pre-administered compound plates in 25 μl (2500, 2000, or 1250 cells per well for MOLM-16, NOMO-1, OCI-AML3 cell lines, respectively) using BioTek MultiFlo FX RAD (5 μl cassette) (Biotek, Winooski, VT, USA) and incubated for 72 hours at 37oC, 5% CO2. Cell viability was then determined by dispensing 25 μl of Cell Titer Glow 2.0 reagent (Promega, Madison, WI, USA). Plates were incubated for 5 min and centrifuged for 5 min (173 × g) before reading luminescence with PHERAstar FS multimode plate reader (BMG Labtech, Ortenberg, Germany).

## Results

### Defining the edge weight of bipartite networks

In the Beat AML dataset, a set of 122 inhibitor drugs were used against 531 patient-derived AML samples. The spectra of low to high potency of drugs were observed across the patient-derived samples. However, this panel of small molecule inhibitors was selected according to their activity against the proteins involved in tyrosine-dependent and non-tyrosine kinase pathways particularly for AML (19). In the first step, we determined a weight value of drug-sample interaction to be used in the bipartite network reconstruction. This value should describe the most potent compounds for inhibiting tumor cells based on the drug sensitivity analysis. In addition to the relative and absolute IC_50_, RI value (39), and AUC, we calculated the median of cell viability in the drug response experiments. The distribution of these measures have been evaluated in terms of normality, skewness, and modality (Fig. 2) to choose the best measure as a weight in the bipartite network. The relationship of Median to AUC was a high positive value (with the highest r Pearson correlation coefficient ~ 0.94). The distribution of Medians was unimodal in contrast to IC_50_ distributions, homoscedastic contrary to RI distribution, and more symmetric (non-skewed) compared to AUC distribution. In addition to investigating the linear relationship, i.e., Pearson correlation analysis, we computed MIC (Maximal Information Coefficient), which measures the relationship strength, and MEV (Maximum Edge Value) to check the closeness of the relationship to being a function. Interestingly, the relationship between Median and AUC displayed higher MAS and MEV (~0.75) compared with the relationship of RI and AUC, meaning that Median has a stronger association with AUC. Therefore, we have chosen the inverse of the actual median as the weight of drug-patient interaction.

**Figure 2:**
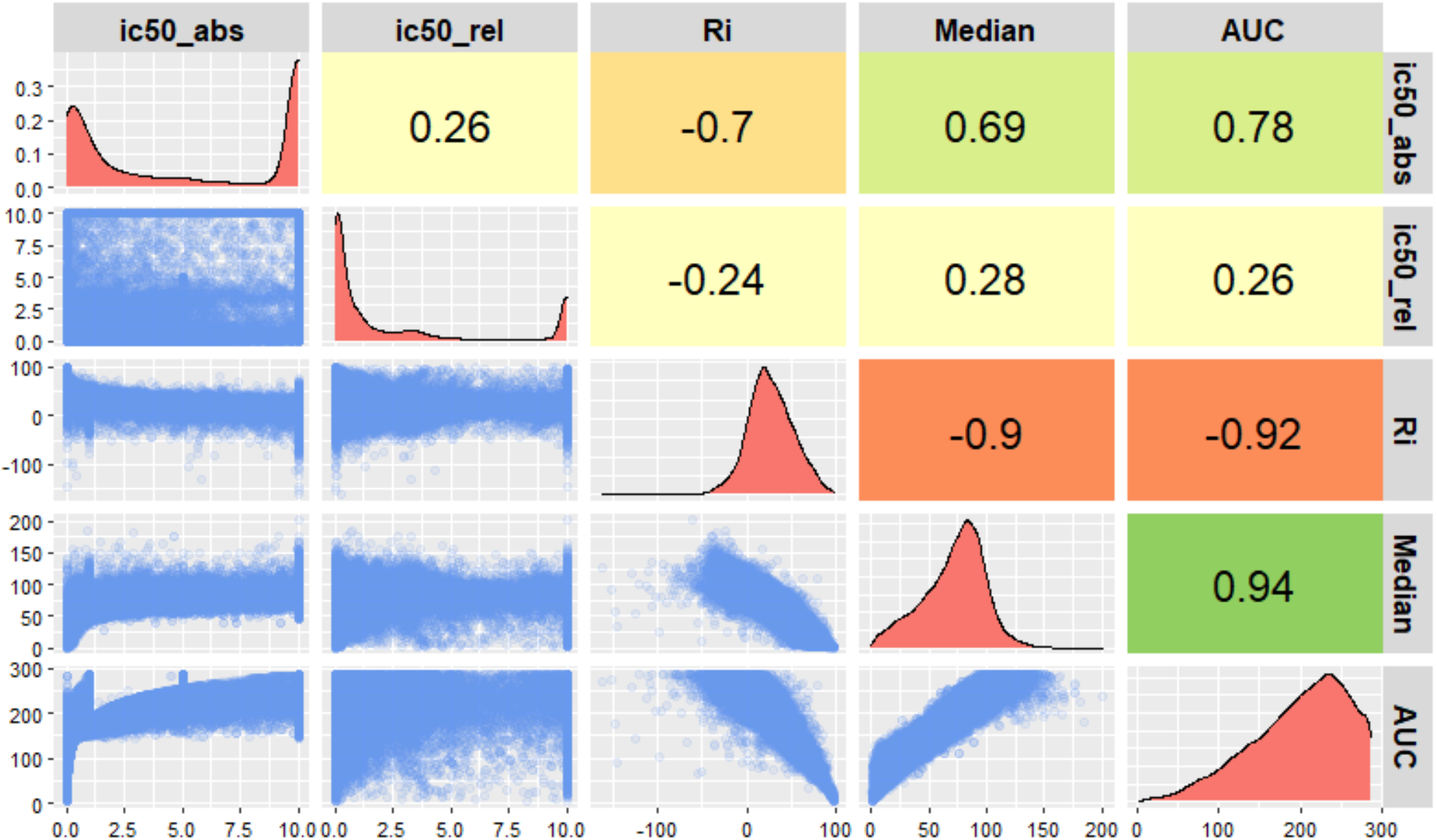
Comparison of different measures for drug response experiments in the Beat AML study. The lower triangle of this pairwise comparison matrix shows the pairwise scatter plots for ic50_abs (Absolute IC_50_), ic50_rel (Relative IC_50_), RI (Relative Inhibition), Median (the median of cell viability), and AUC (Area Under Curve of cell viability fitted line). The diagonal panel describes the histogram of each measure individually. The upper triangle represents the Pearson correlation coefficients of the corresponding pairwise comparisons.

### Analysis of bipartite networks

In the next step, the maximum square submatrix of patient samples and small molecules was used as the incidence matrix of the bipartite network. To be specific, we selected the list-wise deletion strategy to remove missing values and we used the complete cases of both variables. The downstream analysis was done on an undirected weighted bigraph consisting of 176 (88+88) nodes and 7744 edges (Fig. 3A). The distribution of the min-max normalized edge weights indicated positive skewness, indicating that the cells were not highly sensitive to most drugs. All the performed analyses were also carried out for the GDSC dataset as a proof of concept. The undirected weighted bigraph of the GDSC dataset consisted of 532 (266+266) nodes and 70,756 edges (Fig. 3C). The distribution of the min-max normalized edge weights showed positive skewness in this dataset as well (Fig 3D), indicating again low potency for most of the drugs. Therefore, exploring the best combination is not straightforward, and categorizing drug-sample interactions seems to be required. Following the projection of these bigraphs as outlined in Fig. 3, two projected graphs called the patient similarity network (PSN) and drug similarity network (DSN) were reconstructed, such that each edge was obtained by the multiplication of weighted incidence matrix. Thus, the edge weights of the projected graphs indicate the profile similarities of patient samples in PSN and small molecule inhibitors in DSN. Note that the edge weight values in DSNs and PSNs are distinct due to different matrix multiplications.

**Figure 3:**
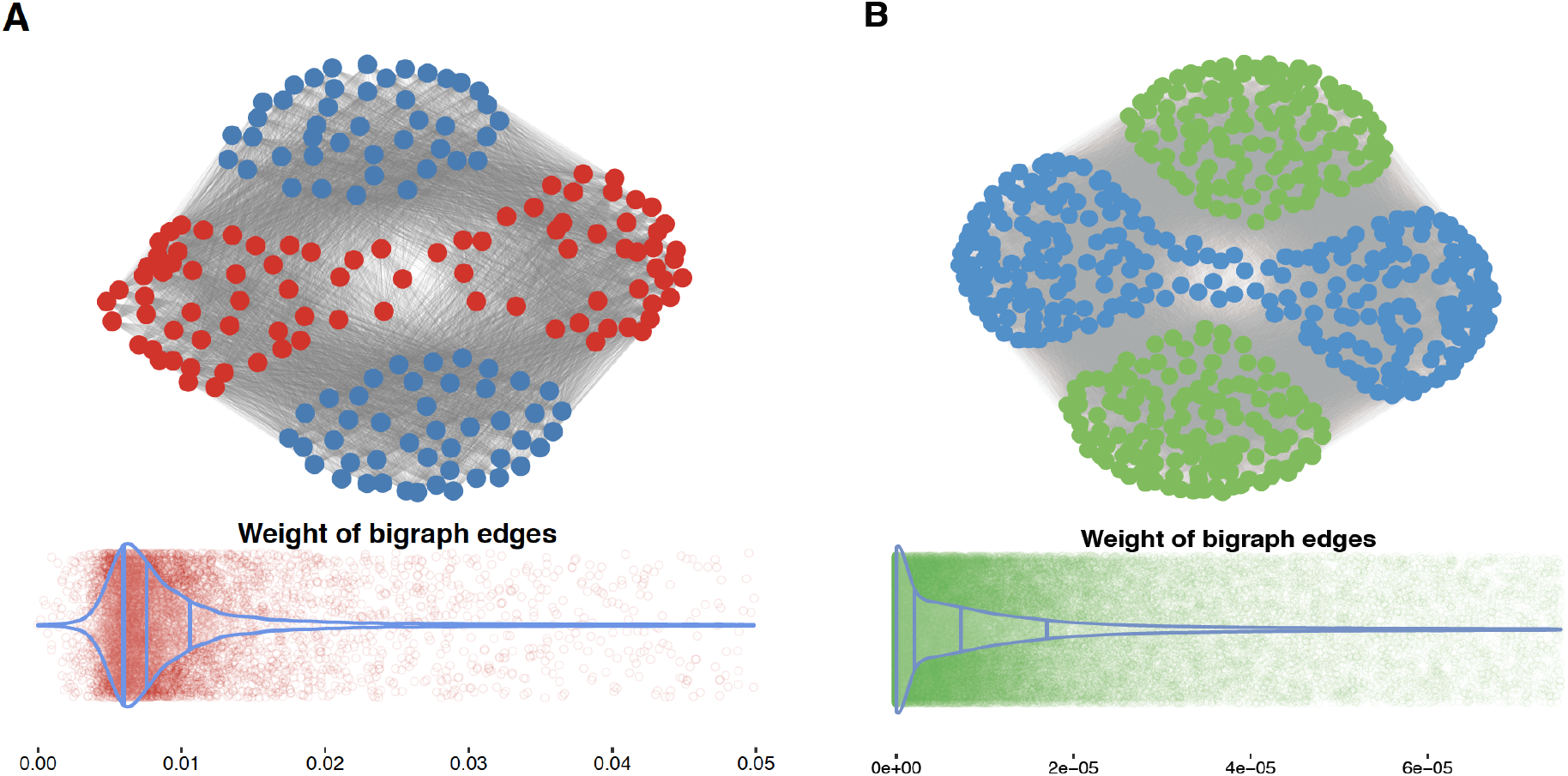
Bigraphs of cancer datasets. The general overview of the bipartite graphs for the Beat AML (A) and GDSC (B) datasets are represented with the blue nodes as small molecule inhibitors, and red and green nodes as patient-derived and cell line samples, respectively. The distributions of edge weight values are also depicted using violin plots with scatter plots.

The PSN and DSN of the Beat AML dataset contained 88 nodes and 3828 edges (Fig. 4), while in the GDSC projected similarity networks, there were 266 nodes and 35,378 edges (data not shown). In Fig. 4, the larger node size, the more sensitive patient-derived samples and more potent drugs. In this subset of Beat AML dataset without missing data, patient 16-00627 was found to be the most sensitive and SNS-032 was the most potent inhibitor (See Supplementary Fig. 1). The community detection was subsequently done for both similarity networks via optimizing a modularity score, resulting in two communities for DSN with 50 and 38 small molecules, and two communities for PSN with 39 and 49 patient samples. In other words, we identified two clusters of patients with distinctive drug response profiles, suggesting two subcategories of the disease. Also, we detected two clusters of small molecules, which pointed disparate inhibiting patterns on the patient samples. In the following steps, we presented evidence of the consistency of cluster members in both networks using prior knowledge.

**Figure 4:**
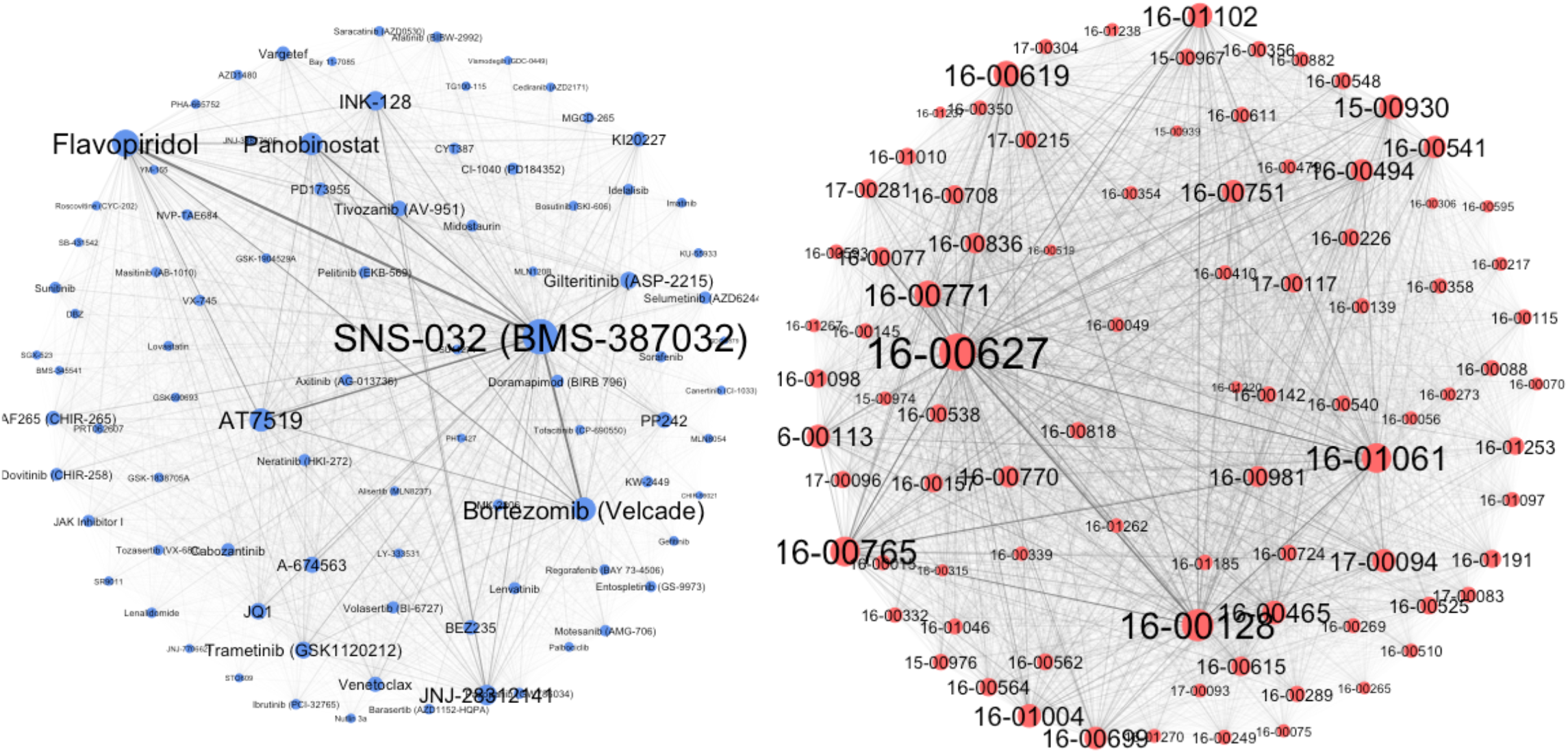
Drug and patient similarity networks of the Beat AML dataset. The force-directed layout was selected to depict both networks. The thickness of edges corresponds to the edge weight of the original bipartite networks after network projection considering the weight values. The edge thickness represents the weight value of similarity between each pair of patient samples or small molecules. The node size is proportional to the strength of each node, which is the sum of the edge weights of the adjacent edges for each node.

### Intra-cluster homogeneity analysis of similarity networks

#### Drug similarity network

Focusing on small molecules, we presumed that inhibitory molecules with correlated effects on cell survival were likely to have similar structures, purposes, and functions (40–43). Therefore, we evaluated the similarity of SMILES structures and the analogy of protein targets and biological pathways of detected clusters in the DSNs against random groupings of molecules. The distribution of the Dice similarity of SMILES structures differed significantly between the random grouping and the clusters based on network topology (Fig. 5A). The statistical test of the median difference also resulted in the lowest p-values for the both pairwise two-sample Wilcoxon and Kruskal-Wallis rank sum test (p-value < 2e-16). To evaluate their target similarities, we explored the protein targets of the small molecule inhibitors and examined the number of the target intersections of small molecule pairs within the clusters. In this analysis, drug target commons (DTC) and OmniPath were applied to explore the binding targets of small molecules and second-order node neighbors (secondary target) in the signaling network, respectively. Assuming that proteins usually correspond to multiple signaling pathways, the KEGG database was used to check the number of pathway intersections of the protein targets for each pair of small molecules. The median similarity measures of the intersections within the network clusters were significantly higher than those for a large set of random pairs of small molecules (Fig. 5B-D) (p-value < 2.2e-16, Kruskal-Wallis rank sum test). Analysis of the GDSC dataset gave similar results (Fig. 6) (p-value < 2.2e-16, Kruskal-Wallis rank sum test), suggesting that our method is also reproducible for the analysis of cell line-based datasets.

**Figure 5:**
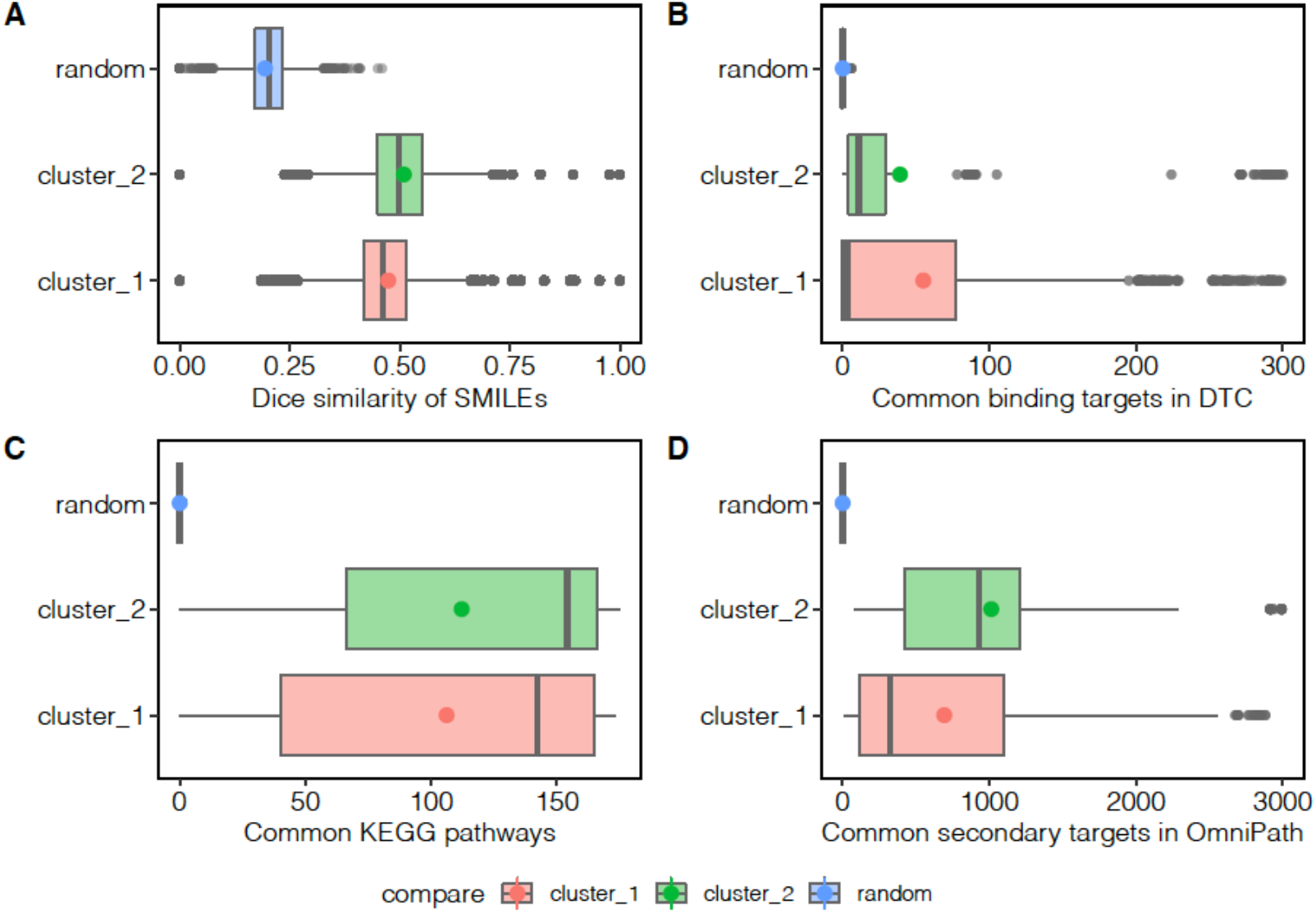
Beat AML intra-cluster homogeneity analysis. (A) The distribution of SMILE structure similarities of DSN clusters compared to random grouping. (B) The distribution of pairwise intersection size of binding protein targets, (C) corresponding KEGG pathways, and (D) secondary targets in OmniPath database.

**Figure 6:**
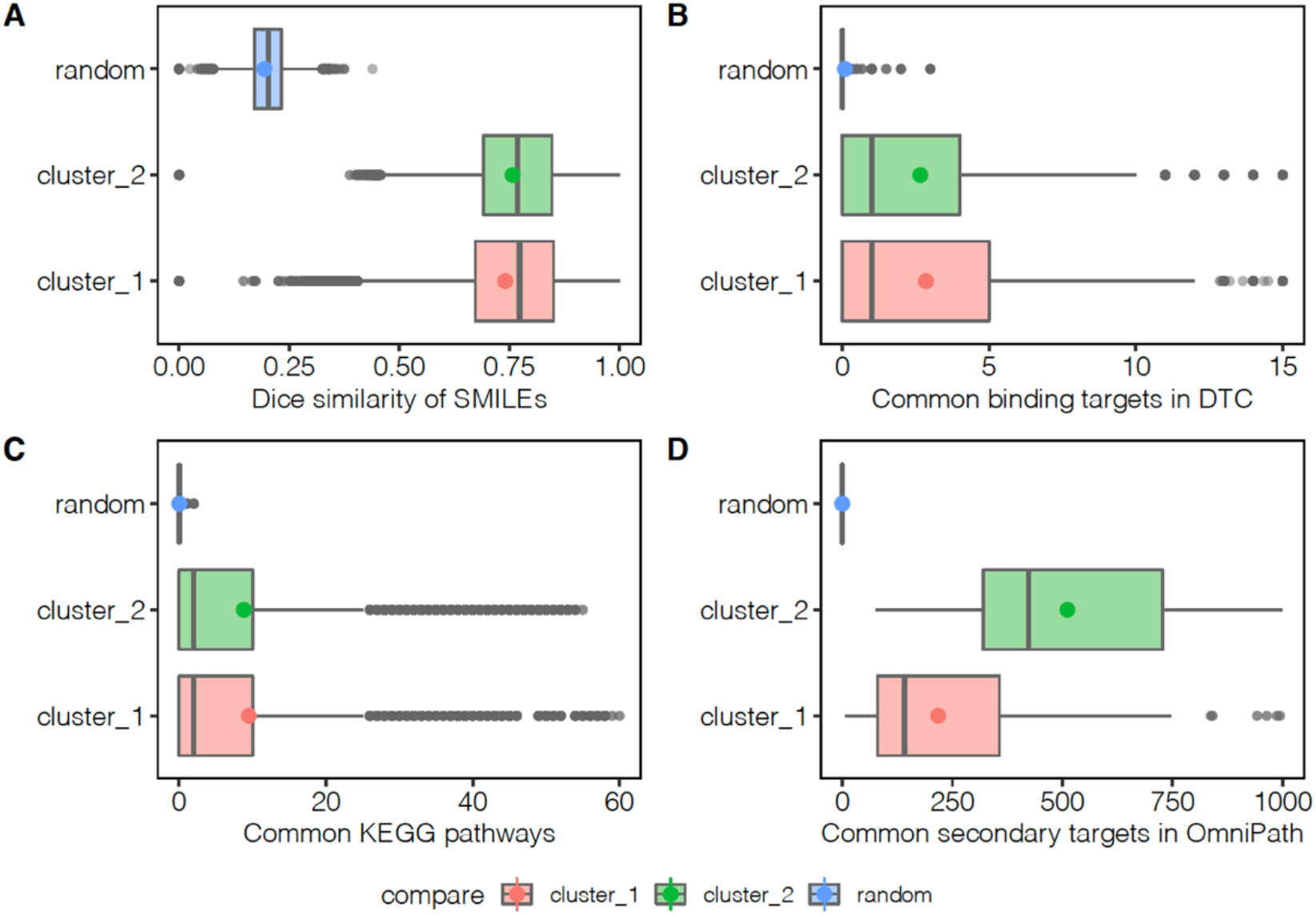
GDSC intra-cluster homogeneity analysis. (A) The distribution of SMILE structure similarities of DSN clusters compared to random grouping. (B) The distribution of pairwise intersection size of immediate protein targets, (C) corresponding KEGG pathways, and (D) second targets in OmniPath database.

#### Patient and cell-line similarity network

Next, we examined the member consistency of the patient clusters in the PSN using other available data from the patient samples in the Beat AML dataset. The gene expression data including RPKM and CPM of the samples were utilized to check pairwise similarity of the cluster members. The similarity measures were also computed for a large set of random pairs of patient samples to compare with our patient stratification using network clustering. When we compared the harmonic mean similarities of the RPKM values, the pairwise similarities of patients within the clusters were significantly larger than those of the randomly selected patients (p-value < 2.2e-16, Kruskal-Wallis rank sum test) (Fig. 7A). For the CPM dataset, the distributions of Jaccard distance were shown, where the distances within the clusters were found to be statistically lower than those in the random group (p-value = 4.655e-05, Kruskal-Wallis rank sum test) (Fig. 7B). For the GDSC dataset, we used the expression profiles of signature genes provided by the SPEED platform (44). Then, differentially expressed genes have been used to provide gene signatures of perturbed cancer-related pathways. In this dataset, there are 11 activity scores to represent the activity level of 11 well-known pathways for each cell line. Therefore, we compared the distance distributions of cell line pairs in the clusters to a set of random pairs of cells lines. Our findings indicated that the distances within the clusters were much lower than those in the random grouping (p-value = 6.94e-08, Kruskal-Wallis rank sum test) (Fig. 7C).

**Figure 7:**
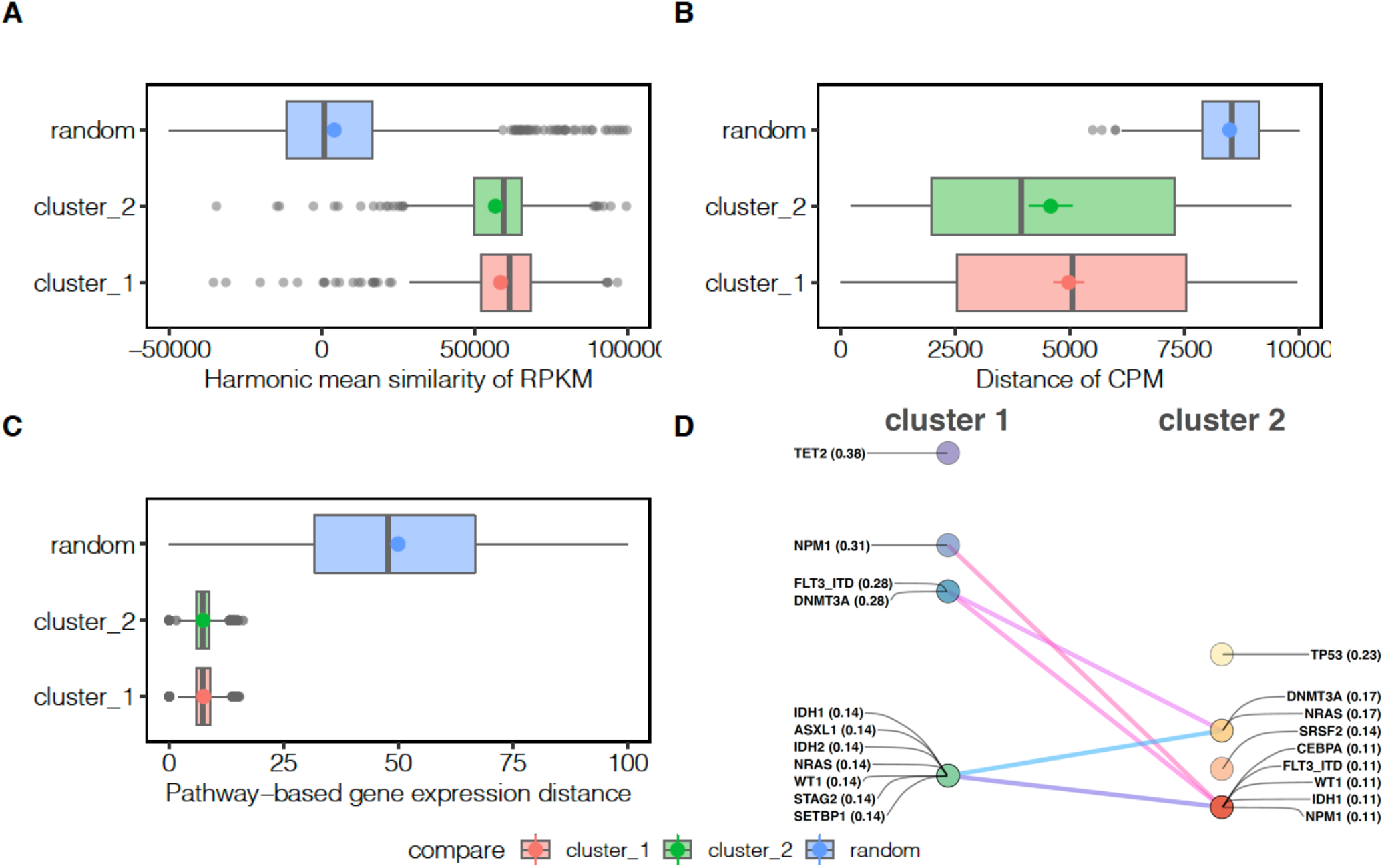
Validation of network communities of PSN. (A) The distribution of similarity of RPKM in Beat AML dataset. (B) The distribution of distances of CPM in Beat AML dataset. (C) The distribution of distances of pathway-based activity scores in GDSC dataset. (D) The frequently mutated genes in the clusters of Beat AML patients. The non-benign mutations with the possibility of being damaging greater than 0.5 were selected to find the intersection of mutated genes. The gene names are shown with the relative frequency of mutated genes in each cluster (e.g., NPM1 – 0.38 indicates 38% of the patients in cluster 1 has this mutation). The lines between mutated genes illustrate the rank shift in two clusters.

The Beat AML study also provided the first detailed view of mutational landscape in AML (19). Here, we used the dataset of non-benign gene mutations to characterize both clusters of patient samples. As shown in Fig. 7D, both clusters of patients demonstrate a distinct profile of gene mutations in terms of involved genes and the ranks of genes based on frequency. Previously, Tyner et al. have highlighted the importance of *TP53* and *ASXL* gene mutations, both responsible for a broad drug resistance pattern. They further showed that mutations of certain genes may identify disease subgroups sensitive to certain inhibitors. For example, they found that patients with FLT3-ITD and *NPM1* mutations were sensitive to SYK inhibitors. Interestingly, our molecular-independent network-based approach to characterize patient samples also captured the significance of the mutations above. Furthermore, our findings indicate that *TP53*, *DNMT3A*, and *NRAS* were the most frequently mutated genes in one of the patient clusters, while *TET2* and *NPM1* were the most frequently mutated genes in the other cluster along with the FLT3-ITD mutation. These results suggested that the phenotype-level of information in drug response data can clearly corroborate the genotype-level information to stratify patients more effectively.

### Inter-cluster design strategy for drug combinations

We assumed that the best drug combination strategy is the selection of one drug from each cluster to block potential drug resistance mechanisms and cancer recurrence. A common drug combination design could be the use of the most effective drugs of each cluster to prohibit cancer cells more effectively. However, other pharmacologic evidence can encourage the choice of the best combination of drugs more specifically. As the focus in drug combination studies also lies in finding the most synergistic drug combinations, previously reported studies were used to explore the synergy values (i.e. the degree of interactions) of drug combinations. We checked first whether combinations of the top five drugs (based on the median values of cell viability) of each cluster in the Beat AML and GDSC datasets (Table1), are found in the DrugComb database.

**Table 1:** Top-five small molecules in each cluster of drug similarity networks.

|  | Cluster 1 | Cluster 2 |
| --- | --- | --- |
| Beat AML | SNS-032 (BMS-387032) | Dovitinib (CHIR-258) |
|  | Flavopiridol | Nintedanib |
|  | Panobinostat | Doramapimod (BIRB 796) |
|  | AT7519 | KI20227 |
|  | Bortezomib (Velcade) | Cabozantinib |
| GDSC | Amuvatinib | Sepantronium bromide (YM-155) |
|  | GSK690693 | Belinostat |
|  | Vinblastine | AT-7519 |
|  | AS605240 | CAY10603 |
|  | HG6-64-1 | AR-42 |

However, there were no reports regarding the 25 possible combinations of these drugs, so we aimed to compare the average of synergy values for these ten drugs in the whole database. Fig. 8 showed the distributions of synergy values in DrugComb, highlighting the mean of synergy of the bottom and top five drugs in each network clusters. This analysis revealed the reasonably high potential of combinations of the top five drugs according to the average median values in both Beat AML and GDSC datasets (p-value = 2.96e-02 and p-value = 3.56e-02, Wilcox rank sum test, respectively).

**Figure 8:**
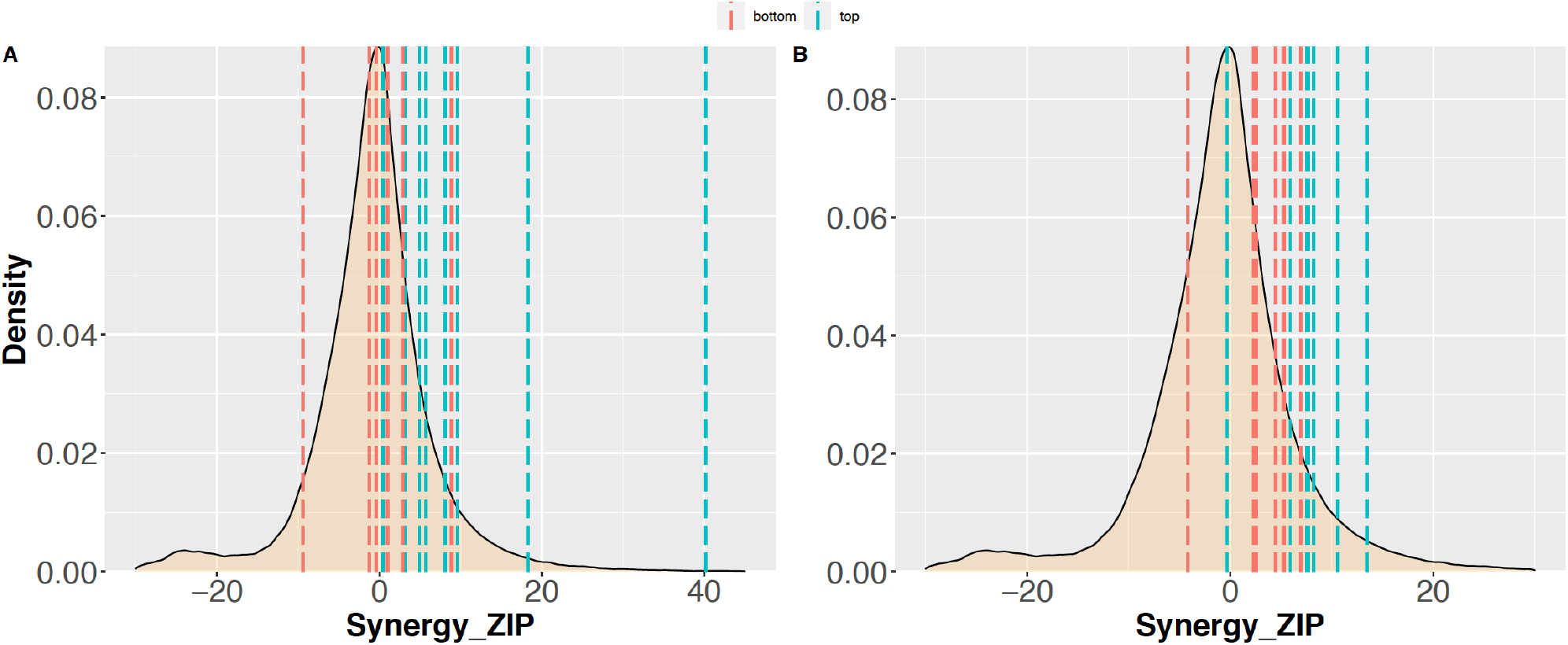
Distribution of drug combination synergy scores in the DrugComb database. The median of synergy zip score for top and bottom five drugs are presented by dashed lines in Beat AML dataset (A) and in the GDSC dataset (B).

#### Synergy analysis of inter-cluster combination of drugs

For further validation of our strategy of predicting synergistic drug combinations using network modeling, we focused on the ALMANAC dataset (21), which has 1,892,650 combinations of 103 inhibitors tested on 60 cell lines. The same procedure as described in Fig. 1 was implemented to extract the drug modules in the drug similarity network according to available single drug experiments in this dataset. The median inhibition values of the single drug responses on cell lines were used as weight values in the bipartite drug-cell line network. Using the projection of the weighted drug similarity network, the clusters of drugs with similar effect profiles on cell lines were extracted.

According to our predefined assumption, the combinations of drugs from different clusters were used as the positive group and the combinations of drugs within the clusters as the negative group. Then, we retrieved the synergy and sensitivity scores of the combinations for both groups using the DrugComb computed values, i.e., highest single agent (HSA), zero-interaction potency (ZIP), Bliss, Loewe, combinational sensitivity score (CSS), and S synergy. As shown in Fig. 9A, the positive group of drug combinations exhibited a significantly higher value of drug synergy than the negative group. This result was evident for all types of synergy measures, indicating the superiority of the strategy of using inter-cluster drug combinations. These data also indicated the efficiency of our proposed network-based modeling to discern drugs with similar profiles of effect on biological samples. Also, our proposed strategy of drug combination using the drugs of contrary clusters is more likely to acquire higher drug synergy and potency.

**Figure 9:**
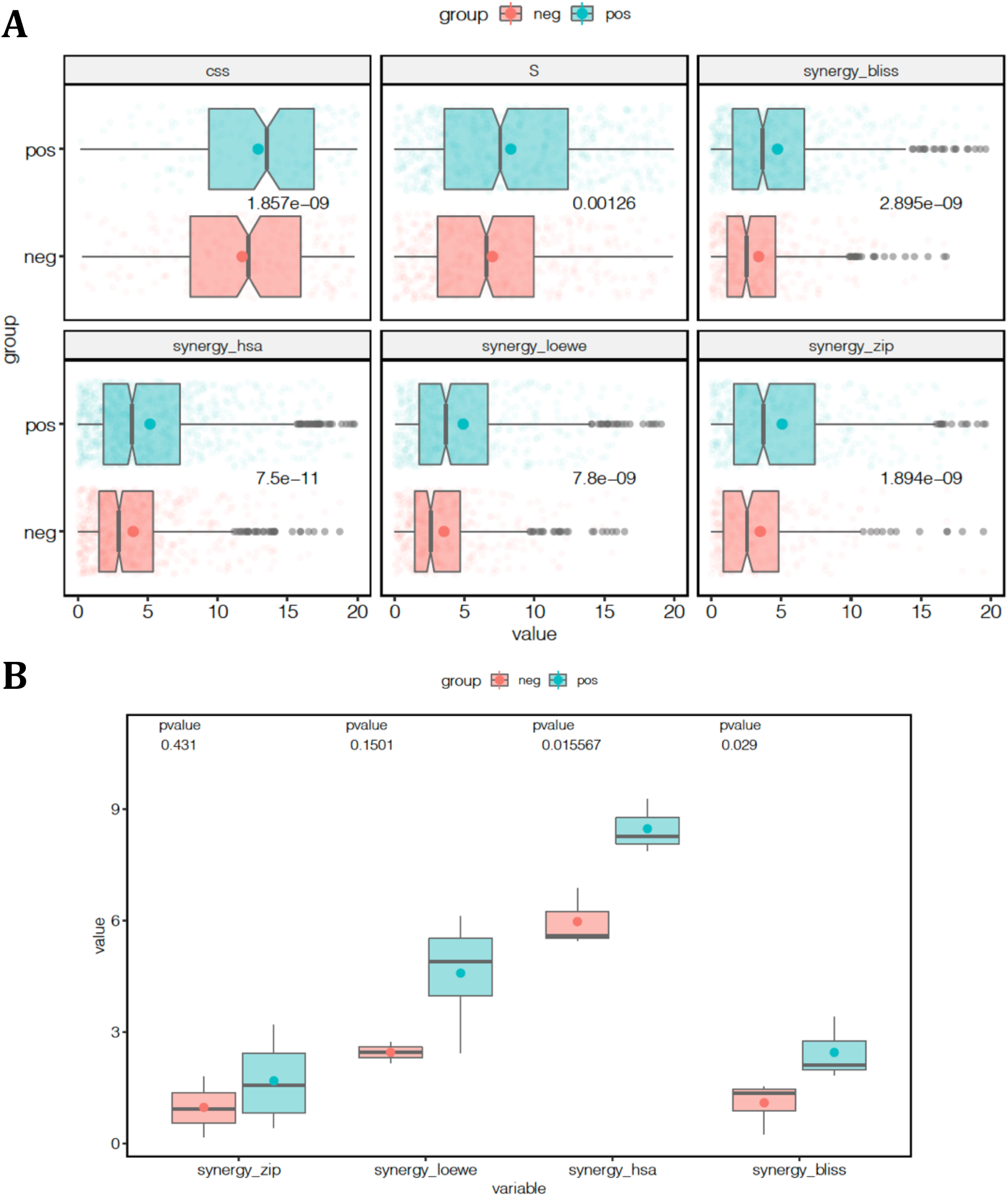
Synergy of drug combinations. (A) The combinational sensitivity (CSS) and synergy scores (S, synergy_bliss, synergy_hsa, synergy_loewe, and synergy_zip) of drug combinations in the ALMANAC dataset. The top five drugs of cluster 1 (Cabazitaxel, 5-FU, Cytarabine hydrochloride, Methotrexate, Bleomycin) and cluster 2 (CHEMBL17639, Gefitinib, Ixabepilone, Dexrazoxane, Eloxatin) for inter-cluster and intra-cluster combinations are shown in blue and red as the positive and negative groups, respectively. Each plot contains a scatter plot, notch box plot, and mean values for each group. The p-value represents the one-sided Student’s t-test significance for each score separately. (B) Measured synergy of drug combination scores in the experimental validation of selected drugs based on the network modeling of the Beat AML data in three AML cell lines. Four measures of synergy, i.e. ZIP, HSA, Bliss, and Loewe, are seen as notch box plots for the experimental confirmation of 25 chosen predictions in three cell lines. Inter-cluster drug combinations are shown in blue as the positive group and the intra-cluster combinations are shown in red as the negative group.

#### High-throughput drug screening for proposed drug combinations in AML cell lines

In order to further demonstrate the ability of our model to predict specific and robust drug combinations, experimental corroboration was carried out on a subset of 45 drug combinations for 3 AML cell lines, MOLM-16, OCI-AML3, and NOMO-1. The 25 out of 45 drug combinations originated from the top five drugs of the two clusters as the positive group, where higher synergy was predicted by our model. The others were the combinations of the top five drugs within each cluster, which transform into 20 combinations as the negative group. The findings of the experimental validation of 135 drug-drug-cell line triplets are depicted in Fig. 9B, using the ZIP, Bliss, HSA, and Loewe models to assess the degree of synergy. The drug combinations predicted by our model in the positive group were validated as more synergistic when considering positive scores as evidence for a degree of synergy (Fig. 10 and Supplementary Fig. 2). These findings were statistically more significant when using Bliss or HSA measures. Taken together, these results demonstrate the robustness of network-based predictions across various experimental setups and synergy scoring models, and the ability of our network-based model to detect new combinations of treatments.

**Figure 10:**
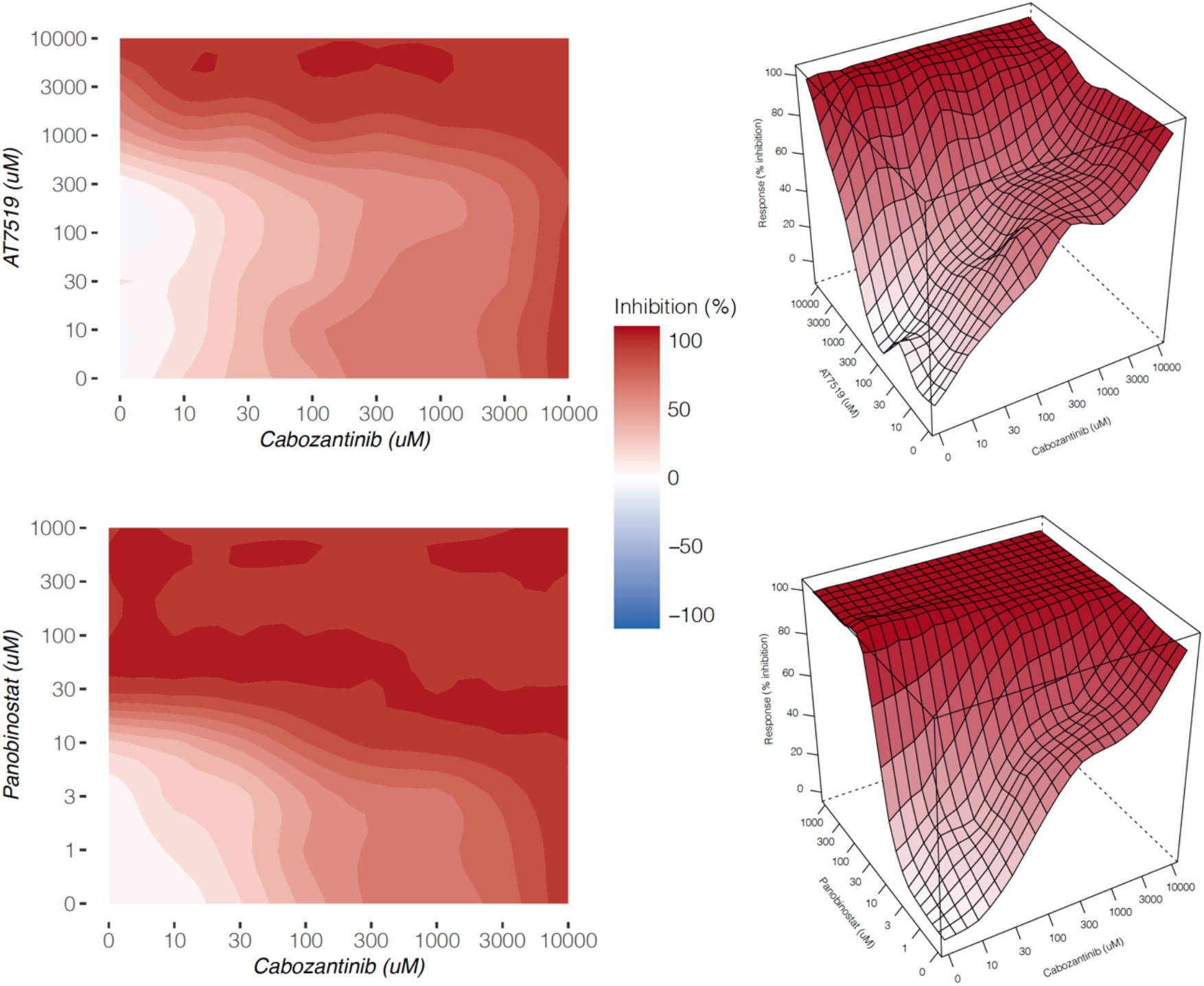
The top synergistic drug combinations identified in positive group. These matrices represent the highest synergistic combination based on four measures of synergy. HSA and LOEWE methods indicated the highest synergy in the cabozanitinb and AT7519 combination. BLISS and ZIP method showed the highest synergy in the cabozanitinb and panobinostat combination. For each combination, the interaction landscapes are shown in both 2D and 3D. The complete interaction landscapes for all the 135 drug combinations can be found in Supplementary Fig. 2.

## Discussion

The availability of single drug response datasets for cancer cell lines has prompted us to develop methods for predicting and selecting the most effective combination therapy. Several AI-based combination prediction approaches have recently been introduced, which combine high-throughput molecular profiling data with drug response data to improve prediction and validation. To reflect the relationships between drug combinations, Narayan *et al.* used dose-response data from pharmacogenomic encyclopedias and represented them as drug atlas (45).

Combining with the pathway/genes ontology data, their approach enables the prediction of combinatorial therapy, i.e., vulnerability when attacked by two drugs that can be related to tumor-driving mutations. They repeated the predicted synergies in several tumors, including glioblastoma, breast cancer, melanoma and leukemia mouse models, highlighting the cancer-independent prediction power of drug combination treatment. Ianevski *et al.* also showed that the bulk viability single-agent screening assays had unexpectedly large predictability for the AML cell subpopulation co-inhibition effects when combined with the scRNA-seq transcriptomic data (18). They developed a machine learning model by combining single-cell RNA sequencing with ex vivo single-agent testing for AML with a different genetic background. They displayed an accurate prediction of synergistic patient-specific combinations while avoiding inhibition of nonmalignant cells. However, our biomarker-independent approach relies only on the phenotypic level of information that is drug-response data. Although, our predictions were consistent with the molecular profiling and biochemical annotations when it came to assessing the intra-cluster homogeneity of drugs, patients and cell lines

Moreover, a training machine learning model for predicting drug combination response, called comboFM, was recently introduced using drug combination screening data as a training dataset (46). comboFM uses a factorization machine to model the cell context-specific drug interactions through higher-order tensors. Julkunen *et al.* demonstrated that comboFM enables leveraging information from previous experiments performed on similar drugs and cells as training data when predicting responses of new combinations in so far untested cells (testing data). They displayed high predictive performance and robust applicability of comboFM in various prediction scenarios using experimental validation of a set of previously untested drug. However, we expounded that predication accuracy of inter-cluster design strategy of drug combinations based on multipartite networks can be achieved independently of the high-quality training dataset.

Strictly speaking, in the present study, we revisited the analysis of nominal variables, namely drug name and sample id, in drug screening results for data mining using graph theory, which we termed the nominal data mining approach. We first considered data quality control, such as outlier detection, outlier treatment, and biological and technical replicates. Because of the discrete explanatory independent variable (i.e., drug doses) (47), we assumed that regression-based measurements might even be discarded; hence, we demonstrated that median values can represent an appropriate weight score to compare drug functionality for network reconstruction. These values were used to quantify and weight the bipartite network, which reflect the interaction strength of drugs and biological samples. Then, two similarity networks were provided by weighted network projection to detect the topological structure of networks, i.e., network communities. We showed that network communities represent a rationale starting point to propose a combinational drug regimen. Our computational and experimental validation steps amplified the logic of our proposed platform. As a result, while training datasets were not required in this method to predict drug combination, drug response data alone was sufficient for prediction without integrating with prior knowledge of biochemical profiling.

Noting that the occurrence of synergistic toxicities, which can arise from additive toxicities when targets are shared by the combined drugs, is a major barrier to applying combination therapy in the clinic (48). If drug screening data on healthy cells is available, we suggest that a similar strategy for predicting toxicity without losing efficacy is also essential before the future translational experiment. Ianevski *et al.* previously illustrated the importance of a desired synergy-efficacy-toxicity balance for predicting patient-customized drug combinations (18). Hence, a drug-response data on healthy cells is demanded to complement synergistic interactions of drug combinations with toxicity predictions Indeed, where drug synergy and toxicity data are optimally matched for combinatorial therapy, stronger and longer-lasting outcomes of drug combinations can be predicted.

Furthermore, prospective works would necessitate the provision of further patient-derived experimental validations. Despite the fact that our prediction is solely dependent on the drug sensitivity dataset, our suggested combinations address the common mutational assigned etiology of AML. Remarkably, this combination was proposed purely on the grounds of the phenotypic response of cancer cells or patient samples to the drugs, with no previous knowledge of the disease’s genetic origin.

## Supplementary figures

**Supplementary figure 1:** Drug and patient nodes in the projected networks of the Beat AML dataset. The nodes are ordered based on the strength of each node, which is the sum of the edge weights of the adjacent edges for each node.

**Supplementary figure 2:** The complete interaction landscapes for all the 135 drug combinations.

## Notes

### Competing Interest Statement

The authors have declared no competing interest.

### Summary of Updates

The discussion added to this version. In this version, we represented more results and edited the whole manuscript.

