## Supplementary figures and images for "Facilitating the design of combination therapy in cancer using multipartite network models: Emphasis on acute myeloid leukemia"

### H8140-C1-301_1_Doramapimod_Panobinostat_MOLM-16.pdf

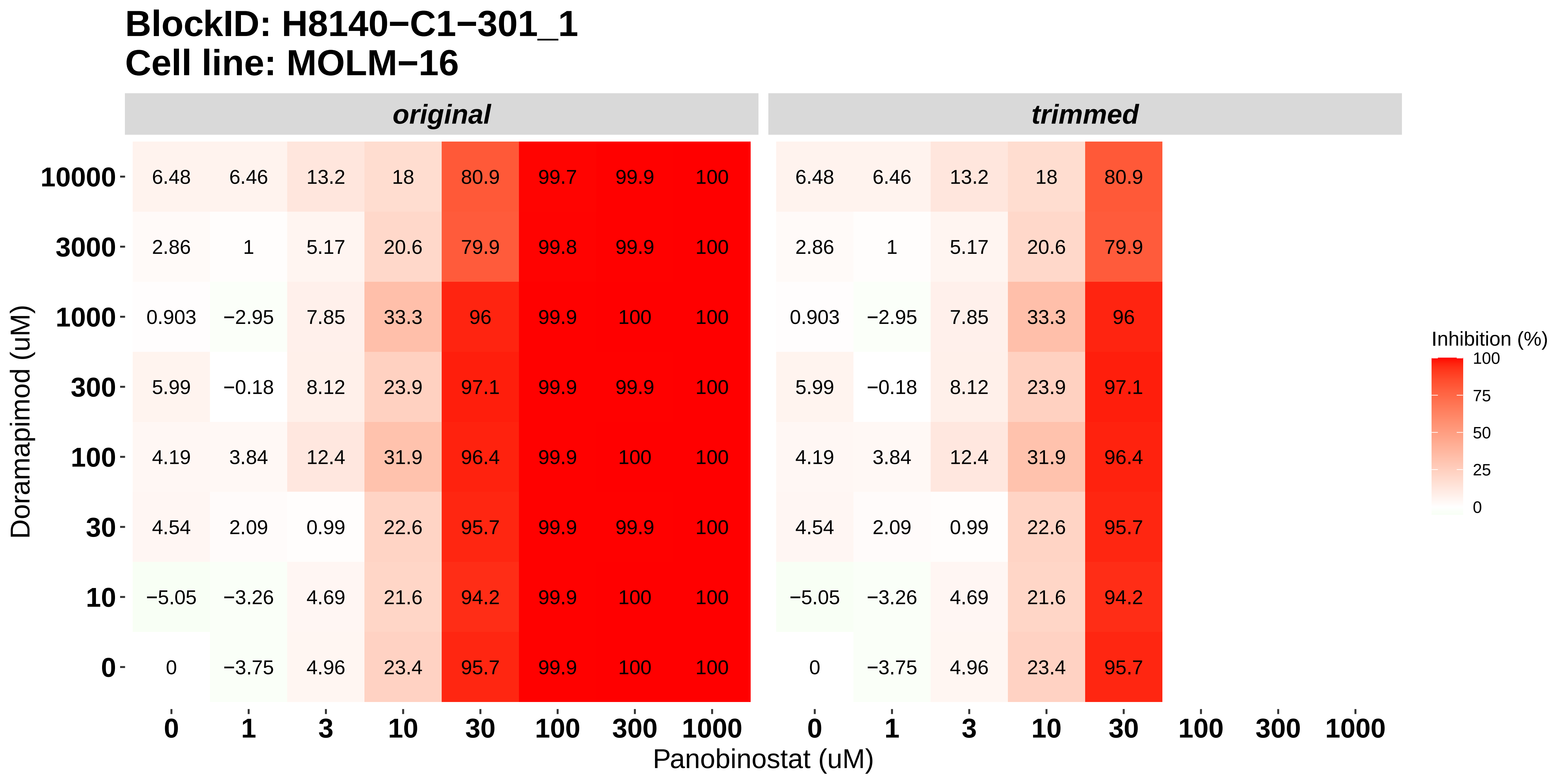

### H8140-C1-301_2_KI20227_Panobinostat_MOLM-16.pdf

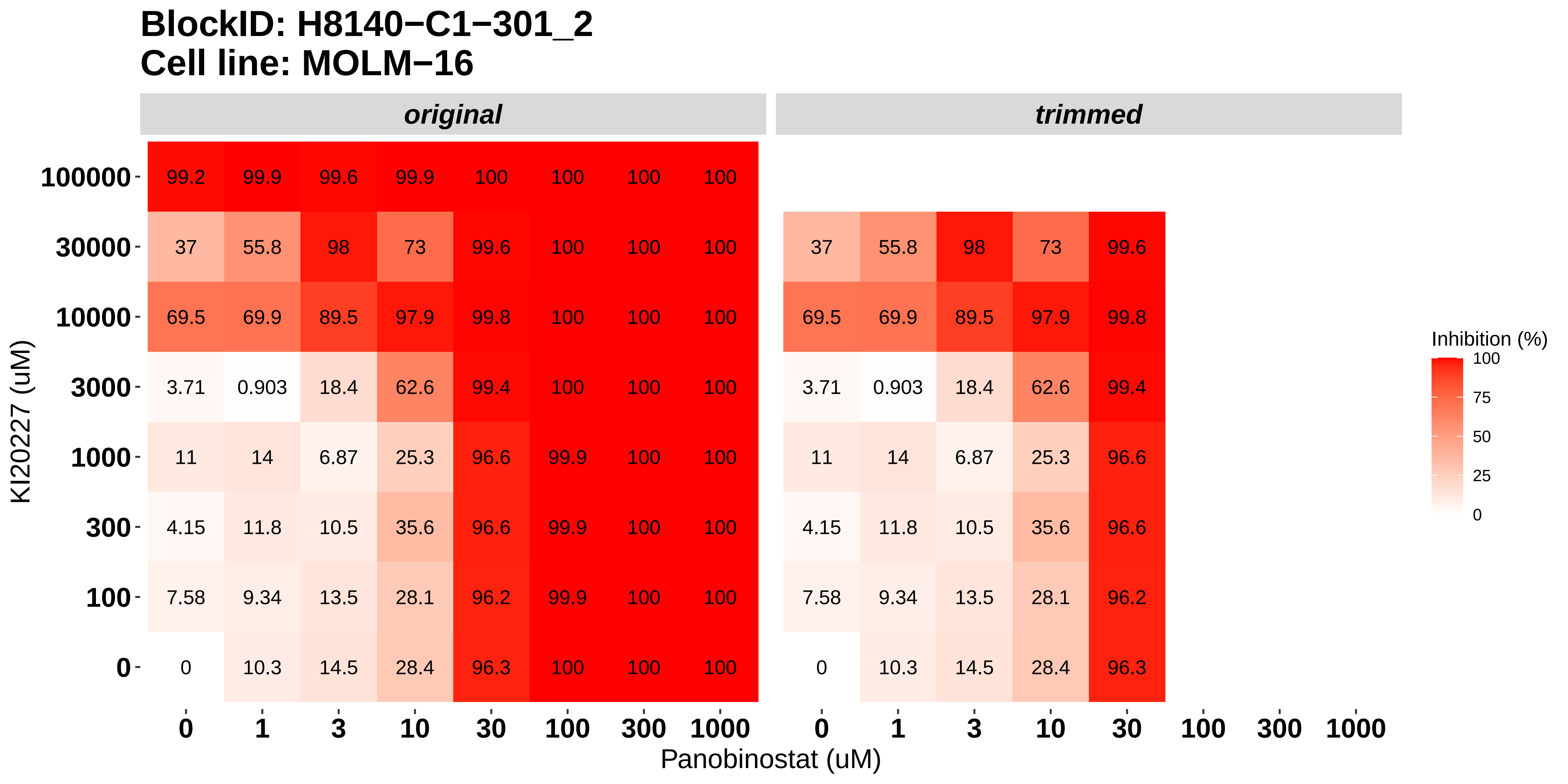

### H8140-C1-301_3_Cabozantinib_Panobinostat_MOLM-16.pdf

#### BlockID: H8140−C1−301_3
#### Cell line: MOLM−16

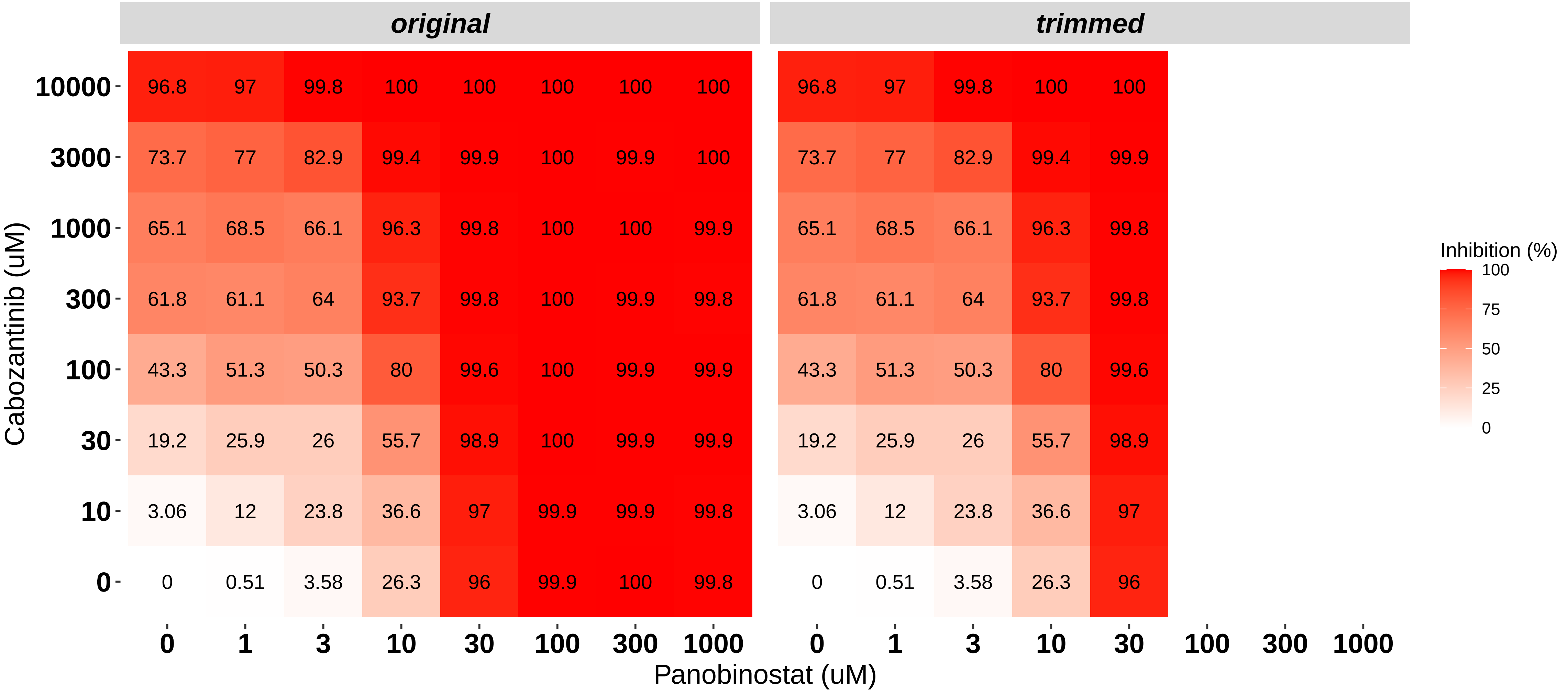

### H8140-C1-301_4_Dovitinib_AT7519_MOLM-16.pdf

##### BlockID: H8140−C1−301_4
##### Cell line: MOLM−16

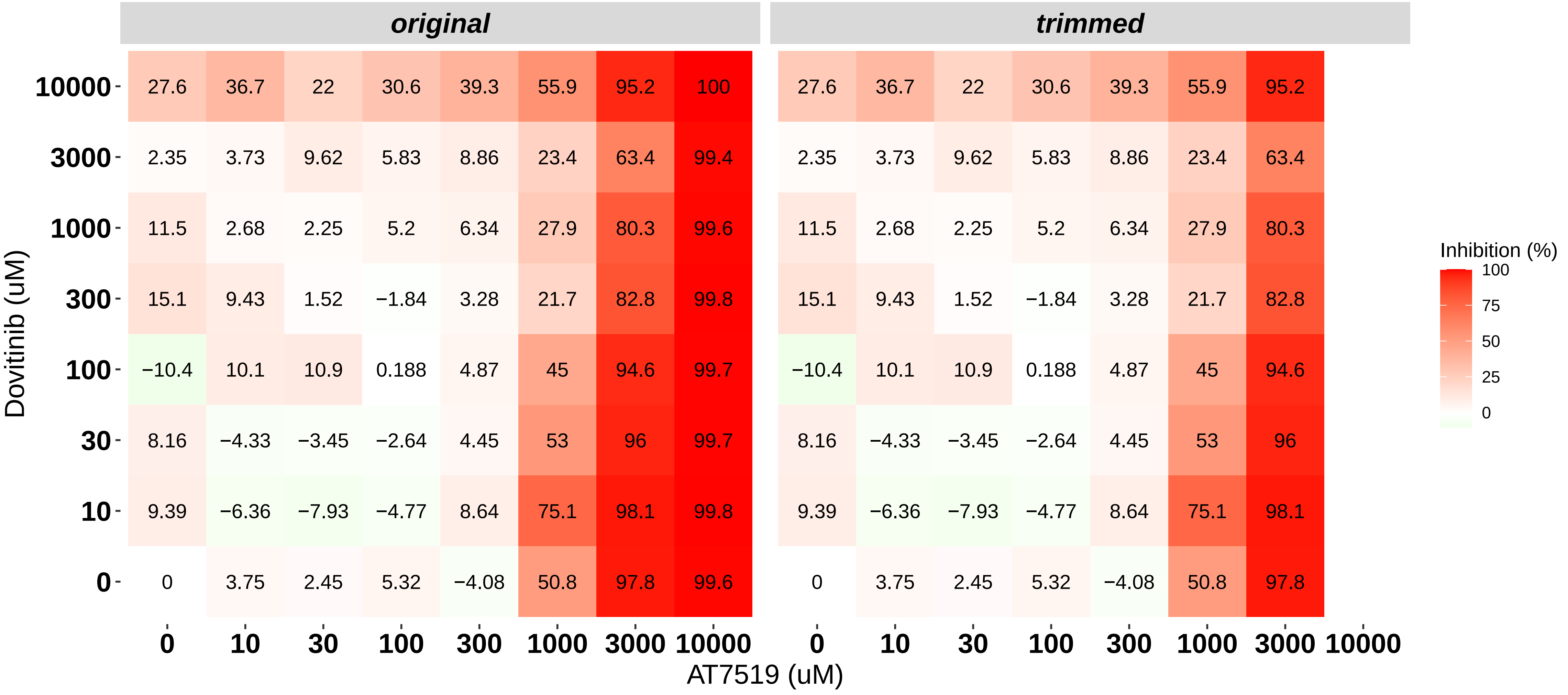

### H8140-C1-301_5_Nintedanib_AT7519_MOLM-16.pdf

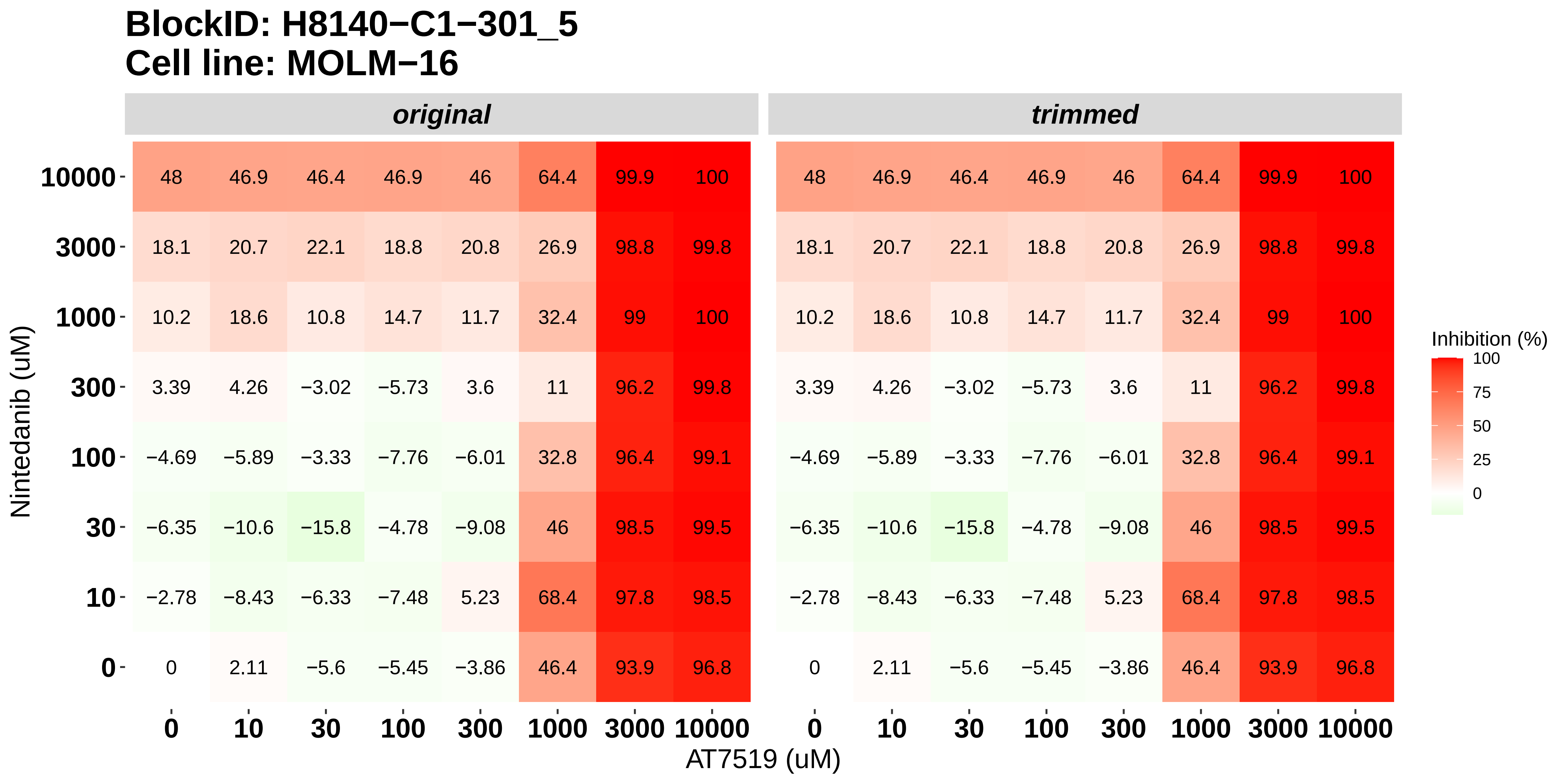

### H8140-C1-301_6_Doramapimod_AT7519_MOLM-16.pdf

#### BlockID: H8140−C1−301_6
#### Cell line: MOLM−16

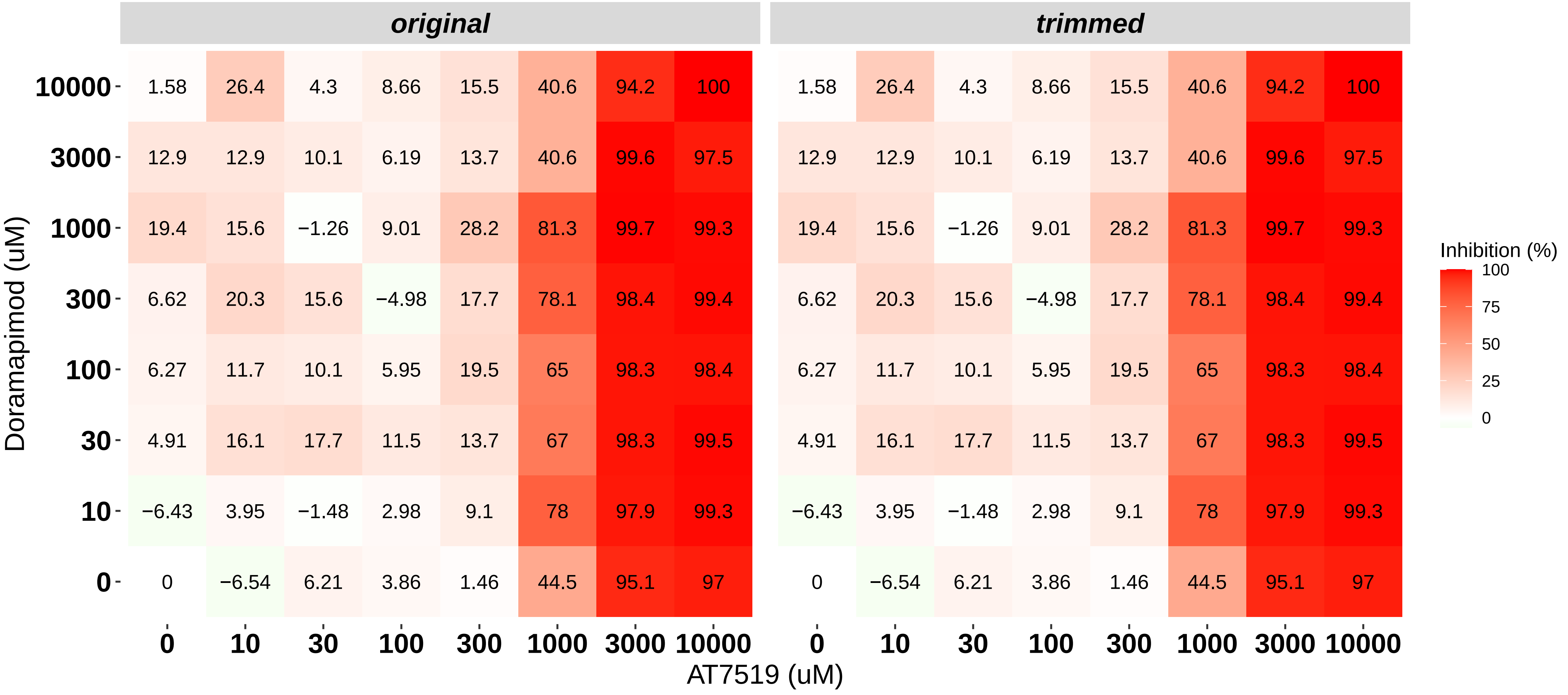

### H8140-C1-302_1_Doramapimod_Panobinostat_NOMO-1.pdf

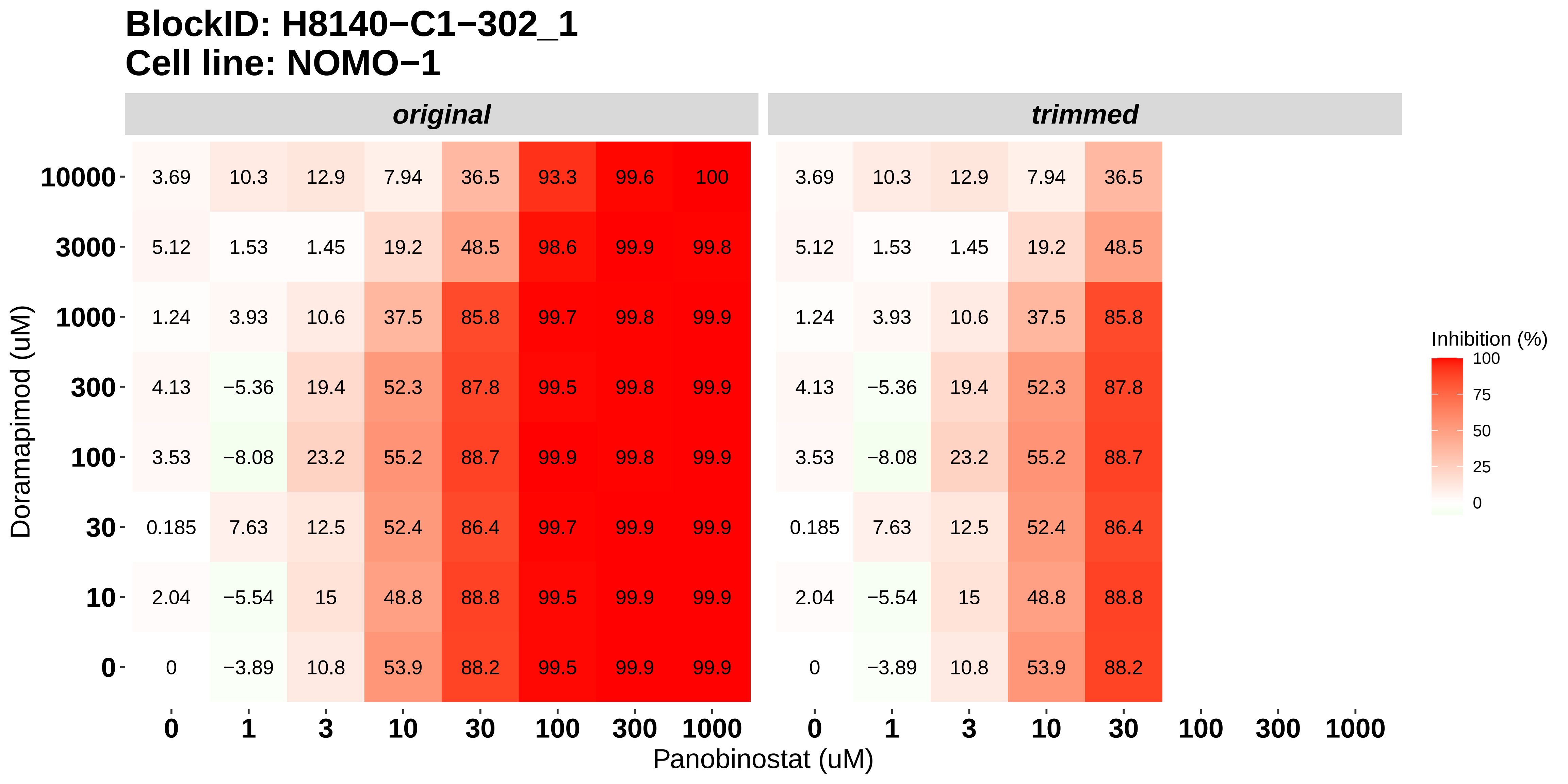

### H8140-C1-302_2_KI20227_Panobinostat_NOMO-1.pdf

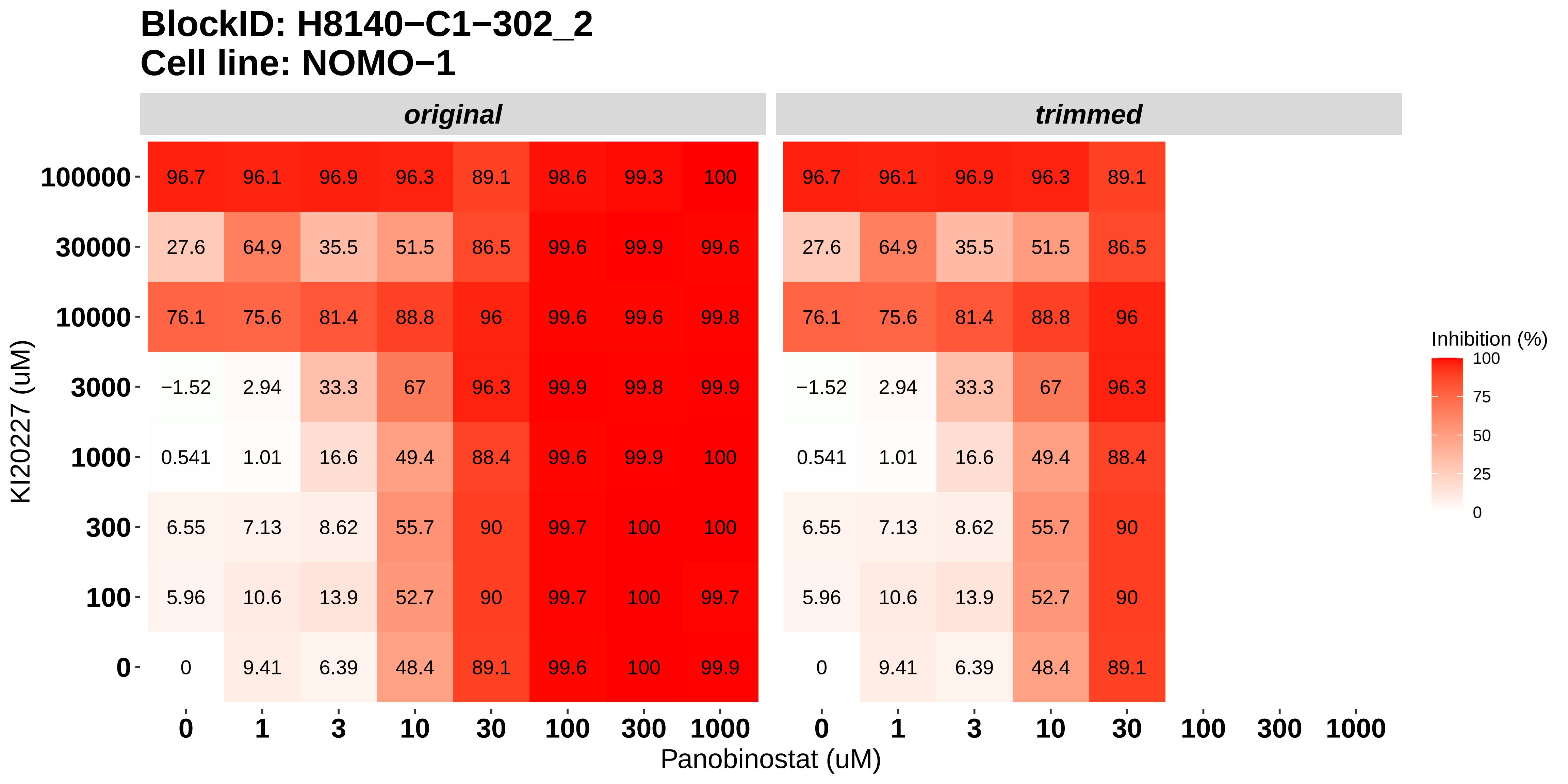

### H8140-C1-302_3_Cabozantinib_Panobinostat_NOMO-1.pdf

#### BlockID: H8140−C1−302_3
#### Cell line: NOMO−1

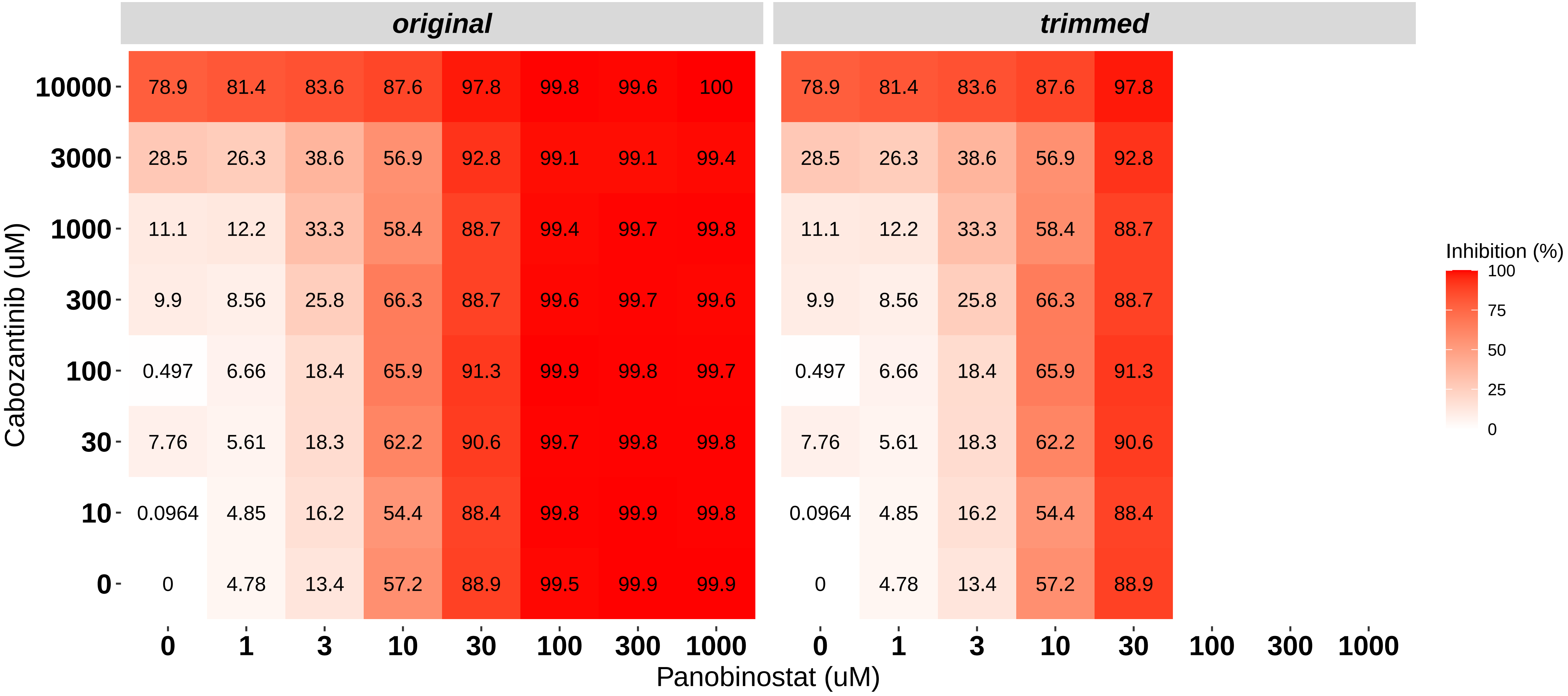

### H8140-C1-302_4_Dovitinib_AT7519_NOMO-1.pdf

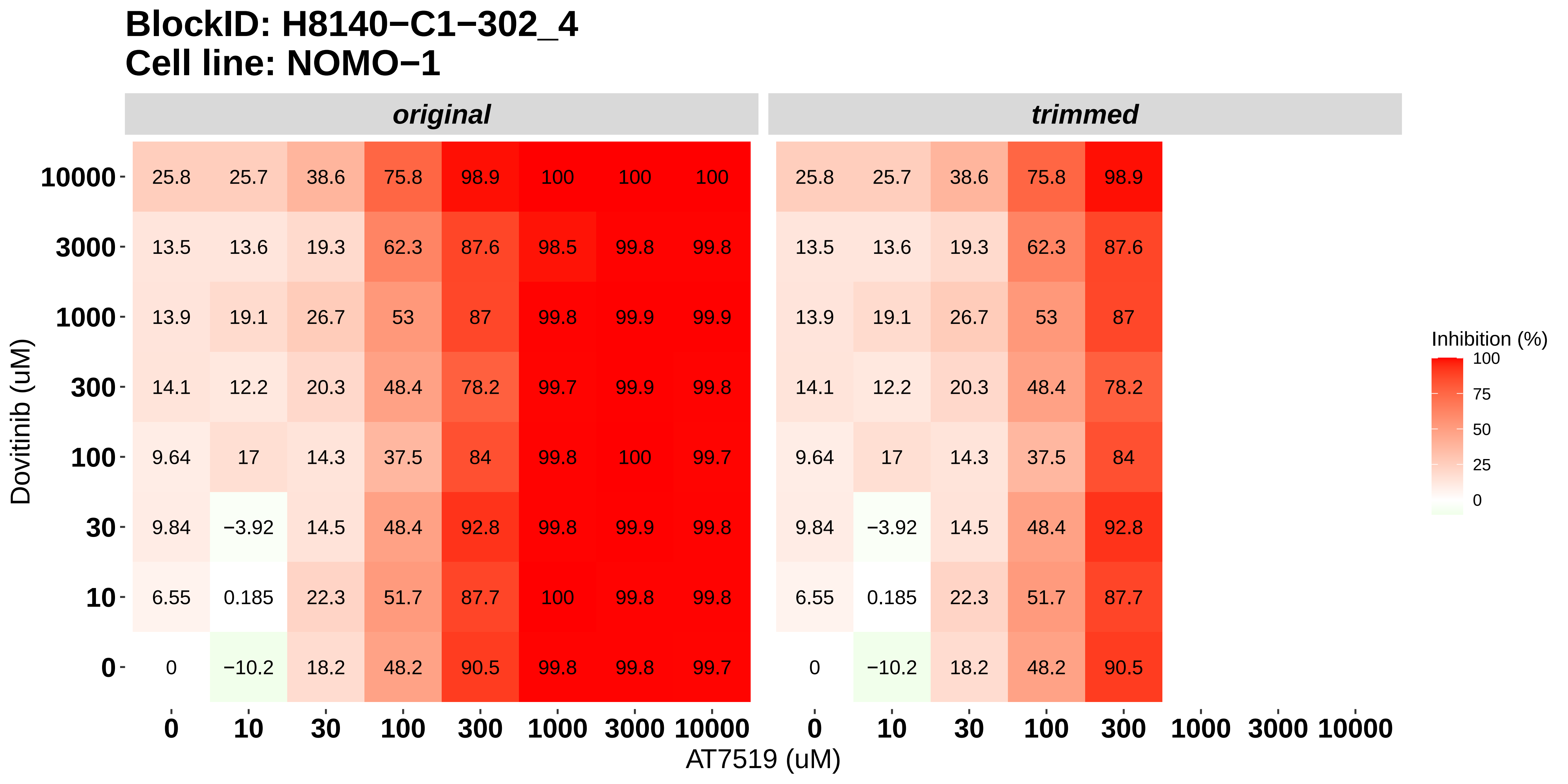

### H8140-C1-302_5_Nintedanib_AT7519_NOMO-1.pdf

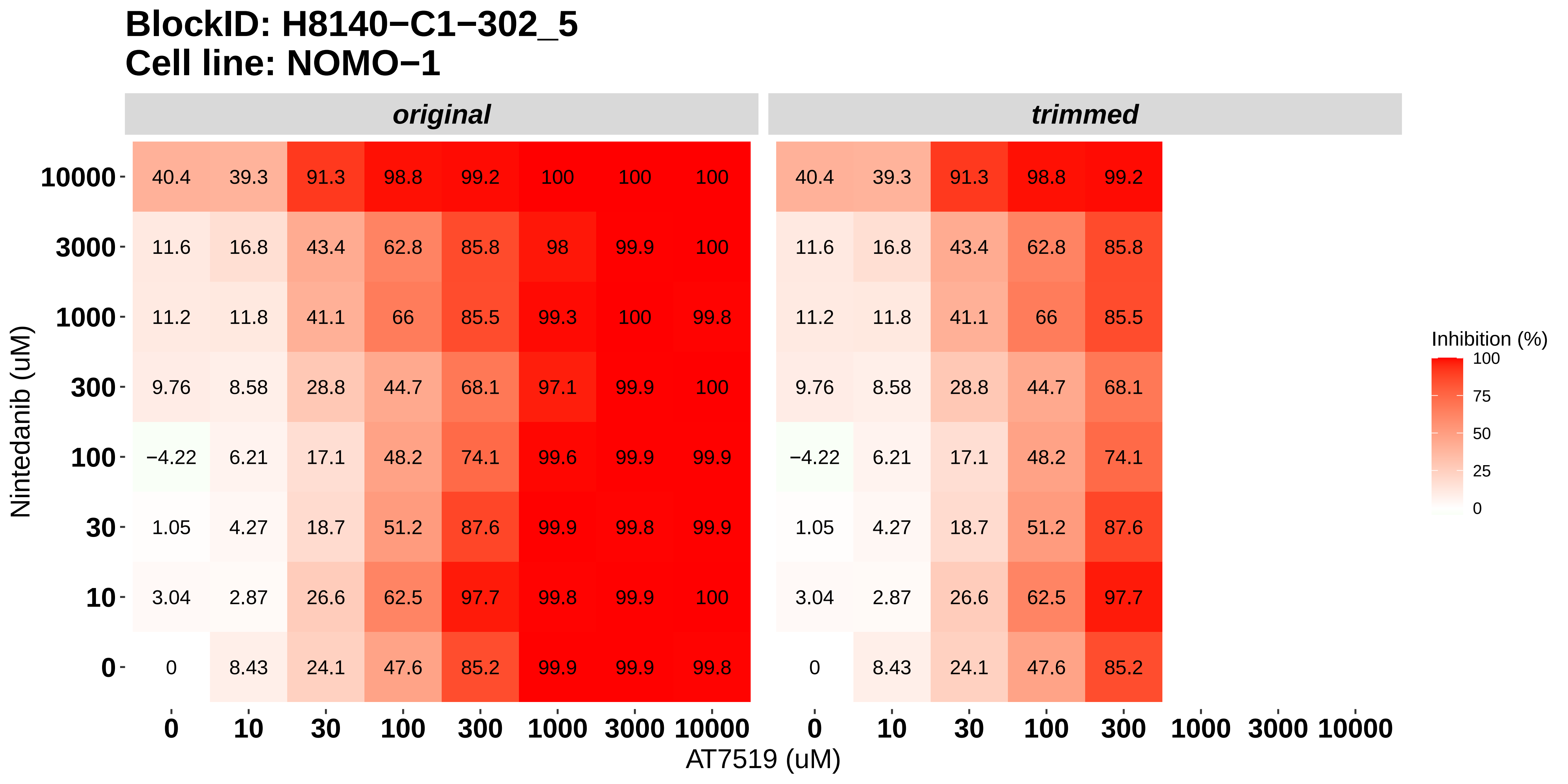

### H8140-C1-302_6_Doramapimod_AT7519_NOMO-1.pdf

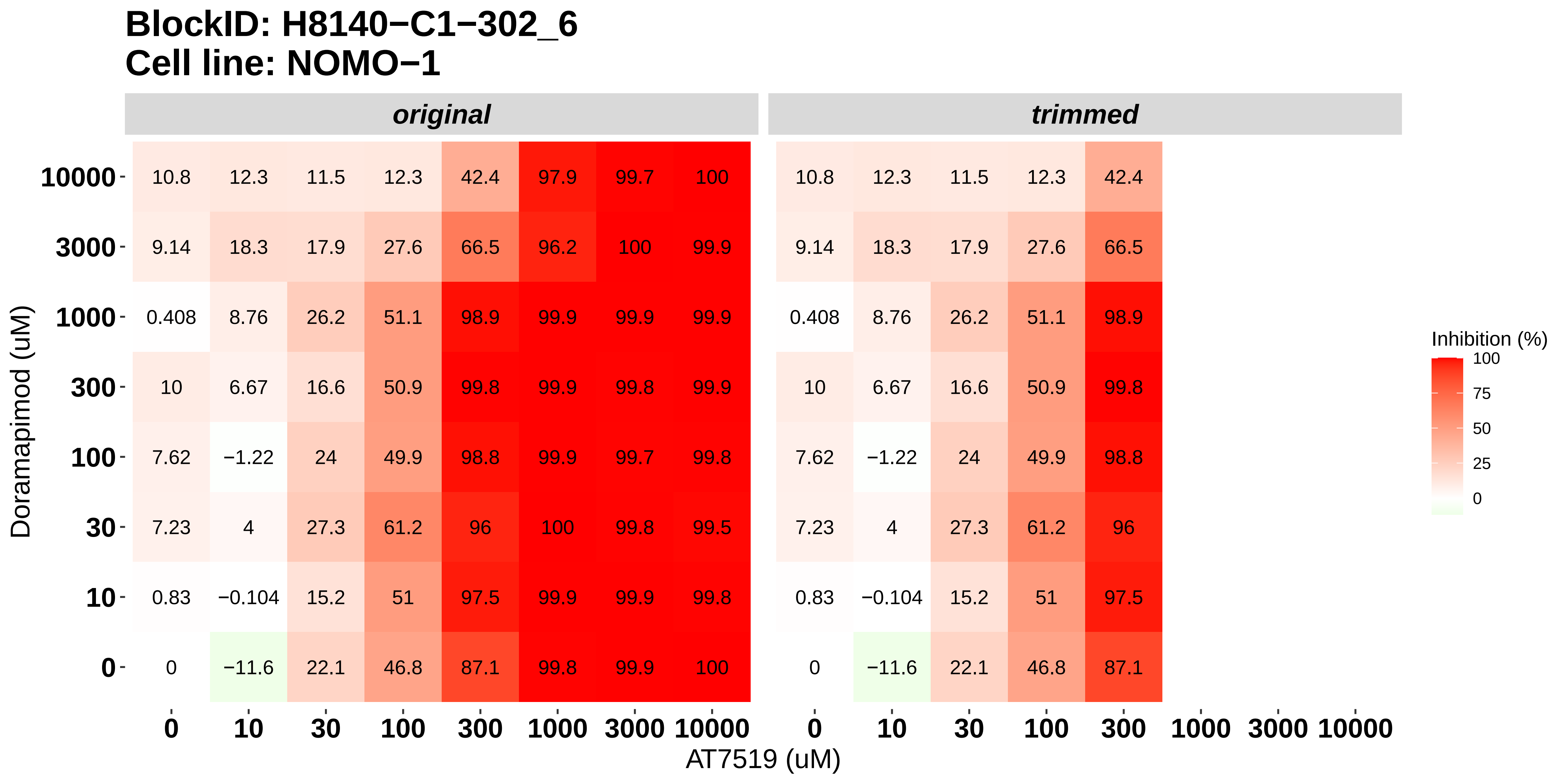

### H8140-C1-303_1_Doramapimod_Panobinostat_OCI-AML3.pdf

#### BlockID: H8140−C1−303_1
#### Cell line: OCI−AML3

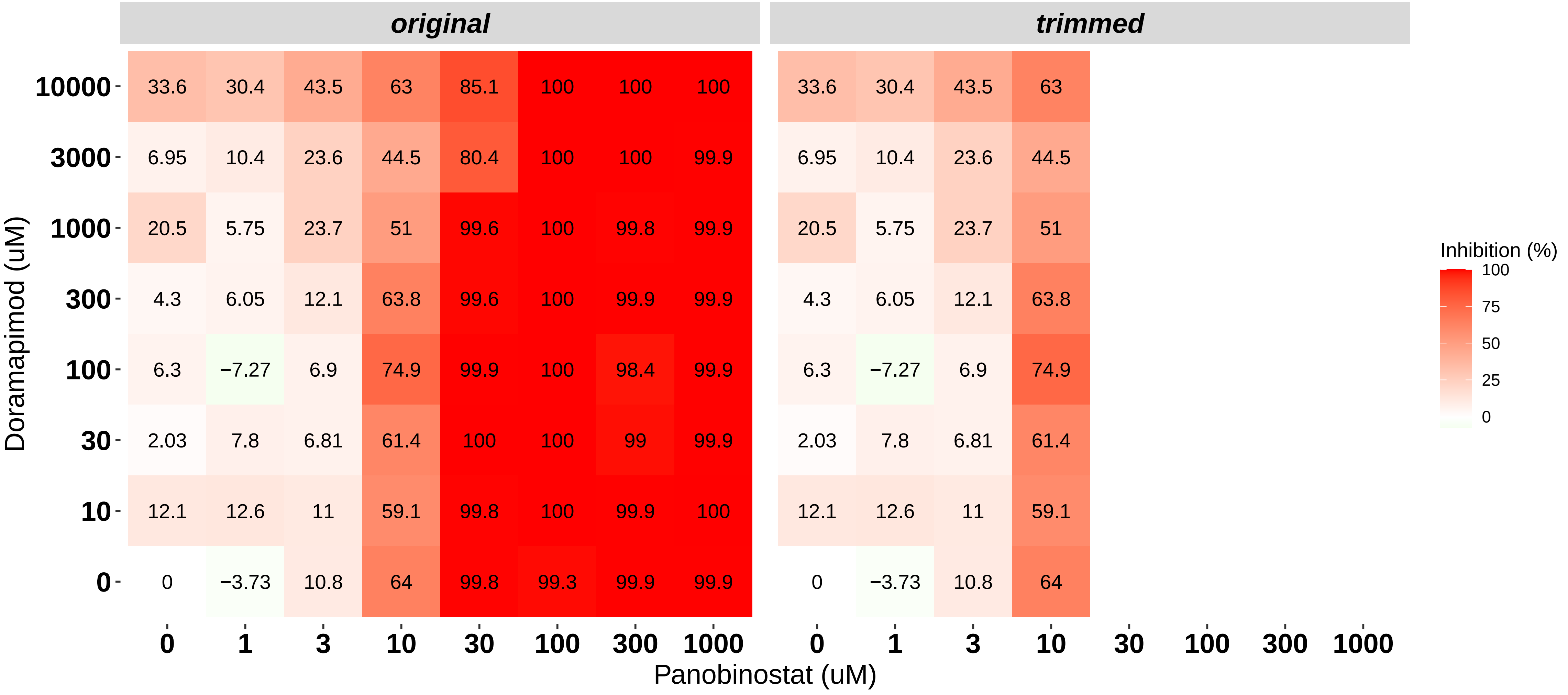

### H8140-C1-303_2_KI20227_Panobinostat_OCI-AML3.pdf

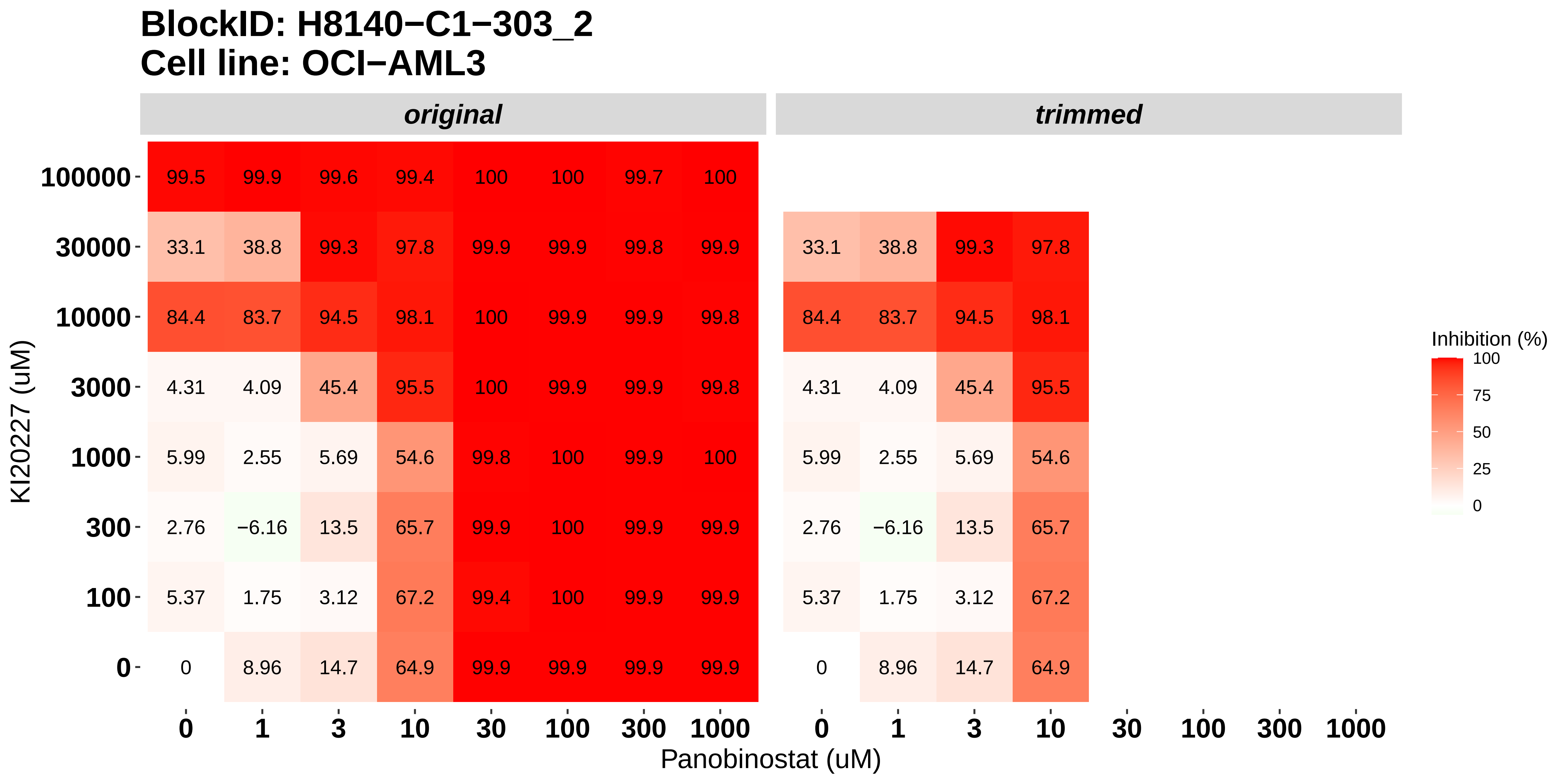

### H8140-C1-303_3_Cabozantinib_Panobinostat_OCI-AML3.pdf

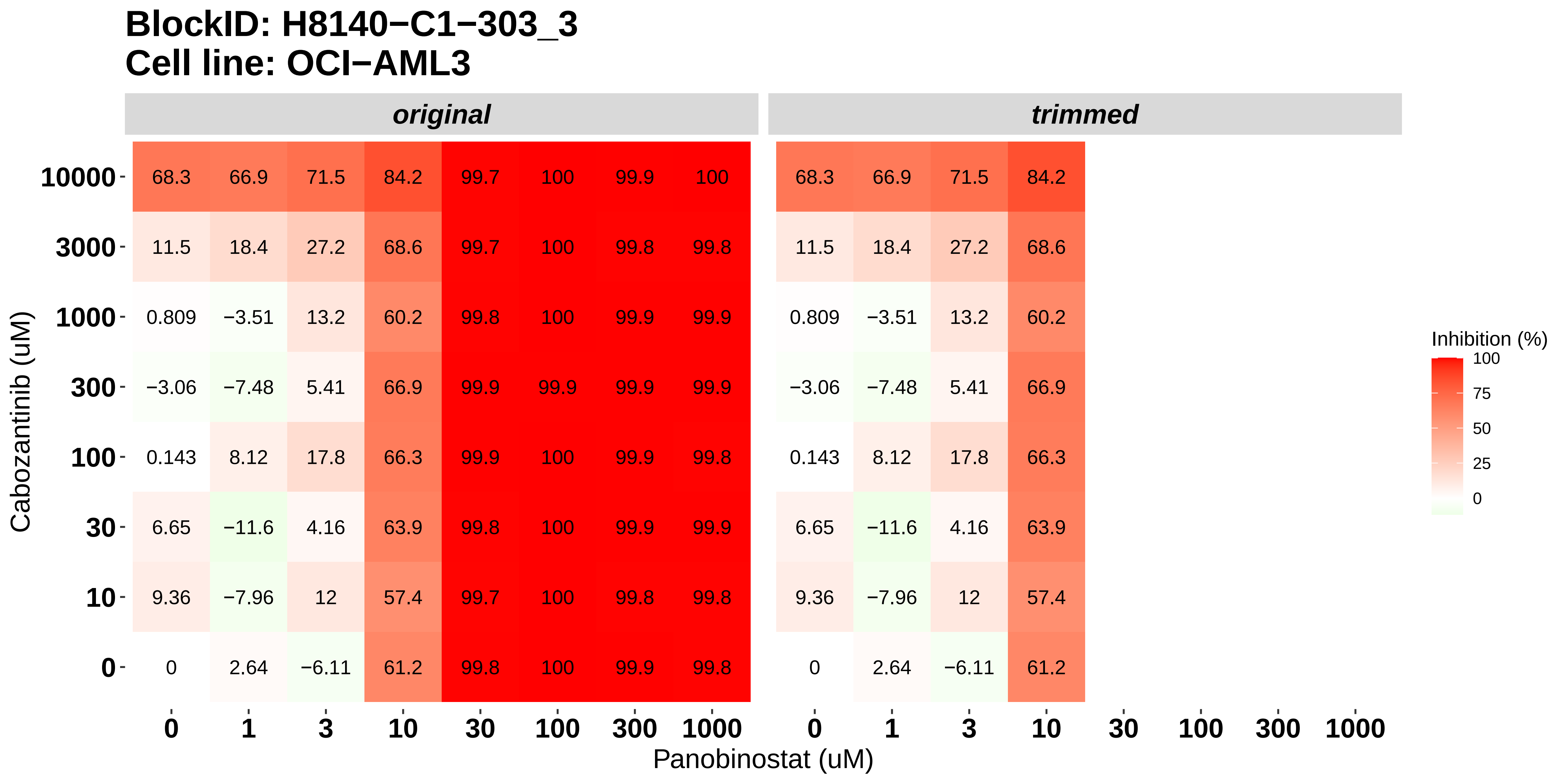

### H8140-C1-303_4_Dovitinib_AT7519_OCI-AML3.pdf

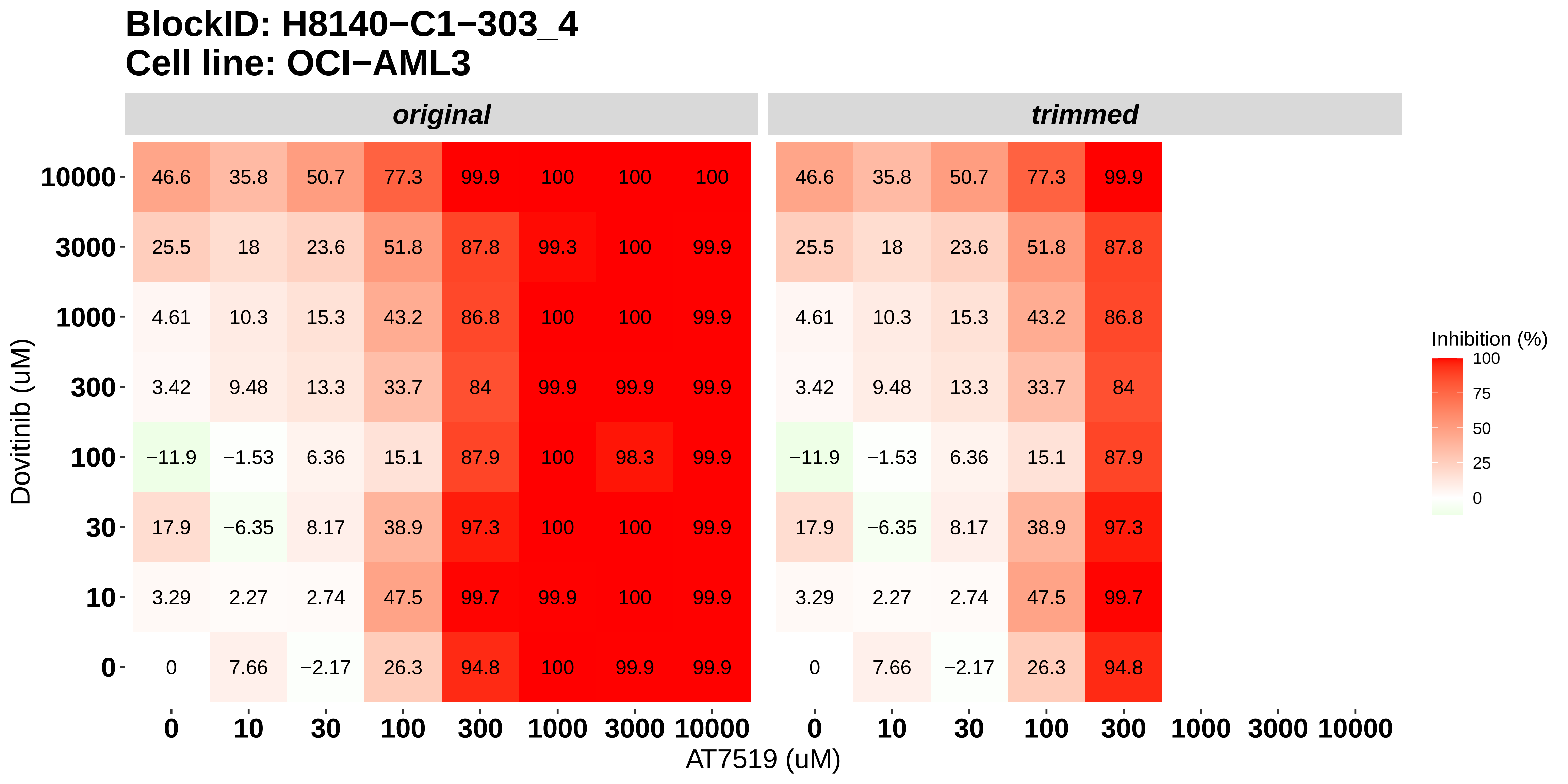

### H8140-C1-303_5_Nintedanib_AT7519_OCI-AML3.pdf

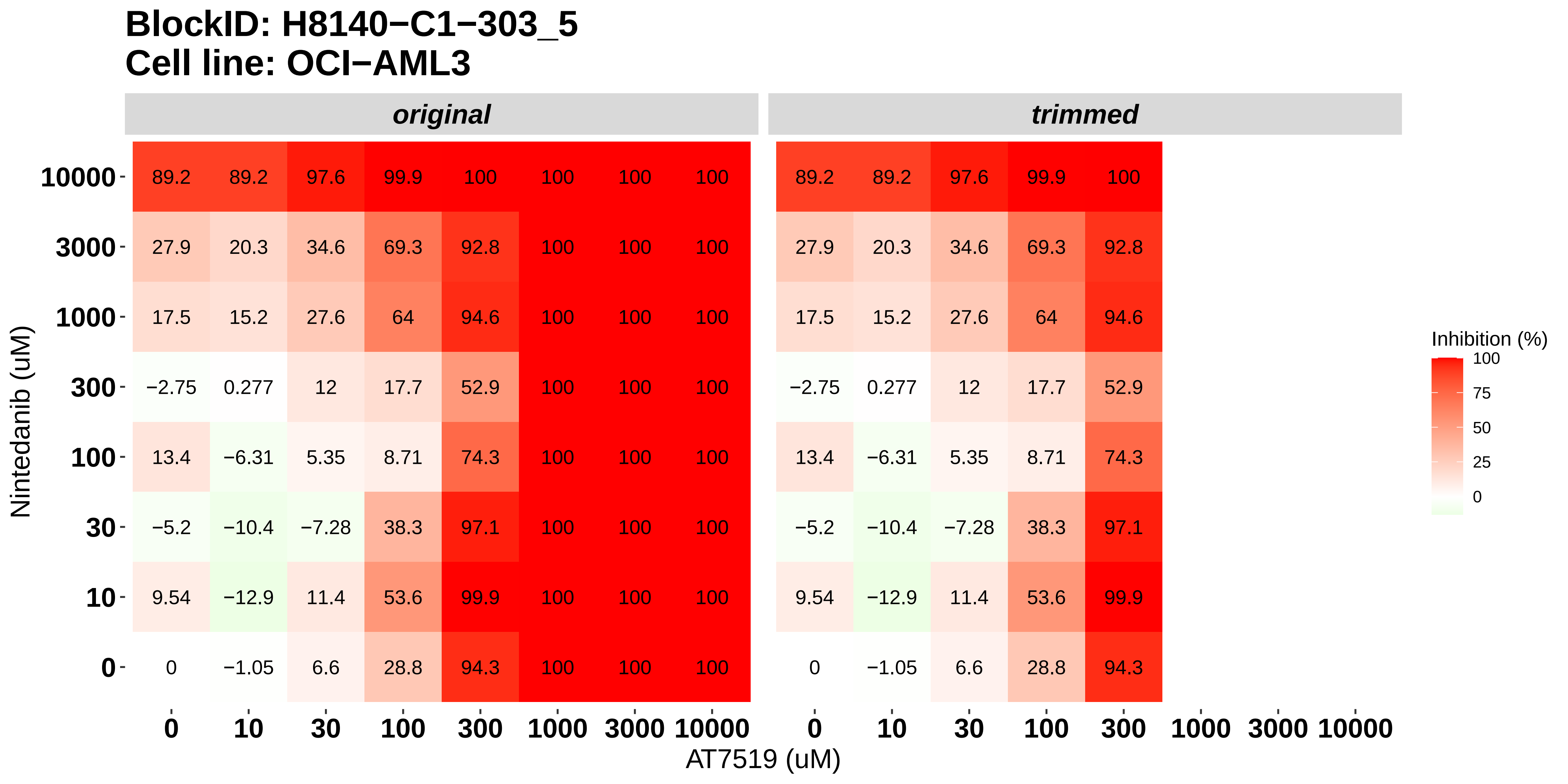

### H8140-C1-303_6_Doramapimod_AT7519_OCI-AML3.pdf

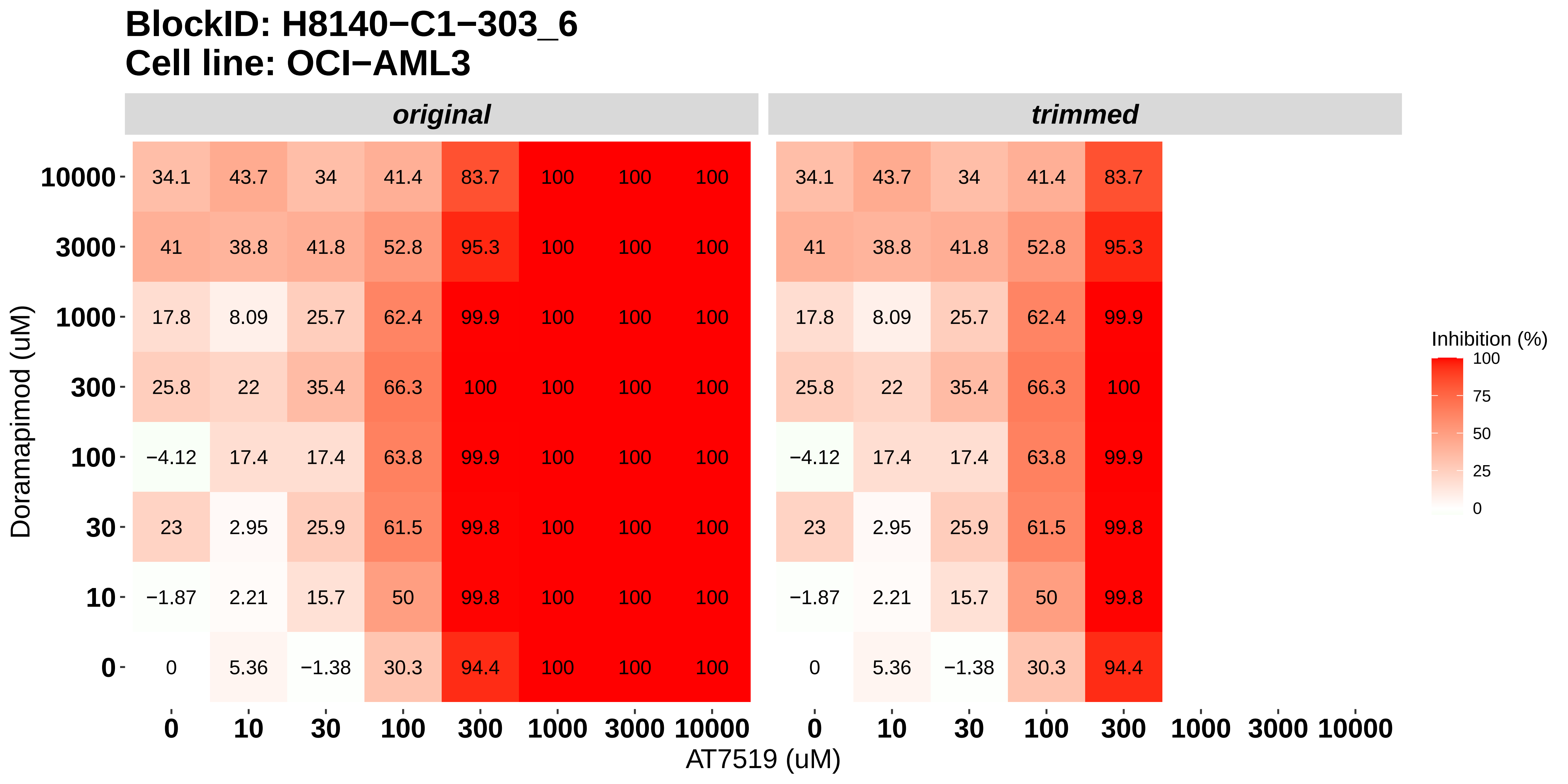

### H8140-C1-401_1_KI20227_AT7519_MOLM-16.pdf

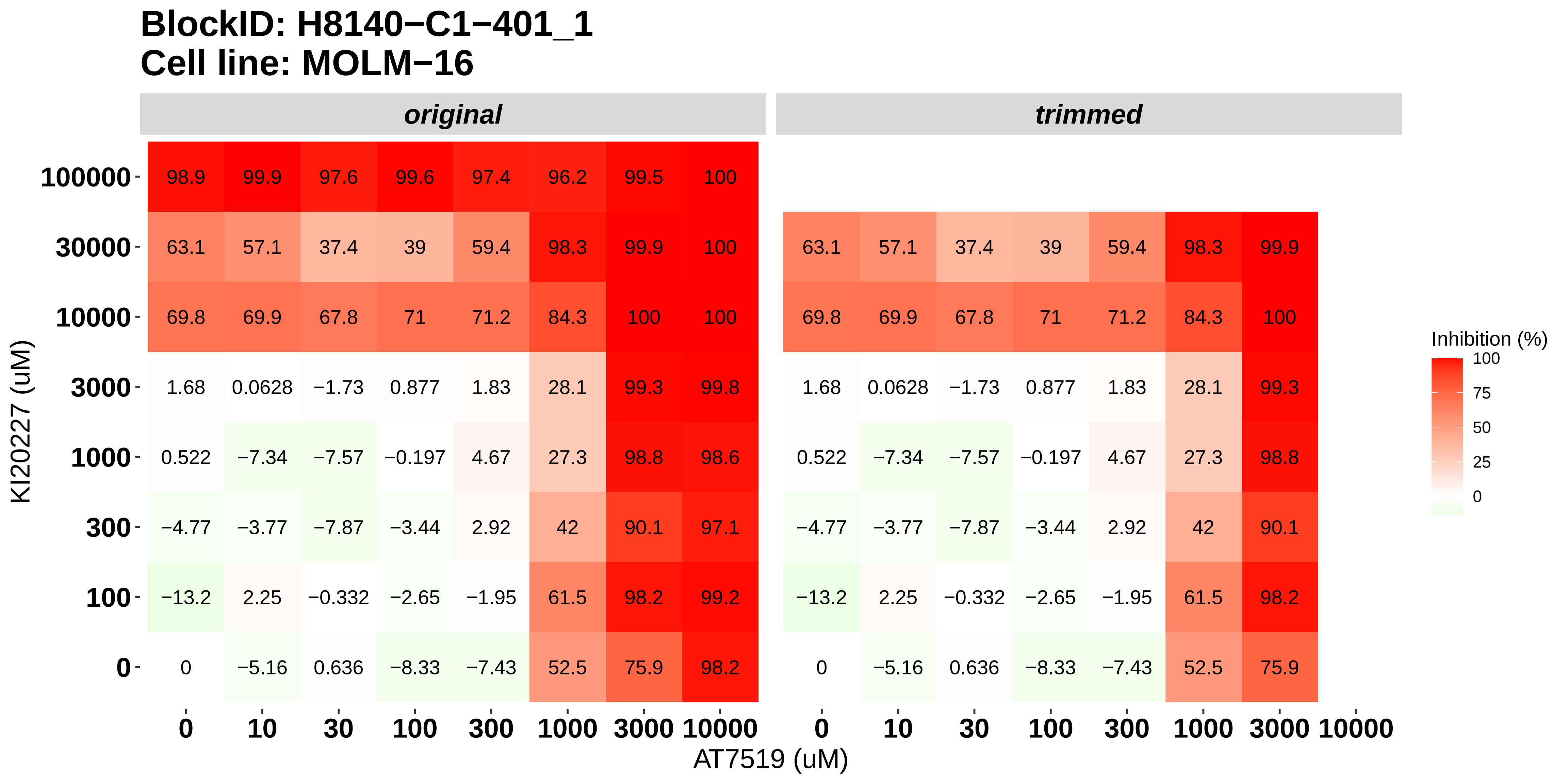

### H8140-C1-401_2_Cabozantinib_AT7519_MOLM-16.pdf

**BlockID: H8140−C1−401_2**
**Cell line: MOLM−16**

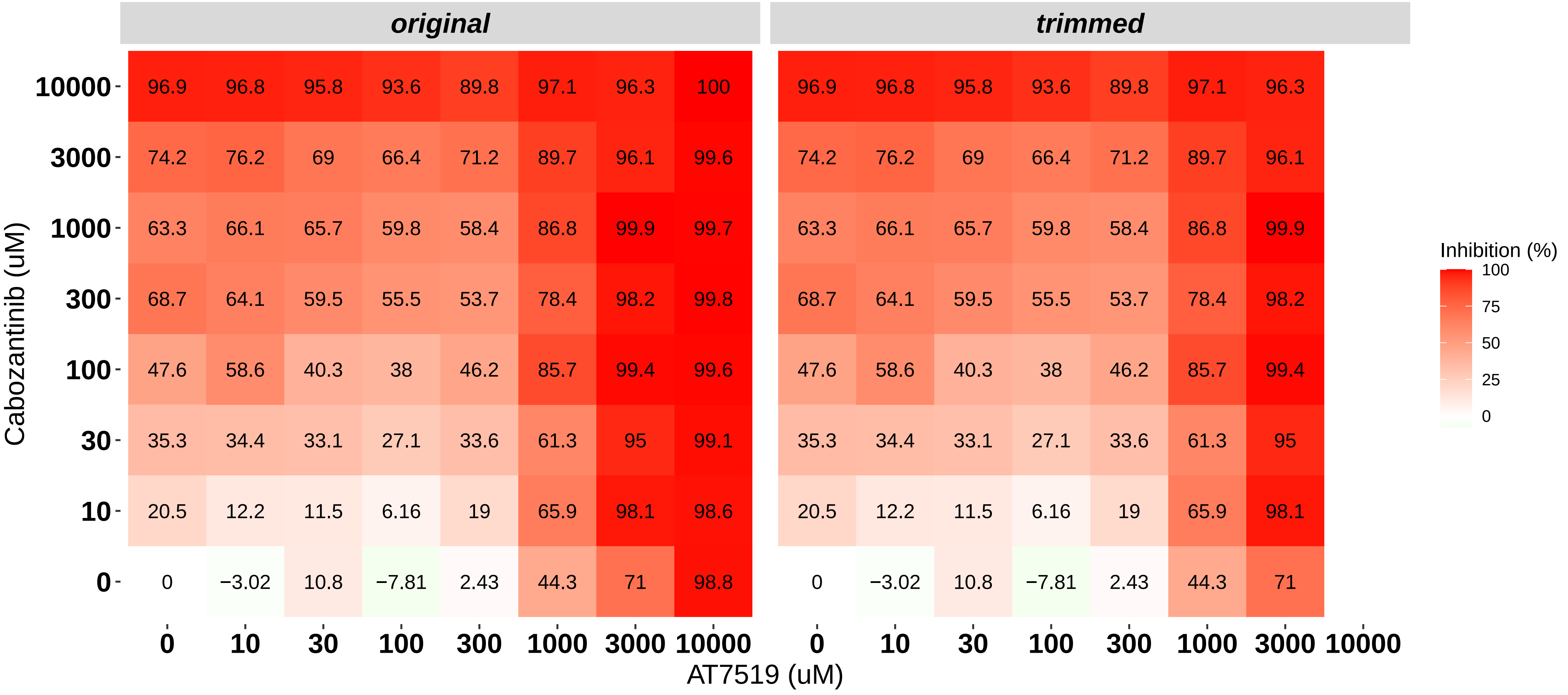

### H8140-C1-401_3_Dovitinib_Bortezomib_MOLM-16.pdf

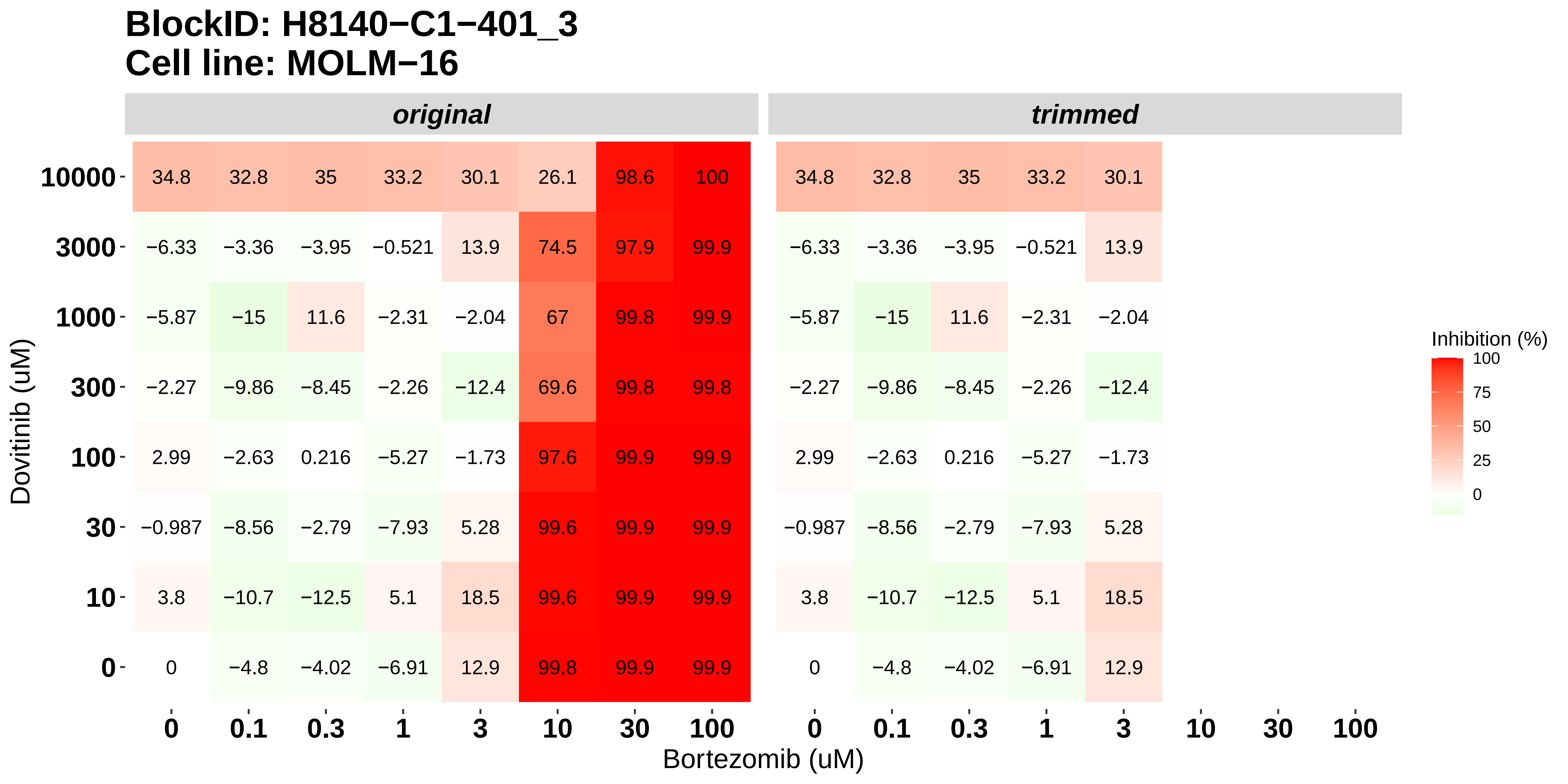

### H8140-C1-401_4_Nintedanib_Bortezomib_MOLM-16.pdf

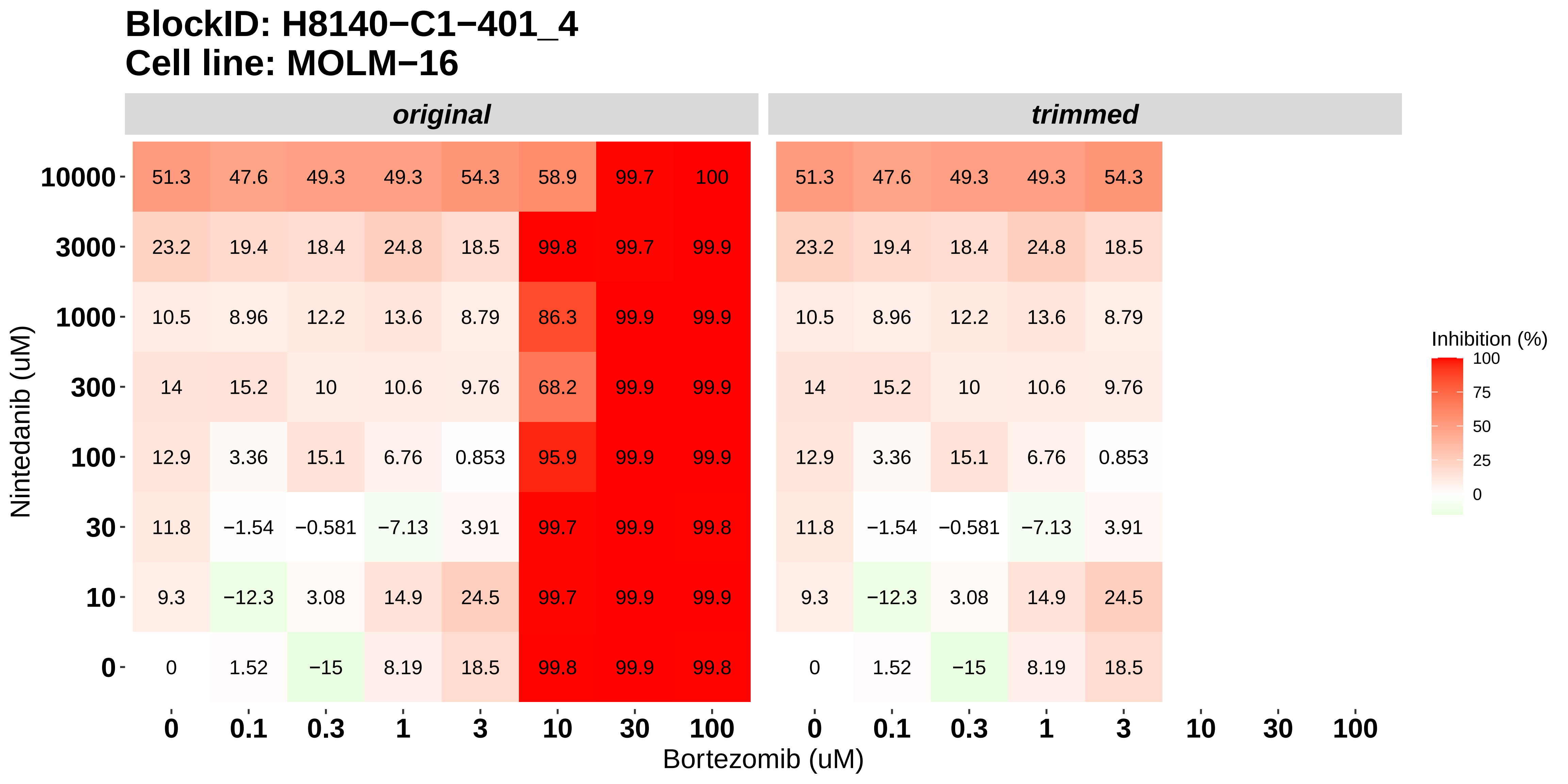

### H8140-C1-401_5_Doramapimod_Bortezomib_MOLM-16.pdf

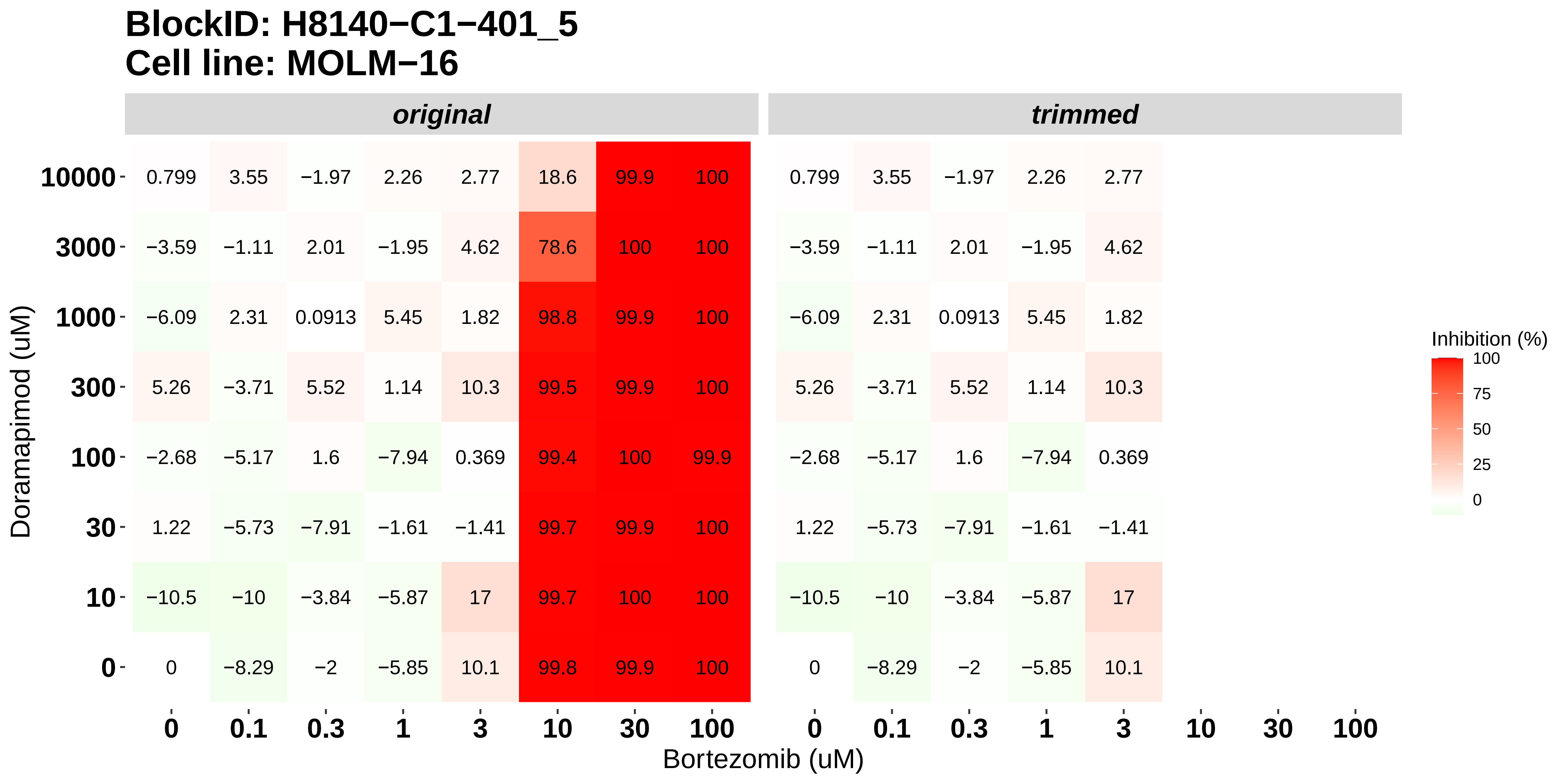

### H8140-C1-401_6_KI20227_Bortezomib_MOLM-16.pdf

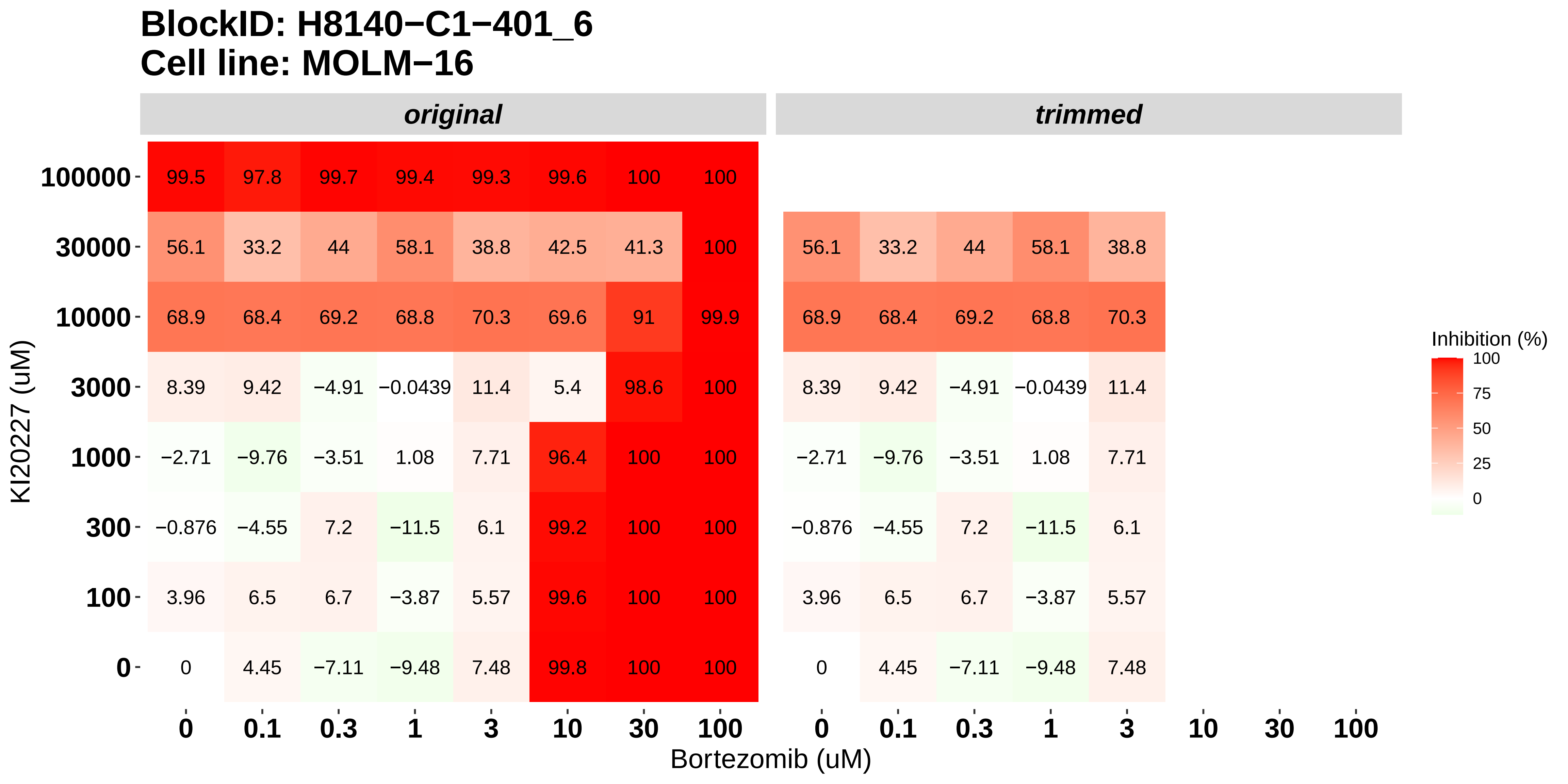

### H8140-C1-402_1_KI20227_AT7519_NOMO-1.pdf

#### BlockID: H8140−C1−402_1
#### Cell line: NOMO−1

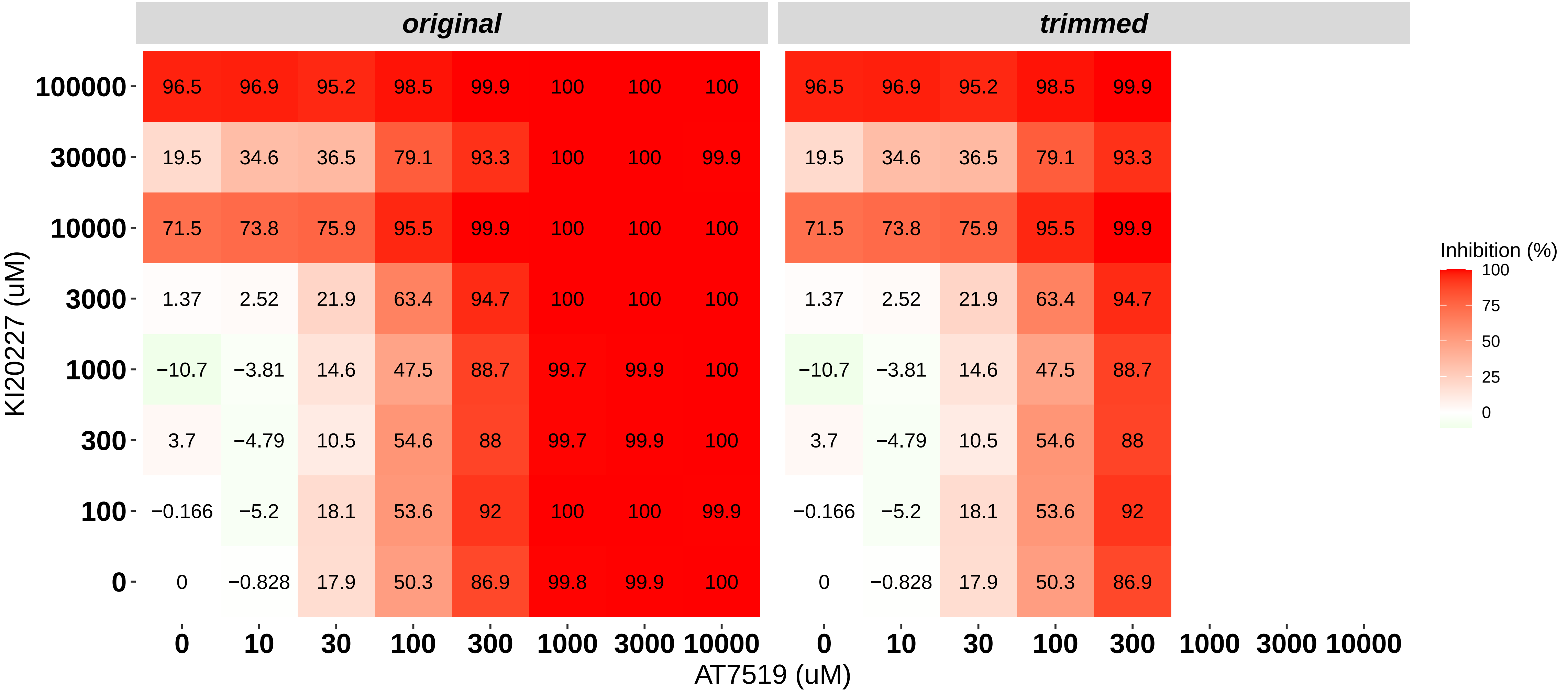

### H8140-C1-402_2_Cabozantinib_AT7519_NOMO-1.pdf

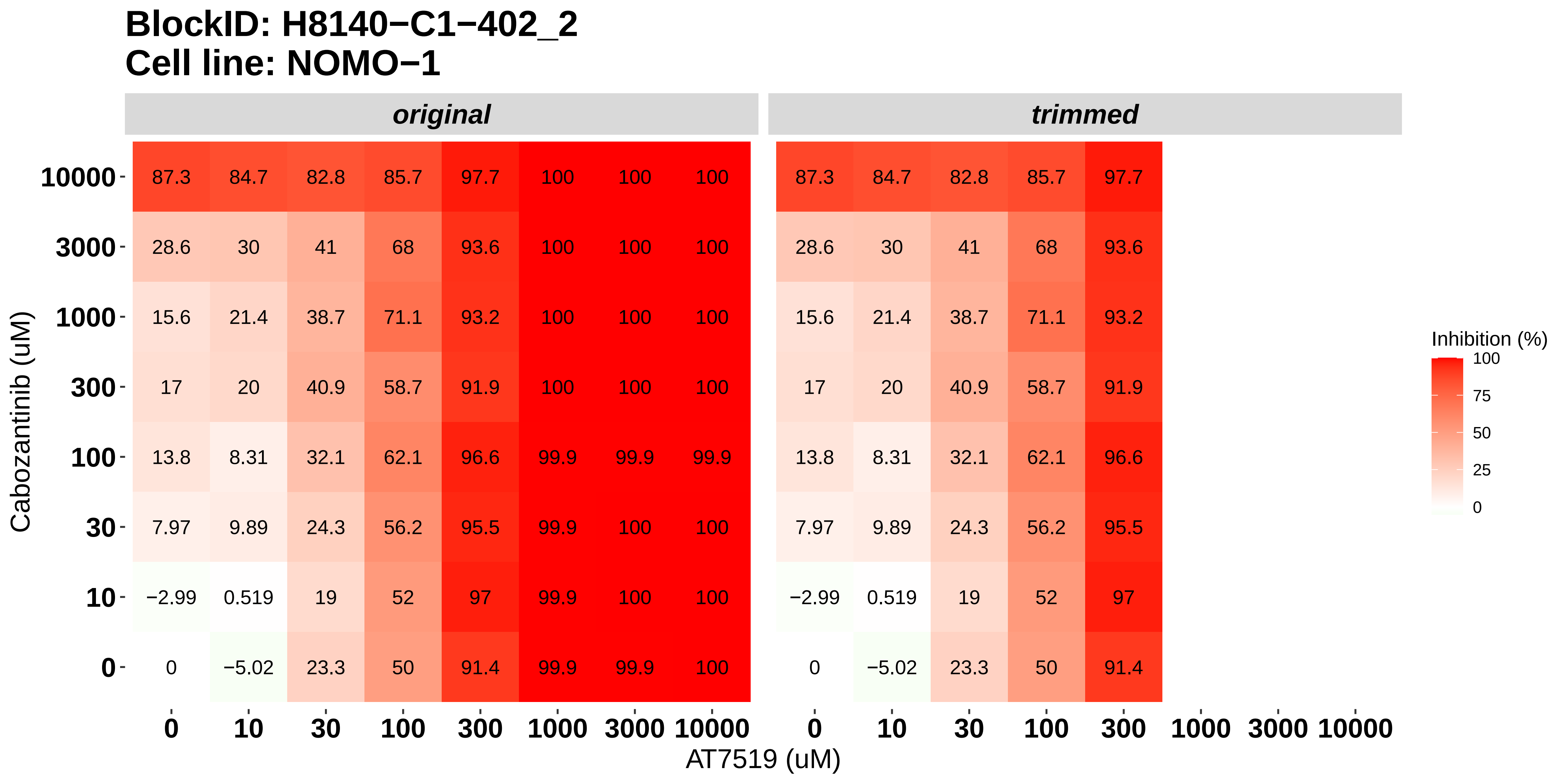

### H8140-C1-402_3_Dovitinib_Bortezomib_NOMO-1.pdf

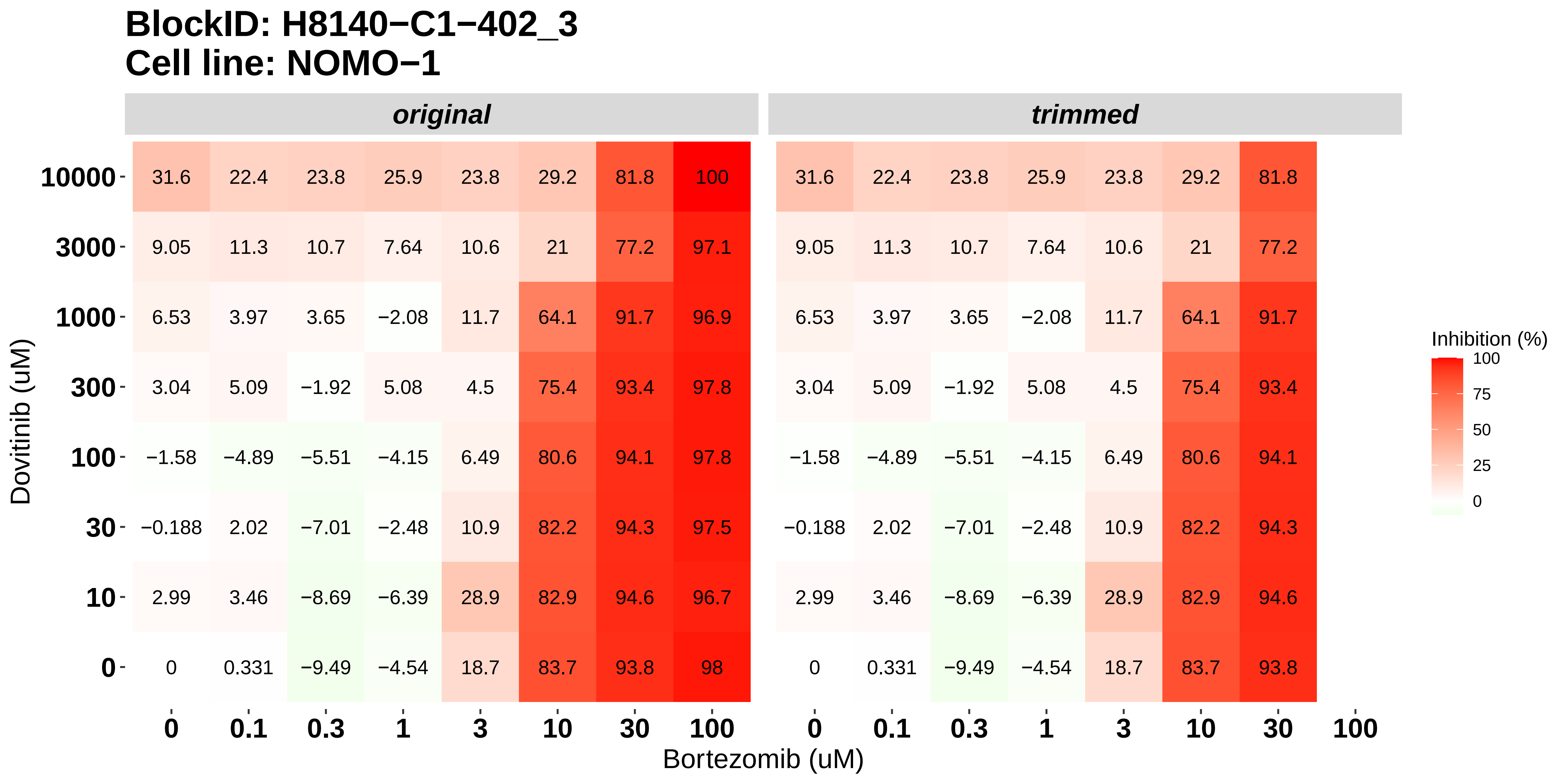

### H8140-C1-402_4_Nintedanib_Bortezomib_NOMO-1.pdf

#### BlockID: H8140−C1−402_4
#### Cell line: NOMO−1

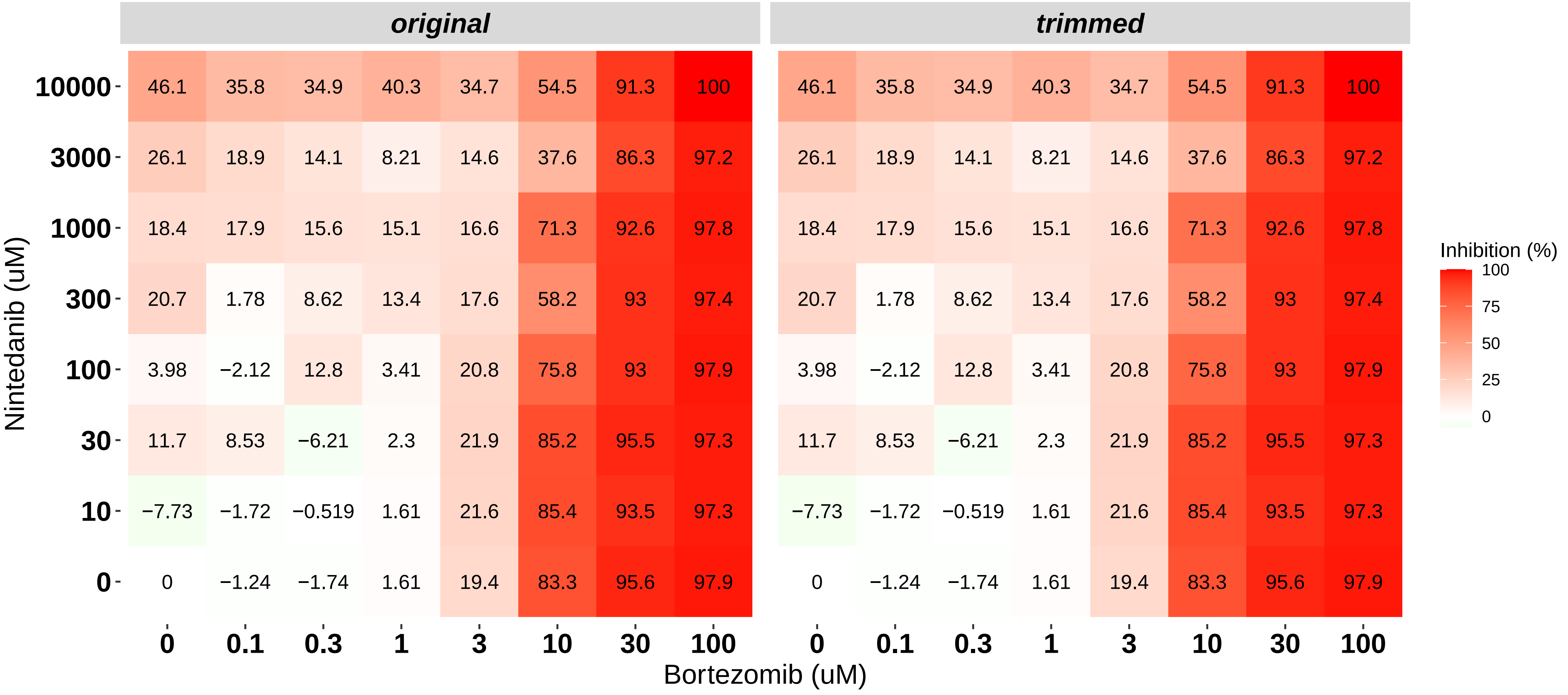

### H8140-C1-402_5_Doramapimod_Bortezomib_NOMO-1.pdf

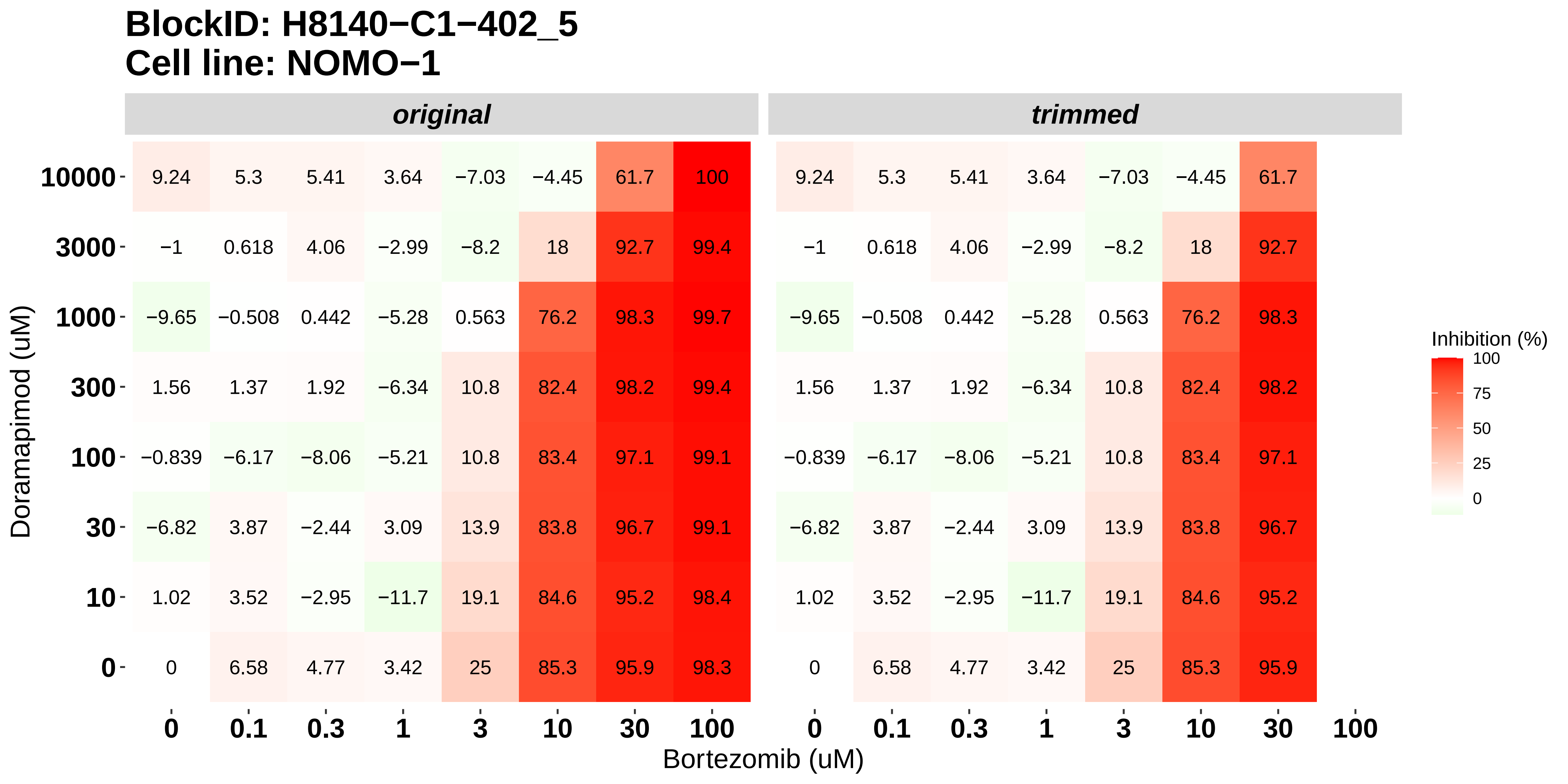

### H8140-C1-402_6_KI20227_Bortezomib_NOMO-1.pdf

#### BlockID: H8140−C1−402_6
#### Cell line: NOMO−1

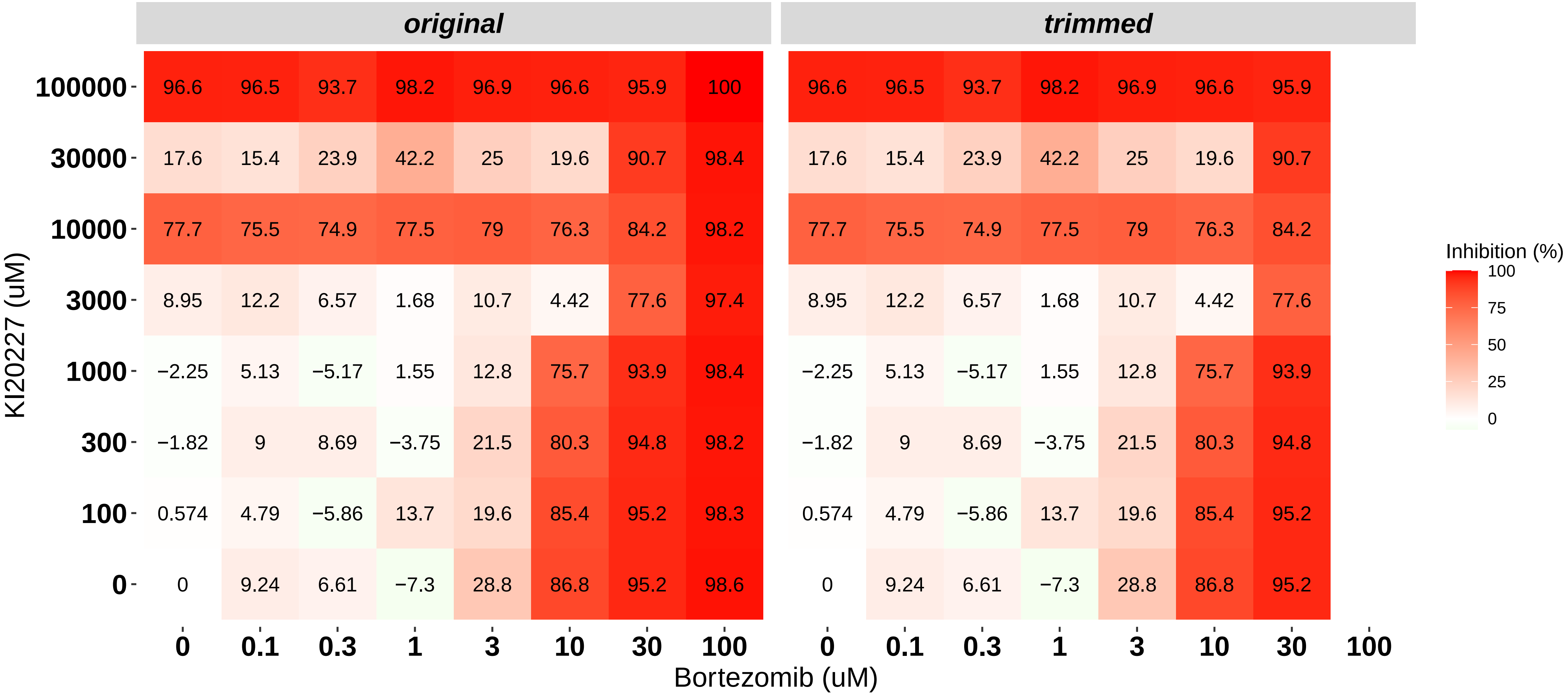

### H8140-C1-403_1_KI20227_AT7519_OCI-AML3.pdf

#### BlockID: H8140−C1−403_1
#### Cell line: OCI−AML3

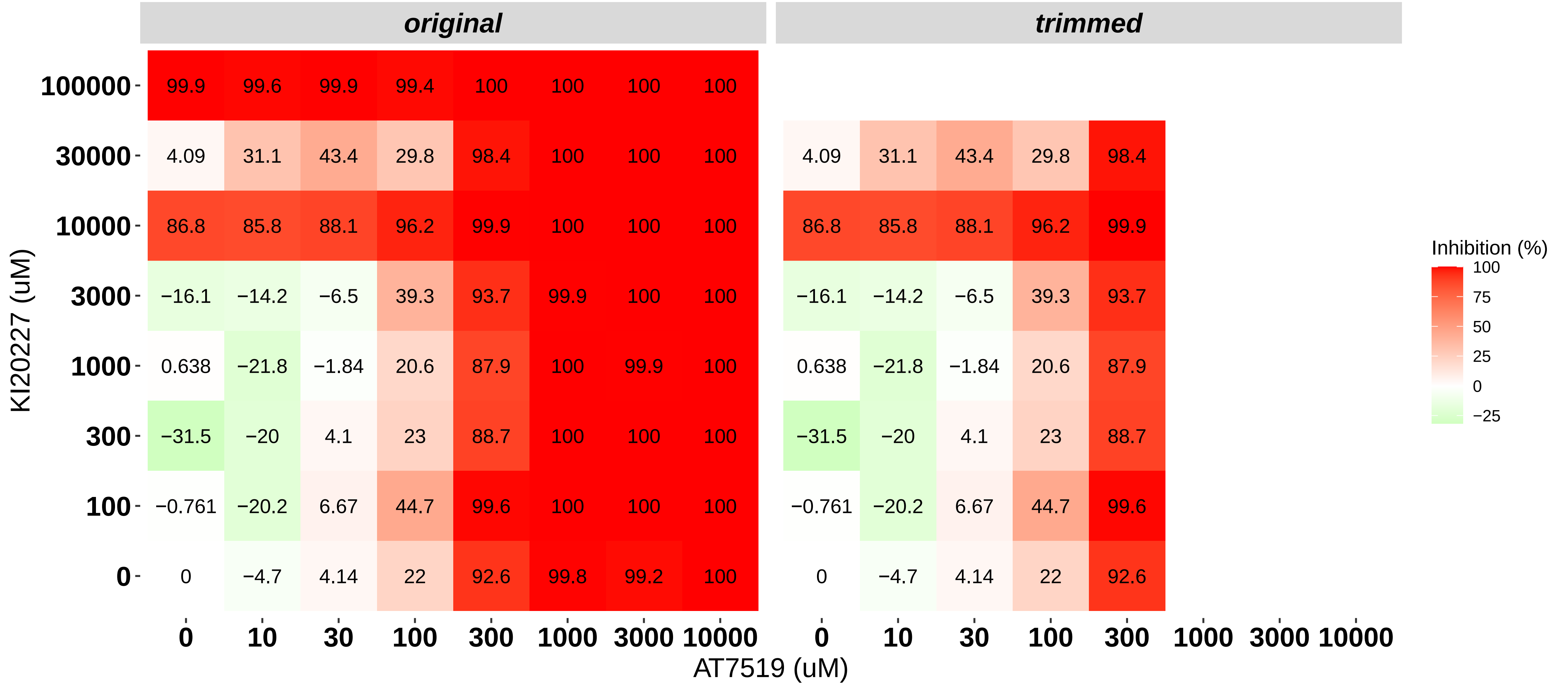

### H8140-C1-403_2_Cabozantinib_AT7519_OCI-AML3.pdf

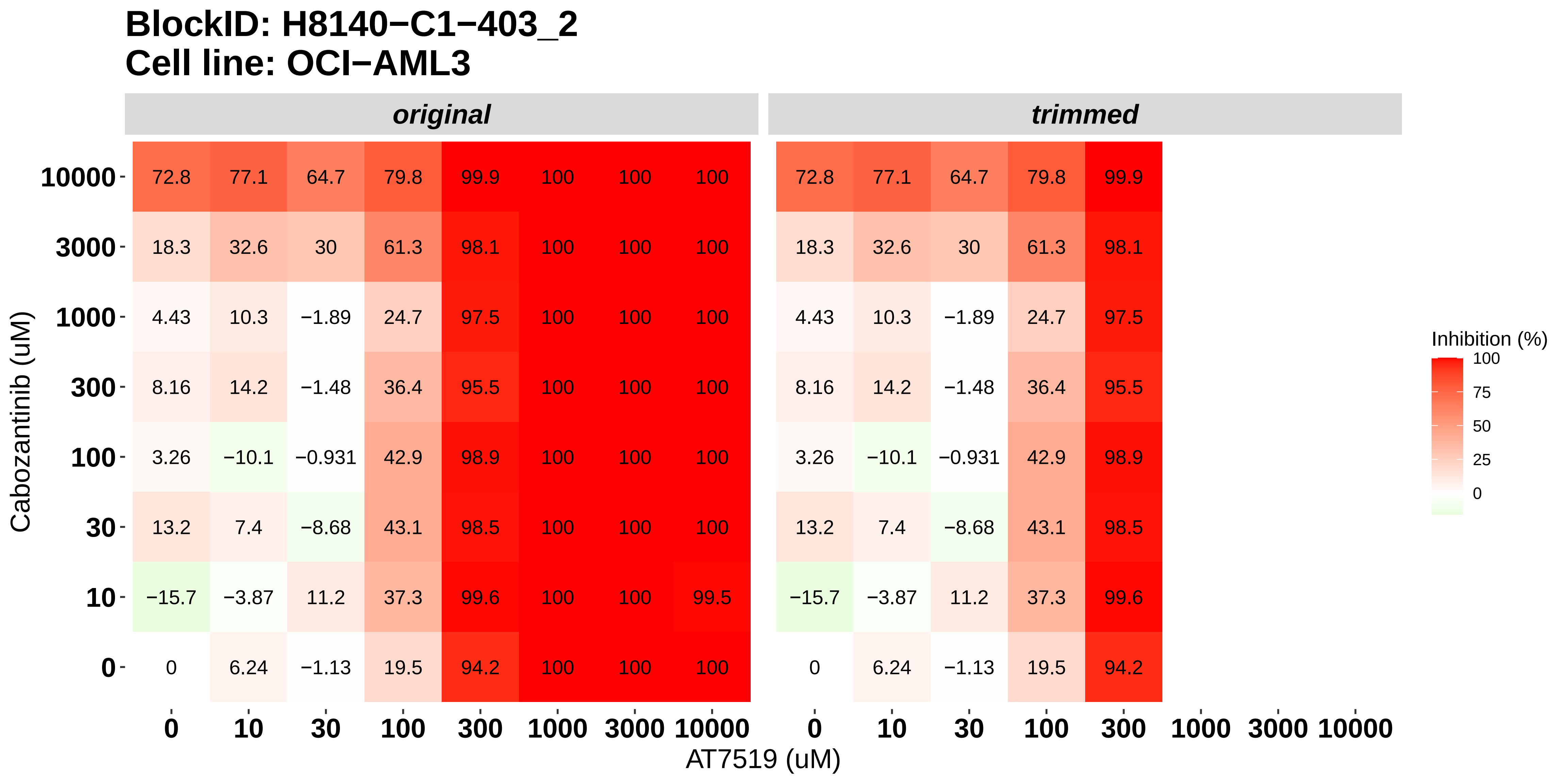

### H8140-C1-403_3_Dovitinib_Bortezomib_OCI-AML3.pdf

#### BlockID: H8140−C1−403_3
#### Cell line: OCI−AML3

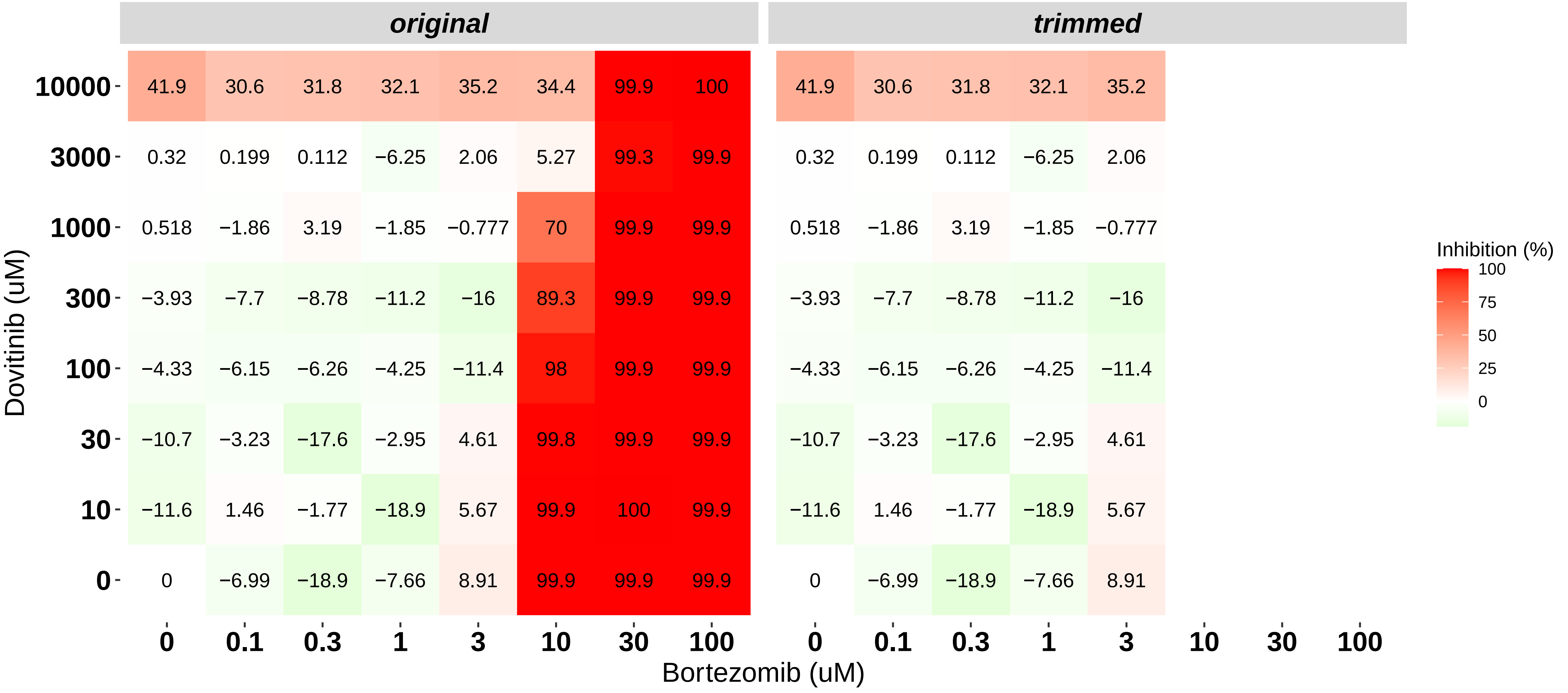

### H8140-C1-403_4_Nintedanib_Bortezomib_OCI-AML3.pdf

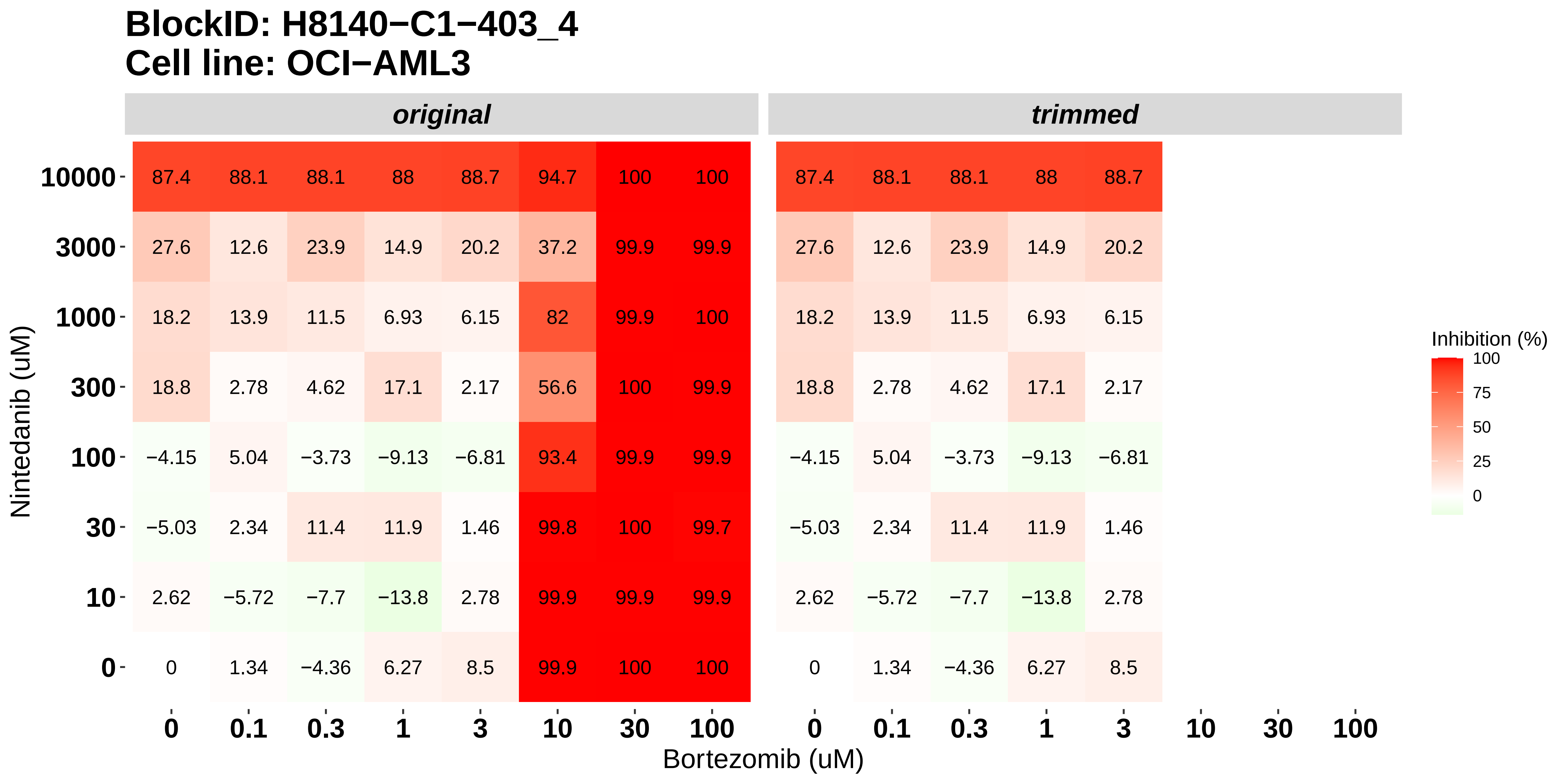

### H8140-C1-403_5_Doramapimod_Bortezomib_OCI-AML3.pdf

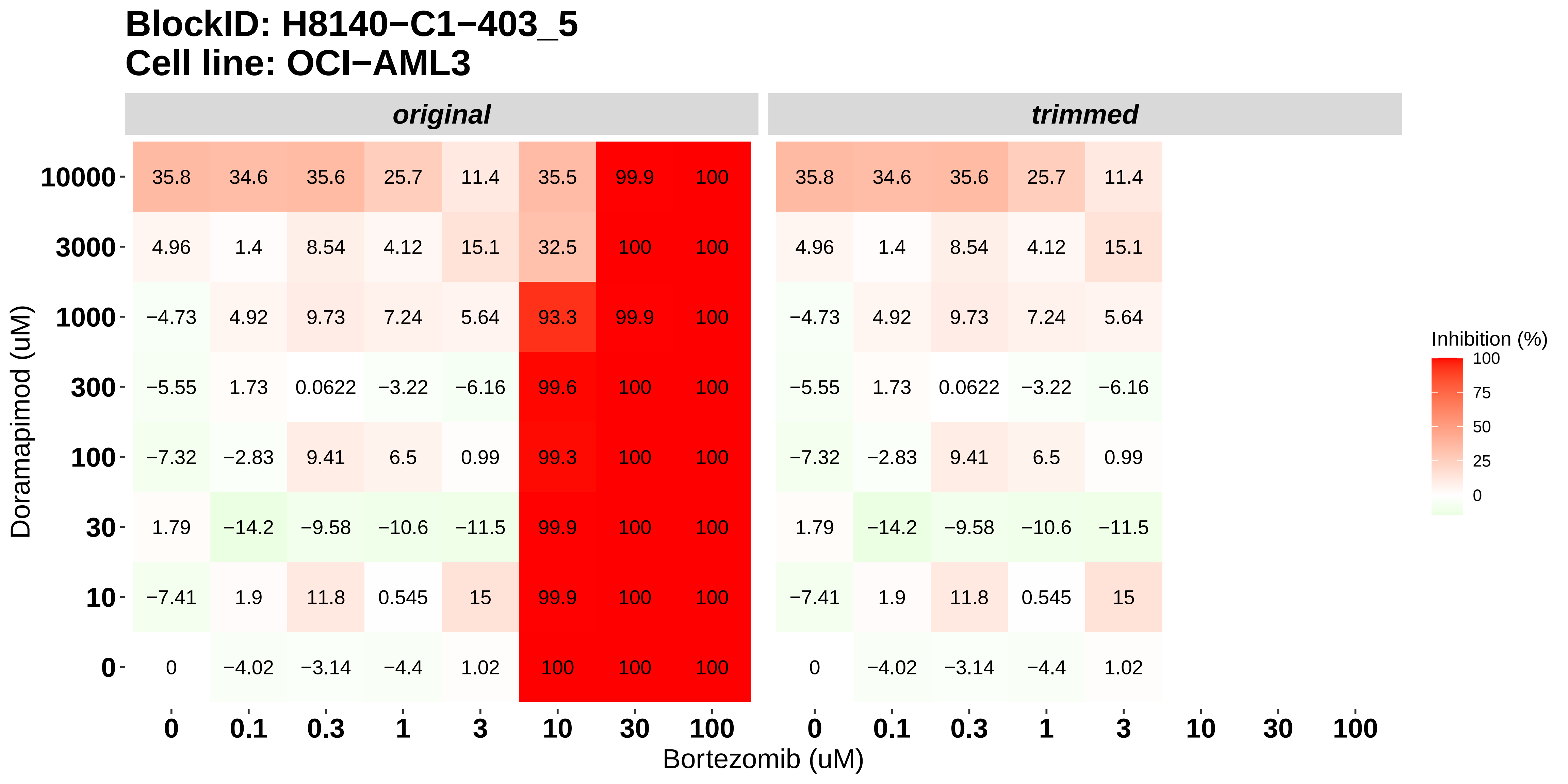

### H8140-C1-403_6_KI20227_Bortezomib_OCI-AML3.pdf

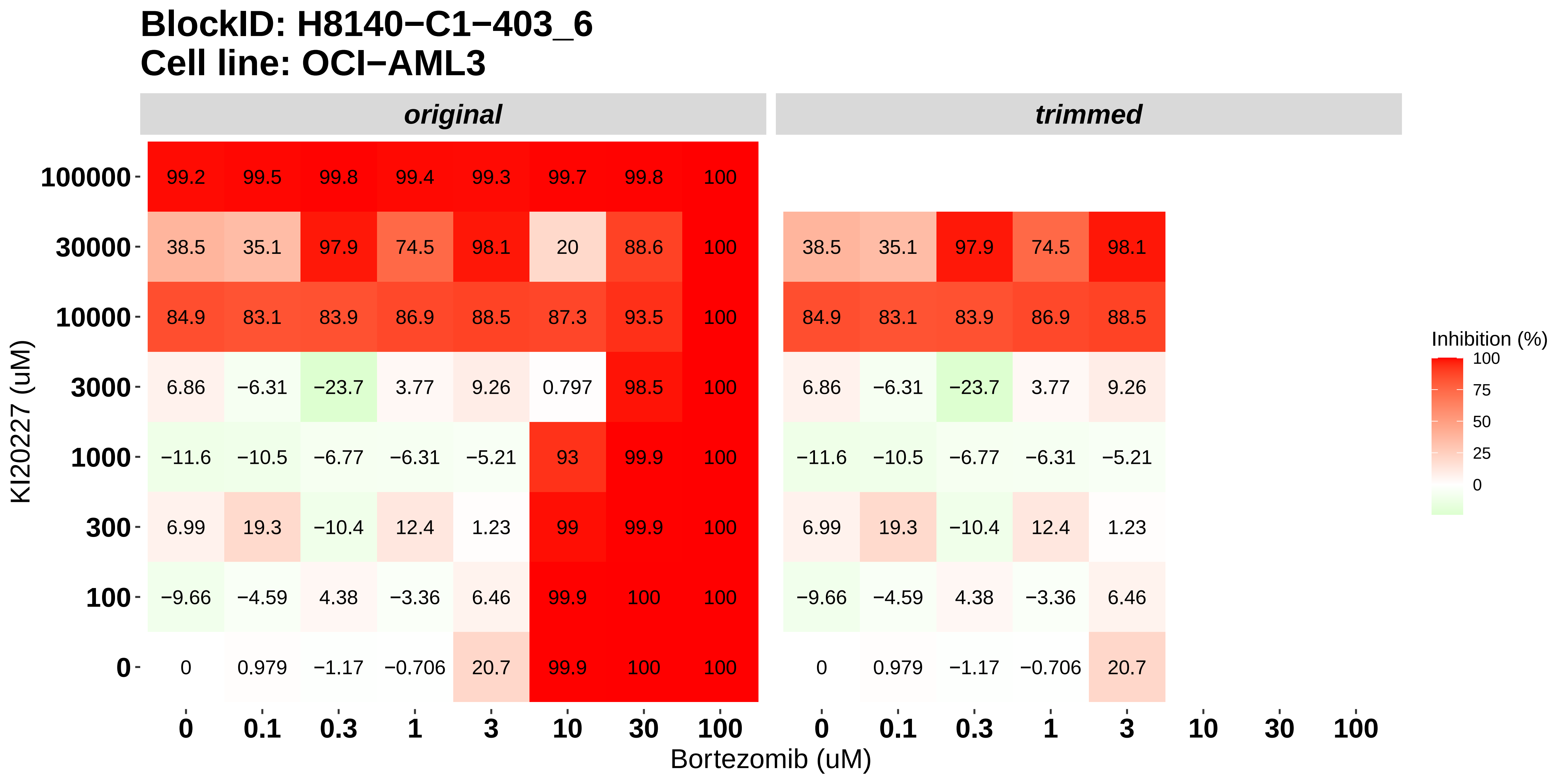

### H8140-C1-501_1_Cabozantinib_Bortezomib_MOLM-16.pdf

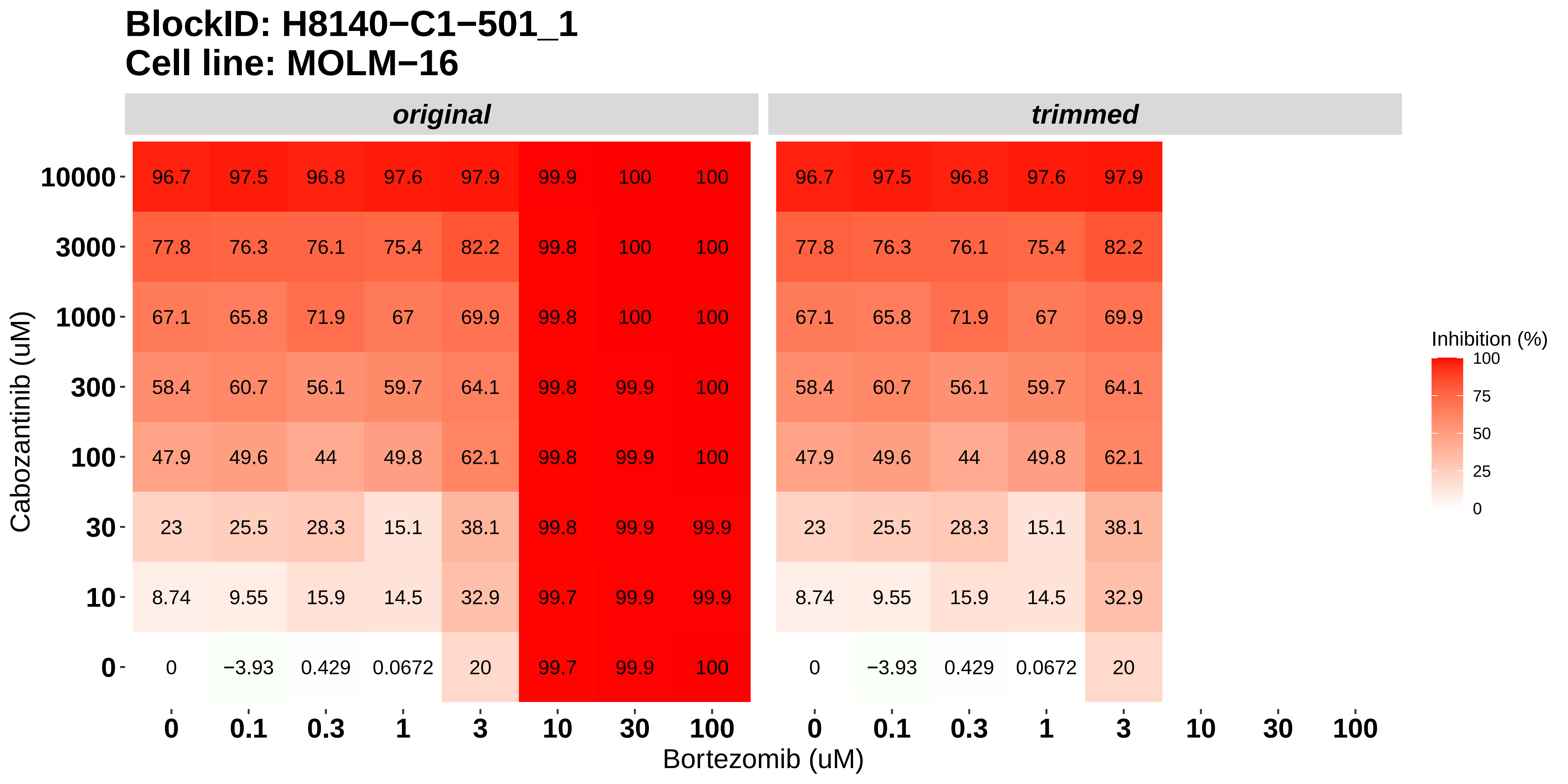

### H8140-C1-501_2_Alvocidib_SNS-032_MOLM-16.pdf

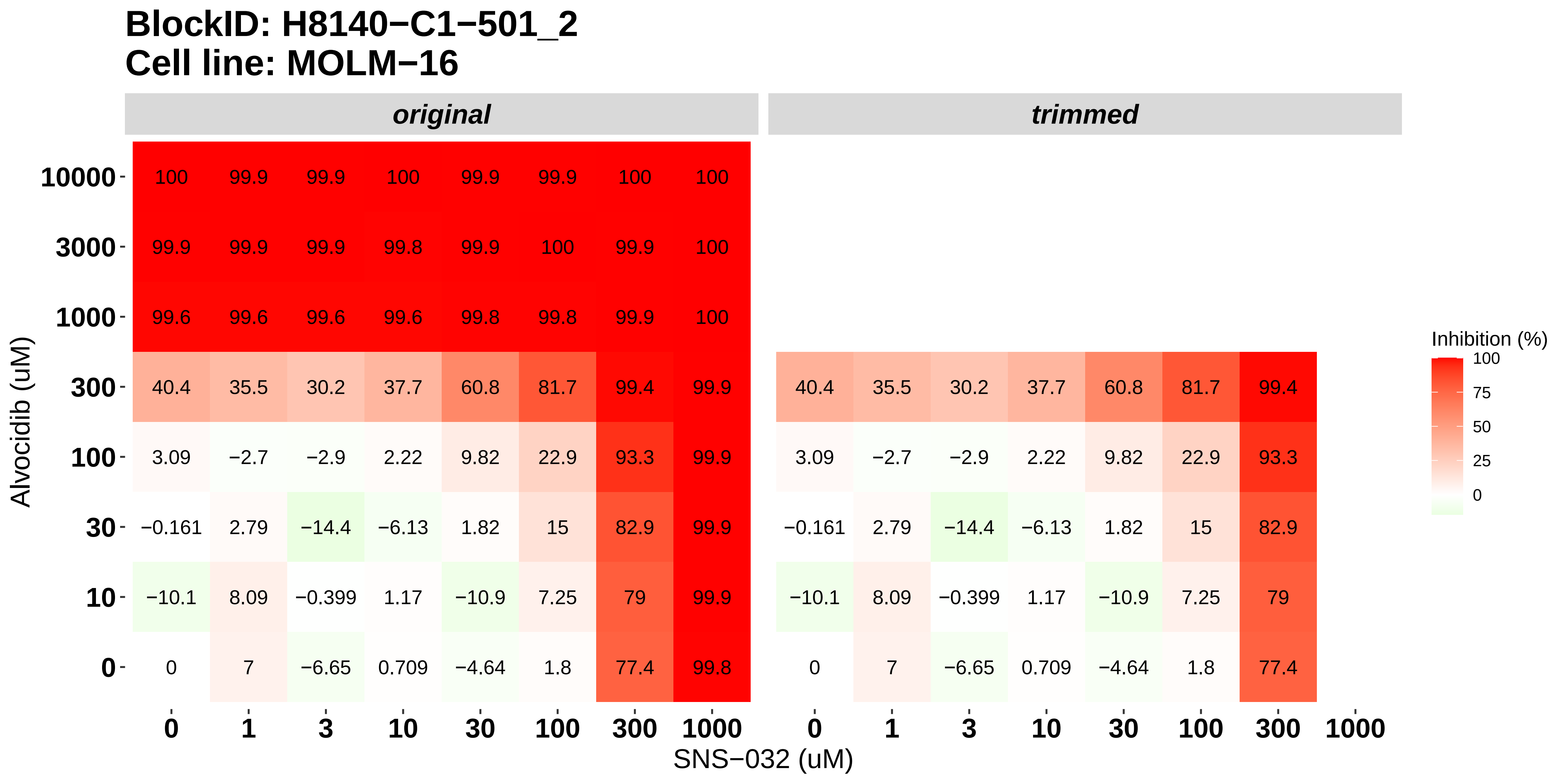

### H8140-C1-501_3_Panobinostat_SNS-032_MOLM-16.pdf

#### BlockID: H8140−C1−501_3
#### Cell line: MOLM−16

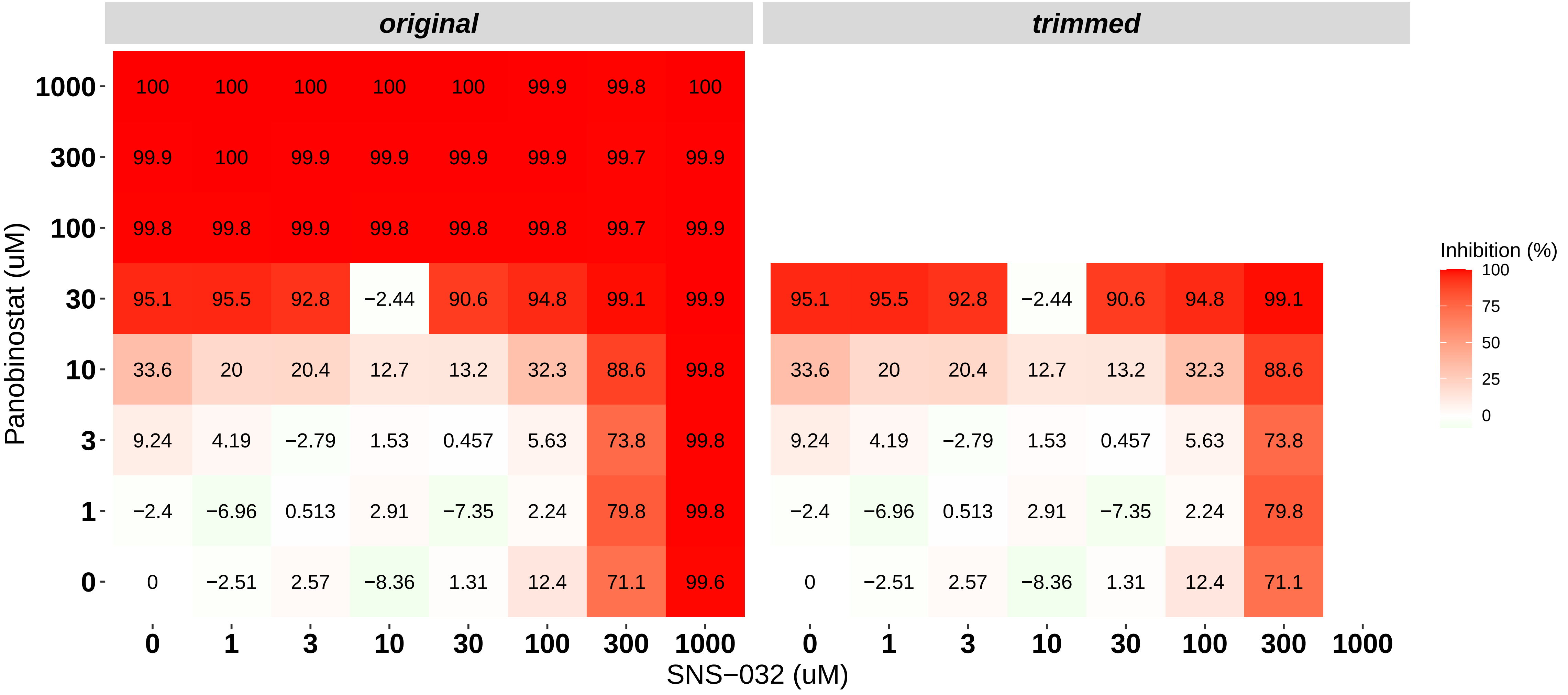

### H8140-C1-501_4_AT7519_SNS-032_MOLM-16.pdf

#### BlockID: H8140−C1−501_4
#### Cell line: MOLM−16

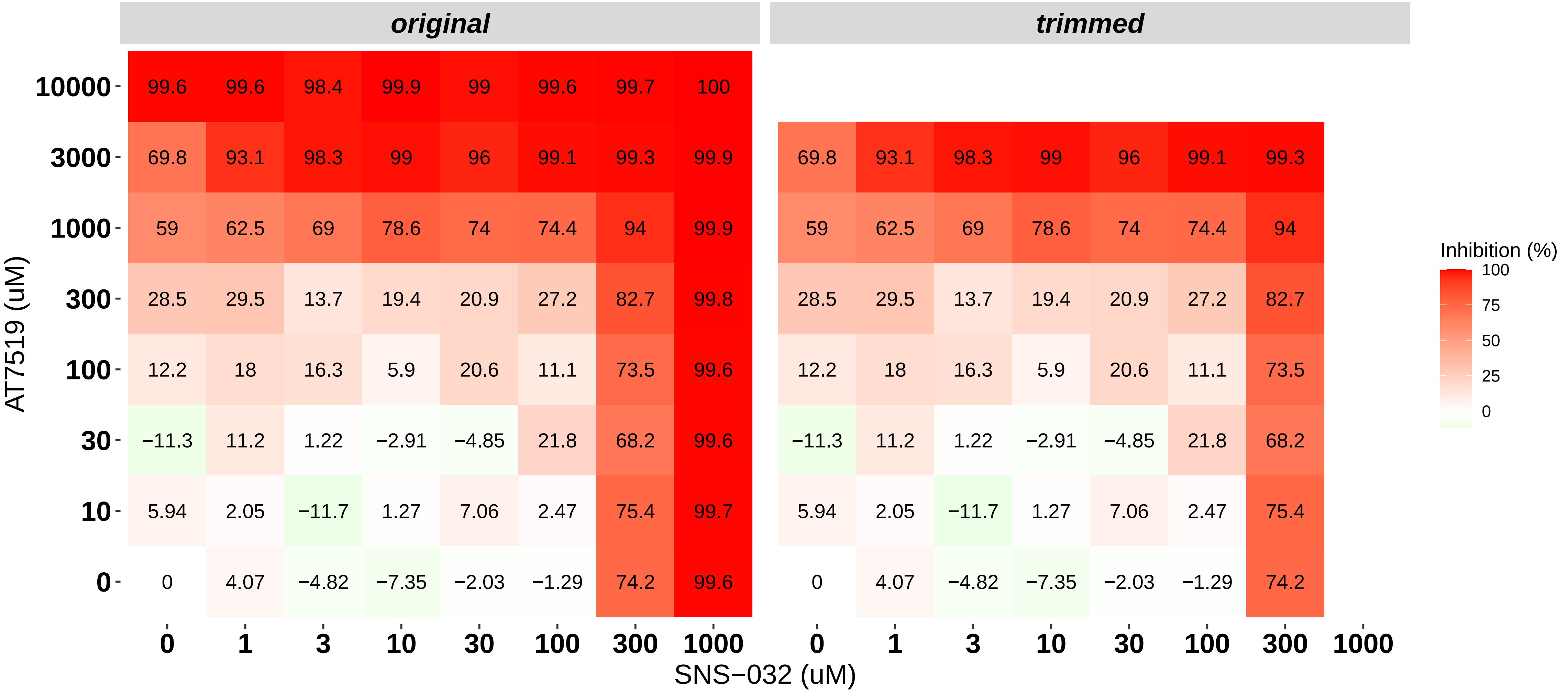

### H8140-C1-501_5_Bortezomib_SNS-032_MOLM-16.pdf

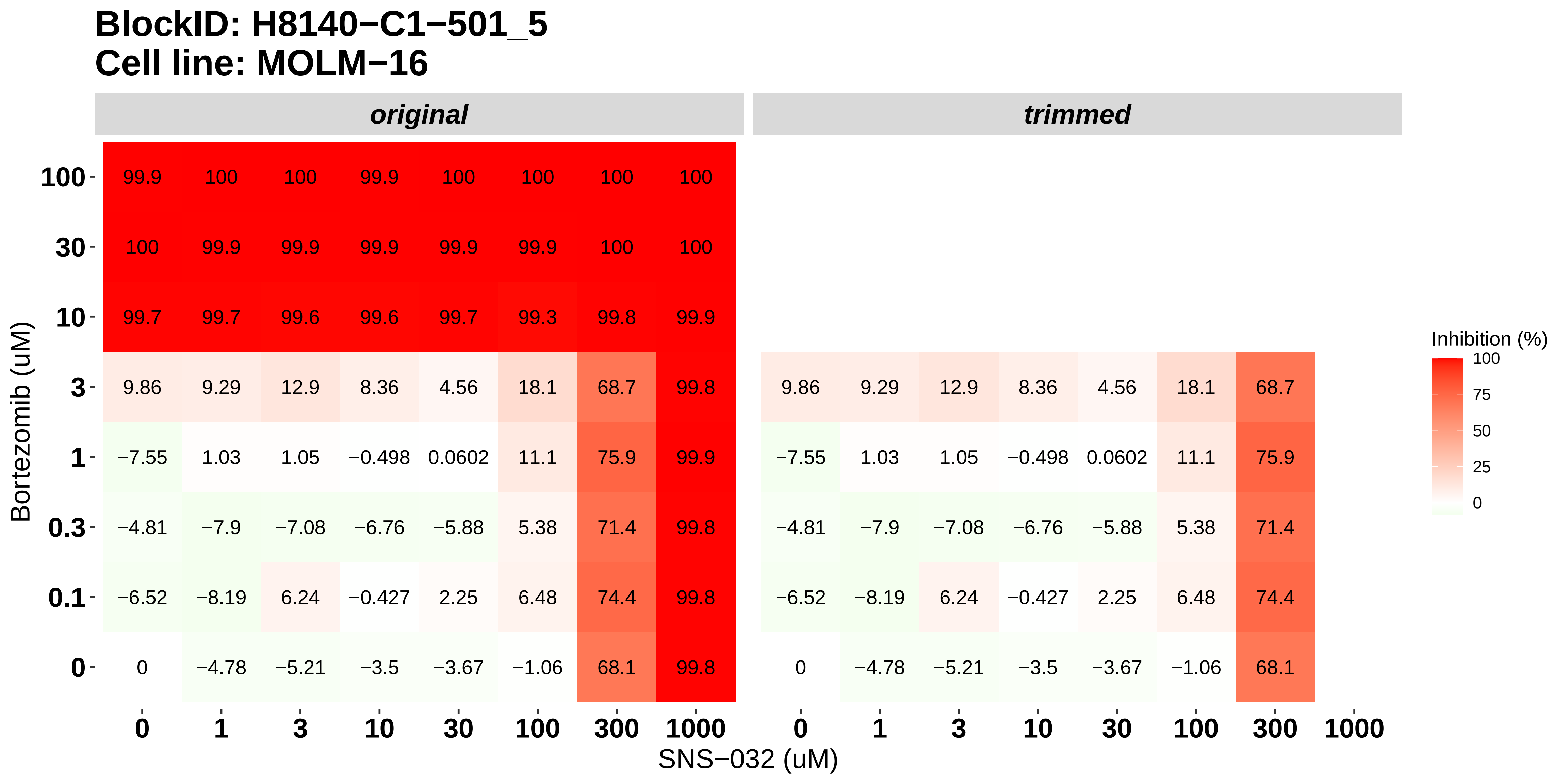

### H8140-C1-501_6_Panobinostat_Alvocidib_MOLM-16.pdf

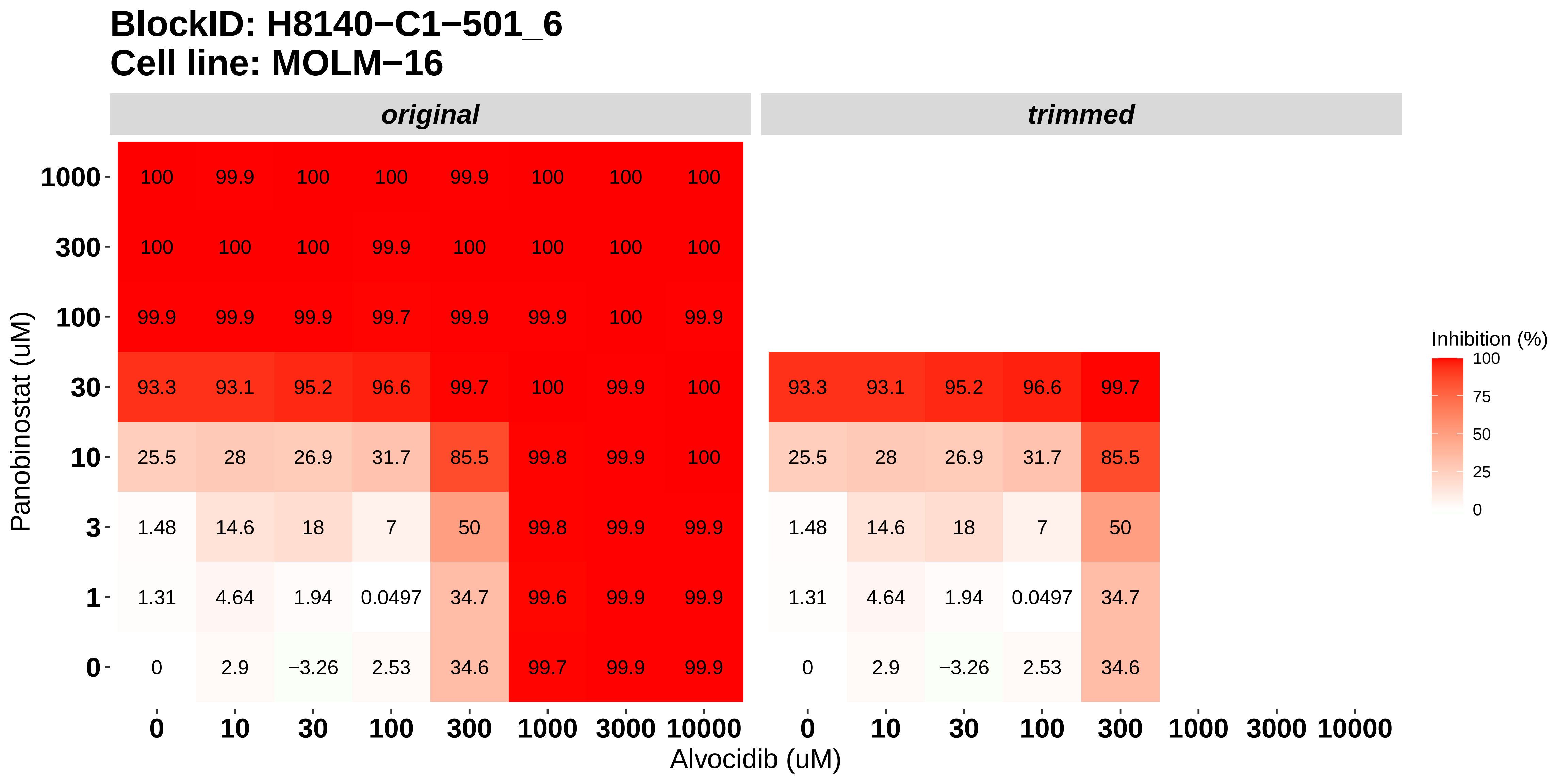

### H8140-C1-502_1_Cabozantinib_Bortezomib_NOMO-1.pdf

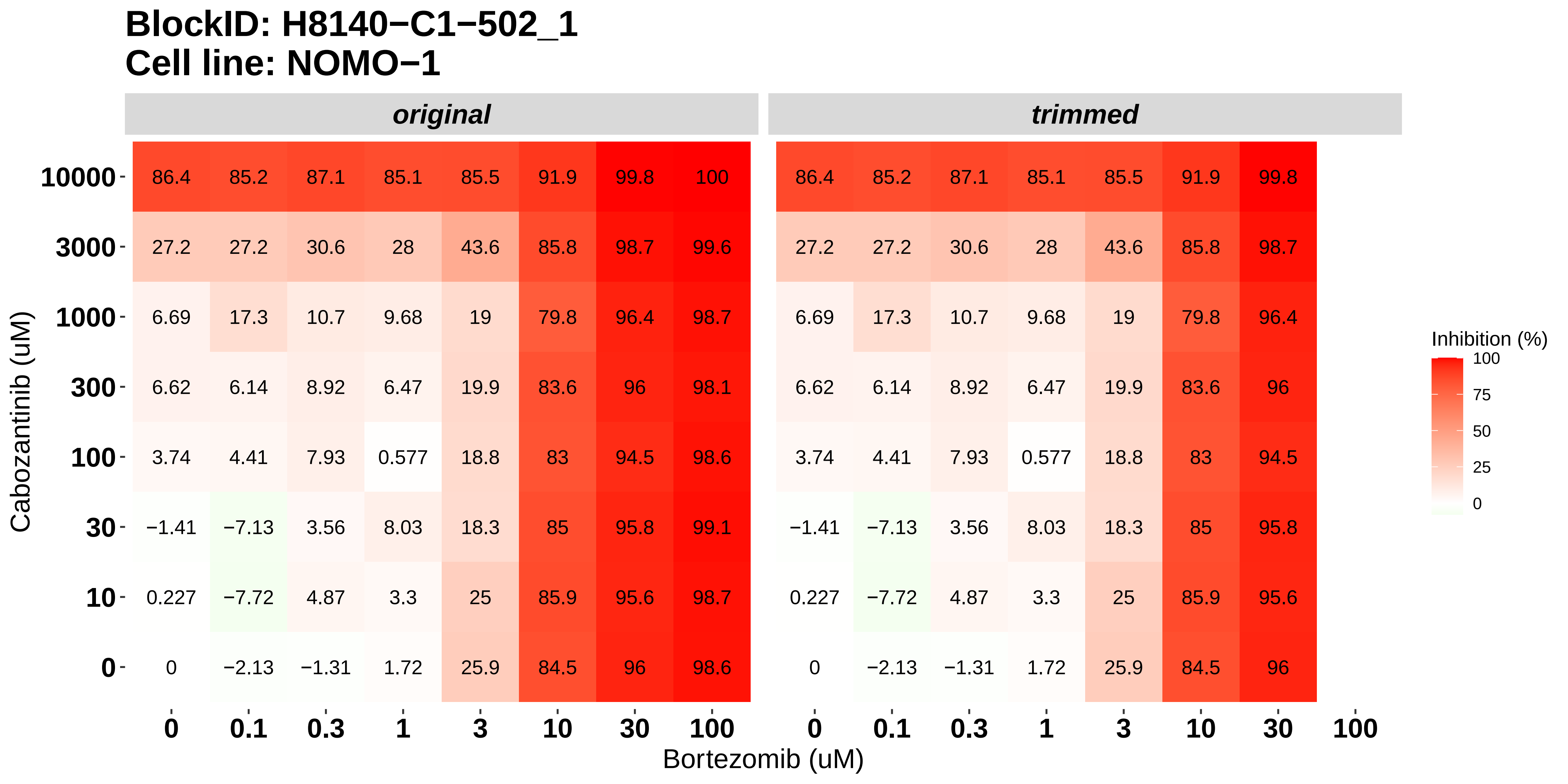

### H8140-C1-502_2_Alvocidib_SNS-032_NOMO-1.pdf

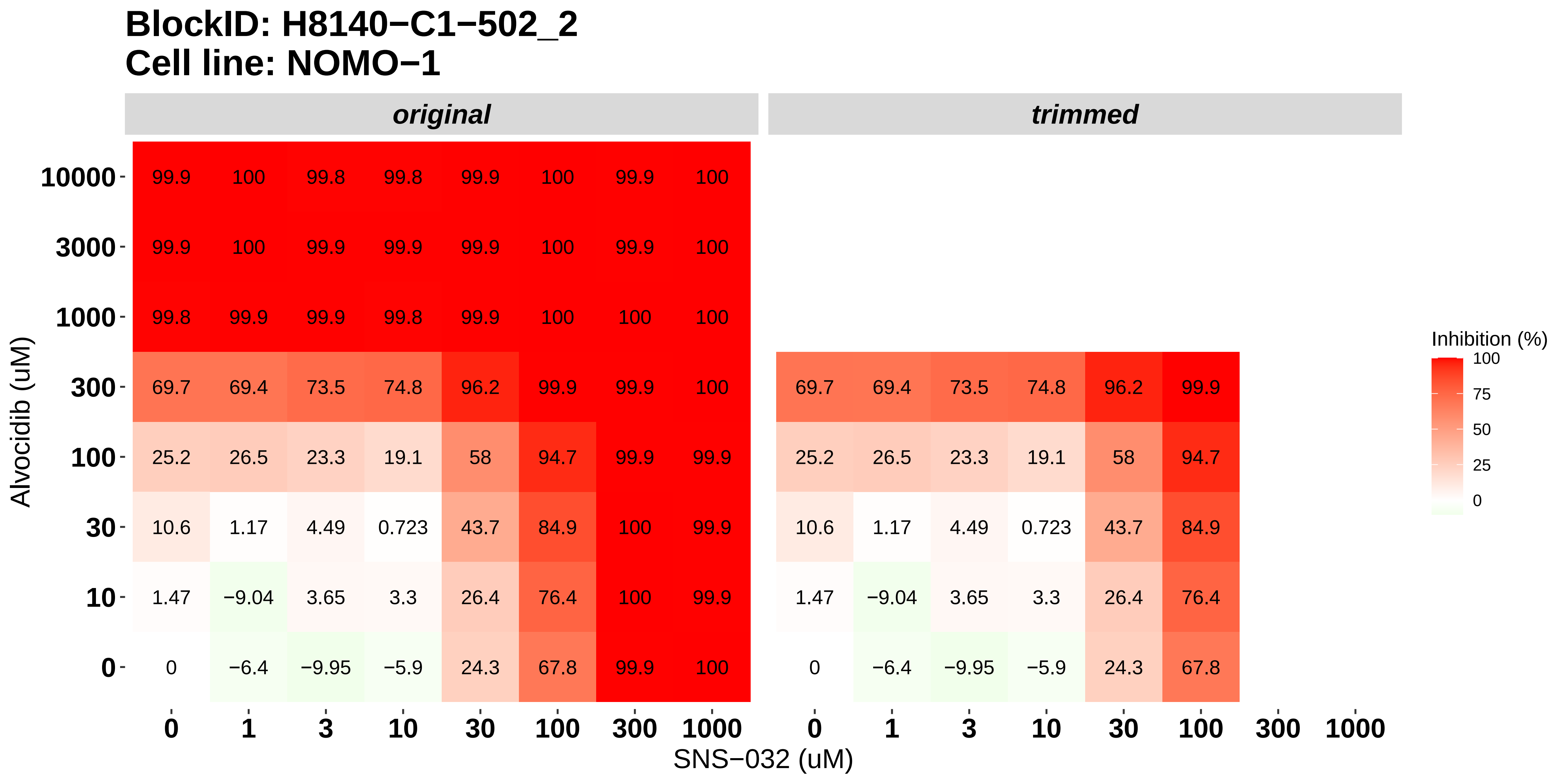

### H8140-C1-502_3_Panobinostat_SNS-032_NOMO-1.pdf

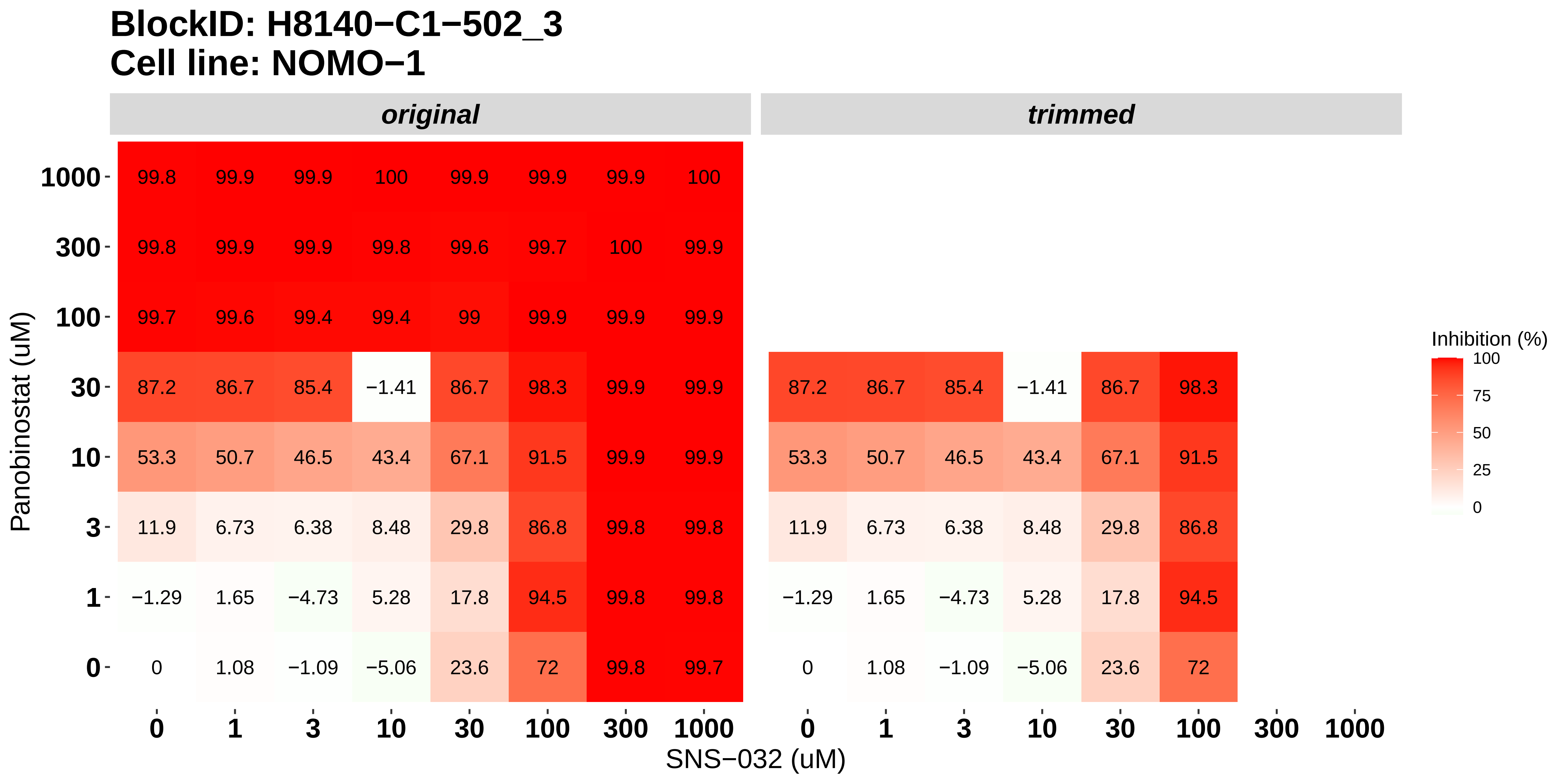

### H8140-C1-502_4_AT7519_SNS-032_NOMO-1.pdf

#### BlockID: H8140−C1−502_4
#### Cell line: NOMO−1

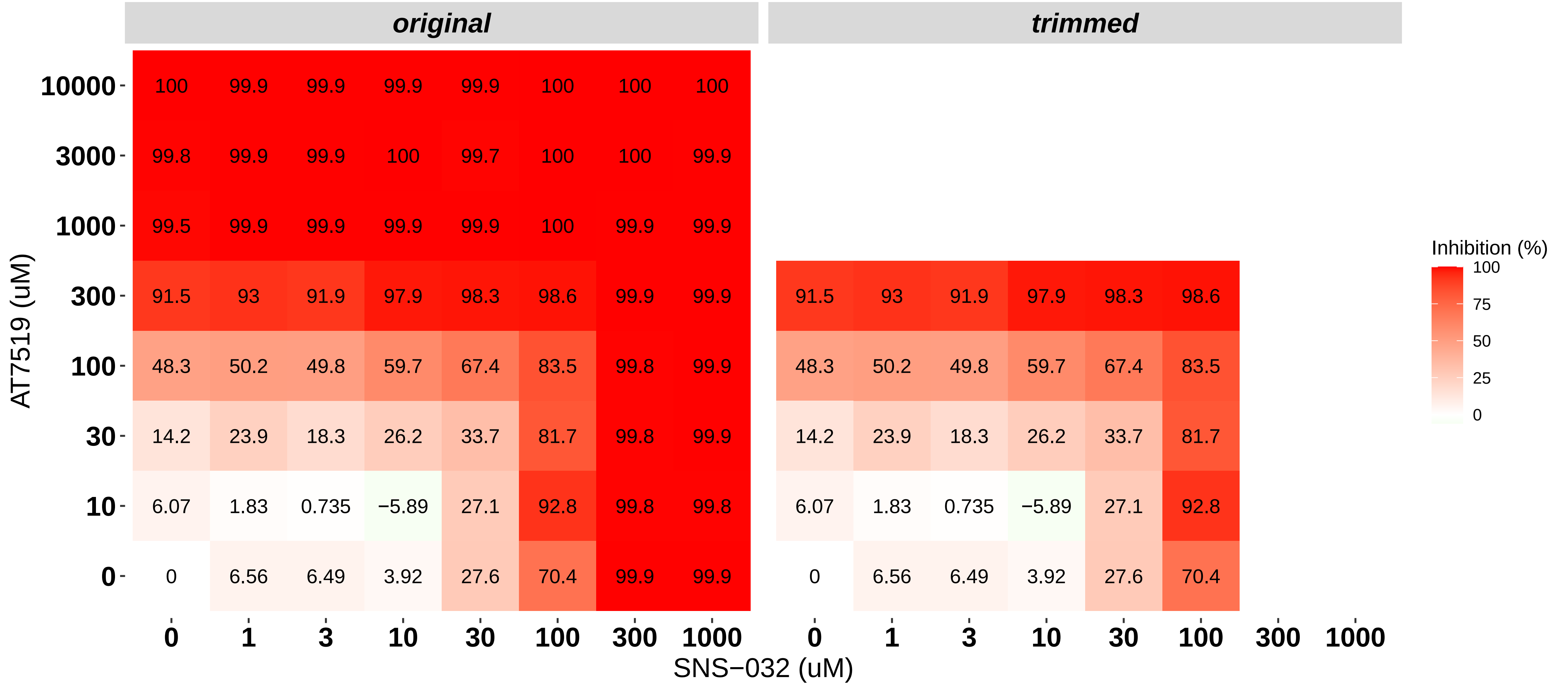

### H8140-C1-502_5_Bortezomib_SNS-032_NOMO-1.pdf

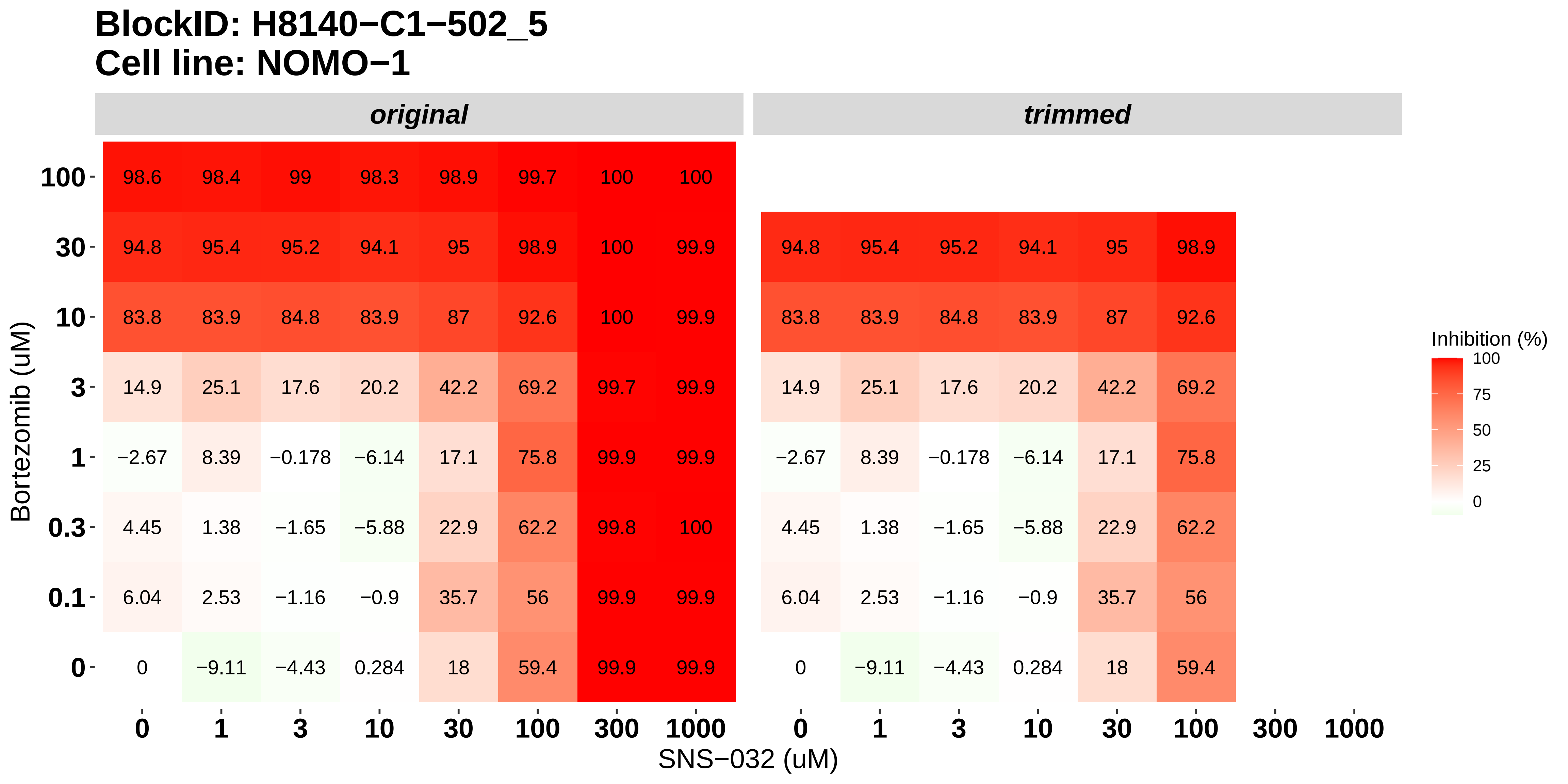

### H8140-C1-502_6_Panobinostat_Alvocidib_NOMO-1.pdf

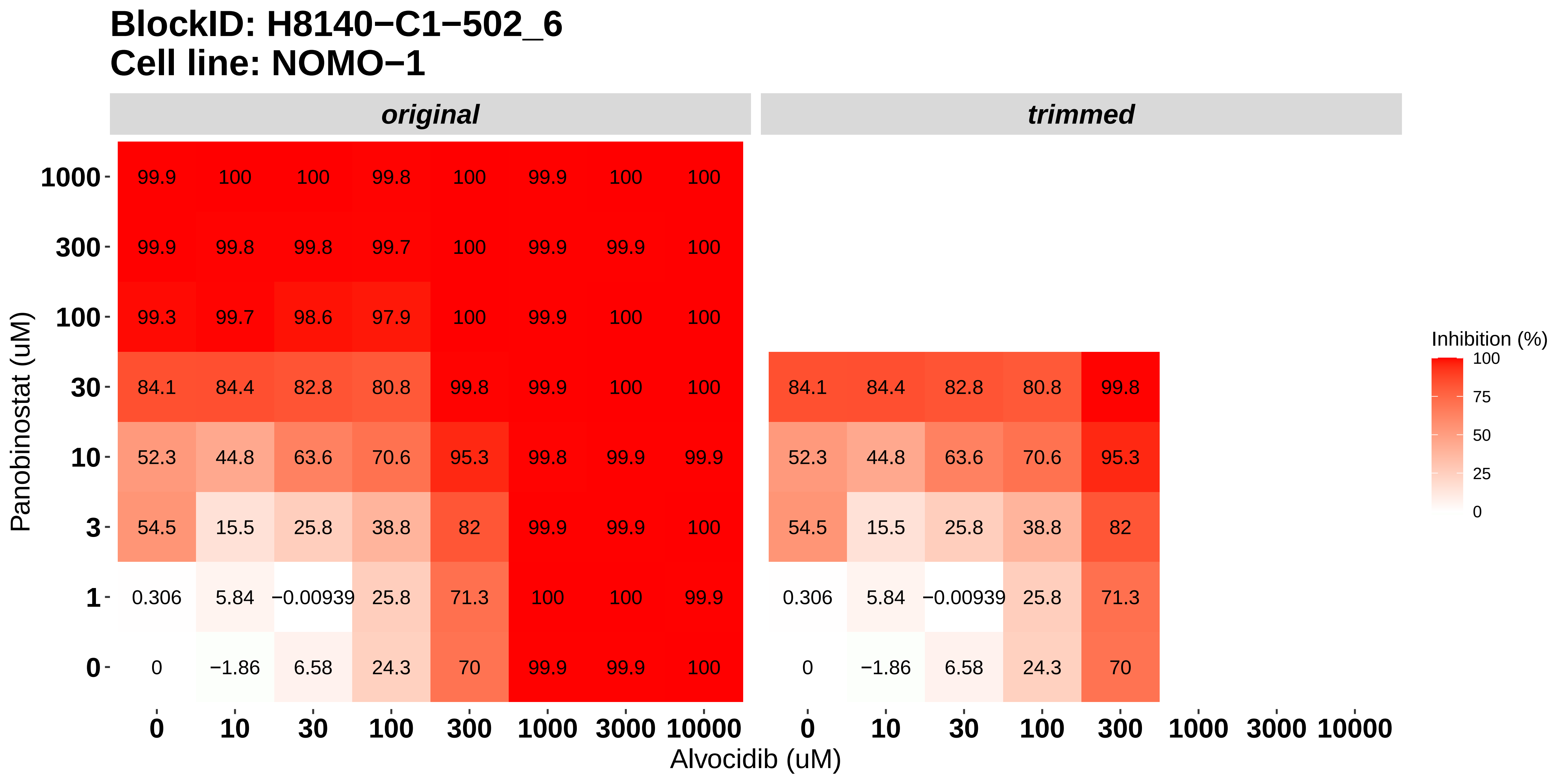

### H8140-C1-503_1_Cabozantinib_Bortezomib_OCI-AML3.pdf

#### BlockID: H8140−C1−503_1
#### Cell line: OCI−AML3

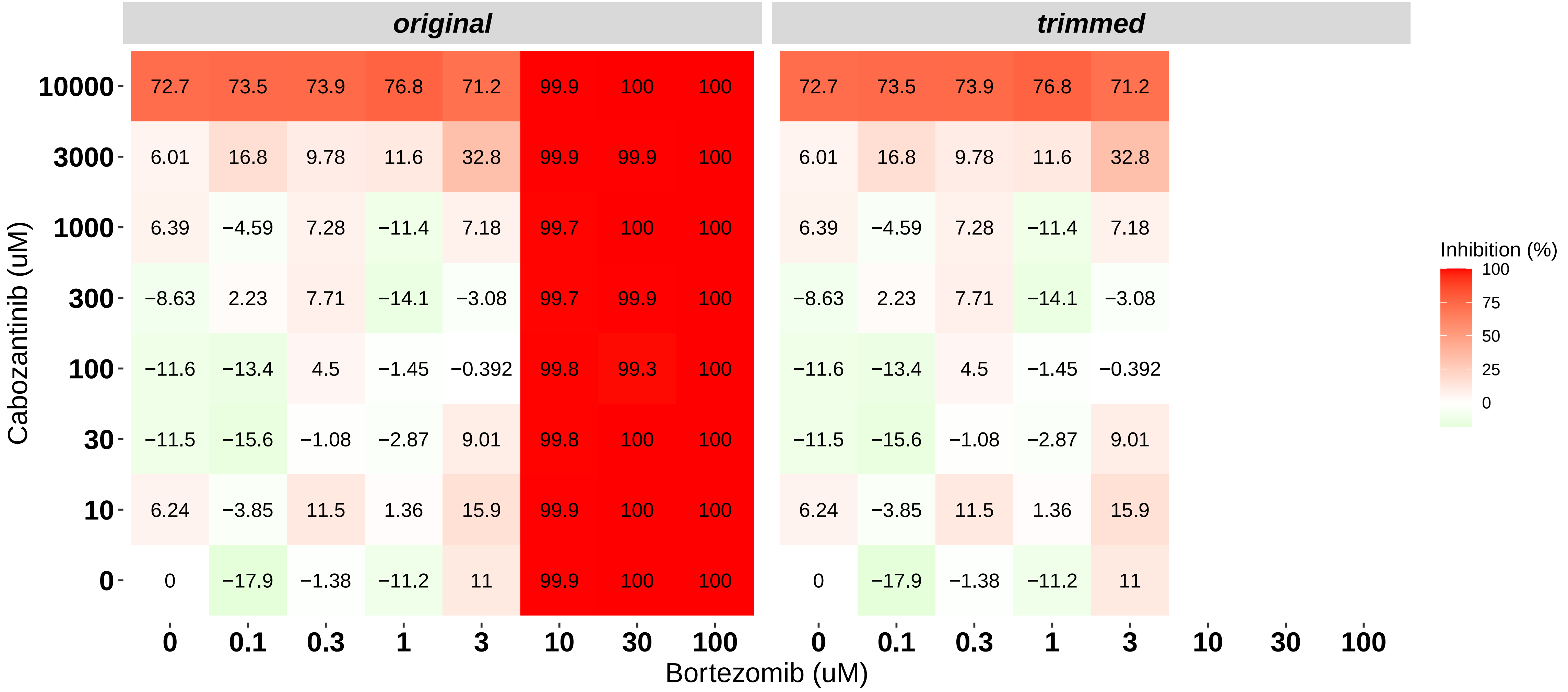

### H8140-C1-503_2_Alvocidib_SNS-032_OCI-AML3.pdf

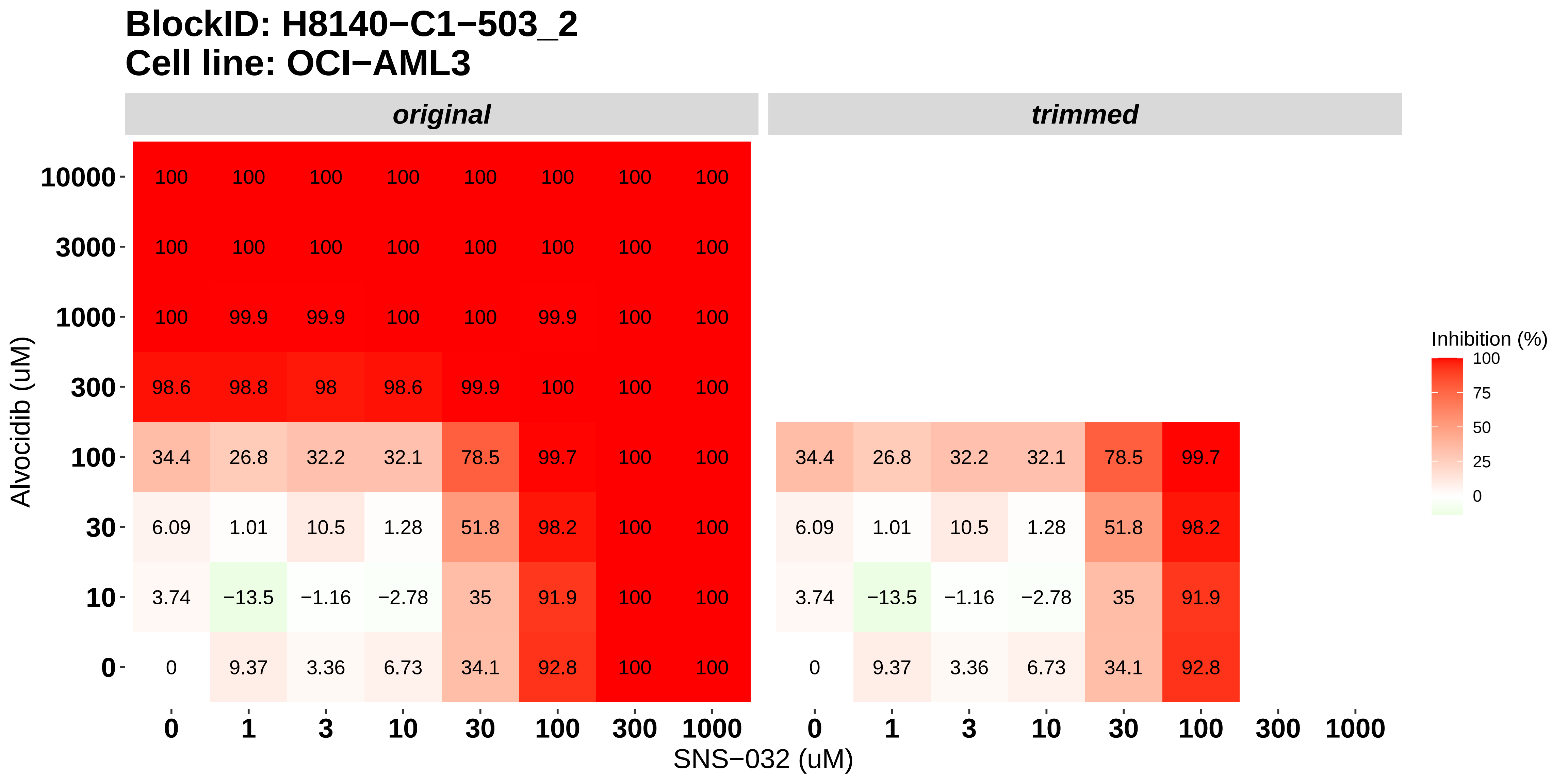

### H8140-C1-503_3_Panobinostat_SNS-032_OCI-AML3.pdf

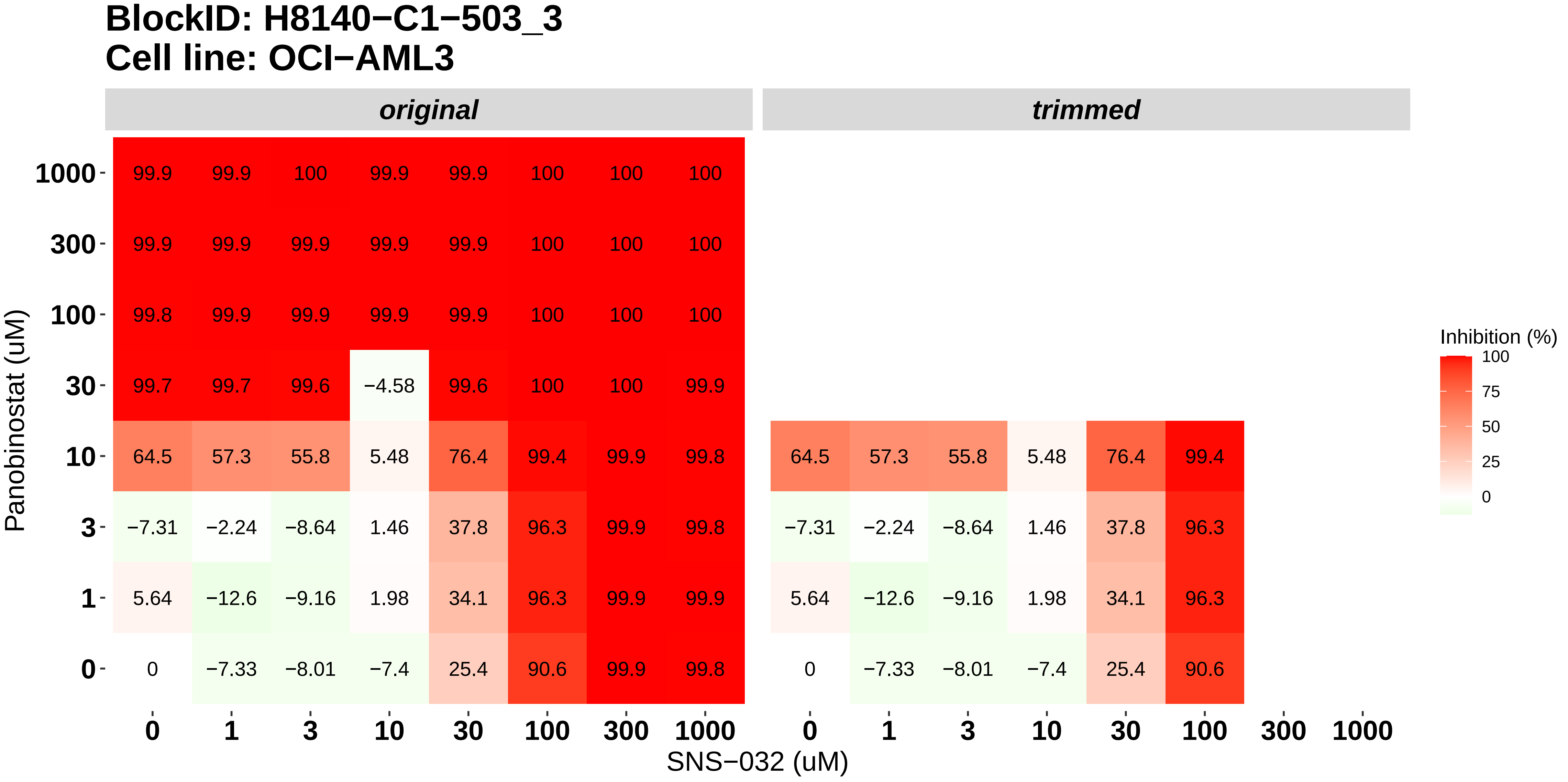

### H8140-C1-503_4_AT7519_SNS-032_OCI-AML3.pdf

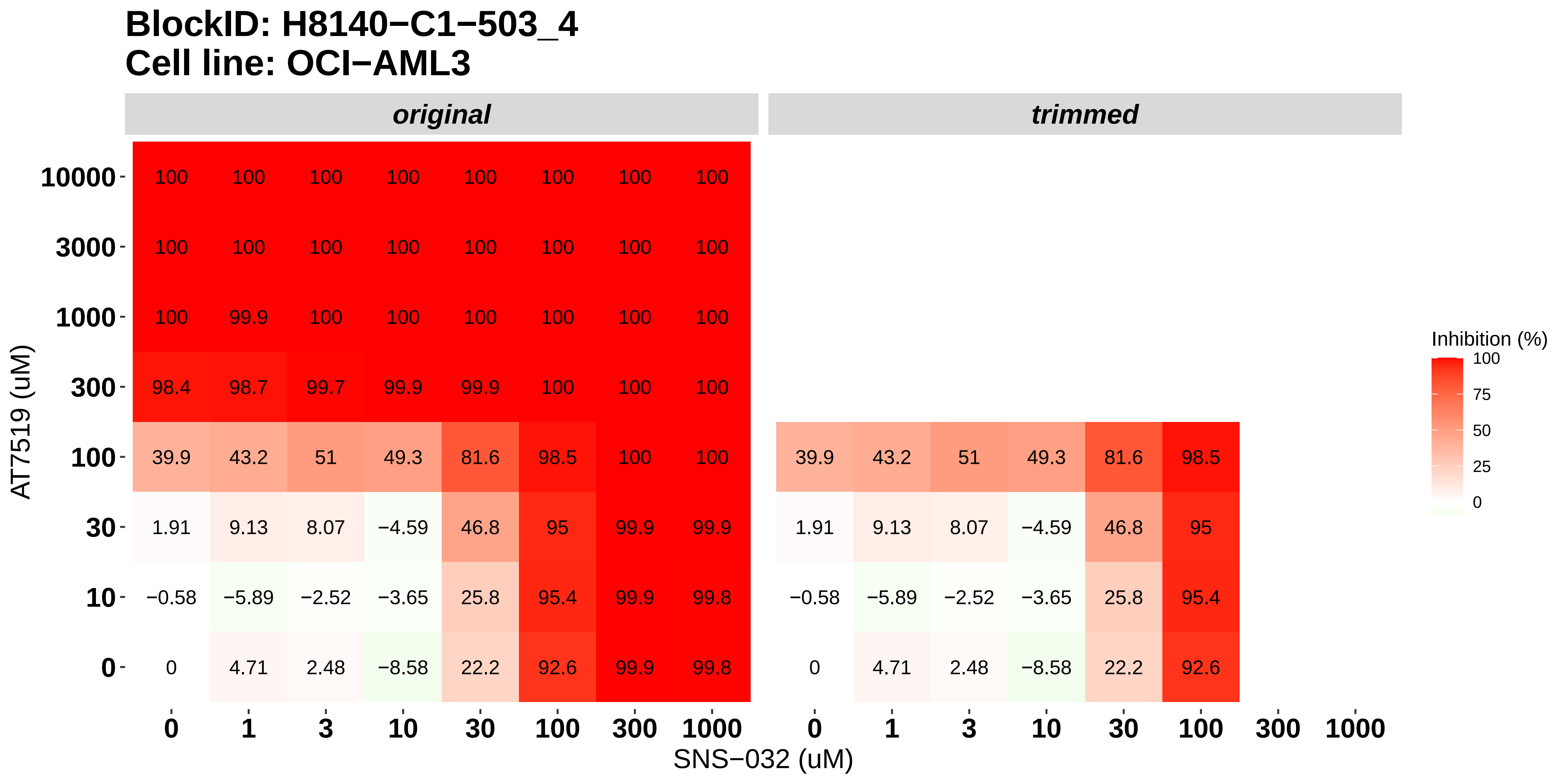

### H8140-C1-503_5_Bortezomib_SNS-032_OCI-AML3.pdf

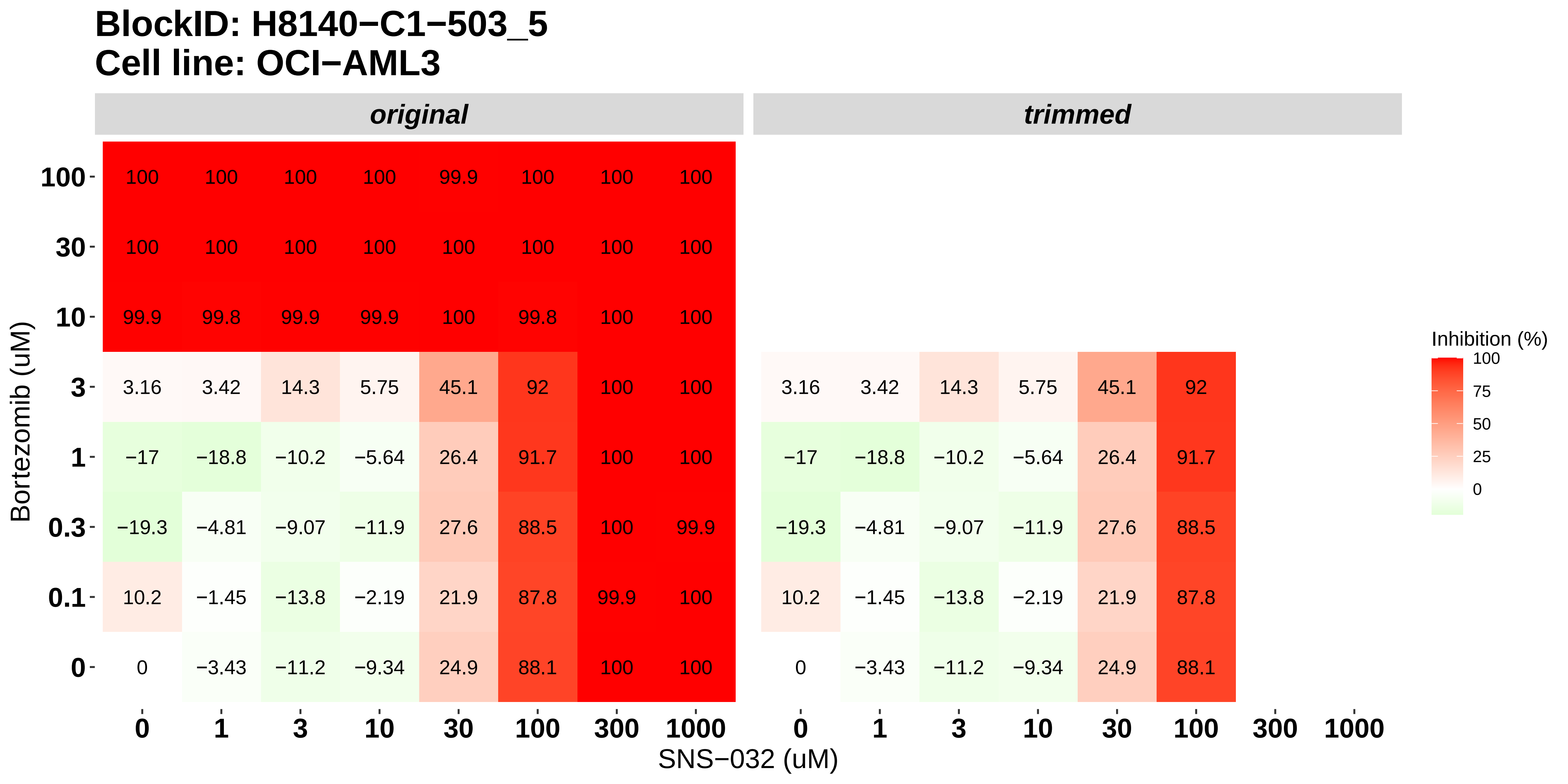

### H8140-C1-503_6_Panobinostat_Alvocidib_OCI-AML3.pdf

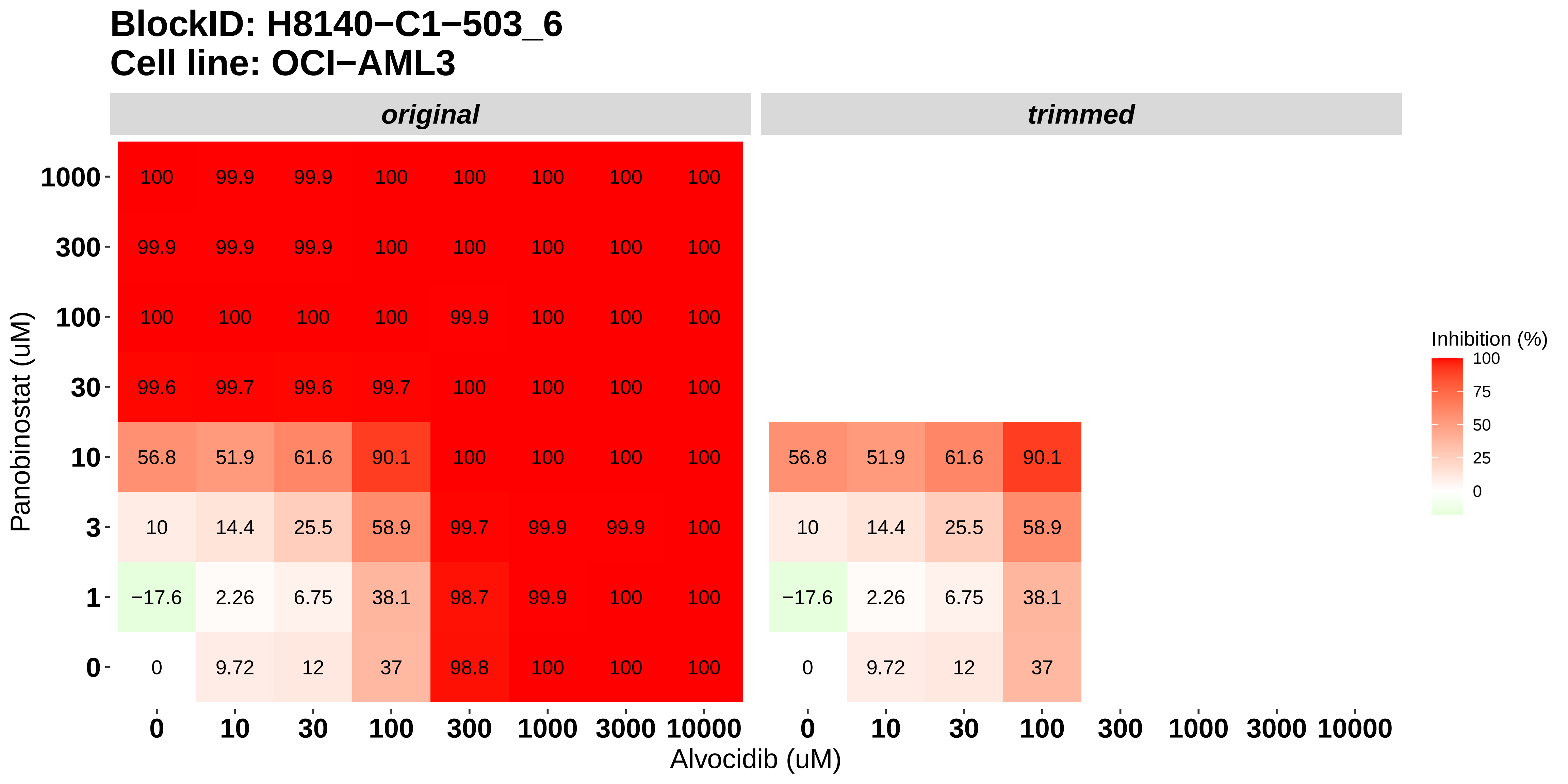

### H8140-C1-601_1_AT7519_Alvocidib_MOLM-16.pdf

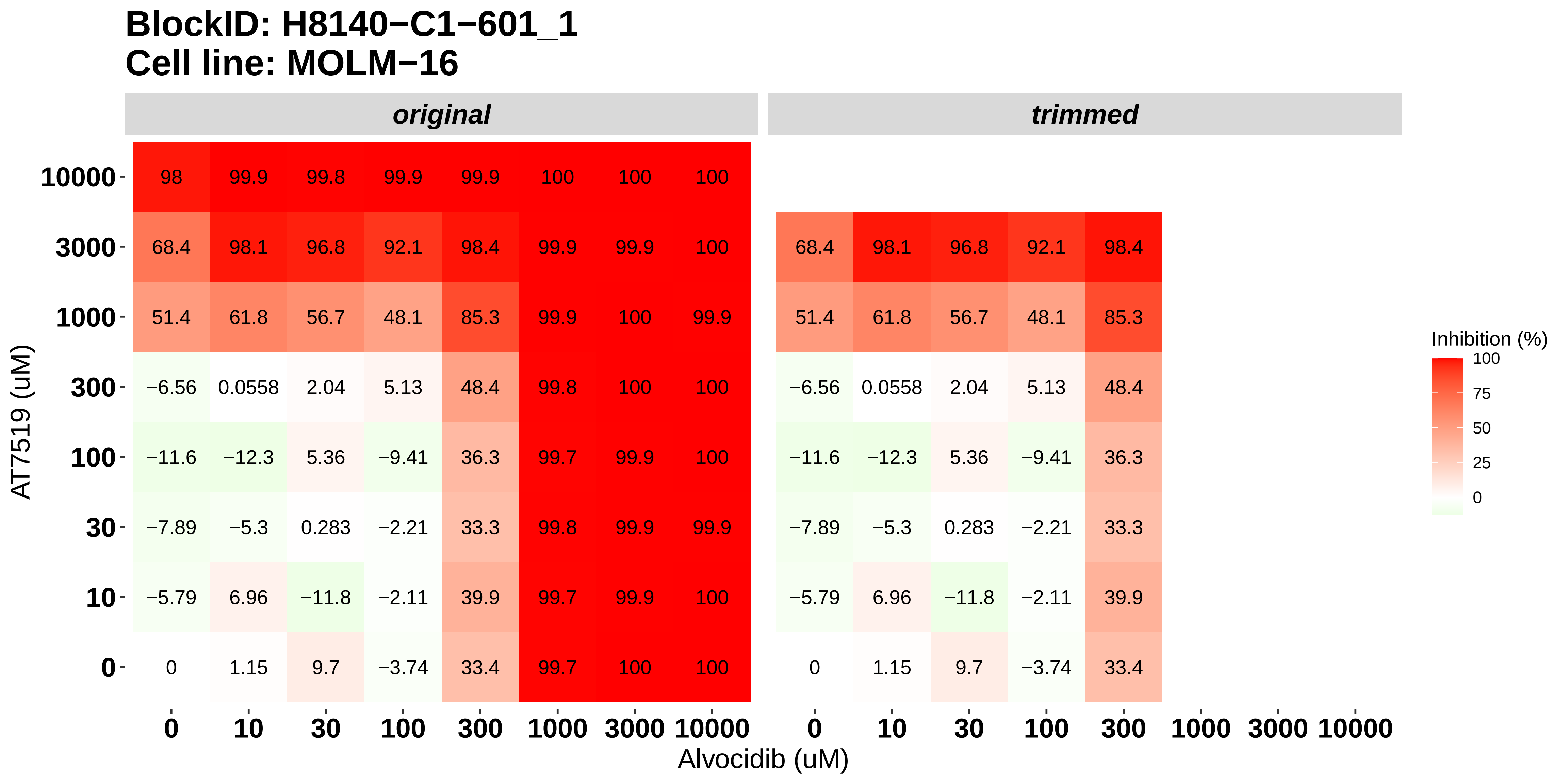

### H8140-C1-601_2_Bortezomib_Alvocidib_MOLM-16.pdf

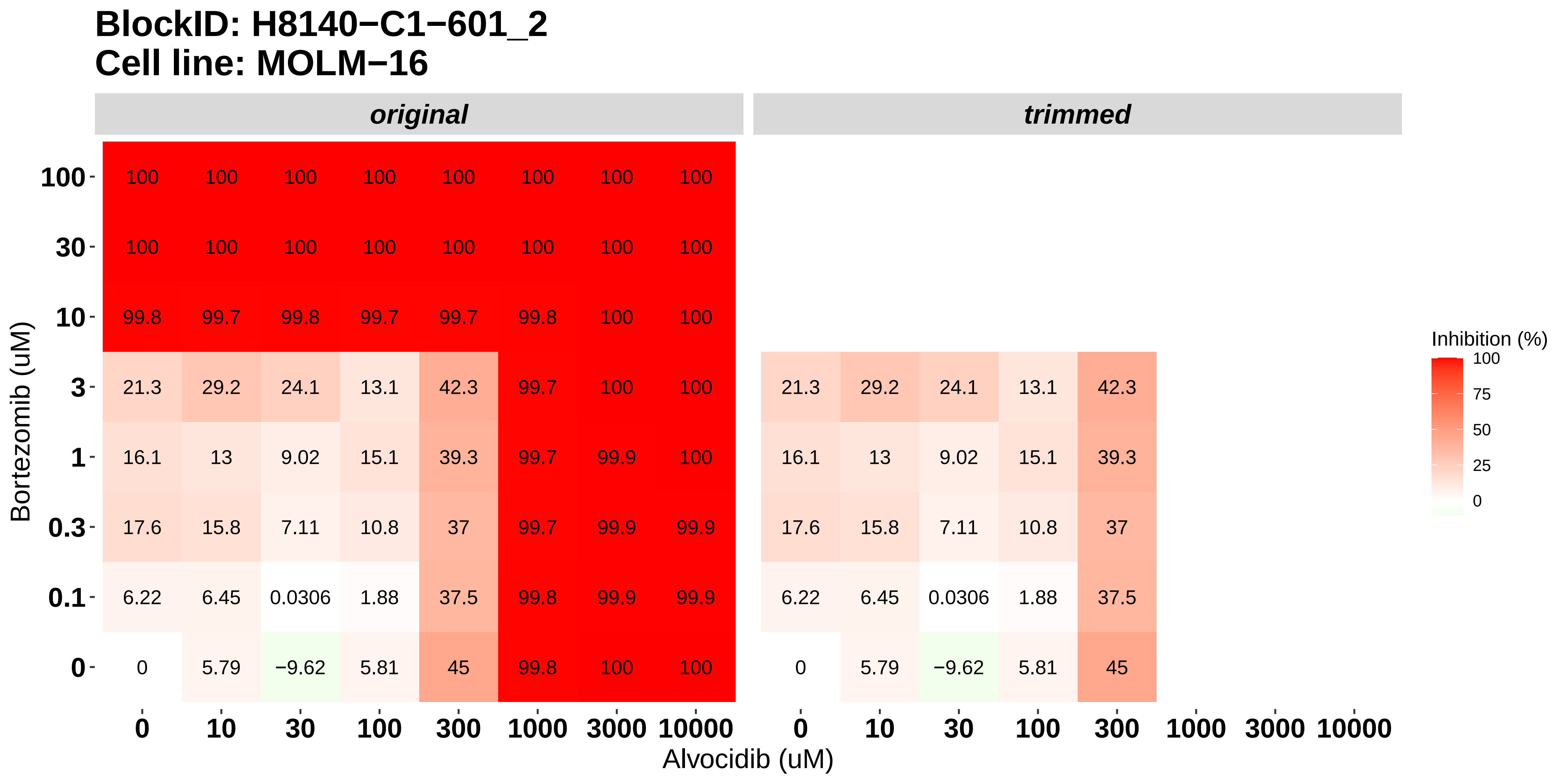

### H8140-C1-601_3_AT7519_Panobinostat_MOLM-16.pdf

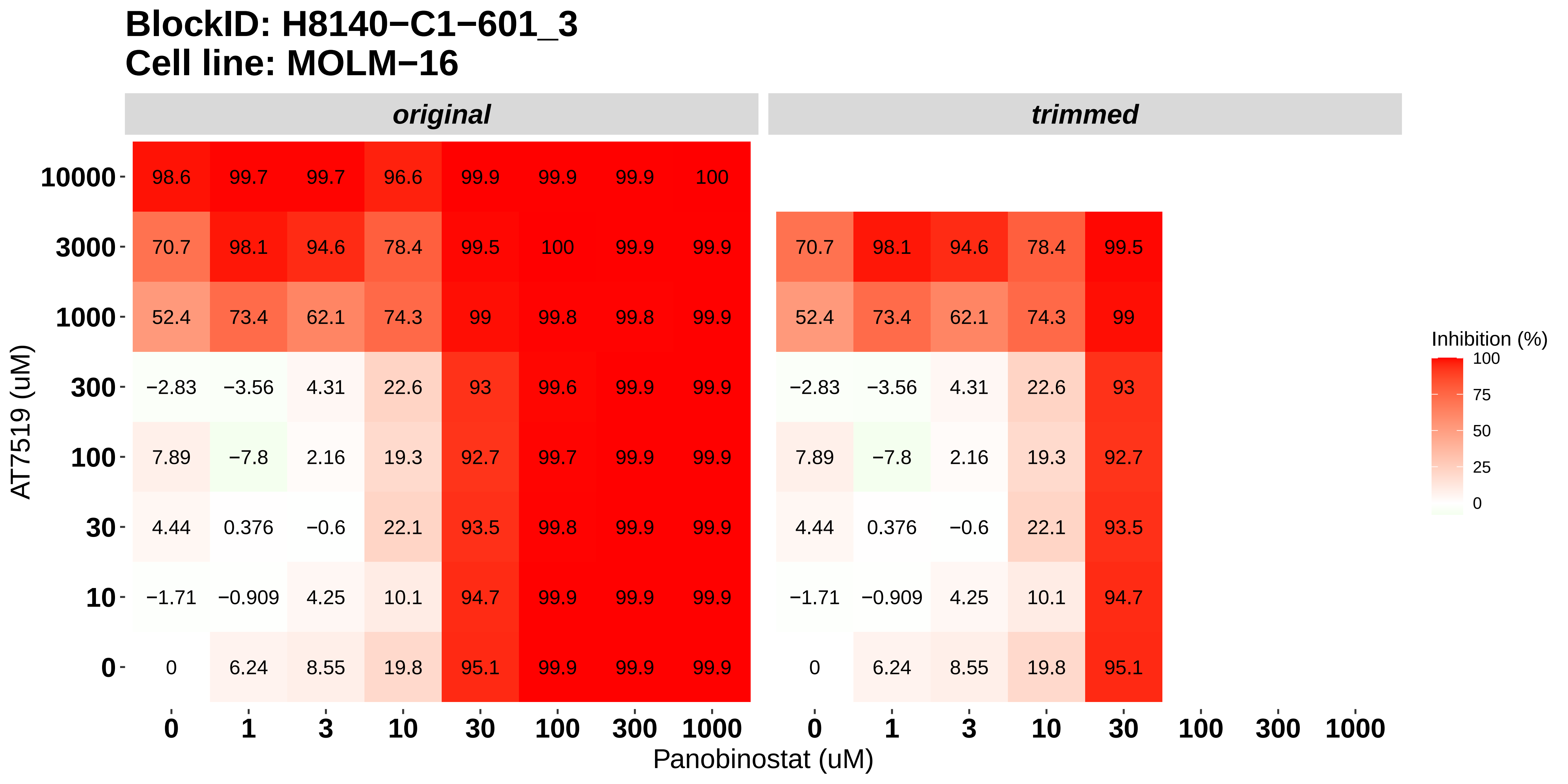

### H8140-C1-601_4_Bortezomib_Panobinostat_MOLM-16.pdf

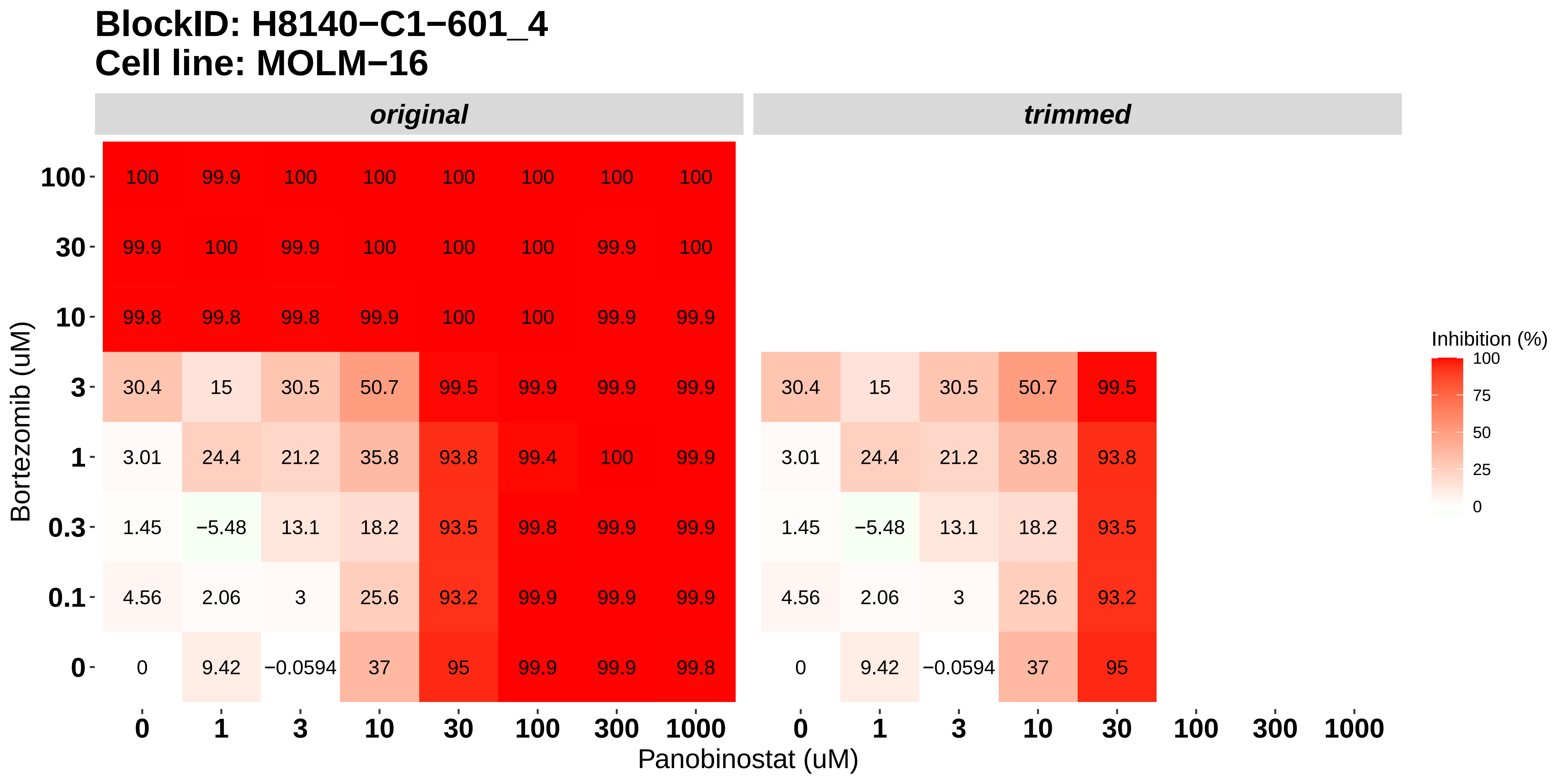

### H8140-C1-601_5_Bortezomib_AT7519_MOLM-16.pdf

#### BlockID: H8140−C1−601_5
#### Cell line: MOLM−16

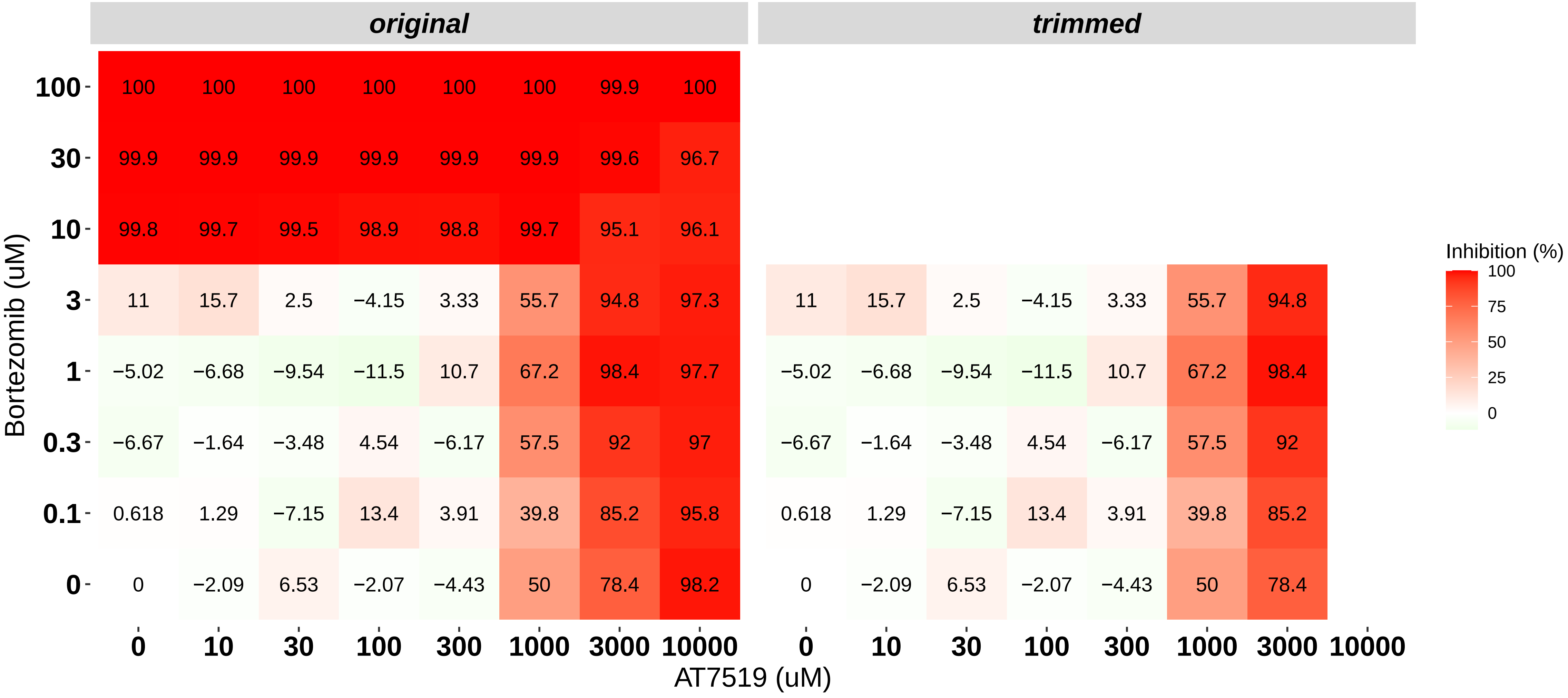

### H8140-C1-601_6_Nintedanib_Dovitinib_MOLM-16.pdf

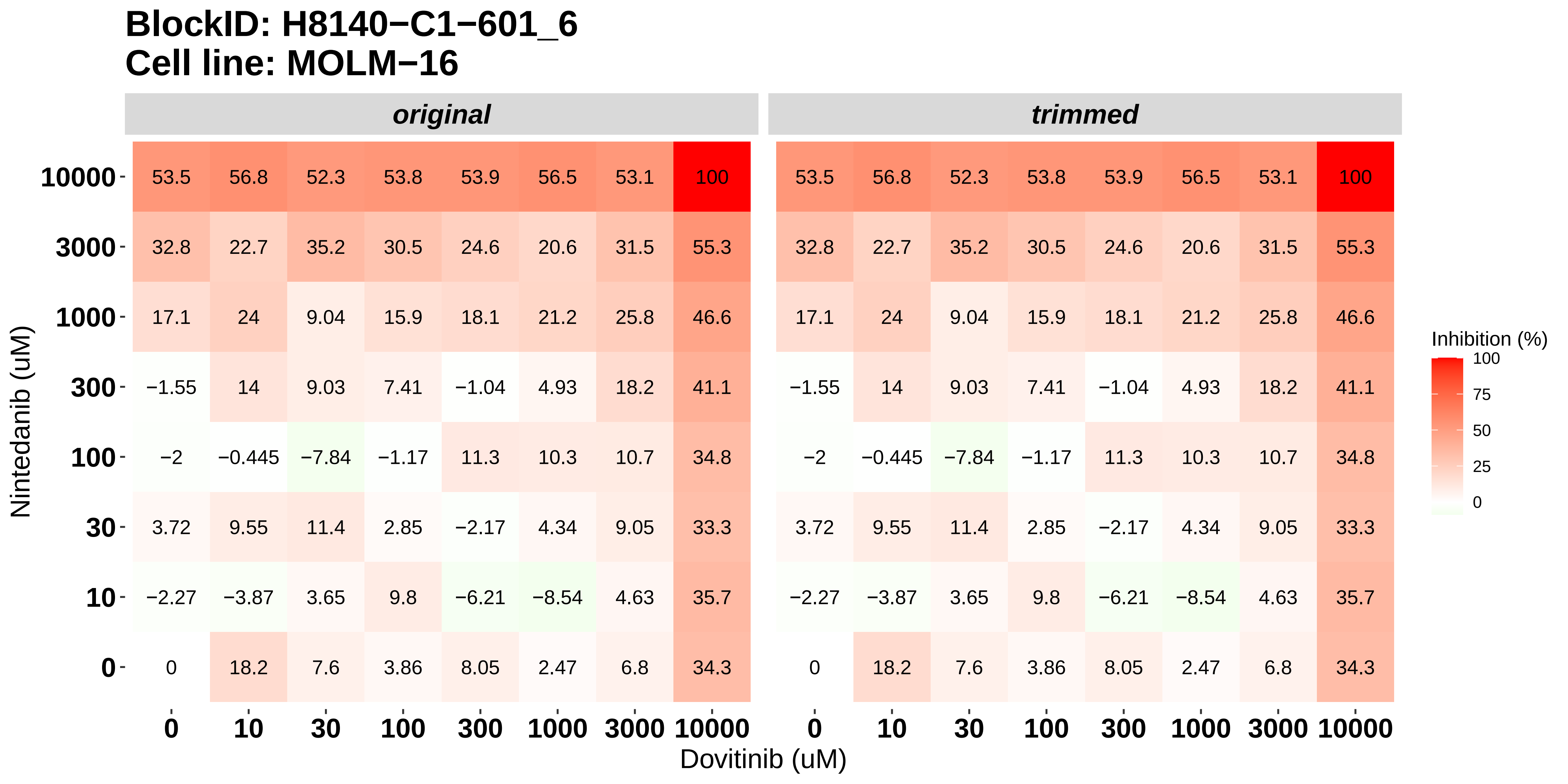

### H8140-C1-602_1_AT7519_Alvocidib_NOMO-1.pdf

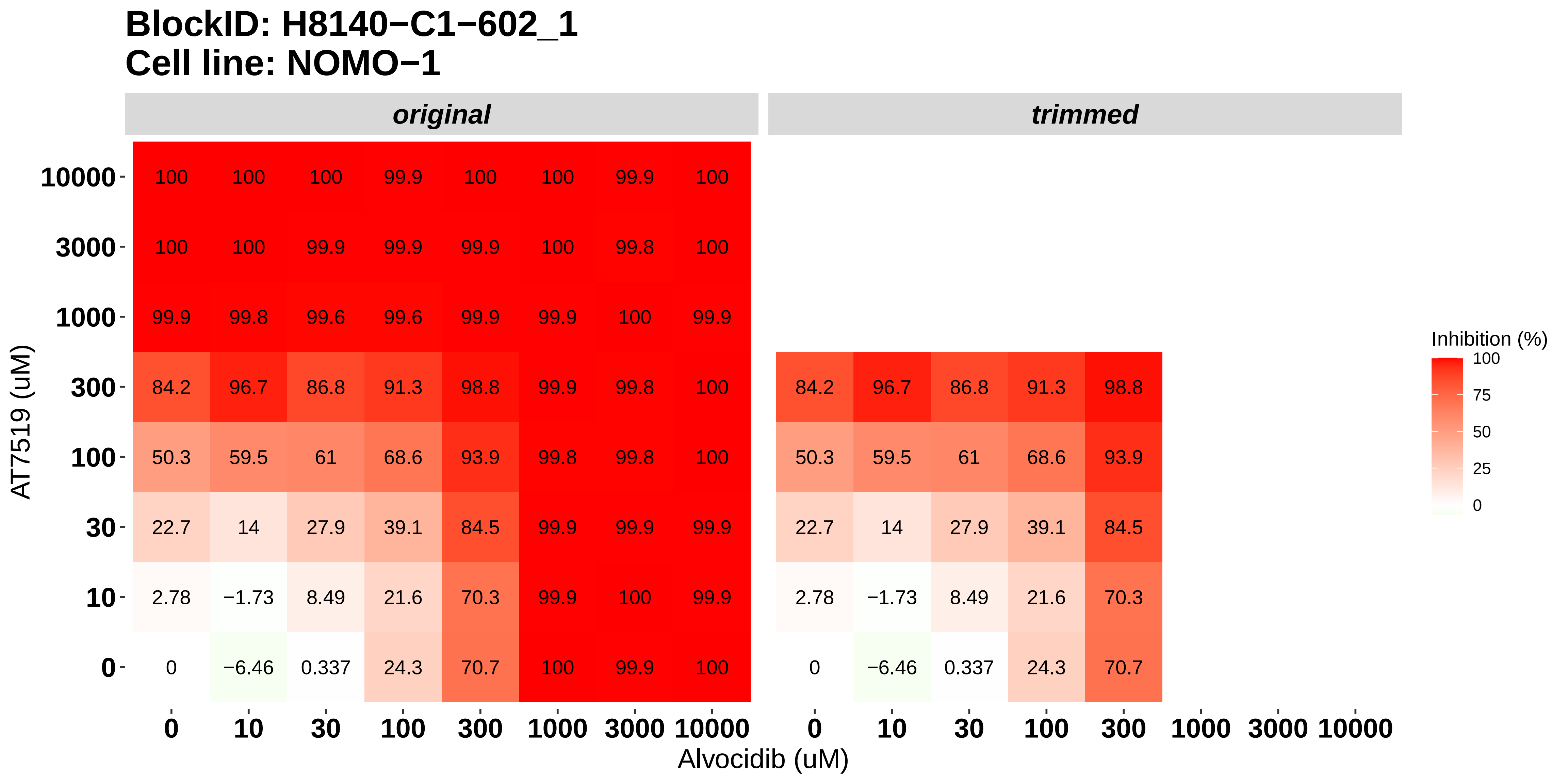

### H8140-C1-602_2_Bortezomib_Alvocidib_NOMO-1.pdf

#### BlockID: H8140−C1−602_2
#### Cell line: NOMO−1

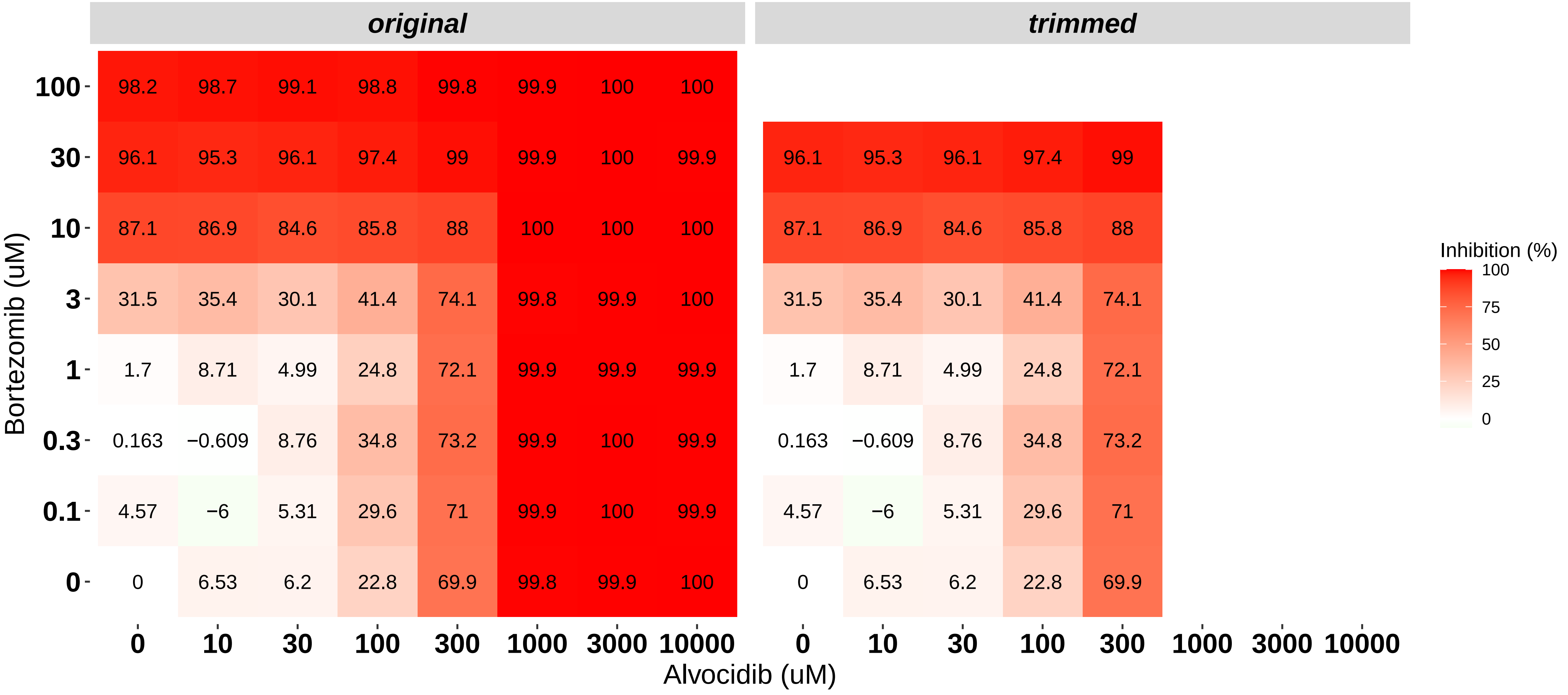

### H8140-C1-602_3_AT7519_Panobinostat_NOMO-1.pdf

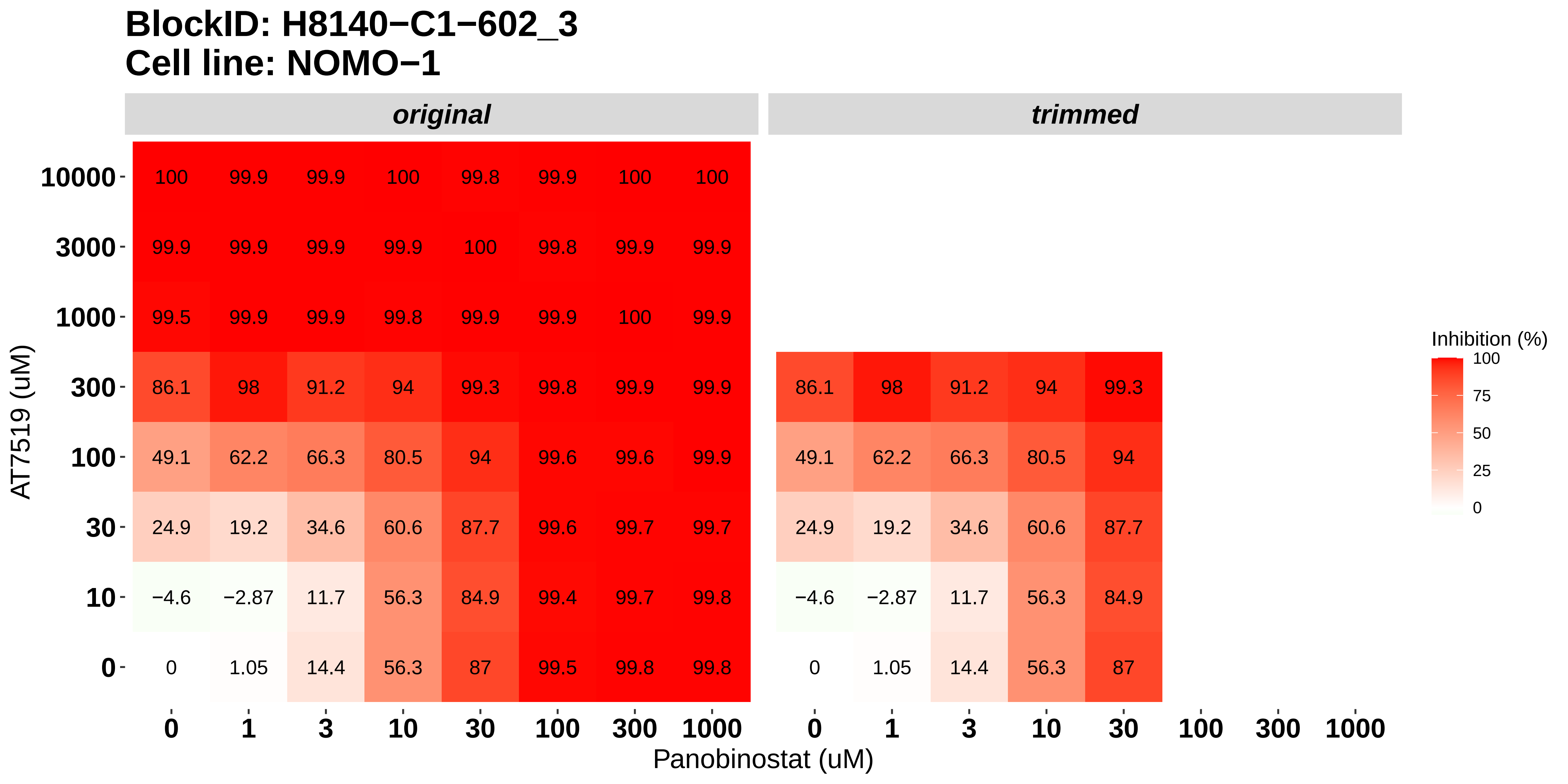

### H8140-C1-602_4_Bortezomib_Panobinostat_NOMO-1.pdf

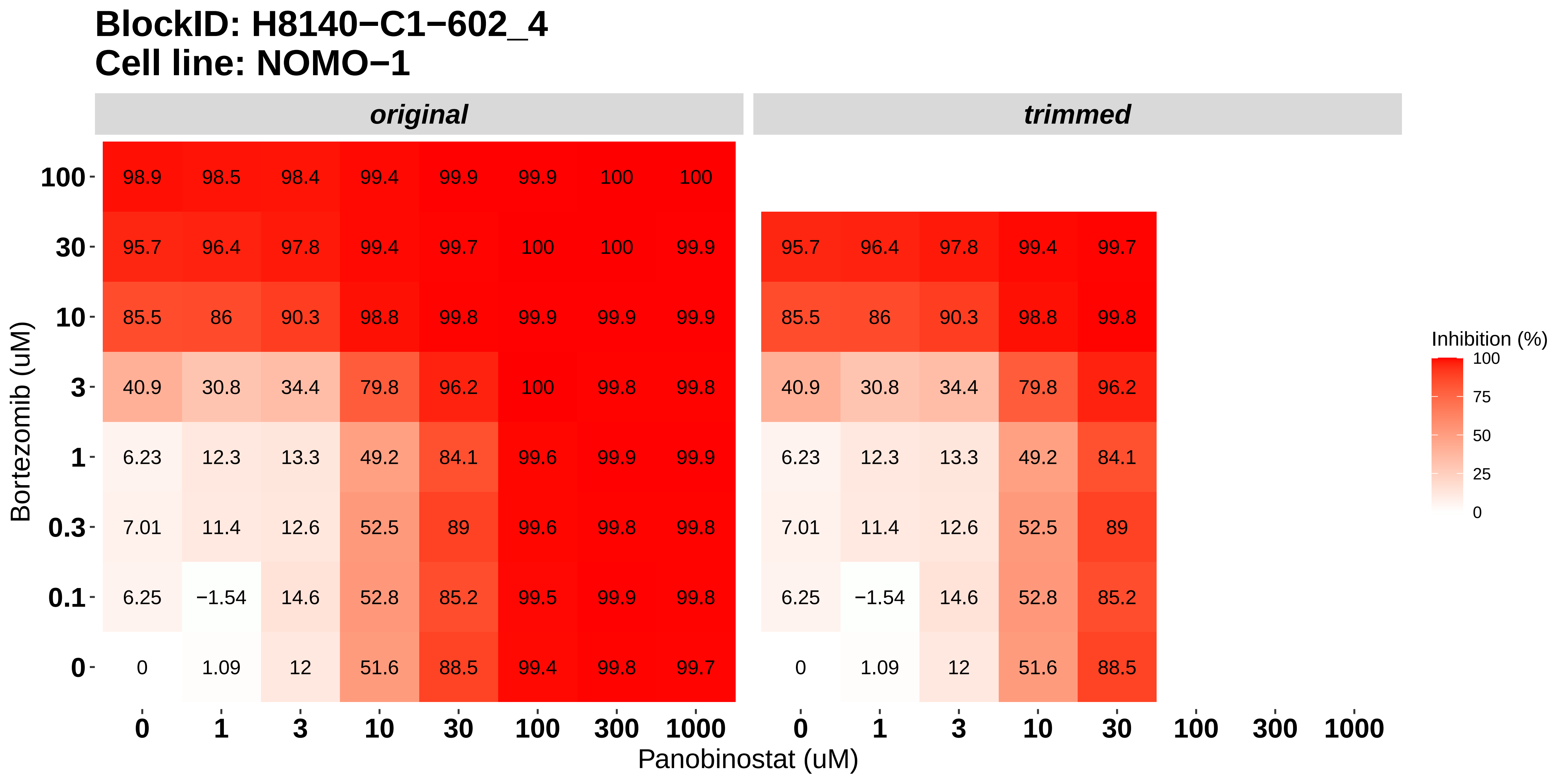

### H8140-C1-602_5_Bortezomib_AT7519_NOMO-1.pdf

**BlockID: H8140−C1−602_5**
**Cell line: NOMO−1**

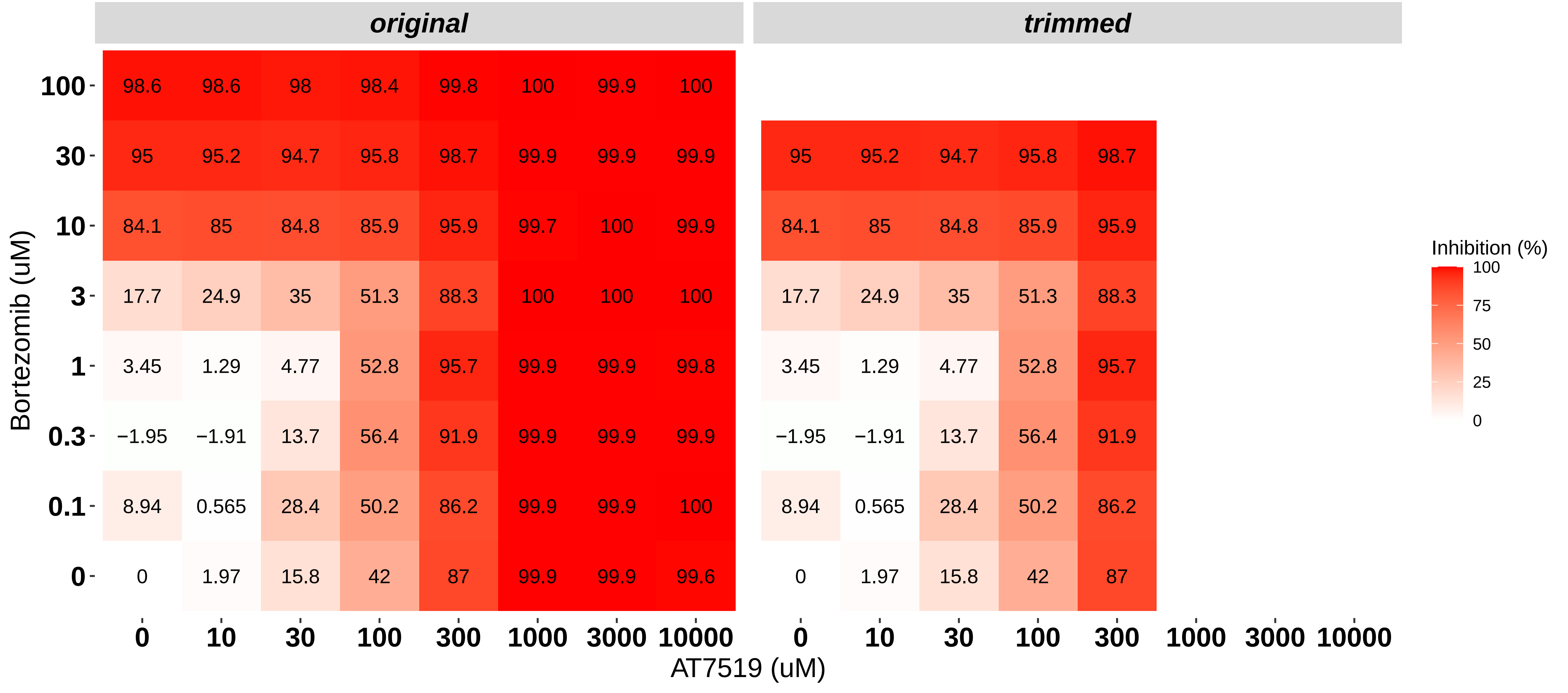

### H8140-C1-602_6_Nintedanib_Dovitinib_NOMO-1.pdf

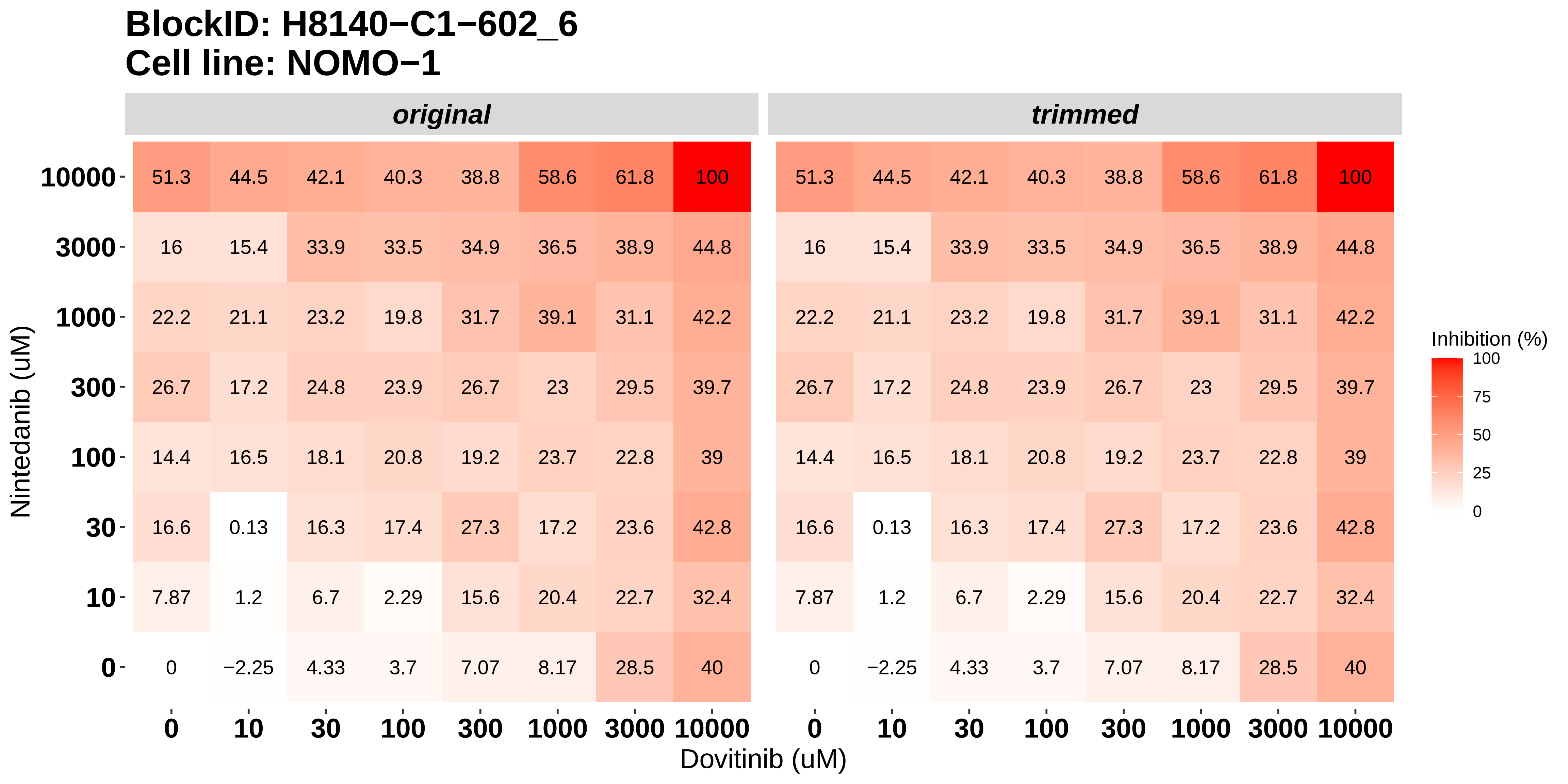

### H8140-C1-603_1_AT7519_Alvocidib_OCI-AML3.pdf

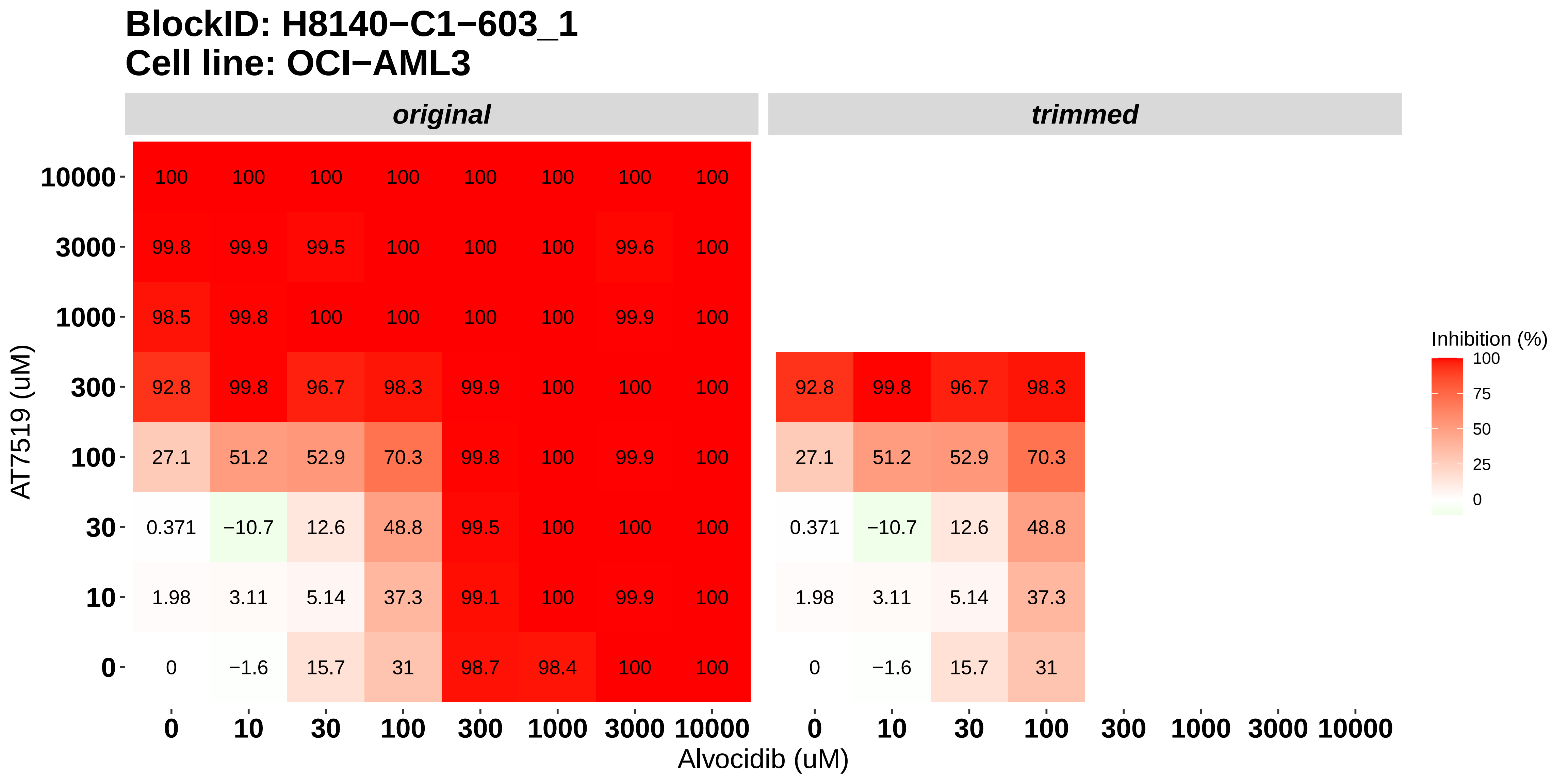

### H8140-C1-603_2_Bortezomib_Alvocidib_OCI-AML3.pdf

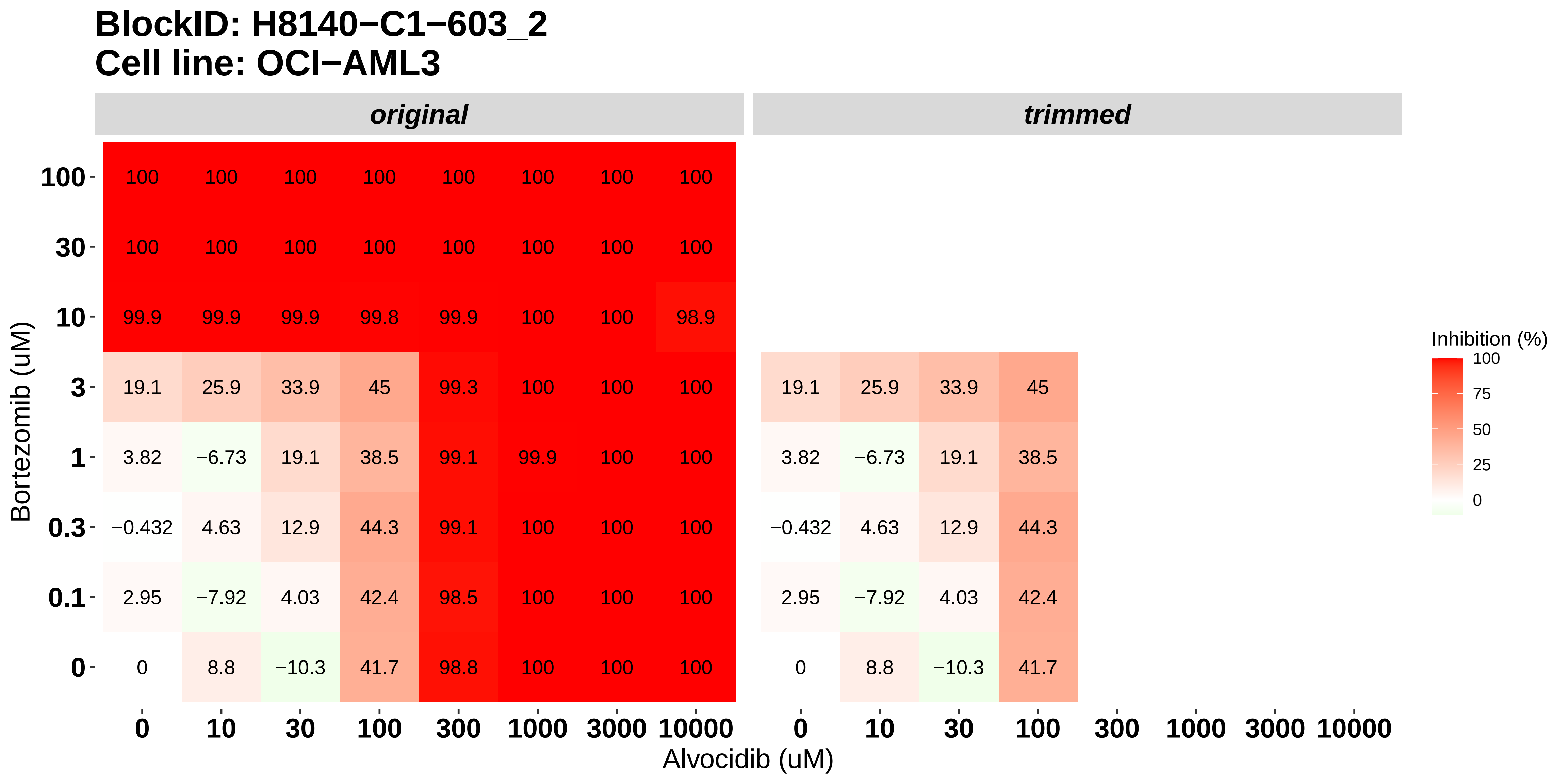

### H8140-C1-603_3_AT7519_Panobinostat_OCI-AML3.pdf

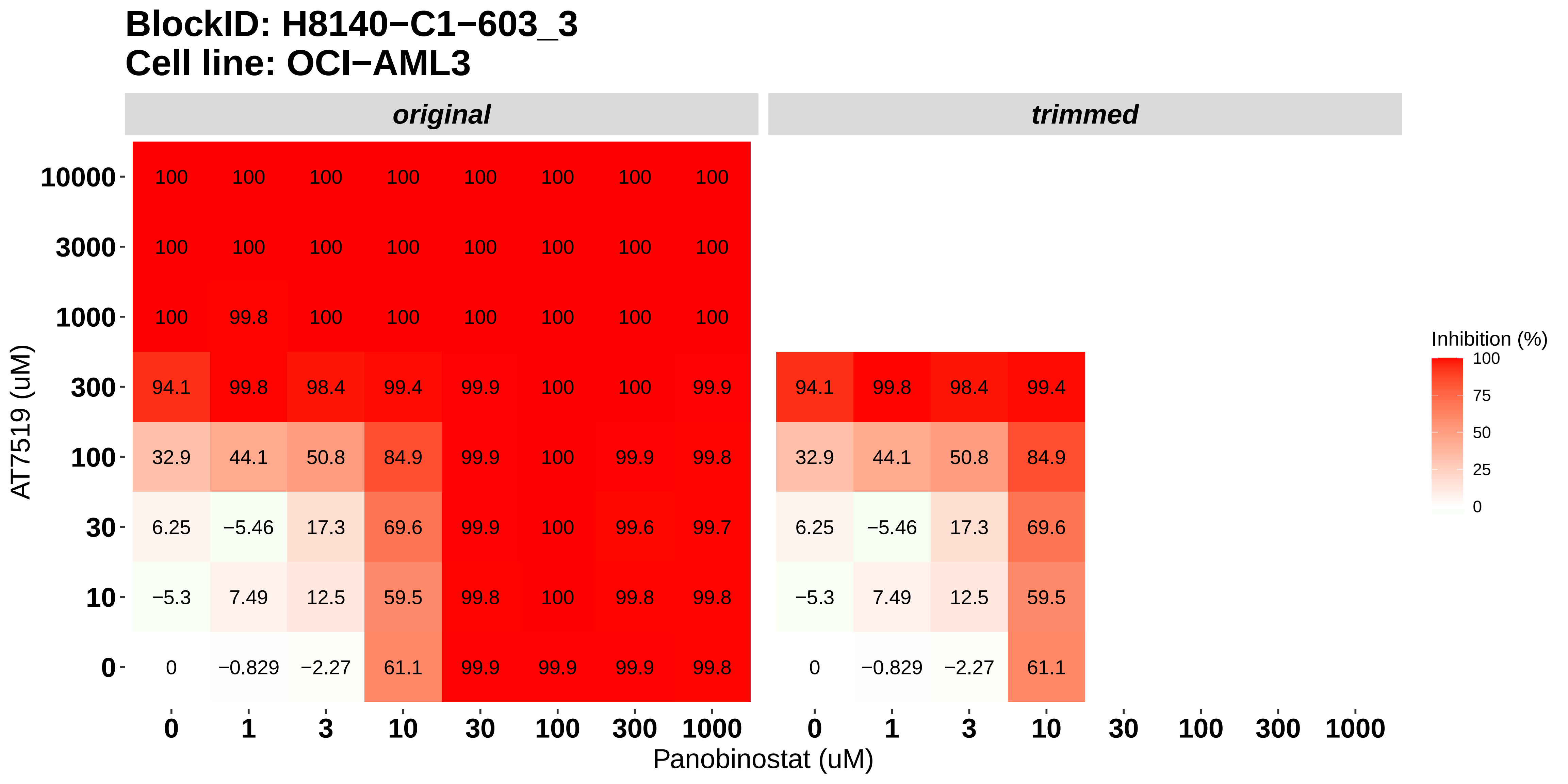

### H8140-C1-603_4_Bortezomib_Panobinostat_OCI-AML3.pdf

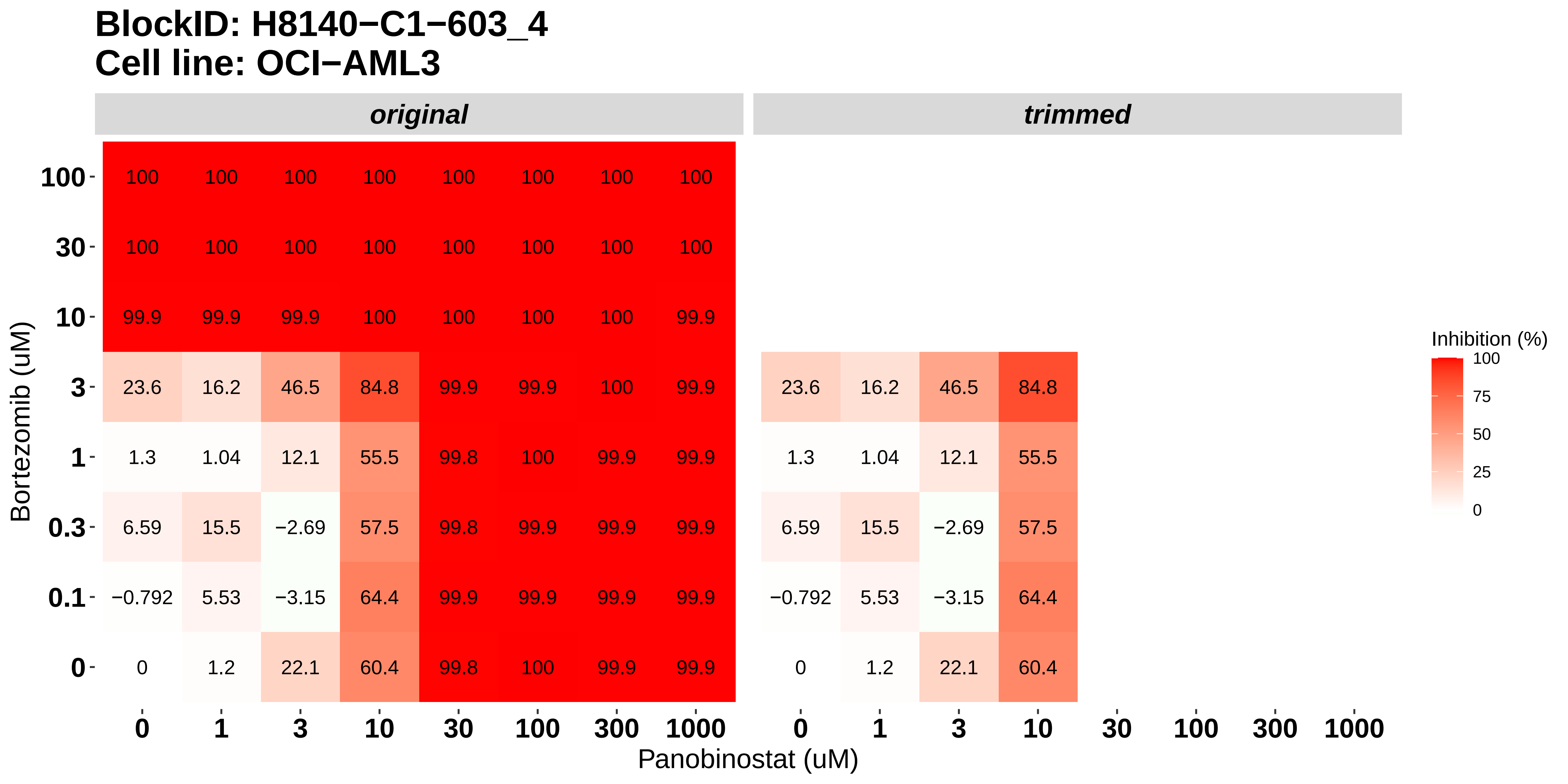

### H8140-C1-603_5_Bortezomib_AT7519_OCI-AML3.pdf

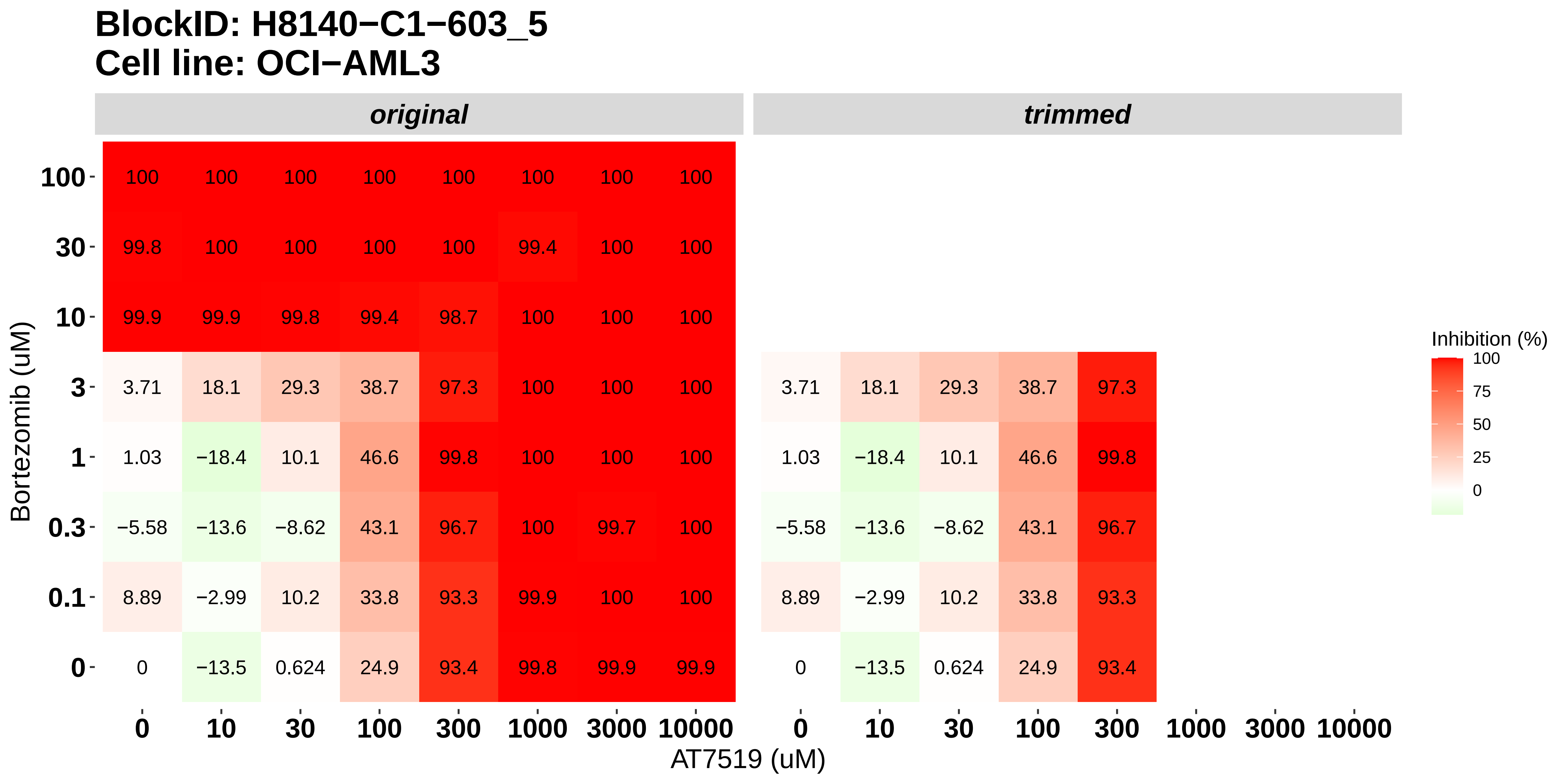

### H8140-C1-603_6_Nintedanib_Dovitinib_OCI-AML3.pdf

#### BlockID: H8140−C1−603_6
#### Cell line: OCI−AML3

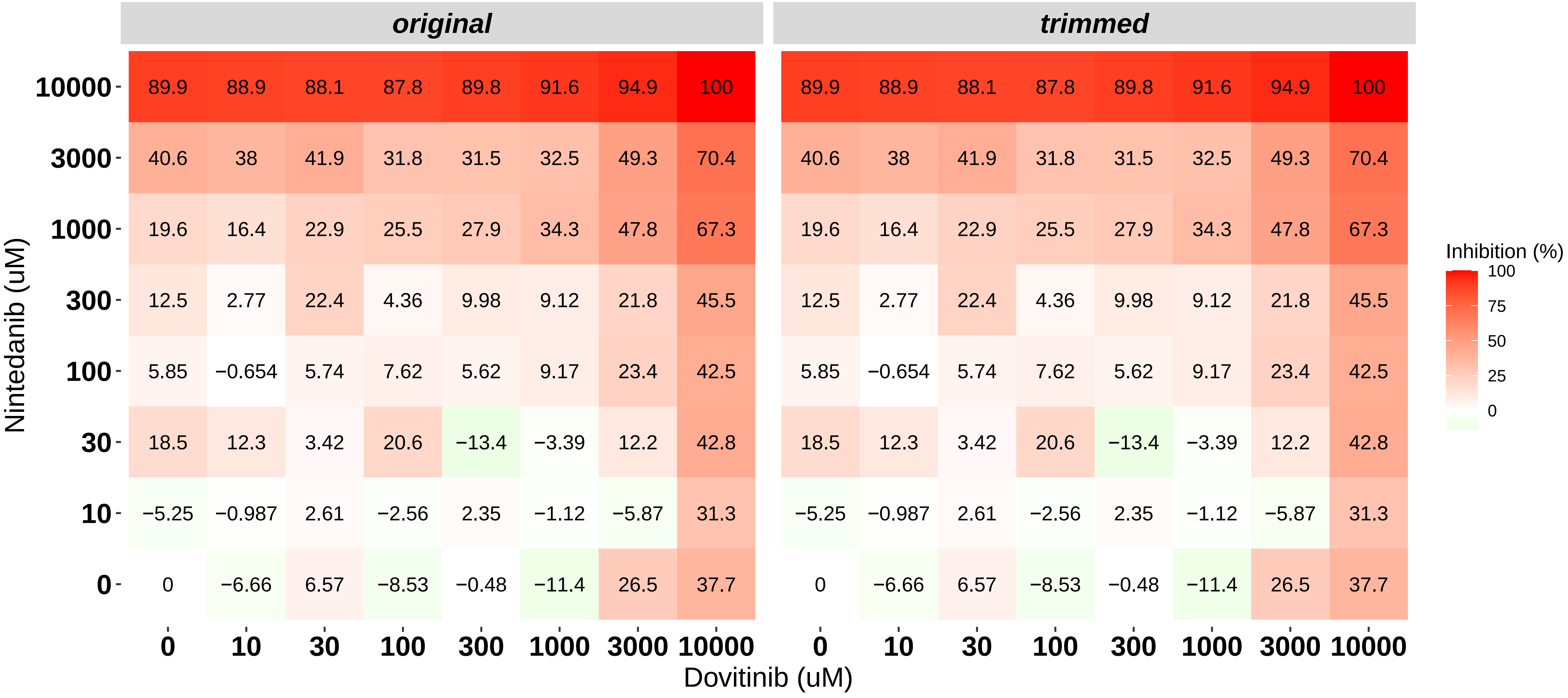

### H8140-C1-701_1_Doramapimod_Dovitinib_MOLM-16.pdf

**BlockID: H8140−C1−701_1**
**Cell line: MOLM−16**

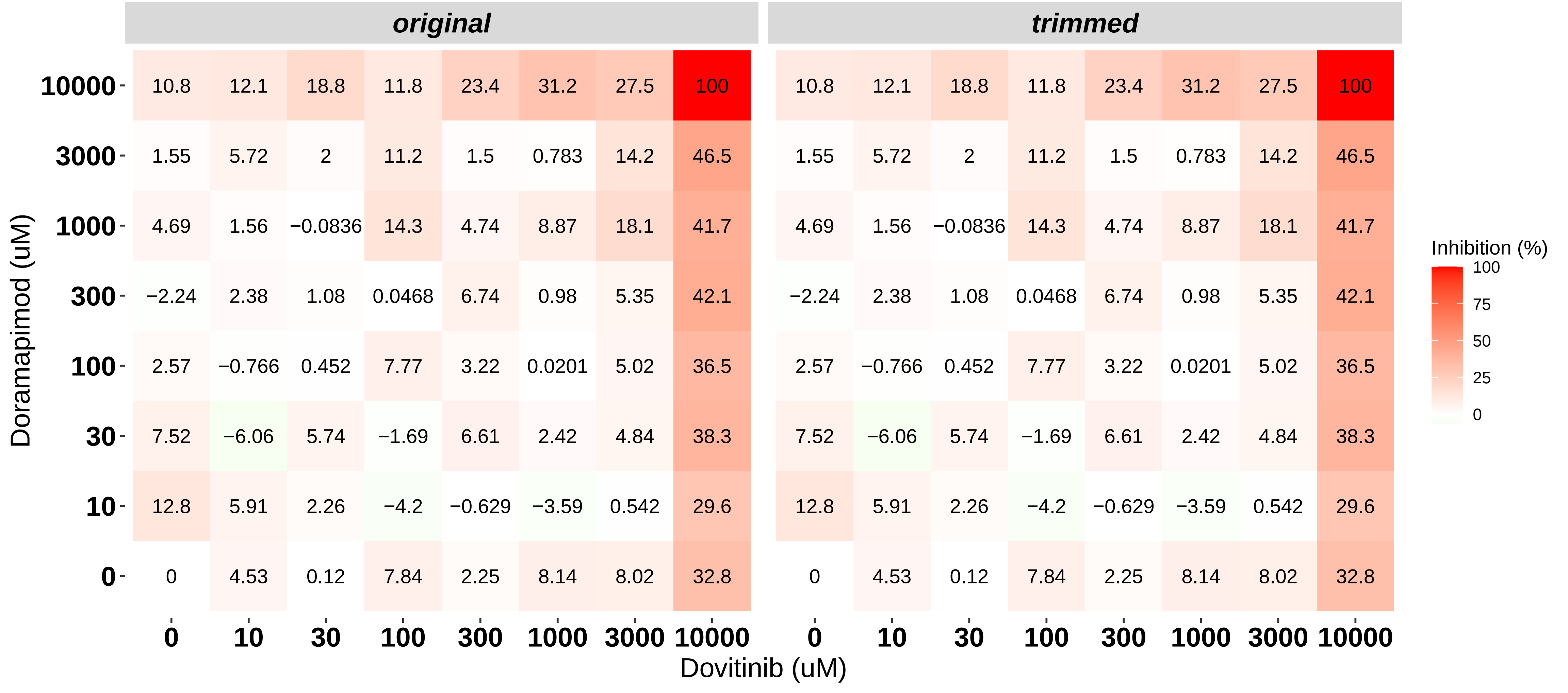

### H8140-C1-701_2_KI20227_Dovitinib_MOLM-16.pdf

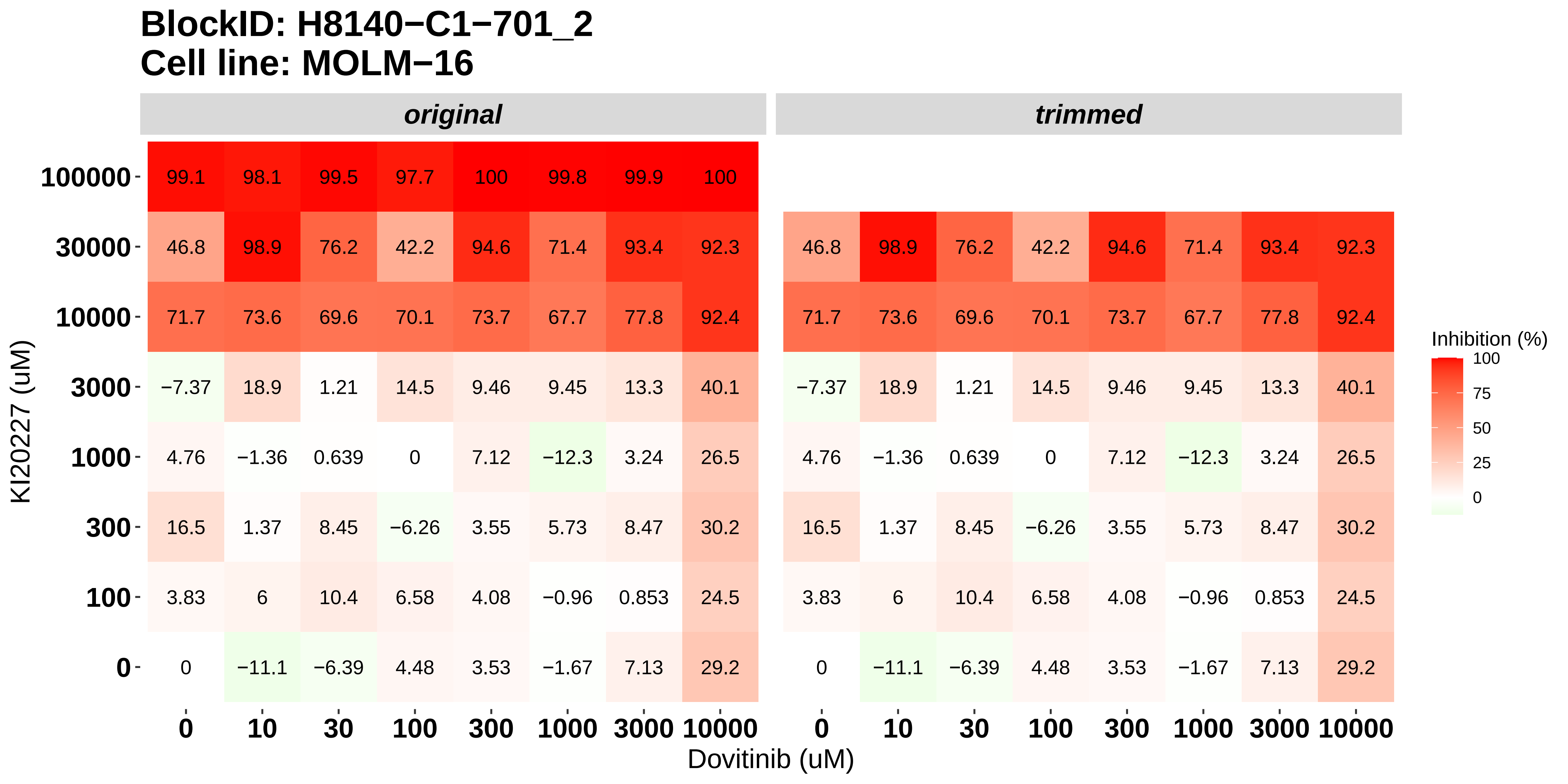

### H8140-C1-701_3_Cabozantinib_Dovitinib_MOLM-16.pdf

#### BlockID: H8140−C1−701_3
#### Cell line: MOLM−16

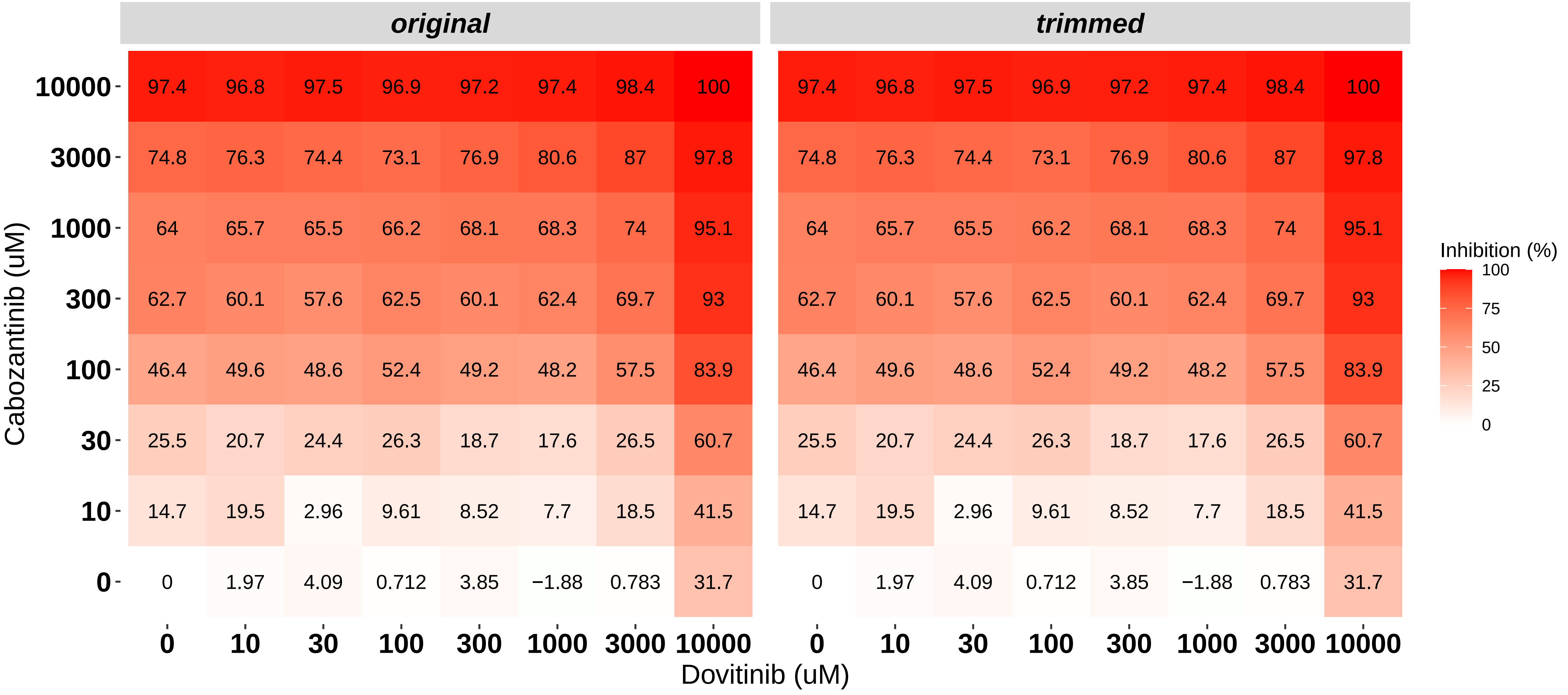

### H8140-C1-701_4_Doramapimod_Nintedanib_MOLM-16.pdf

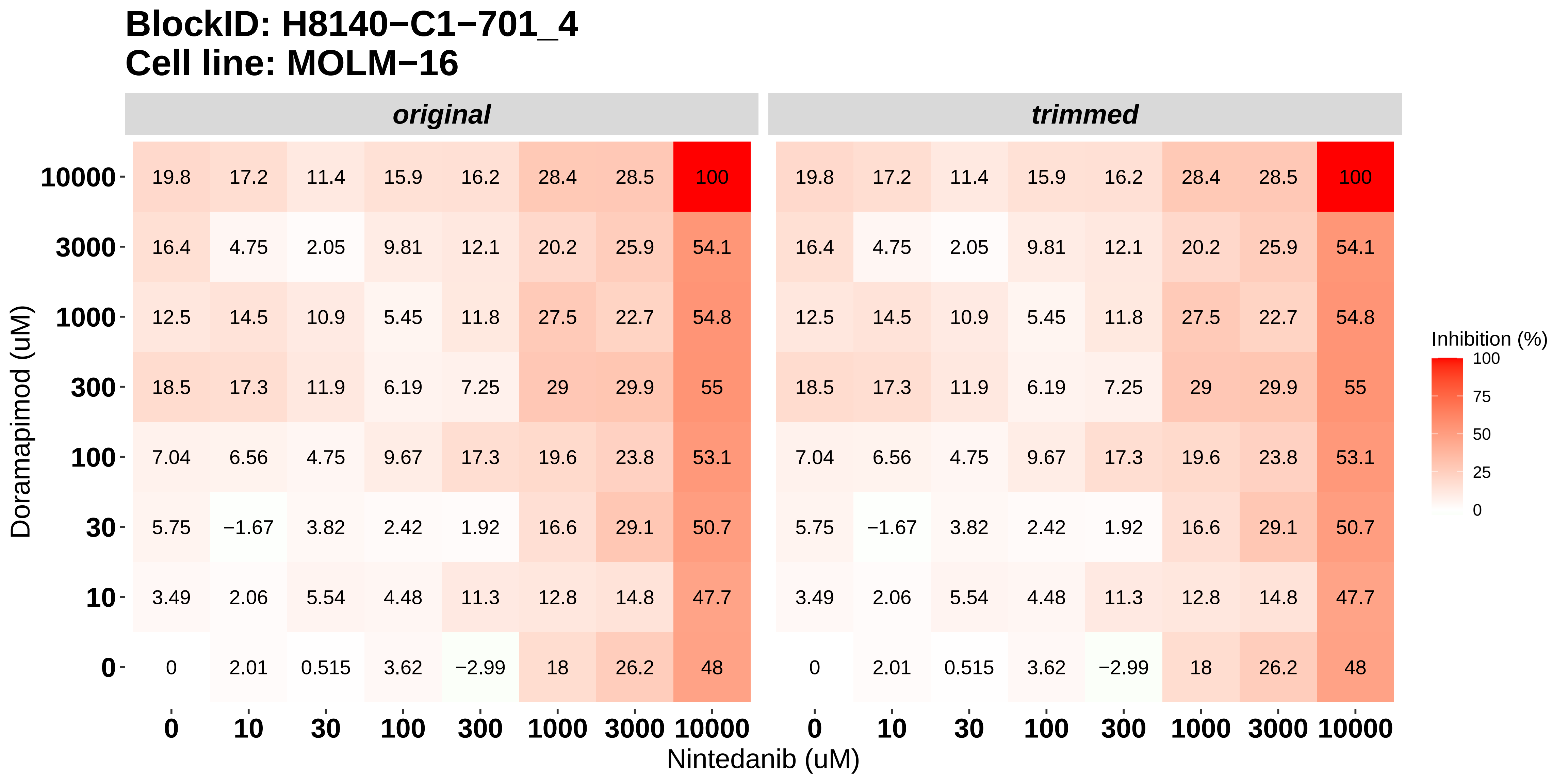

### H8140-C1-701_5_KI20227_Nintedanib_MOLM-16.pdf

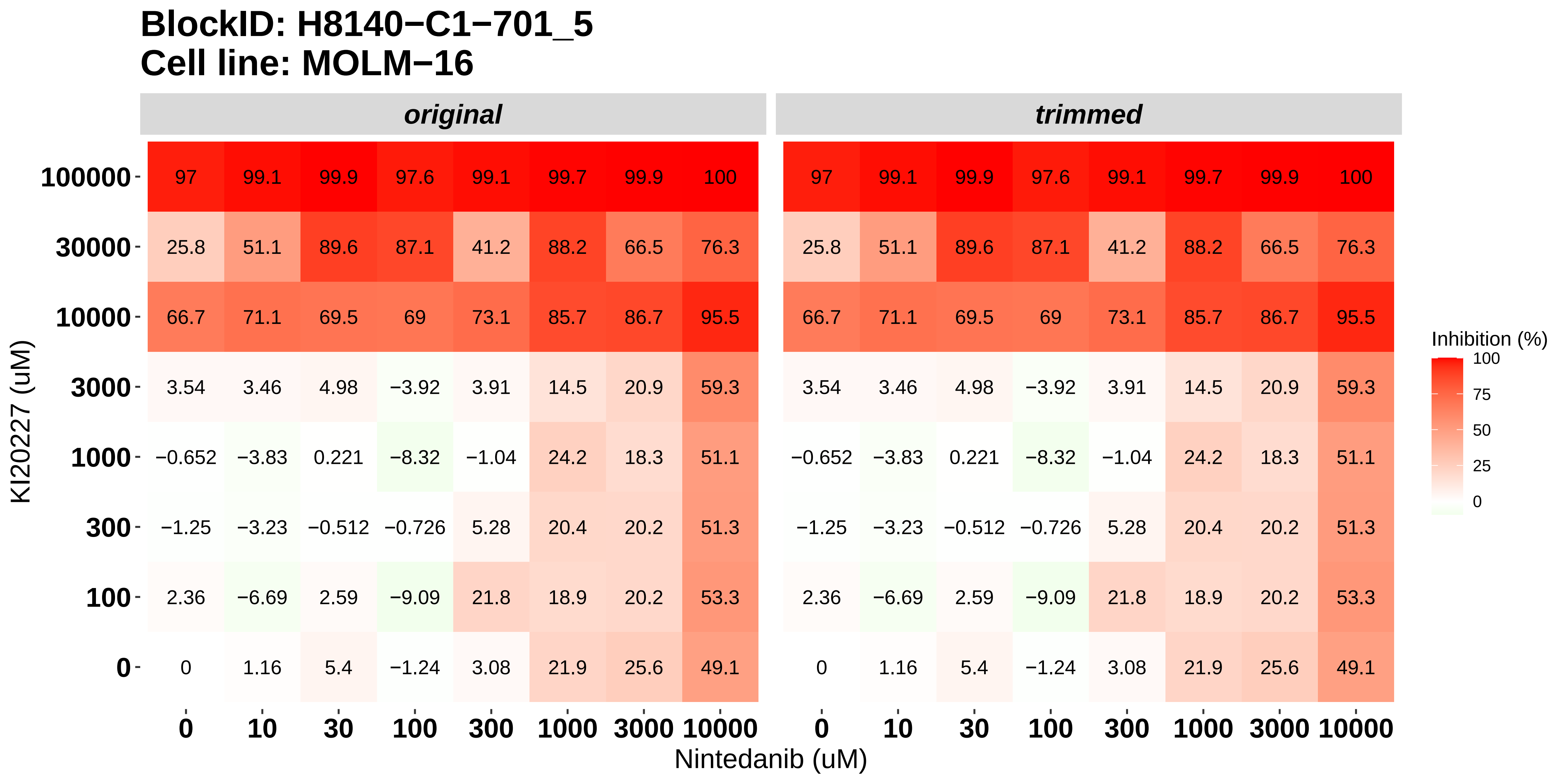

### H8140-C1-701_6_Cabozantinib_Nintedanib_MOLM-16.pdf

#### BlockID: H8140−C1−701_6
#### Cell line: MOLM−16

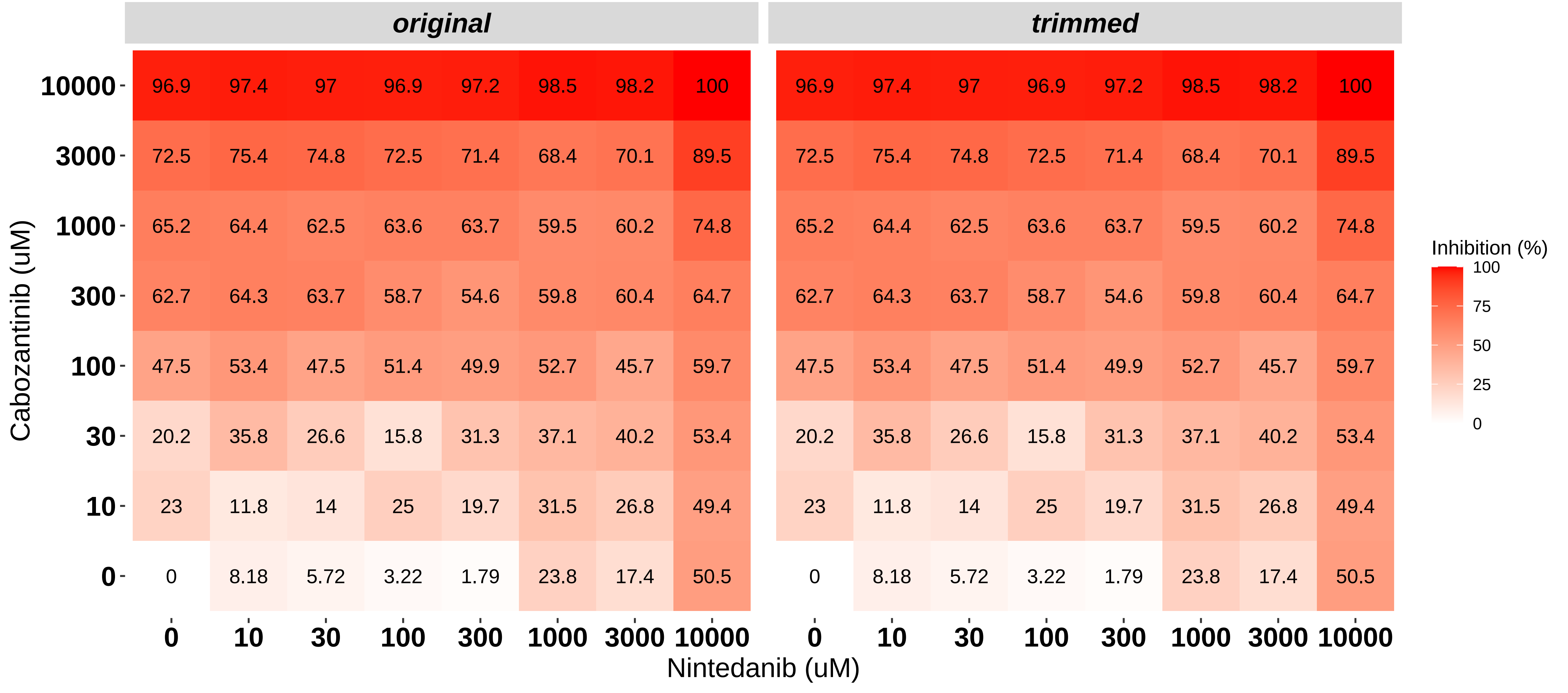

### H8140-C1-702_1_Doramapimod_Dovitinib_NOMO-1.pdf

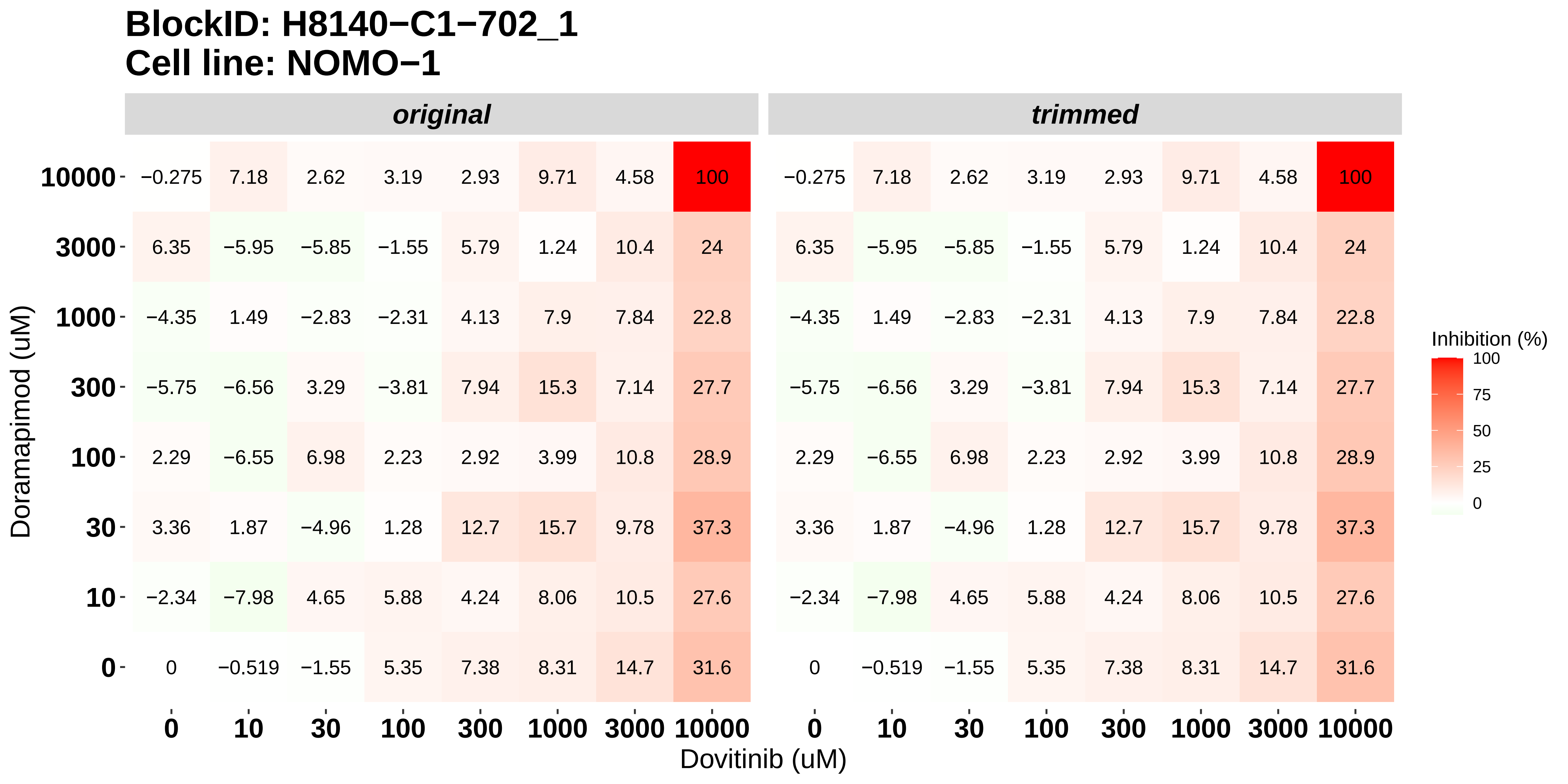

### H8140-C1-702_2_KI20227_Dovitinib_NOMO-1.pdf

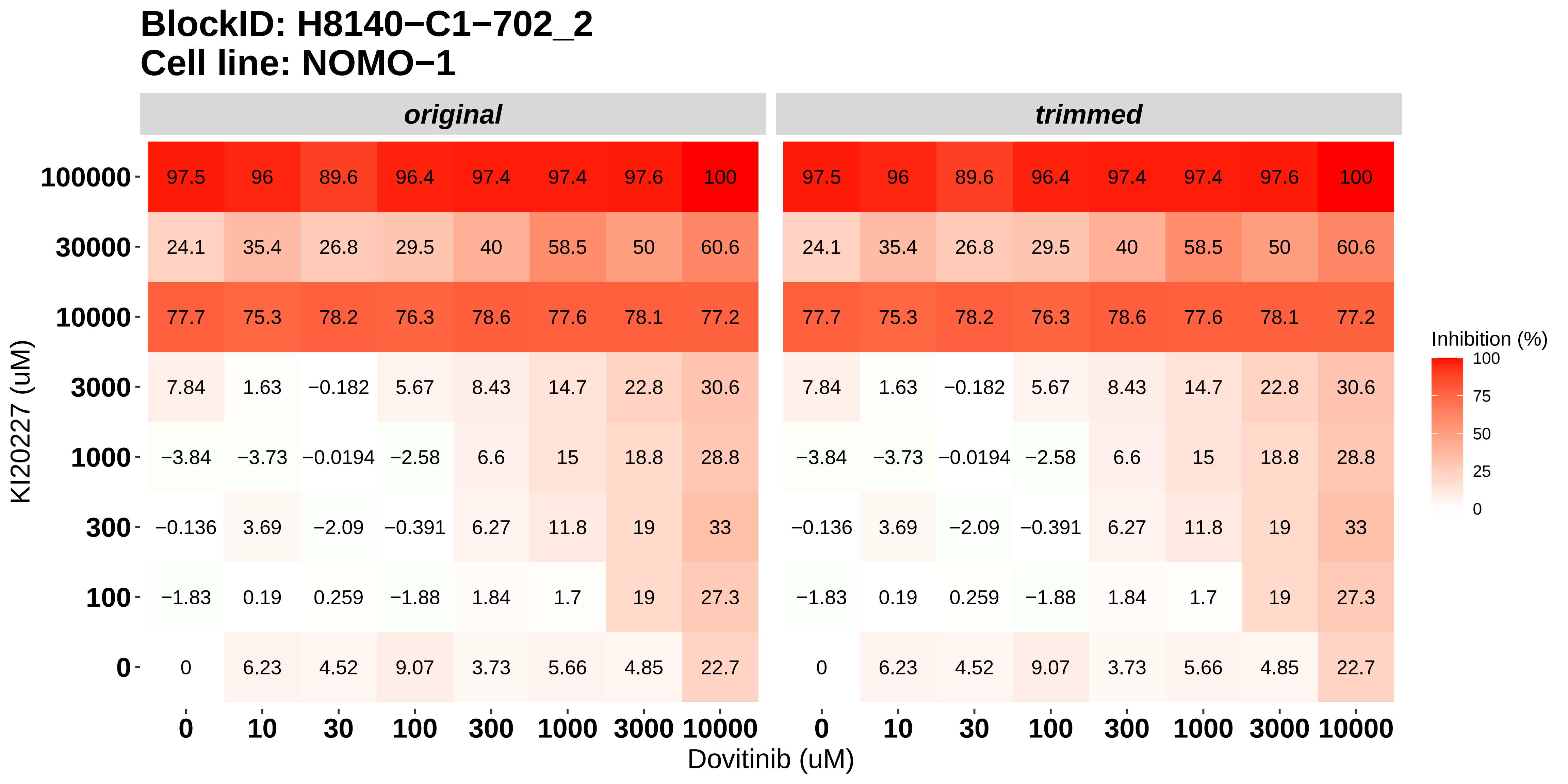

### H8140-C1-702_3_Cabozantinib_Dovitinib_NOMO-1.pdf

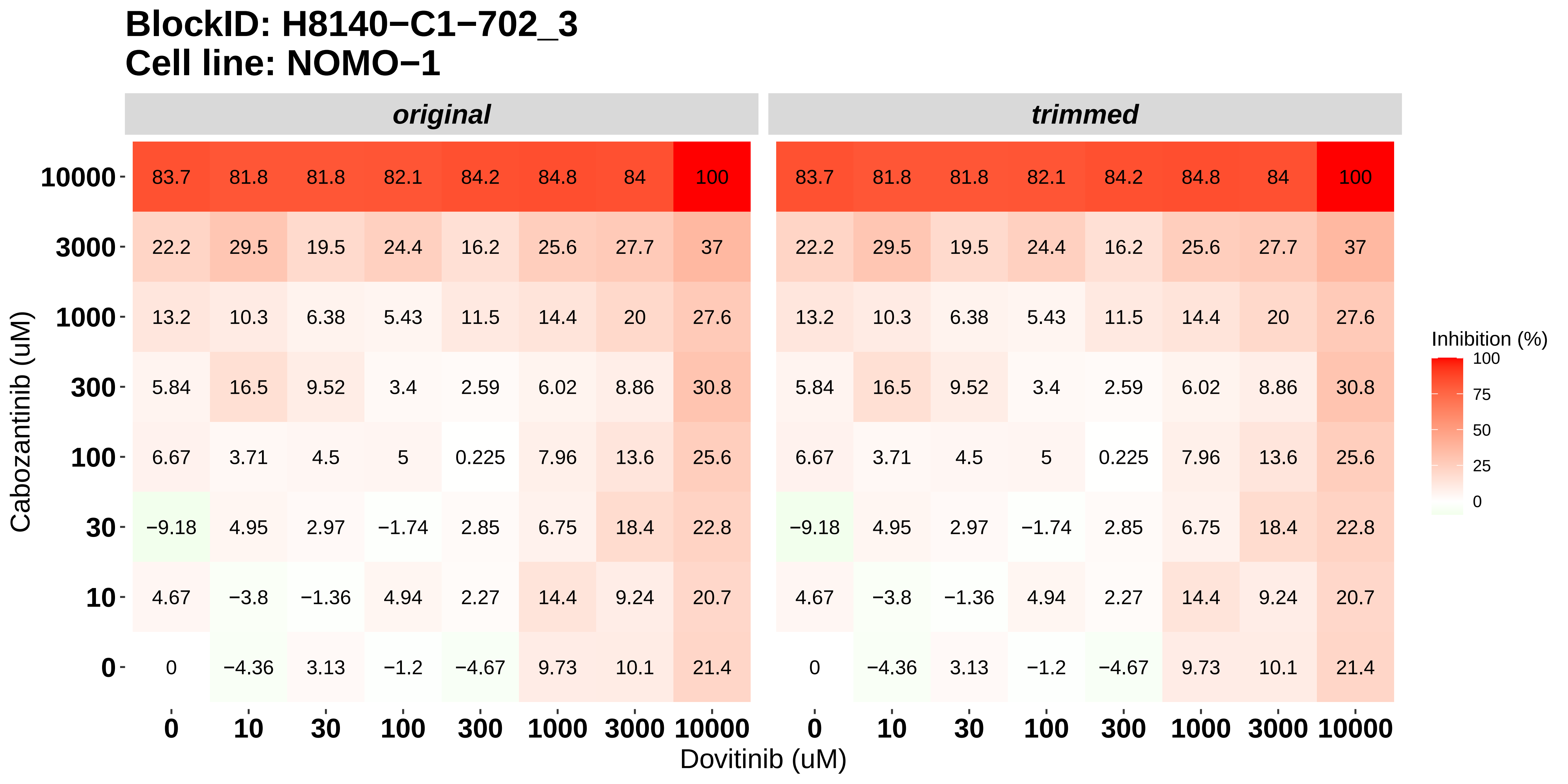

### H8140-C1-702_4_Doramapimod_Nintedanib_NOMO-1.pdf

#### BlockID: H8140−C1−702_4
#### Cell line: NOMO−1

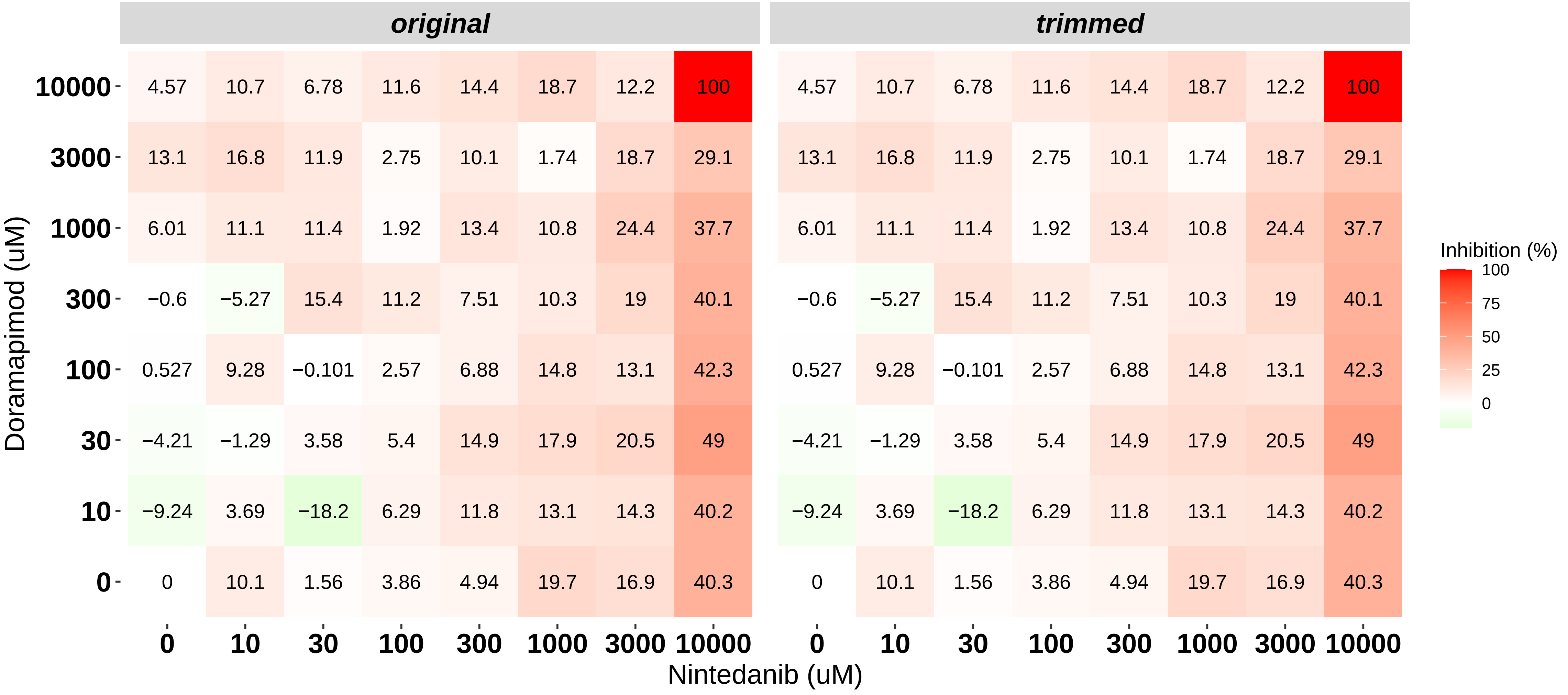

### H8140-C1-702_5_KI20227_Nintedanib_NOMO-1.pdf

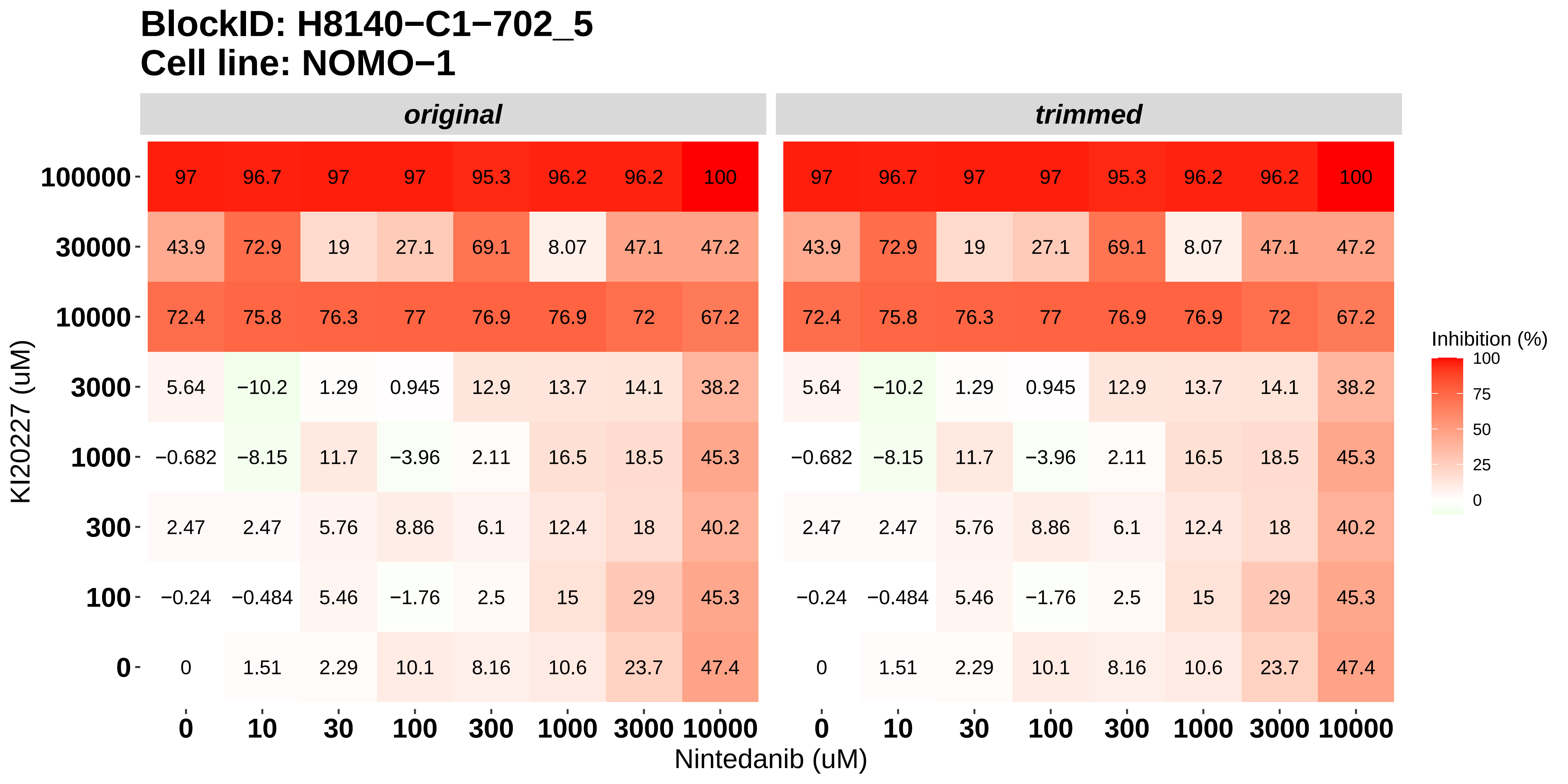

### H8140-C1-702_6_Cabozantinib_Nintedanib_NOMO-1.pdf

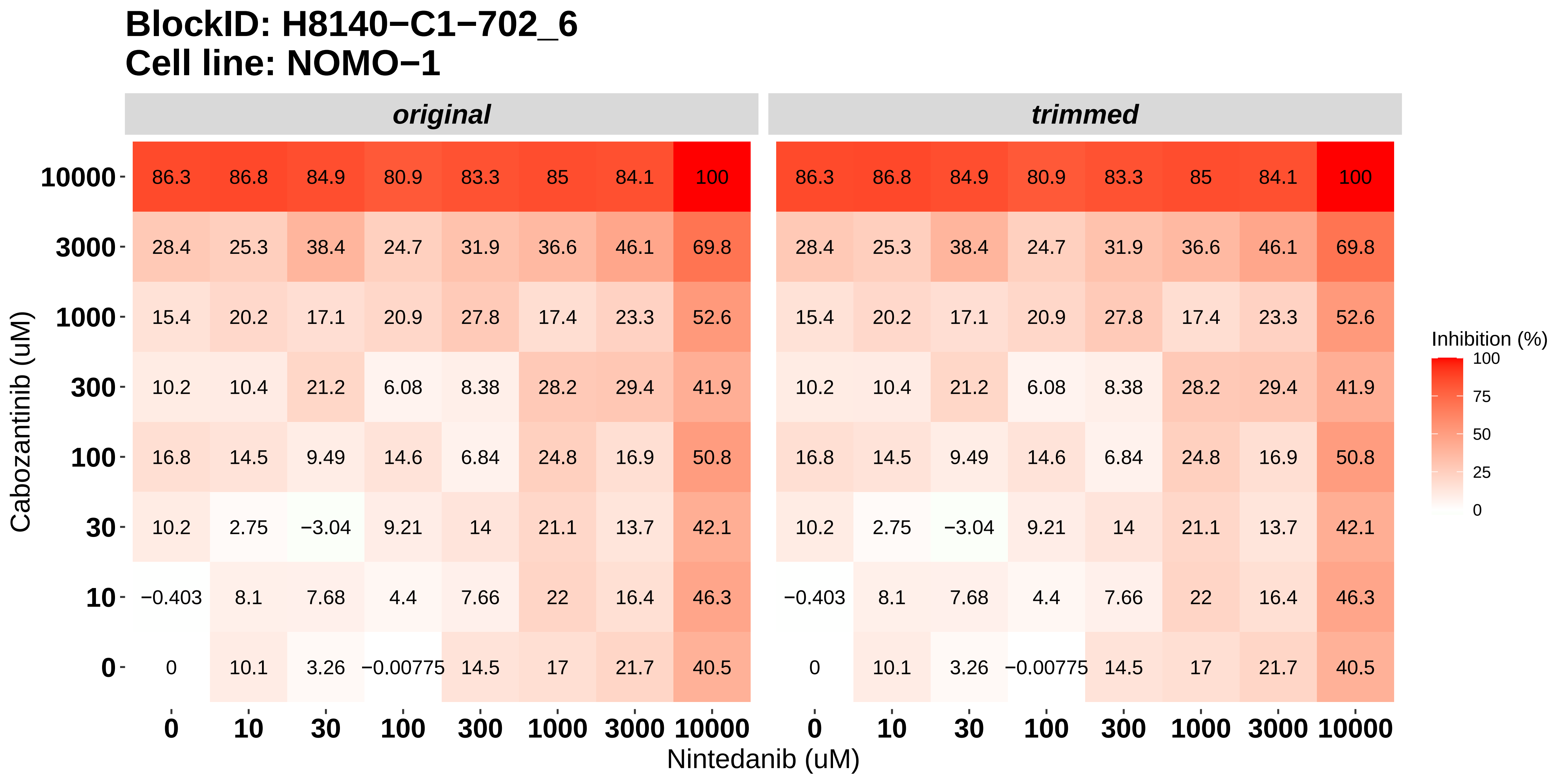

### H8140-C1-703_1_Doramapimod_Dovitinib_OCI-AML3.pdf

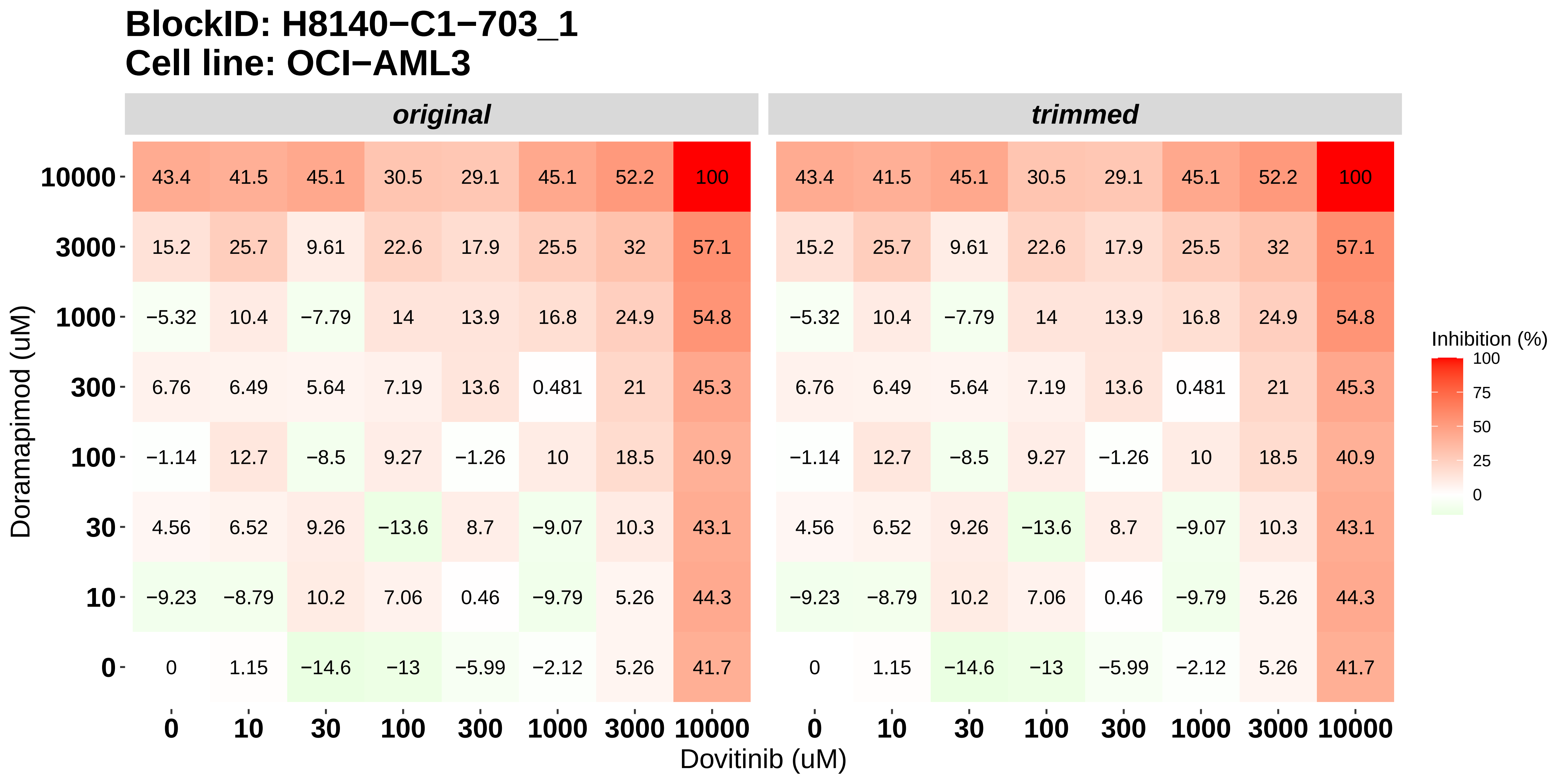

### H8140-C1-703_2_KI20227_Dovitinib_OCI-AML3.pdf

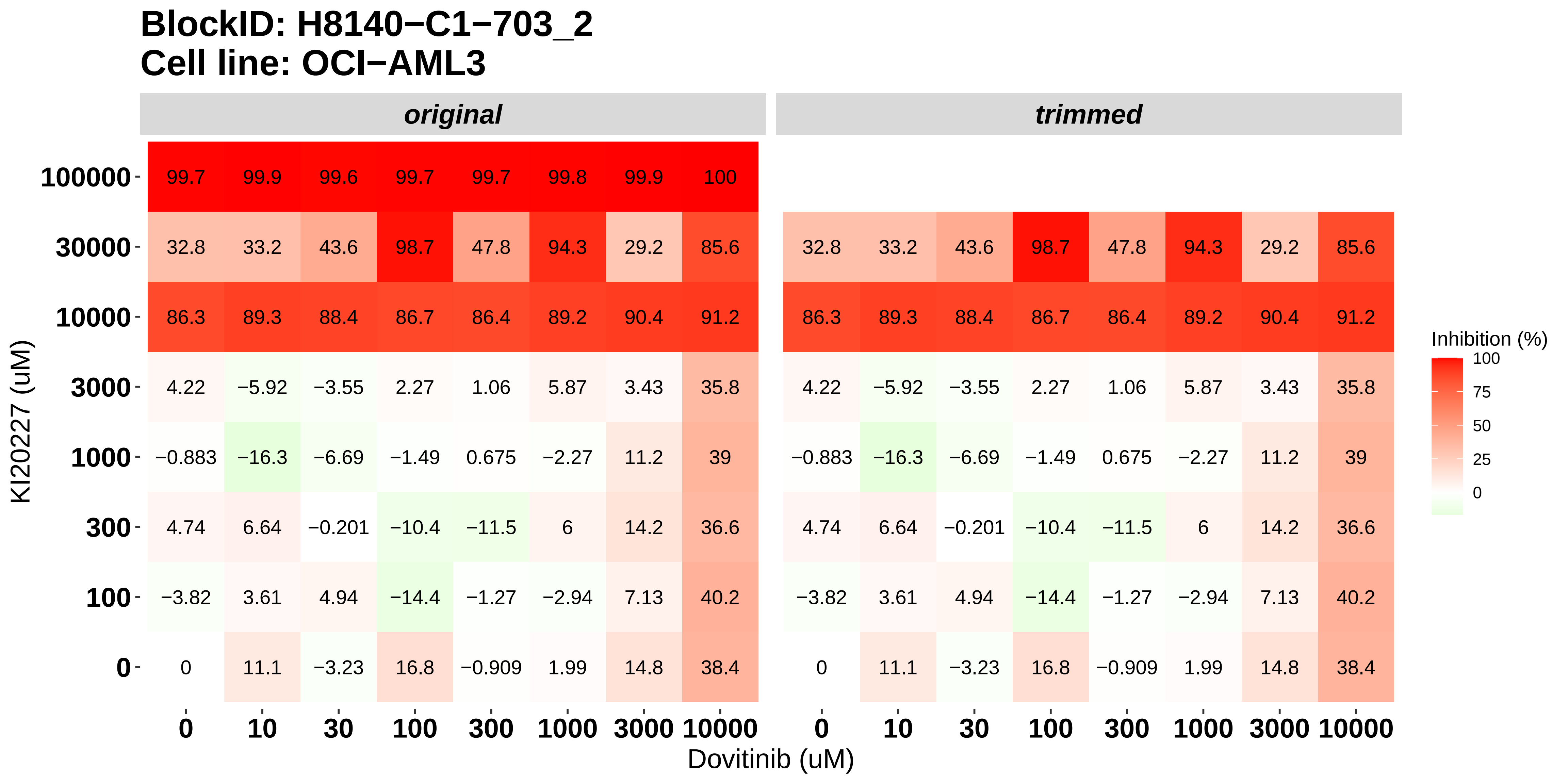

### H8140-C1-703_3_Cabozantinib_Dovitinib_OCI-AML3.pdf

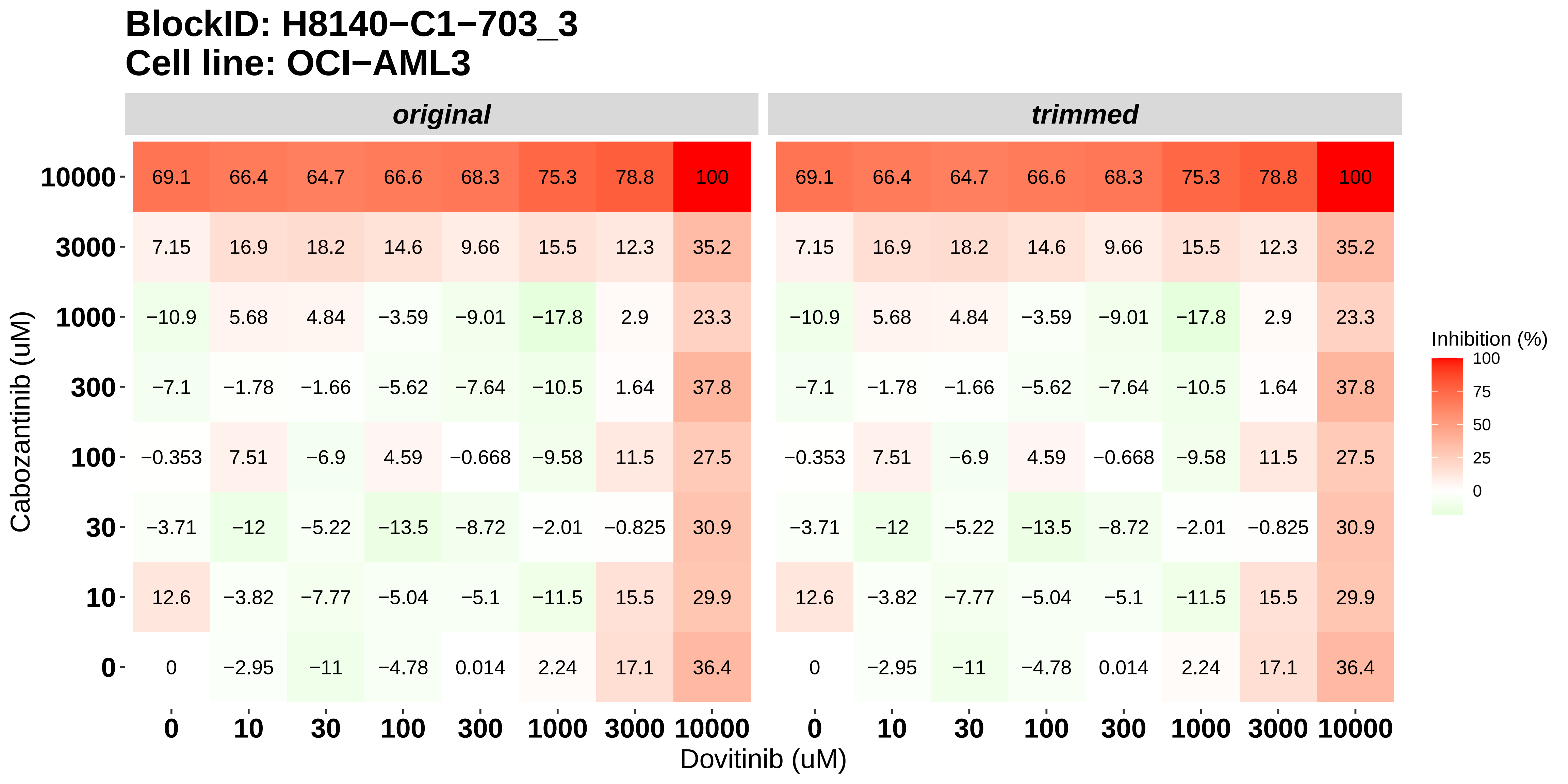

### H8140-C1-703_4_Doramapimod_Nintedanib_OCI-AML3.pdf

#### BlockID: H8140−C1−703_4
#### Cell line: OCI−AML3

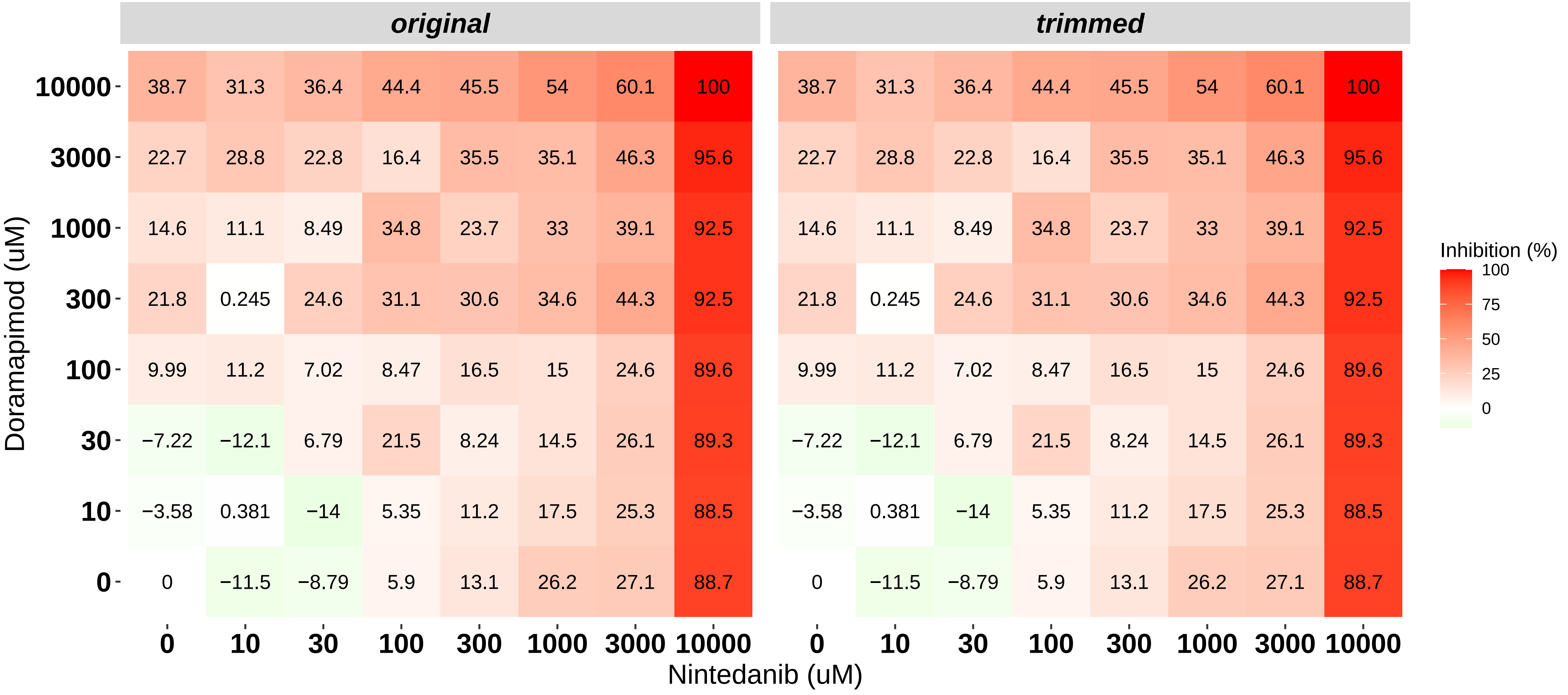

### H8140-C1-703_5_KI20227_Nintedanib_OCI-AML3.pdf

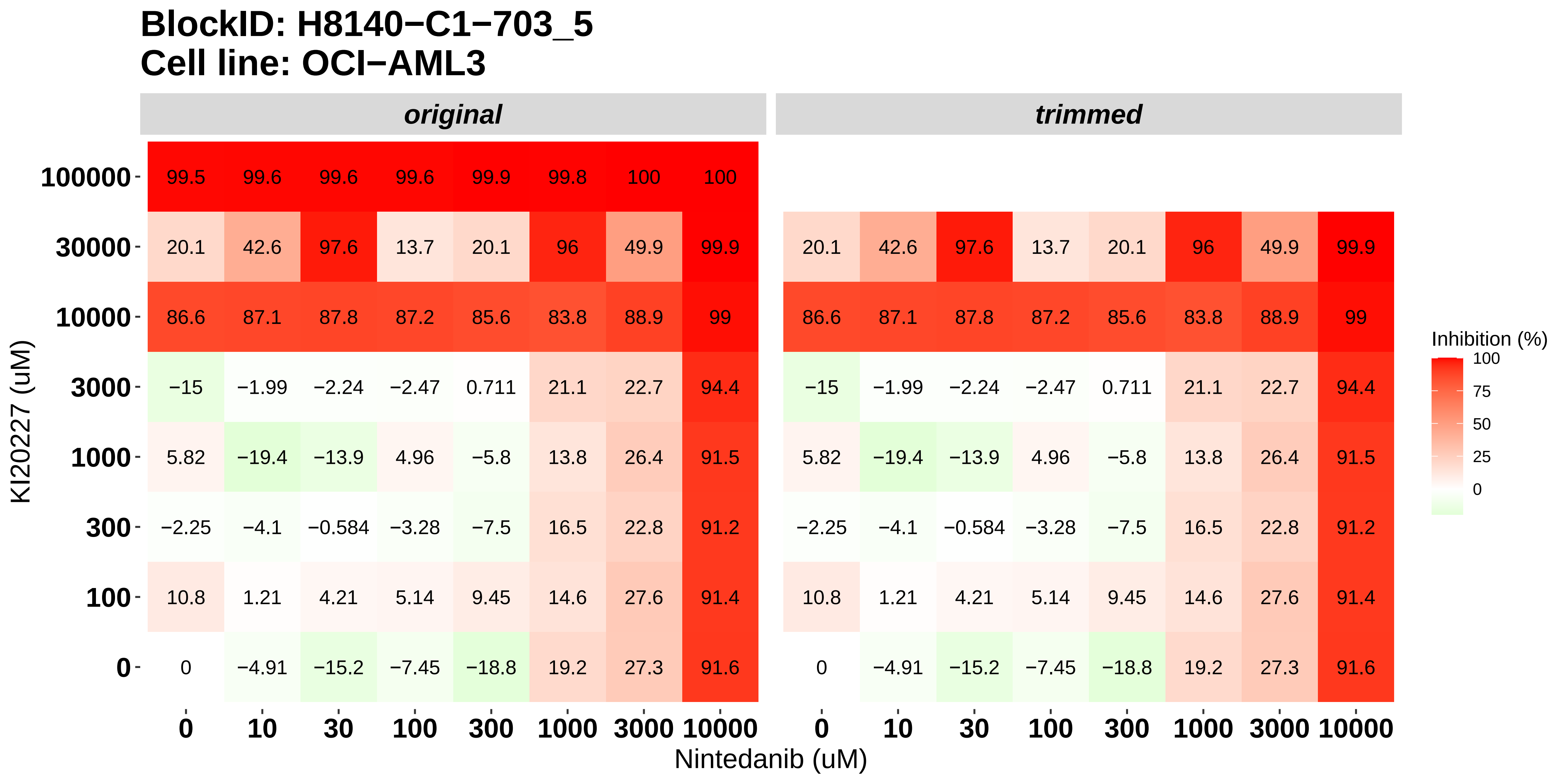

### H8140-C1-703_6_Cabozantinib_Nintedanib_OCI-AML3.pdf

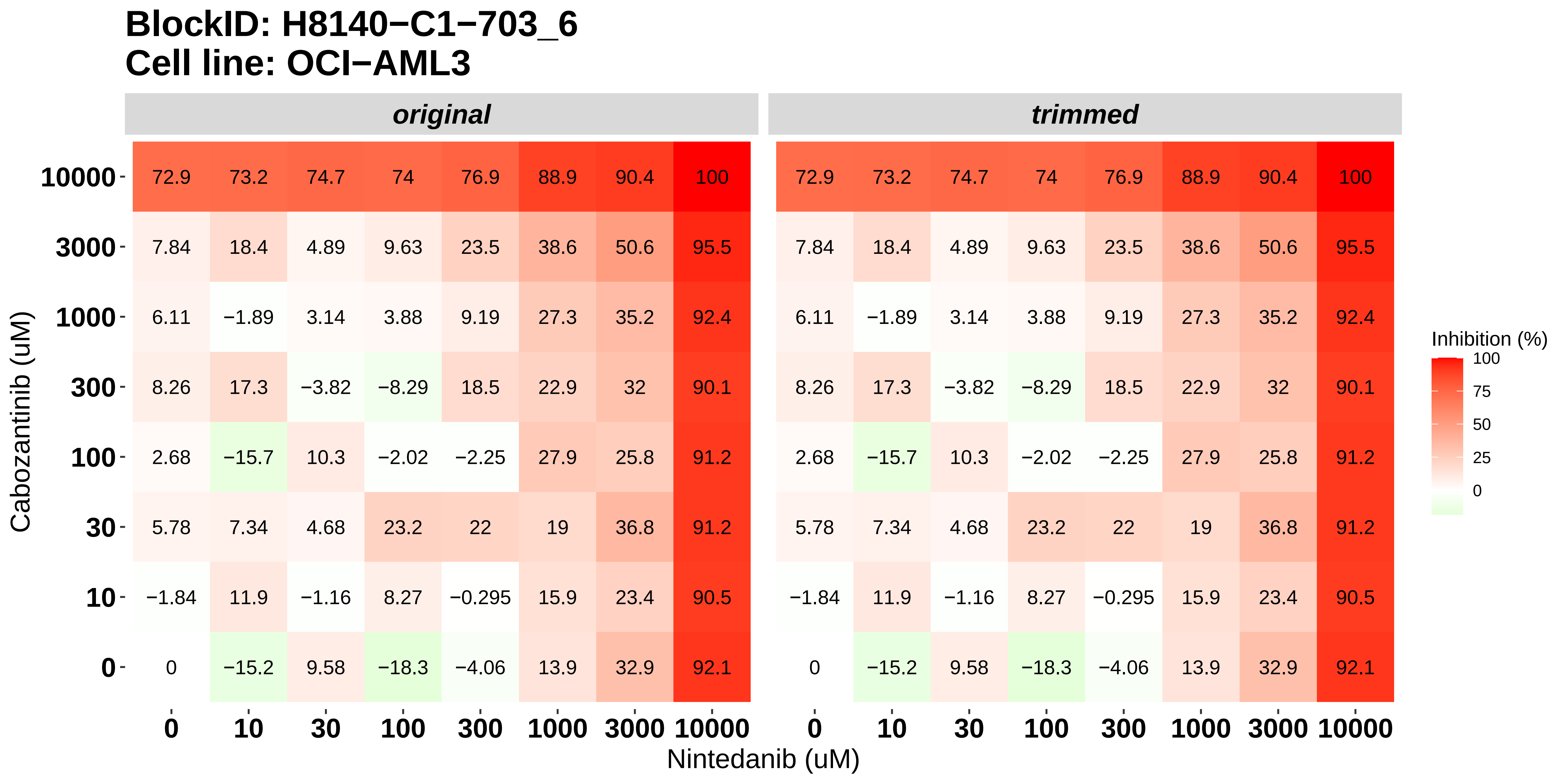

### H8140-C1-801_1_KI20227_Doramapimod_MOLM-16.pdf

#### BlockID: H8140−C1−801_1
#### Cell line: MOLM−16

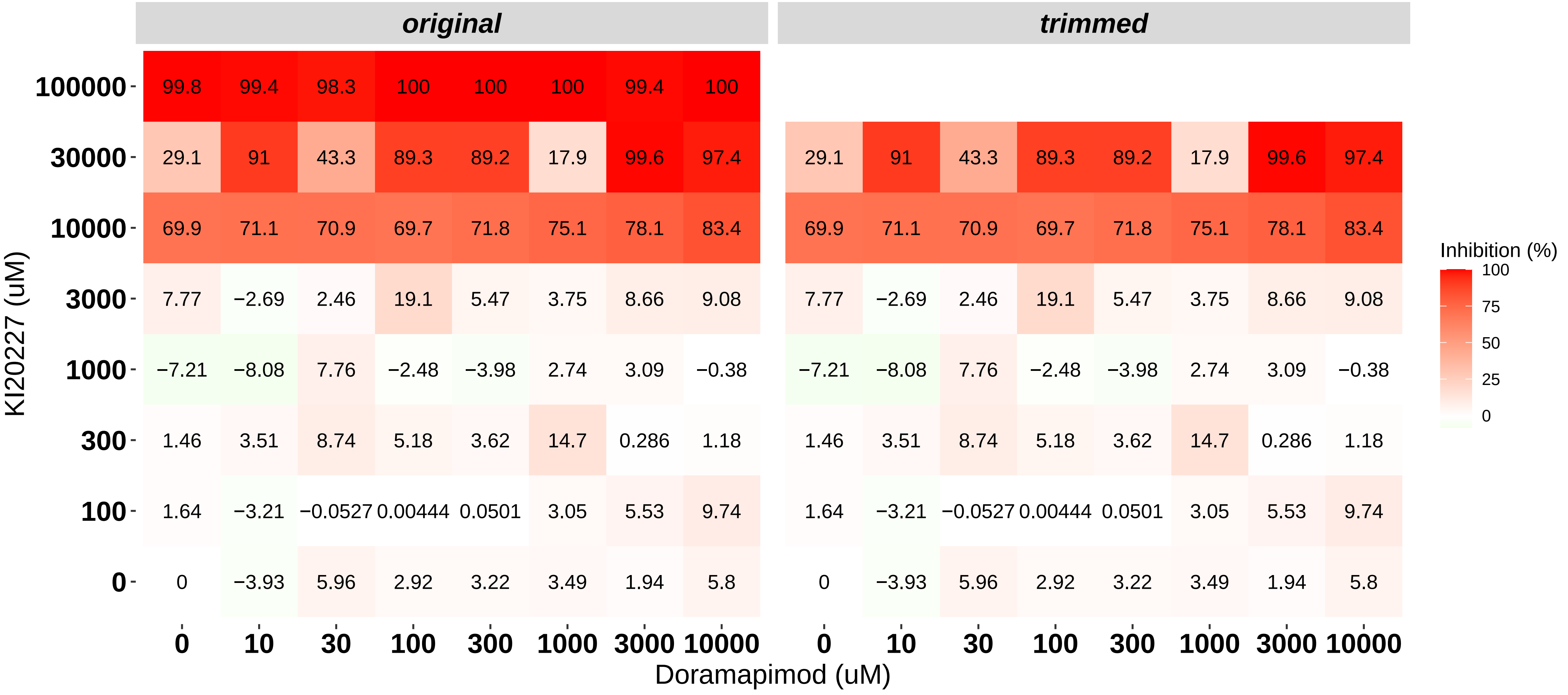

### H8140-C1-801_2_Cabozantinib_Doramapimod_MOLM-16.pdf

#### BlockID: H8140−C1−801_2
#### Cell line: MOLM−16

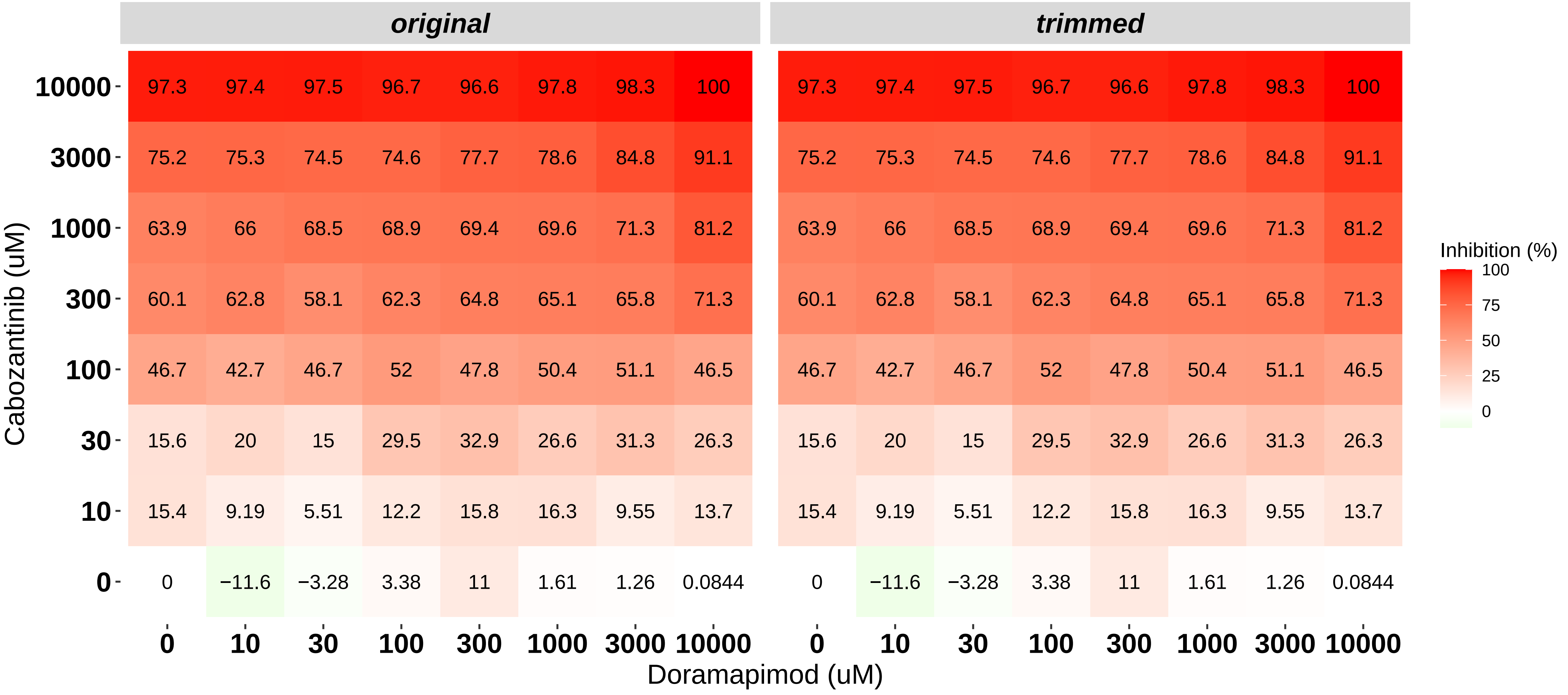

### H8140-C1-801_3_Cabozantinib_KI20227_MOLM-16.pdf

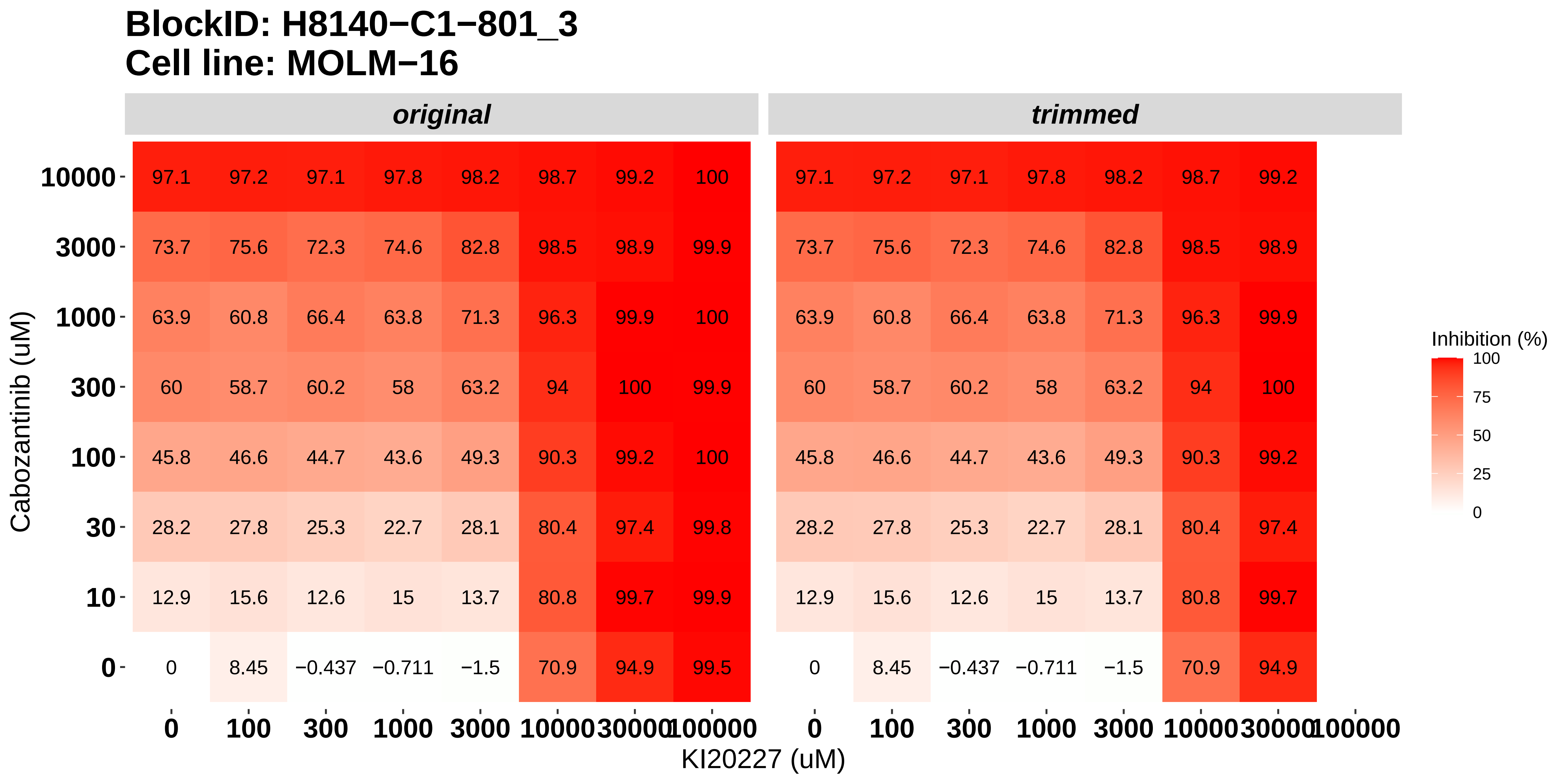

### H8140-C1-802_1_KI20227_Doramapimod_NOMO-1.pdf

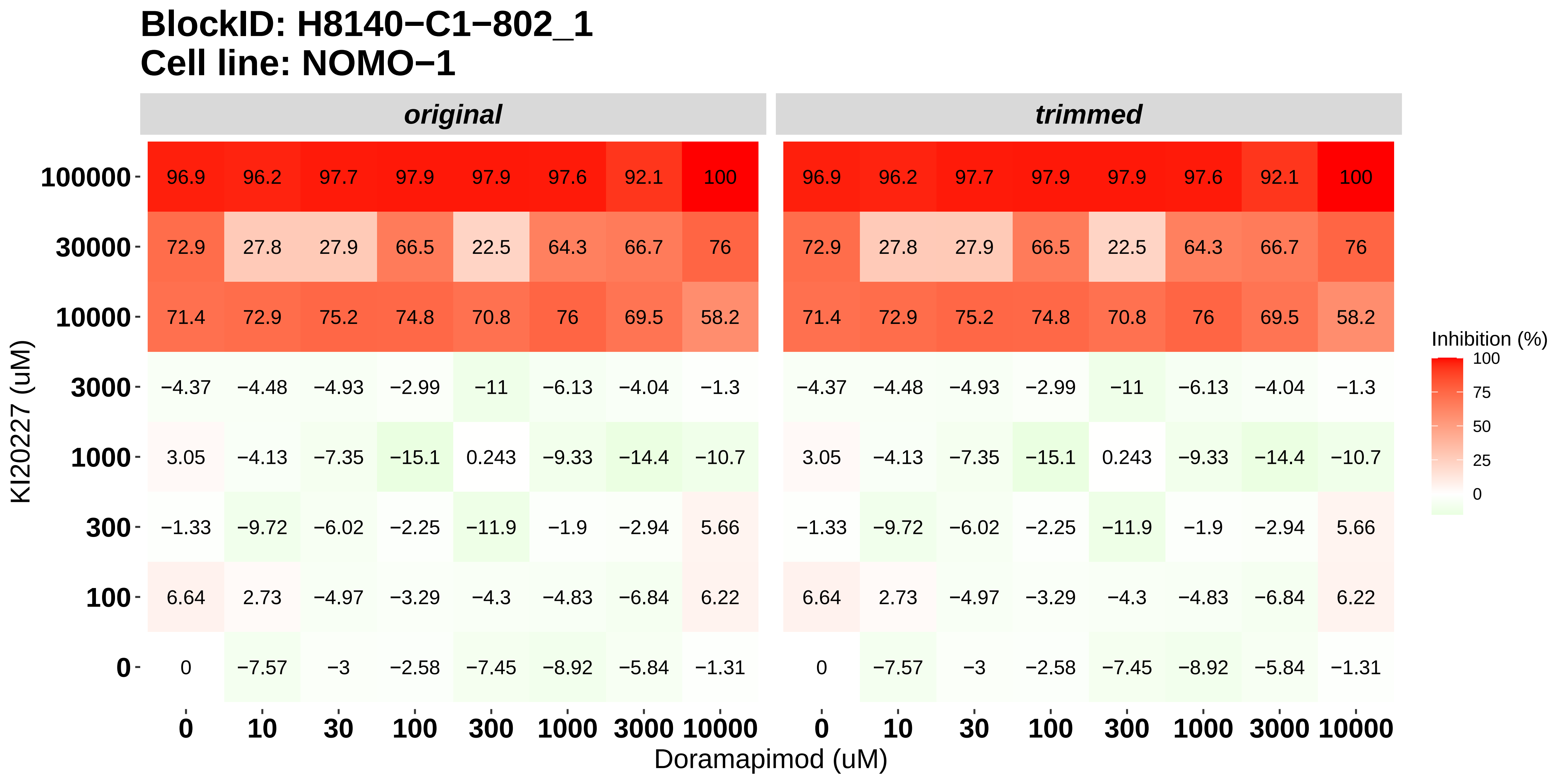

### H8140-C1-802_2_Cabozantinib_Doramapimod_NOMO-1.pdf

**BlockID: H8140−C1−802_2**
**Cell line: NOMO−1**

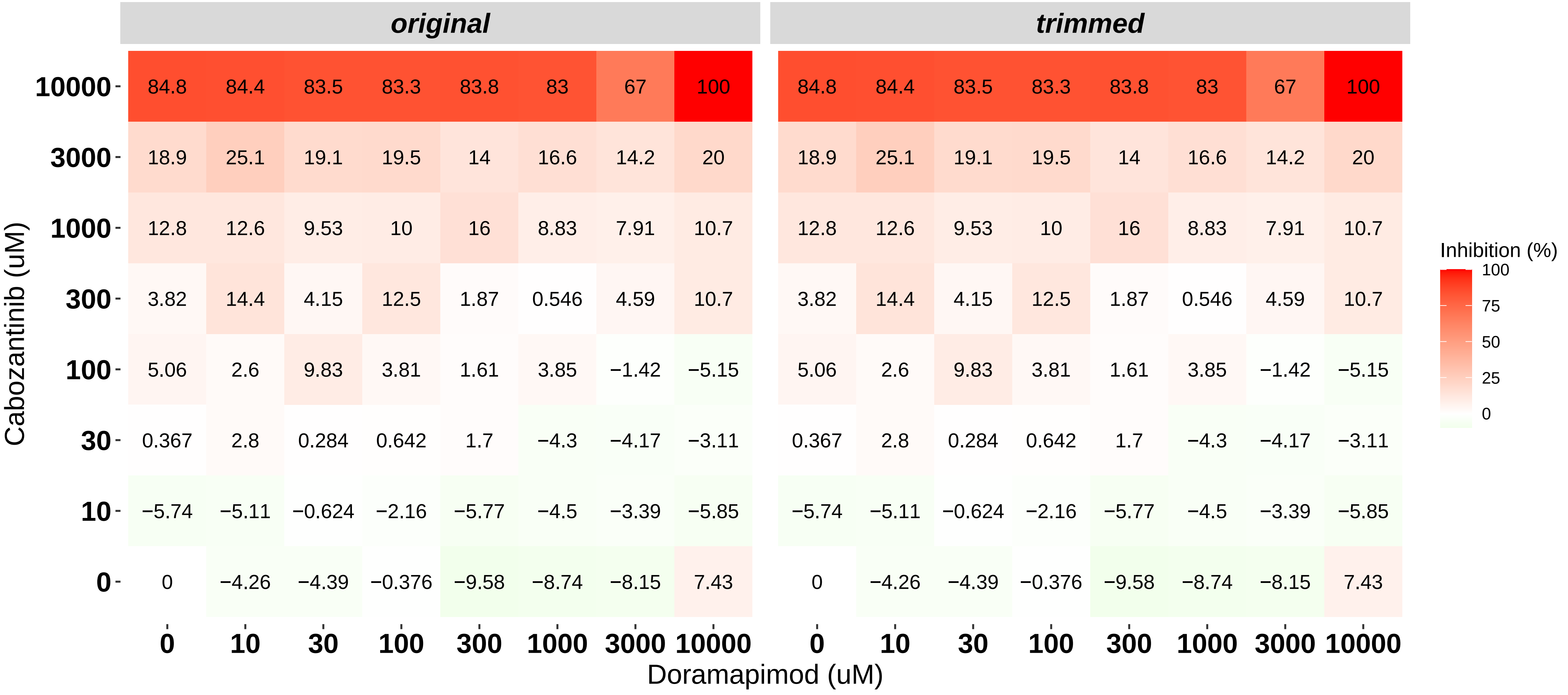

### H8140-C1-802_3_Cabozantinib_KI20227_NOMO-1.pdf

#### BlockID: H8140−C1−802_3
#### Cell line: NOMO−1

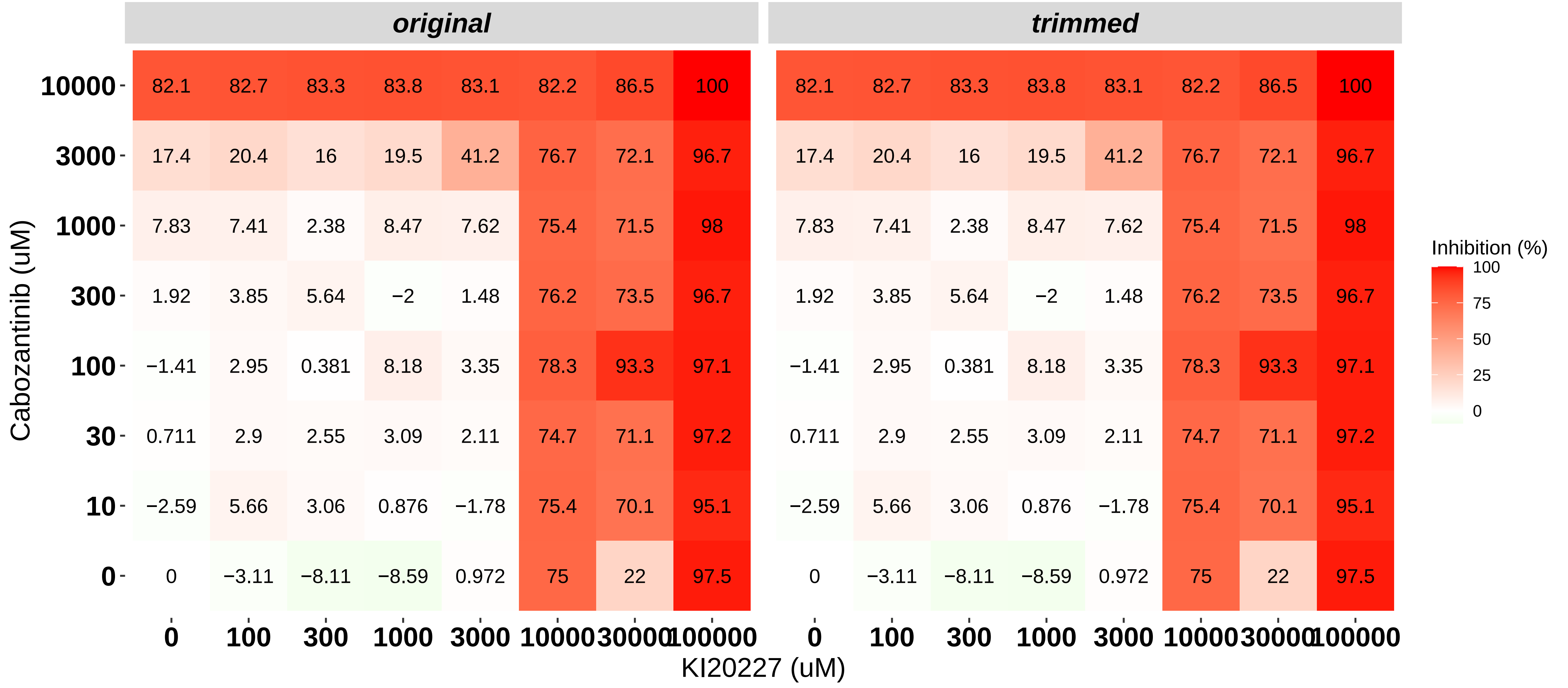

### H8140-C1-803_1_KI20227_Doramapimod_OCI-AML3.pdf

#### BlockID: H8140−C1−803_1
#### Cell line: OCI−AML3

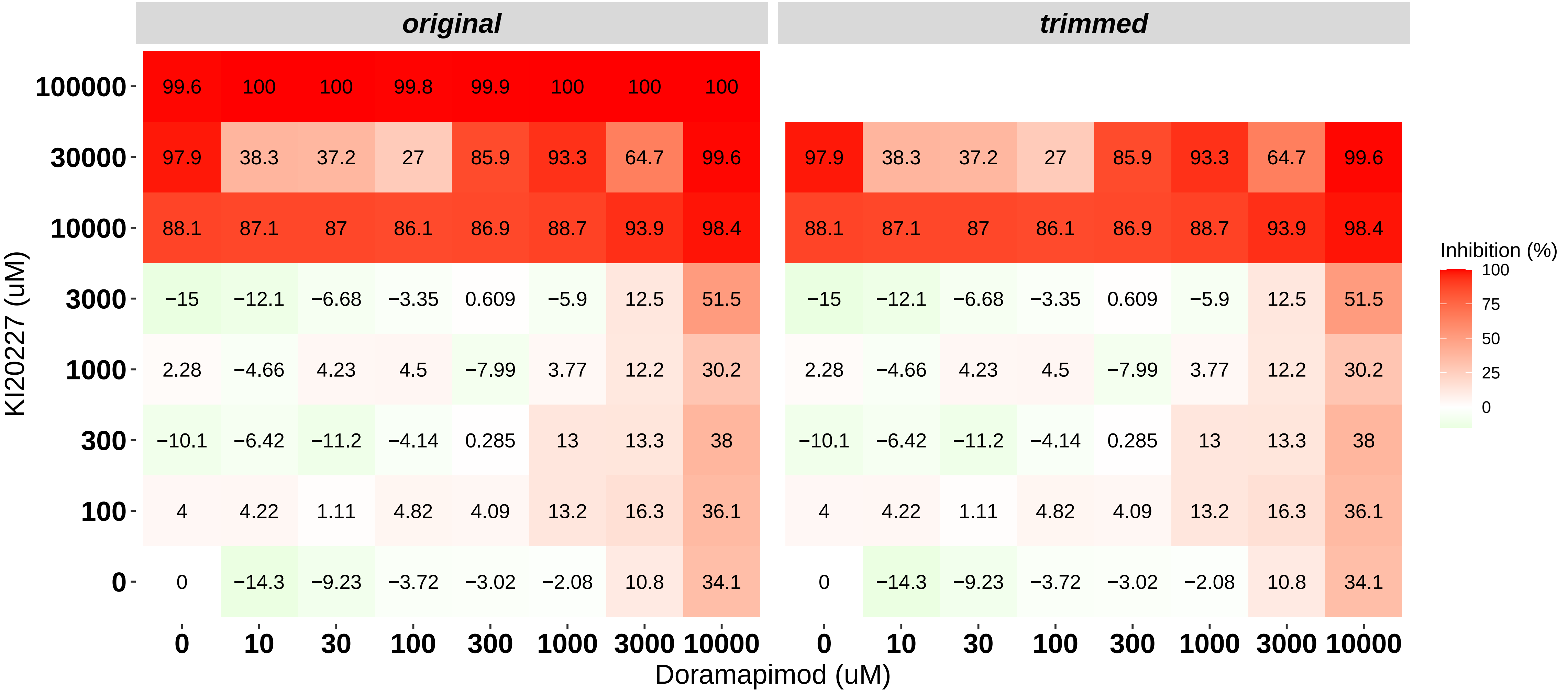

### H8140-C1-803_2_Cabozantinib_Doramapimod_OCI-AML3.pdf

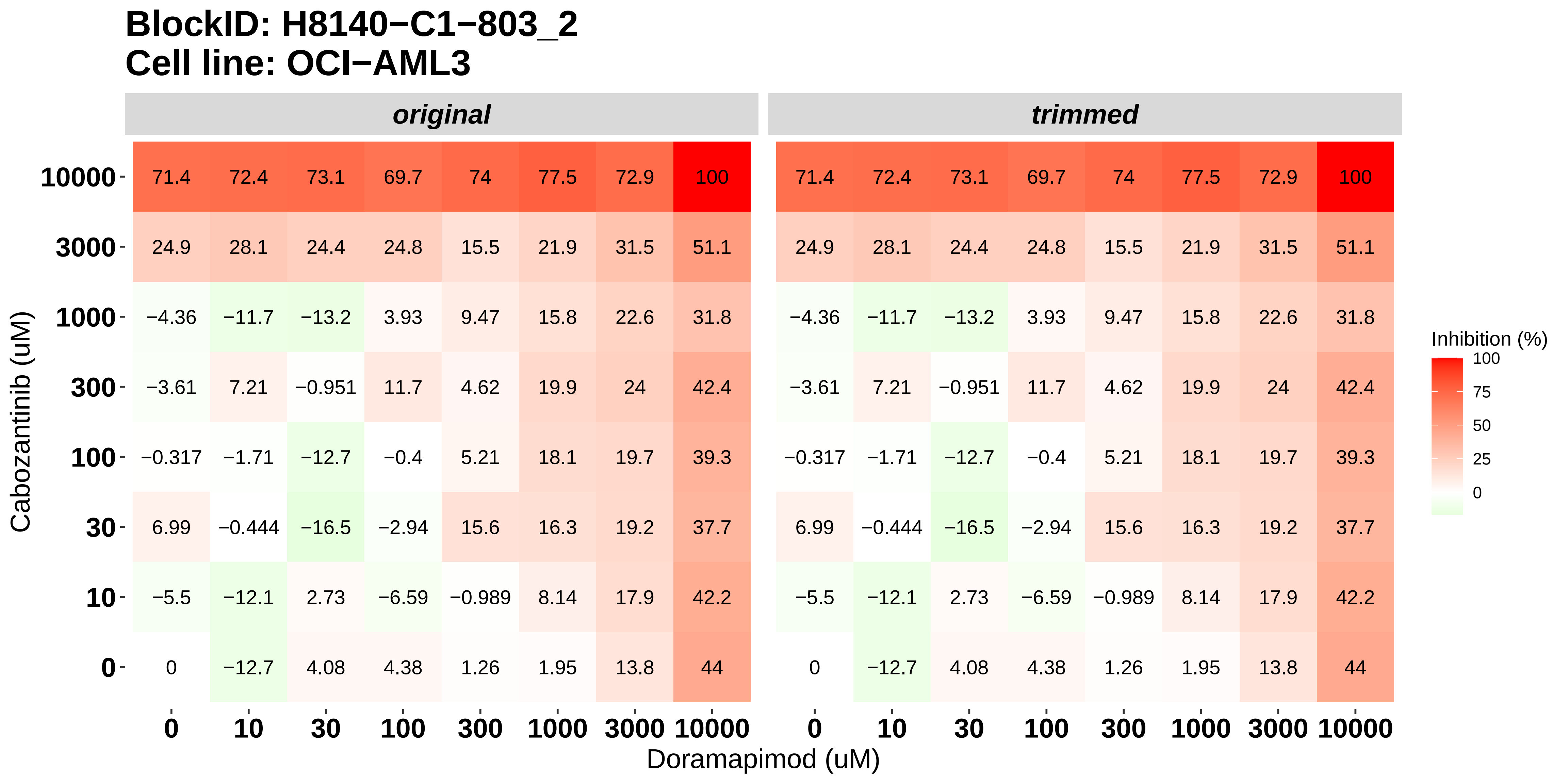

### H8140-C1-803_3_Cabozantinib_KI20227_OCI-AML3.pdf

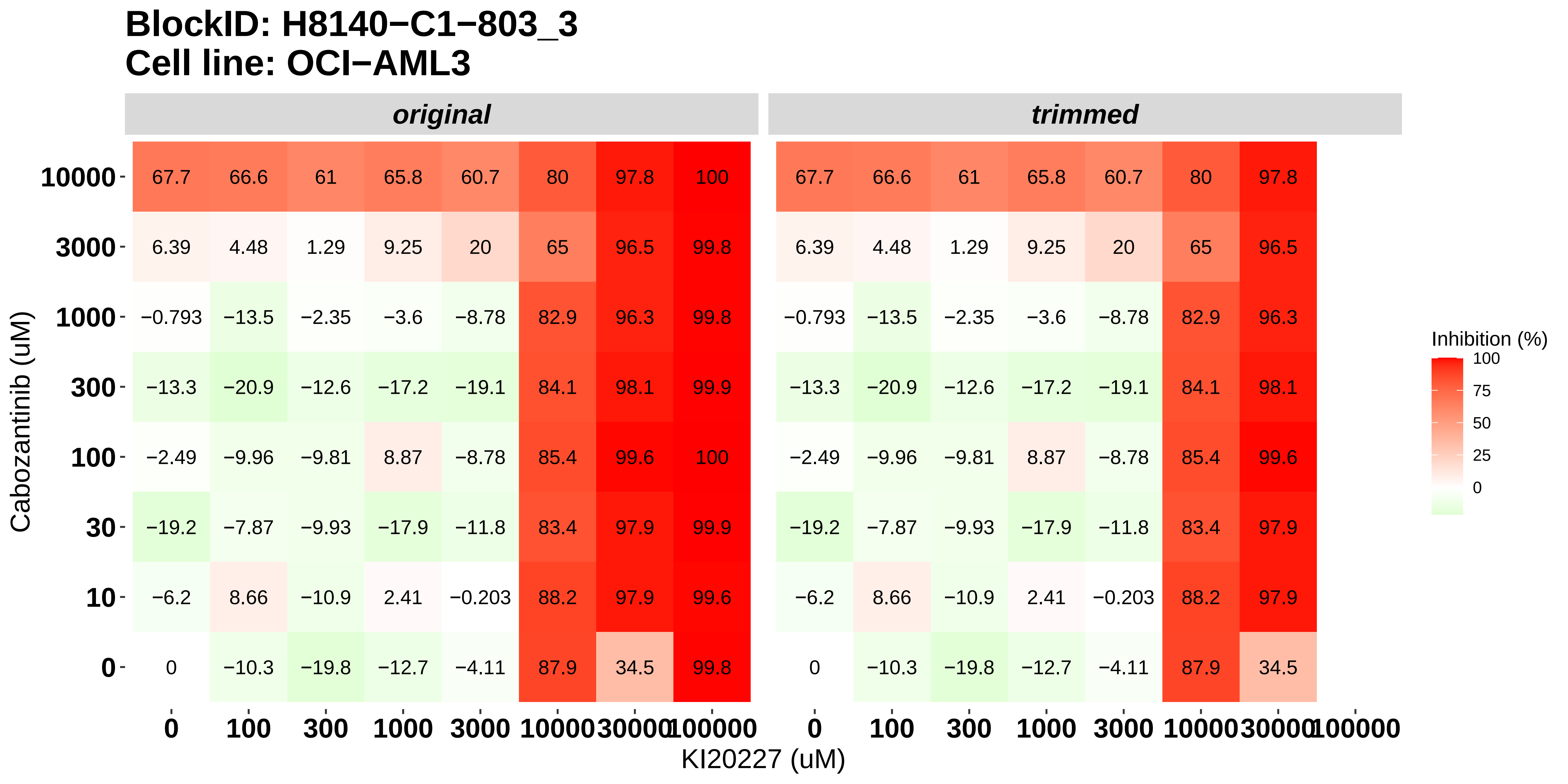

### Supplementary Fig. 1

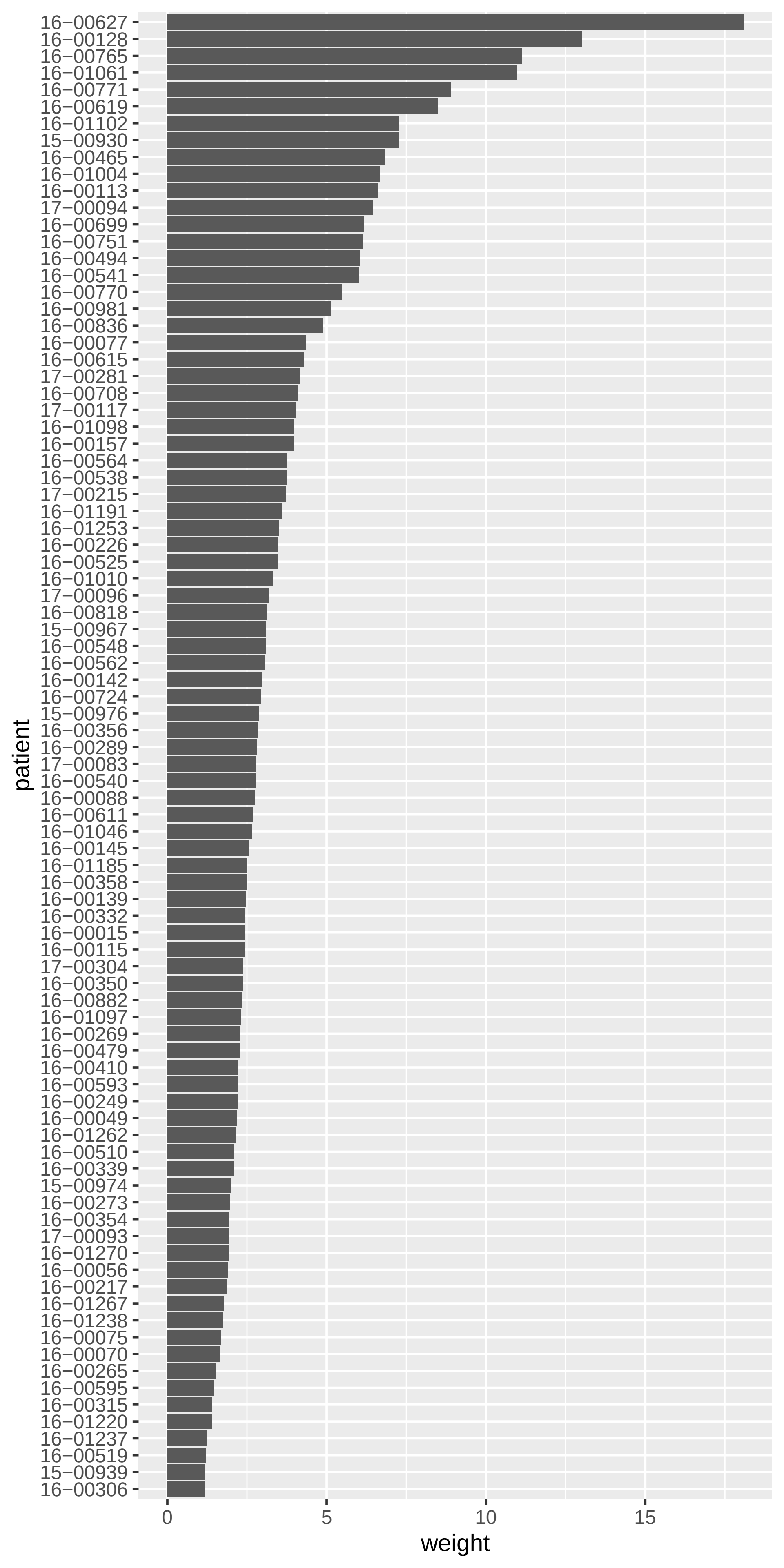

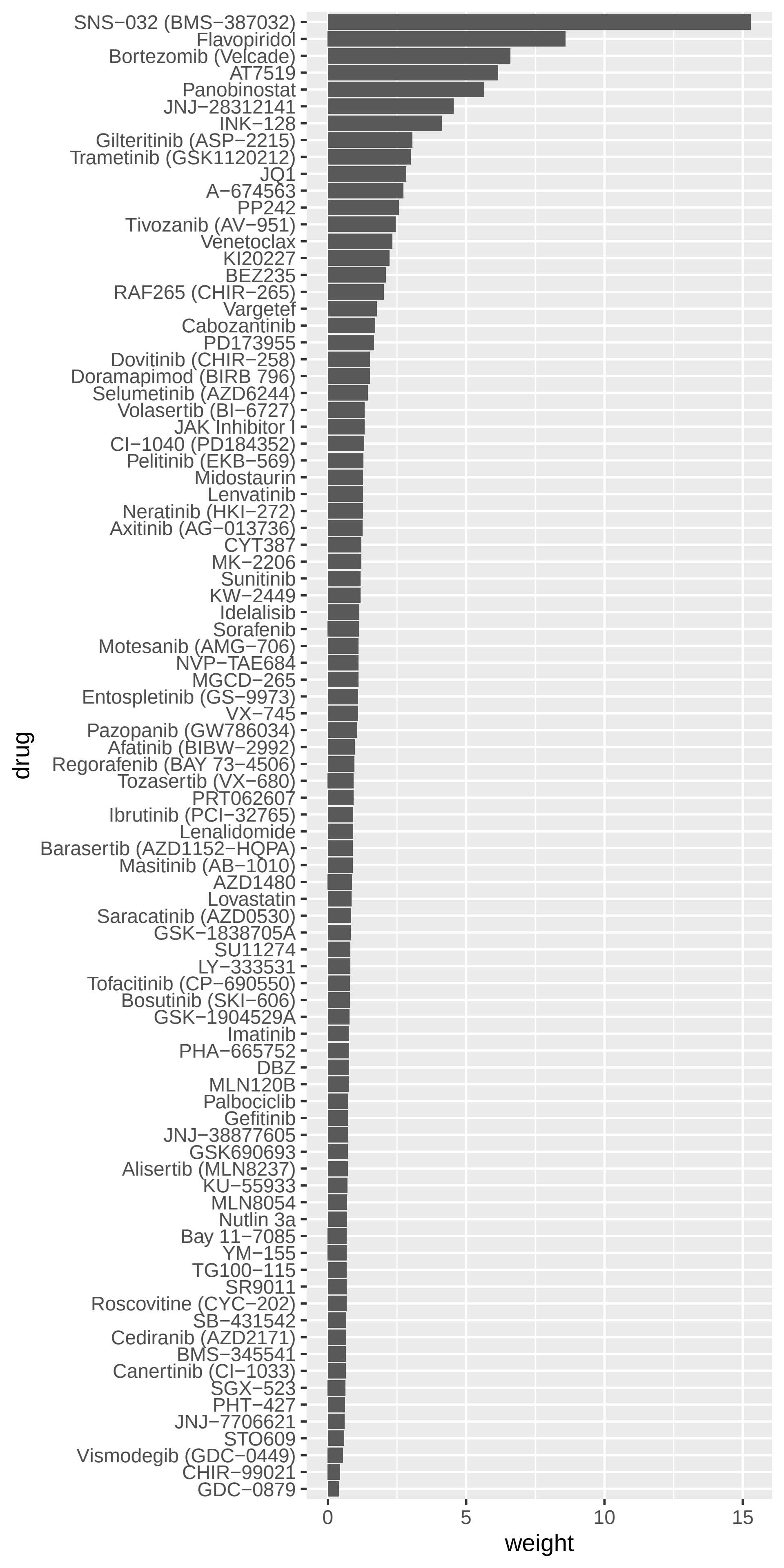
