## Supplementary figures and images for "Facilitating the design of combination therapy in cancer using multipartite network models: Emphasis on acute myeloid leukemia"

### H8140-C1-101_1_Dovitinib_SNS-032_MOLM-16.pdf

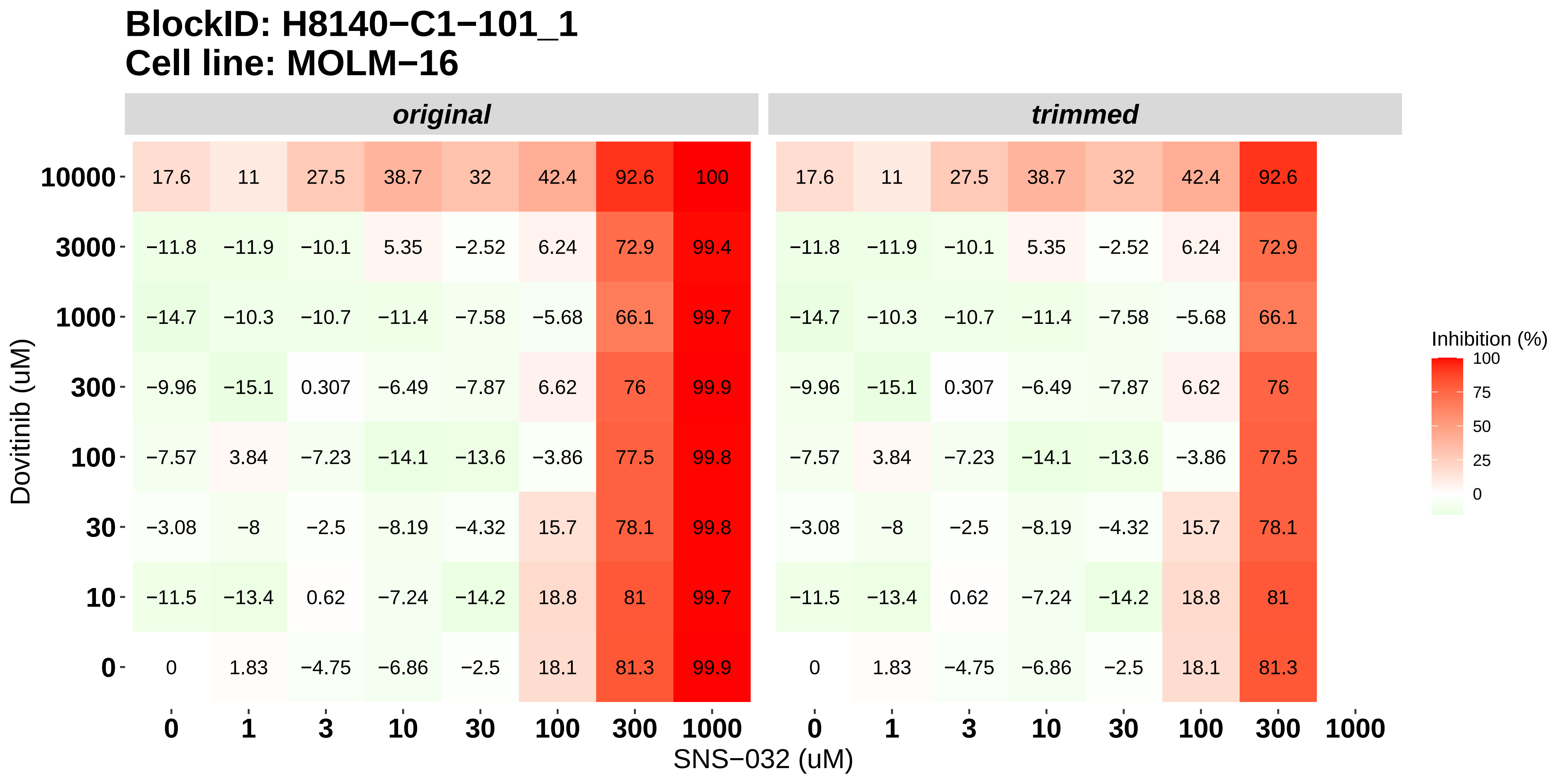

### H8140-C1-101_2_Nintedanib_SNS-032_MOLM-16.pdf

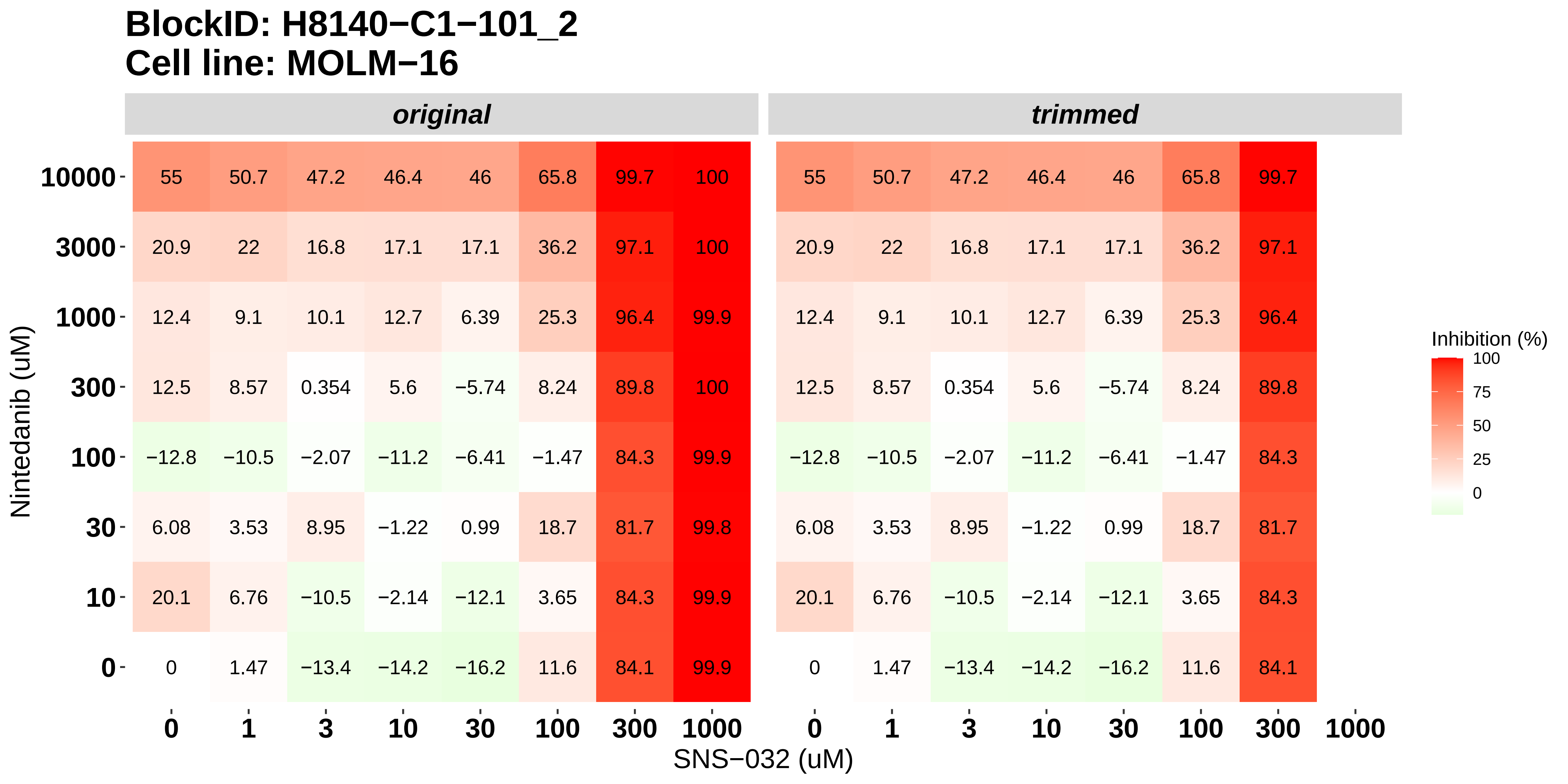

### H8140-C1-101_3_Doramapimod_SNS-032_MOLM-16.pdf

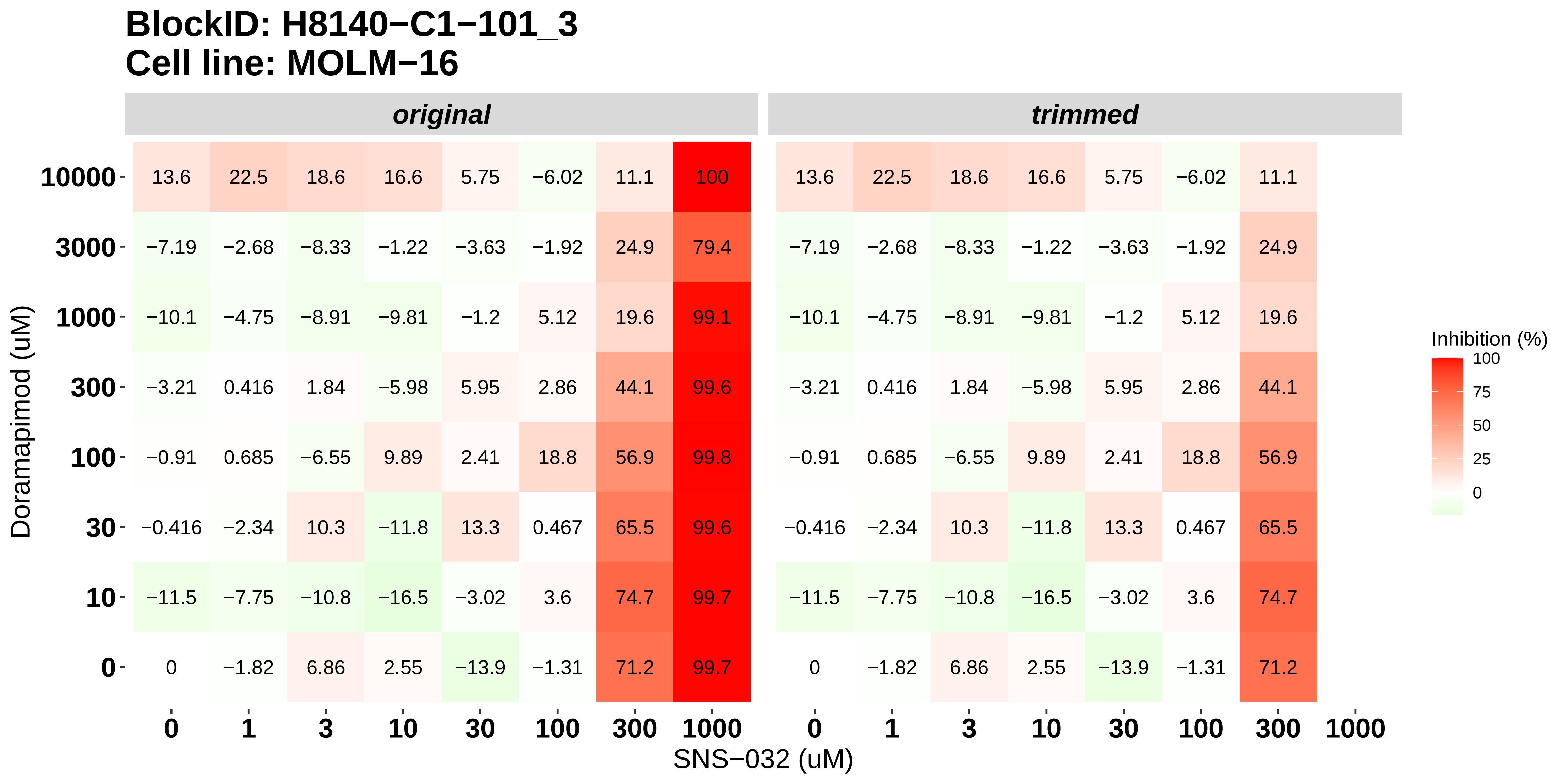

### H8140-C1-101_4_KI20227_SNS-032_MOLM-16.pdf

#### BlockID: H8140−C1−101_4
##### Cell line: MOLM−16

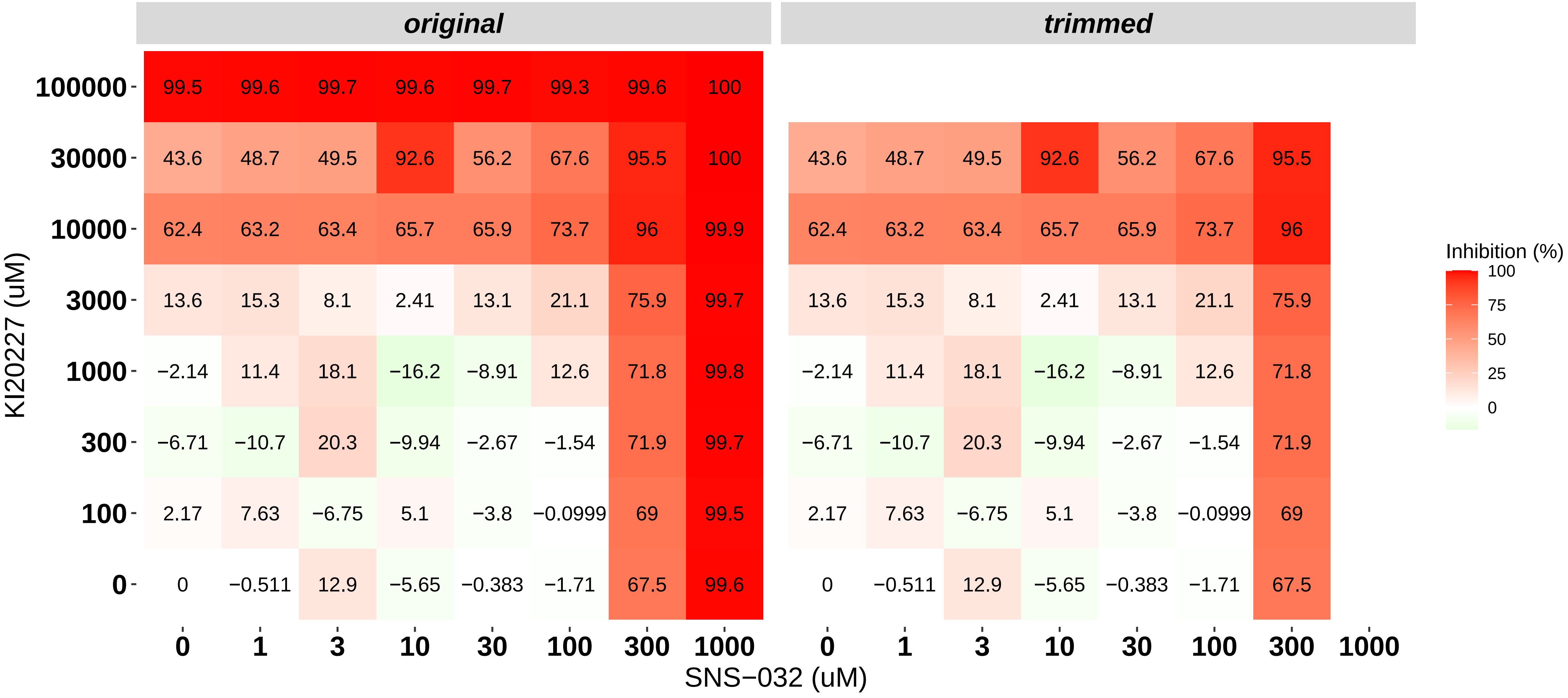

### H8140-C1-101_5_Cabozantinib_SNS-032_MOLM-16.pdf

#### BlockID: H8140−C1−101_5
#### Cell line: MOLM−16

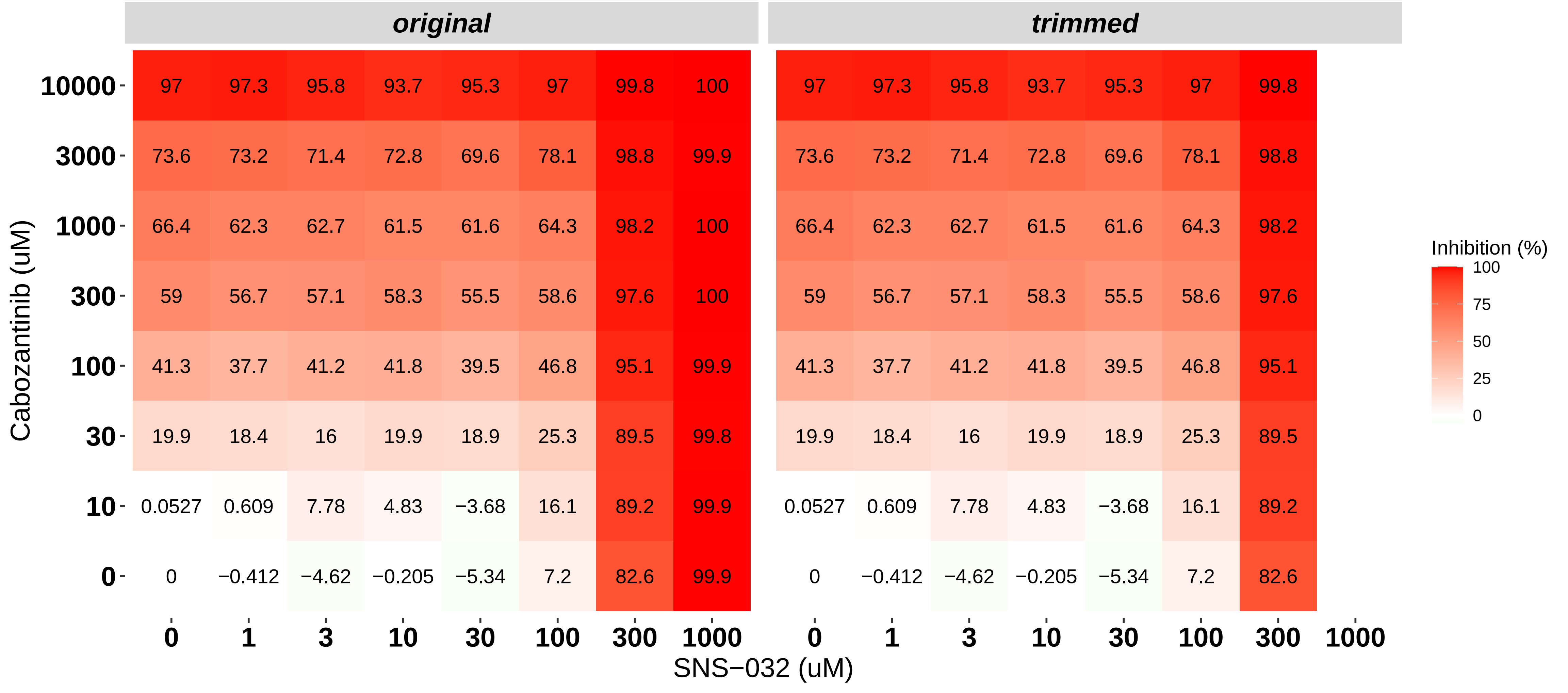

### H8140-C1-101_6_Dovitinib_Alvocidib_MOLM-16.pdf

#### BlockID: H8140−C1−101_6
#### Cell line: MOLM−16

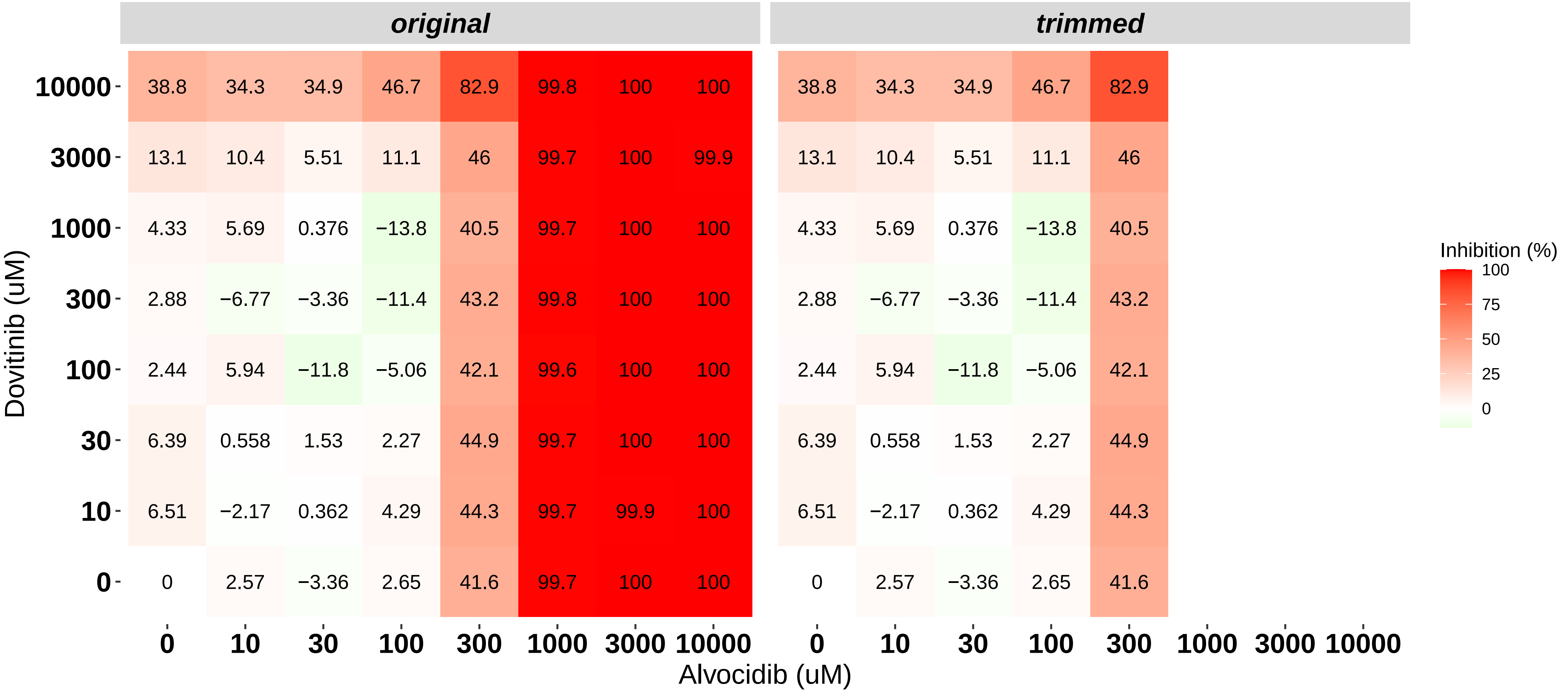

### H8140-C1-102_1_Dovitinib_SNS-032_NOMO-1.pdf

**BlockID: H8140−C1−102_1**
**Cell line: NOMO−1**

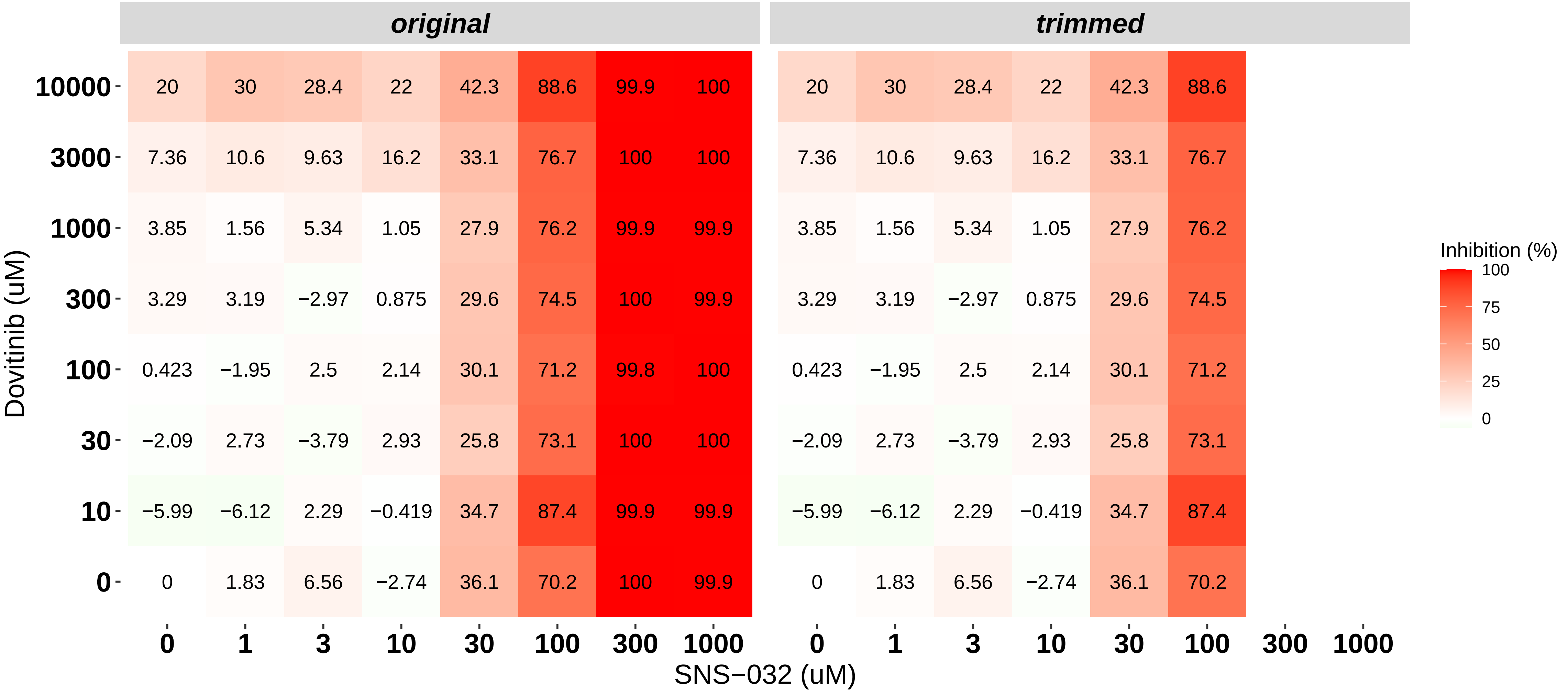

### H8140-C1-102_2_Nintedanib_SNS-032_NOMO-1.pdf

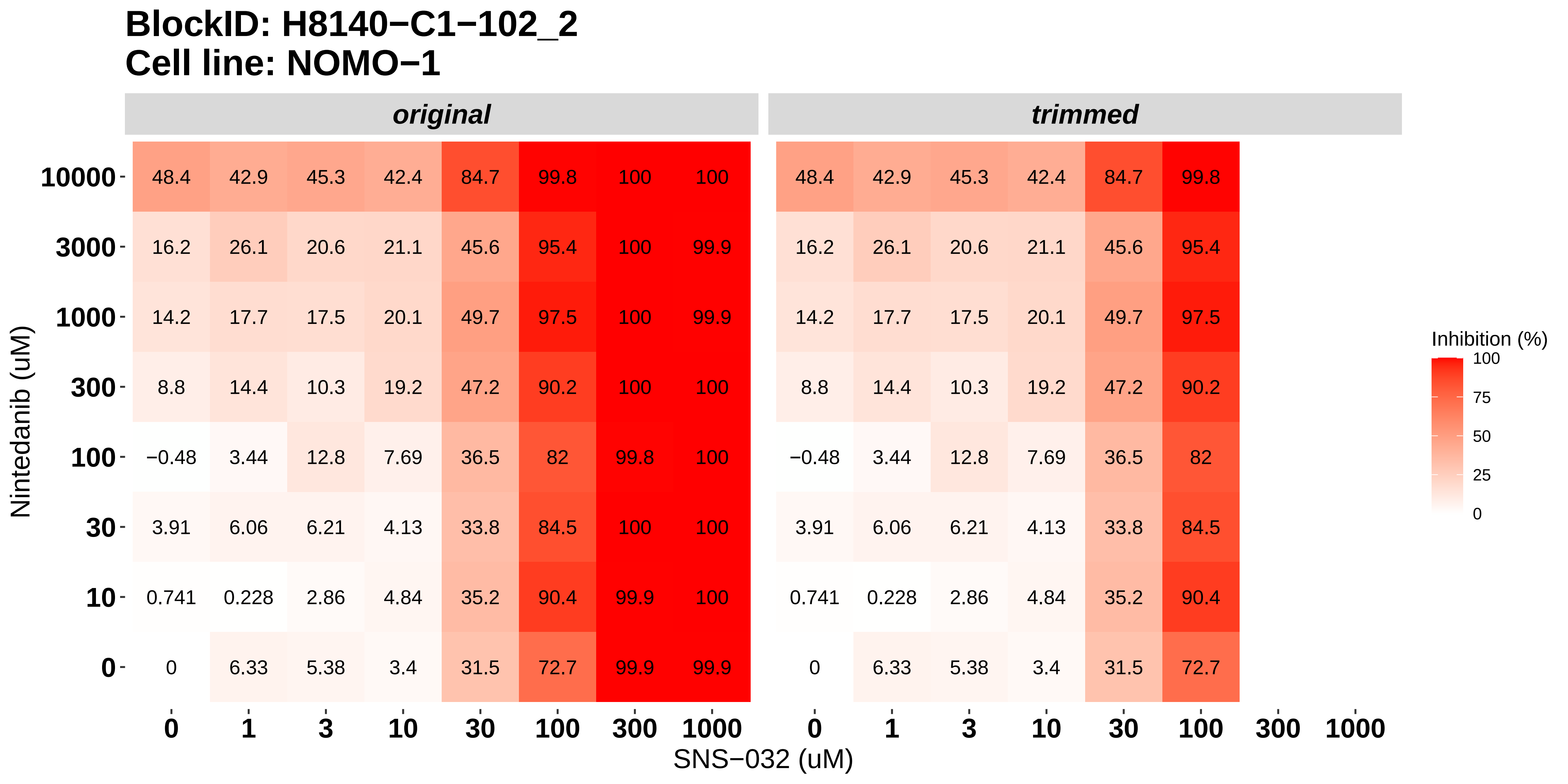

### H8140-C1-102_3_Doramapimod_SNS-032_NOMO-1.pdf

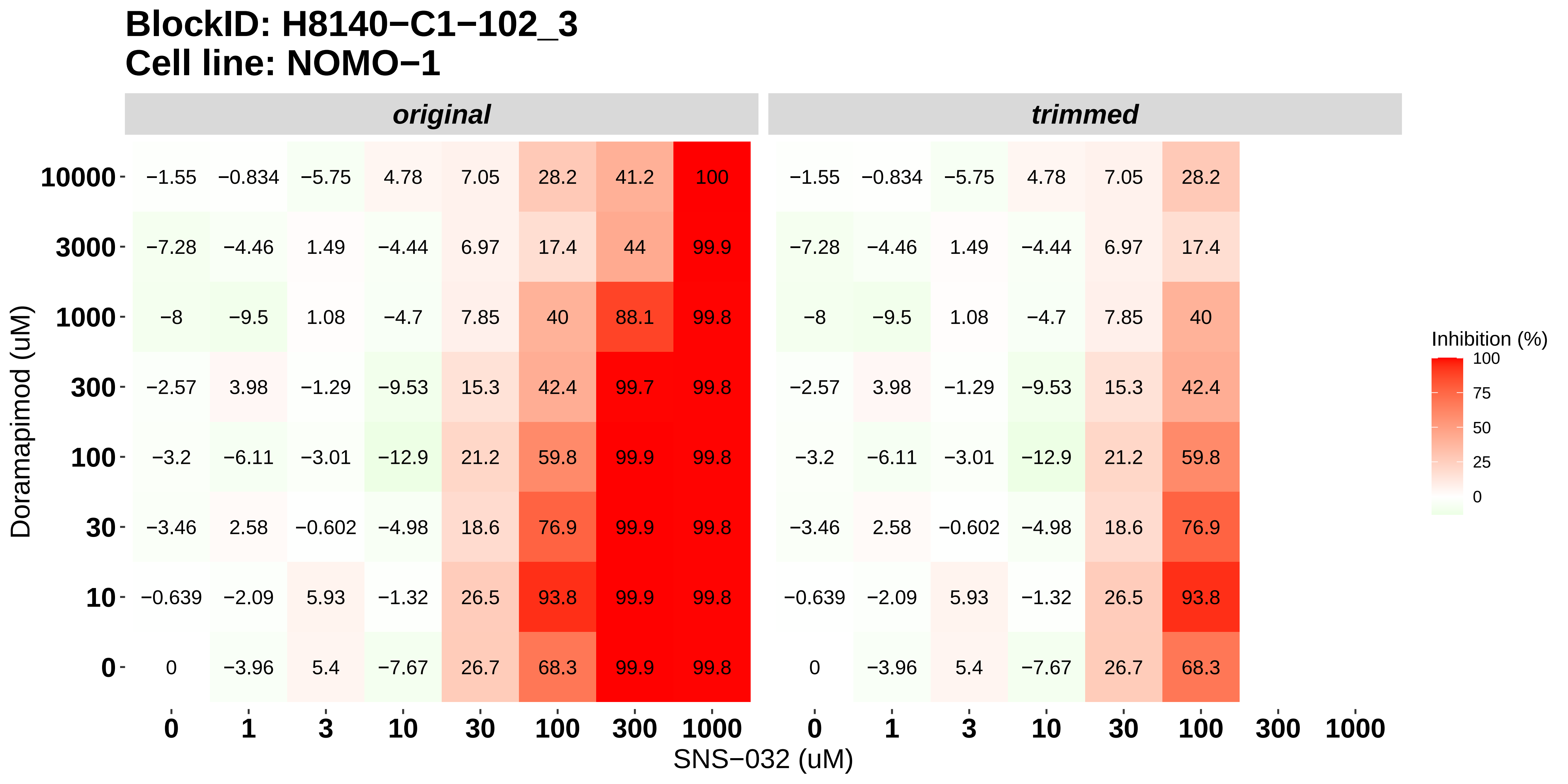

### H8140-C1-102_4_KI20227_SNS-032_NOMO-1.pdf

#### BlockID: H8140−C1−102_4
#### Cell line: NOMO−1

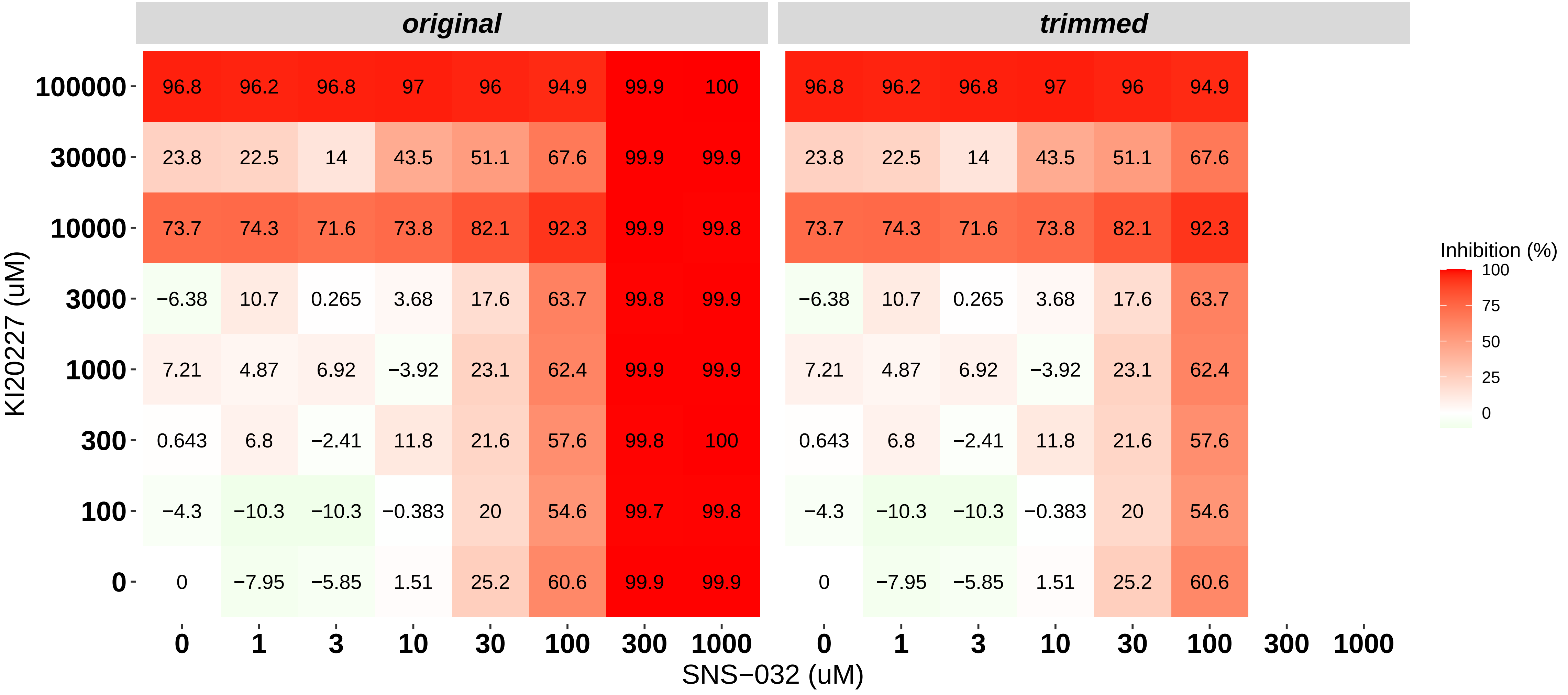

### H8140-C1-102_5_Cabozantinib_SNS-032_NOMO-1.pdf

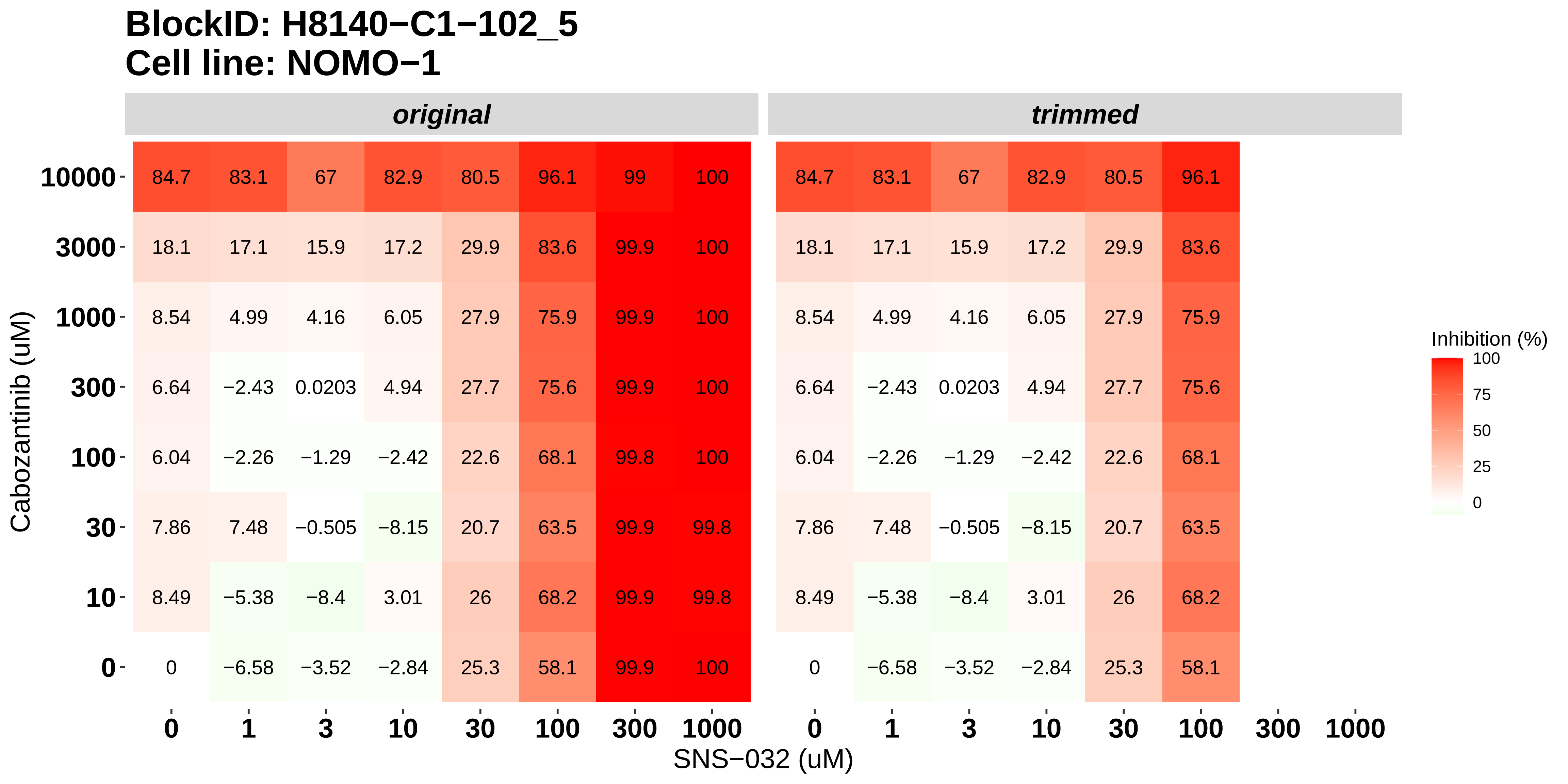

### H8140-C1-102_6_Dovitinib_Alvocidib_NOMO-1.pdf

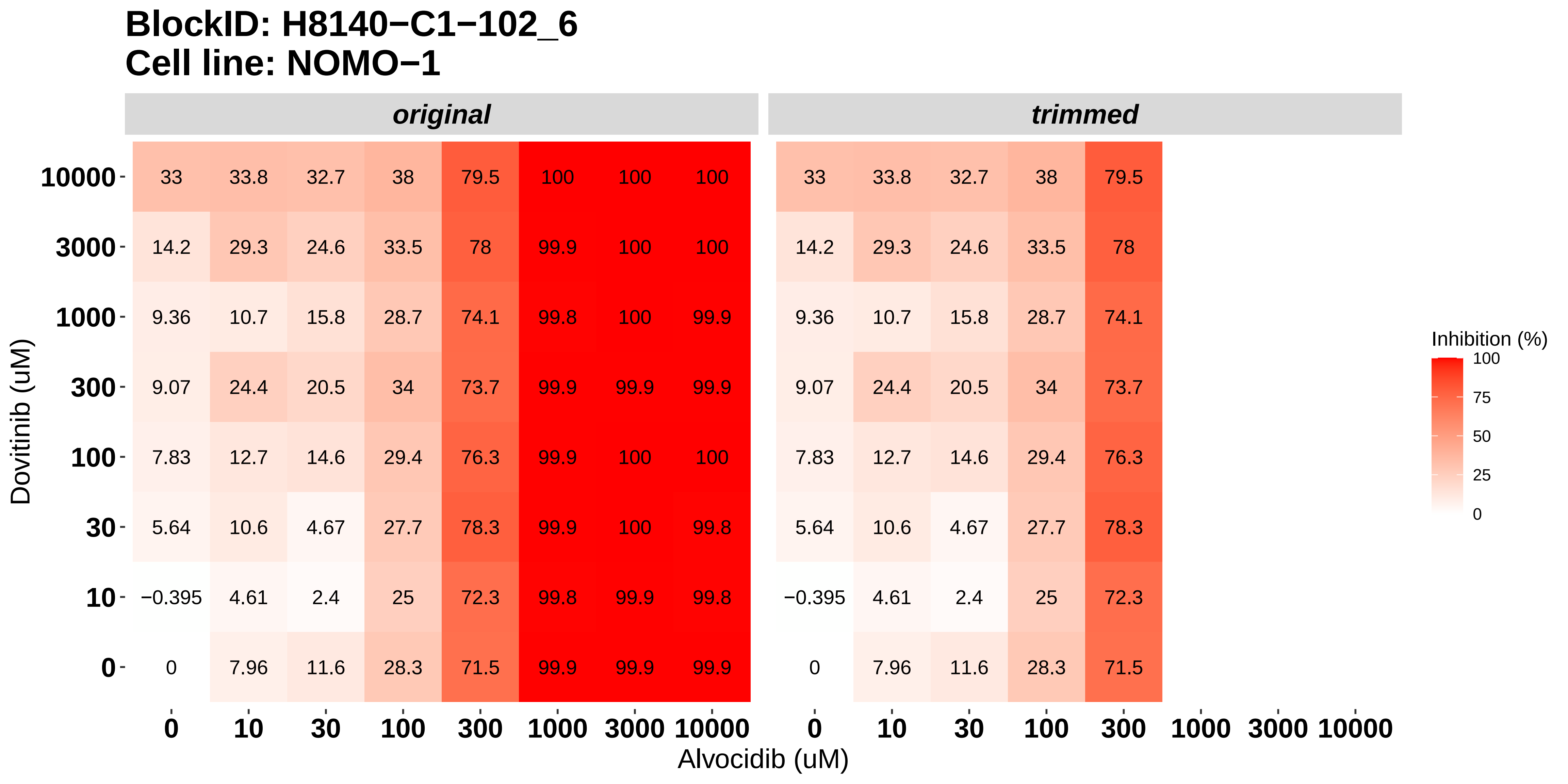

### H8140-C1-103_1_Dovitinib_SNS-032_OCI-AML3.pdf

#### BlockID: H8140−C1−103_1
#### Cell line: OCI−AML3

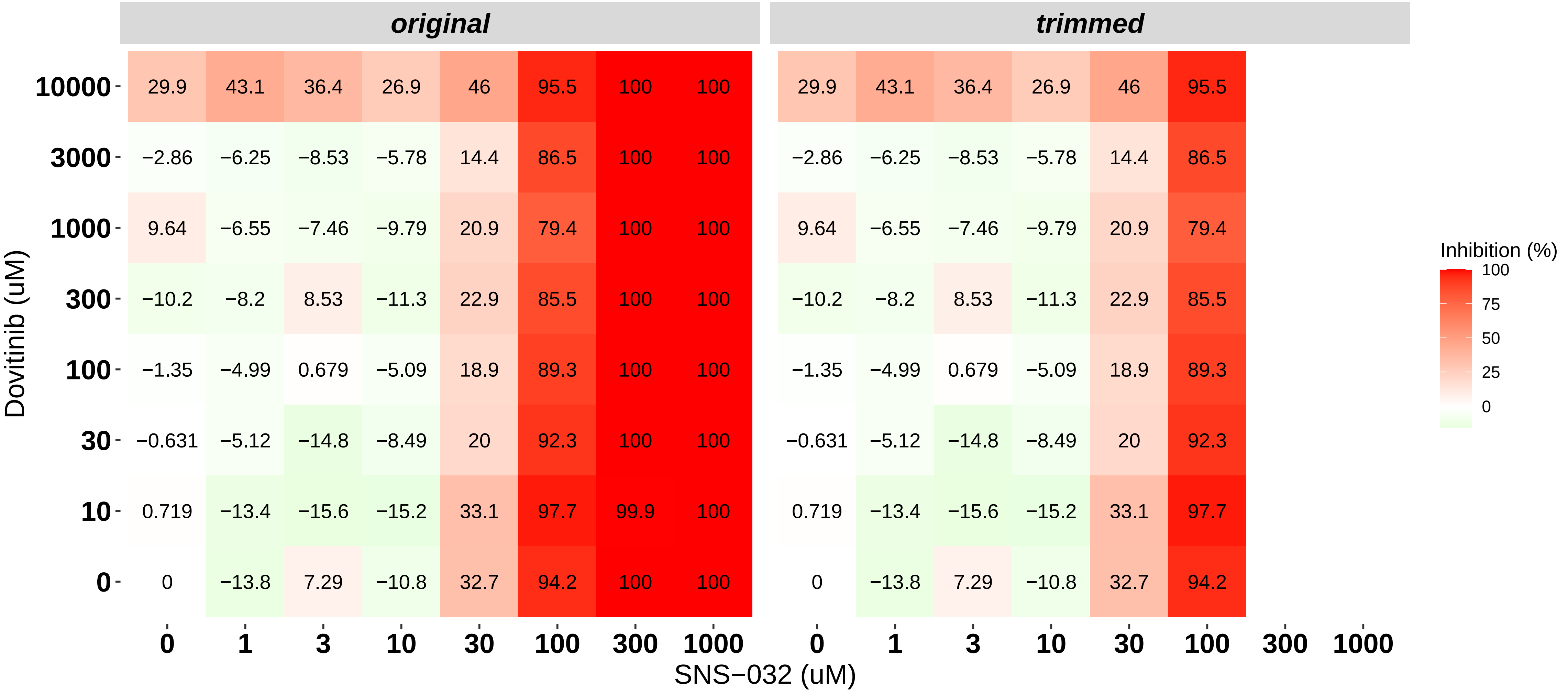

### H8140-C1-103_2_Nintedanib_SNS-032_OCI-AML3.pdf

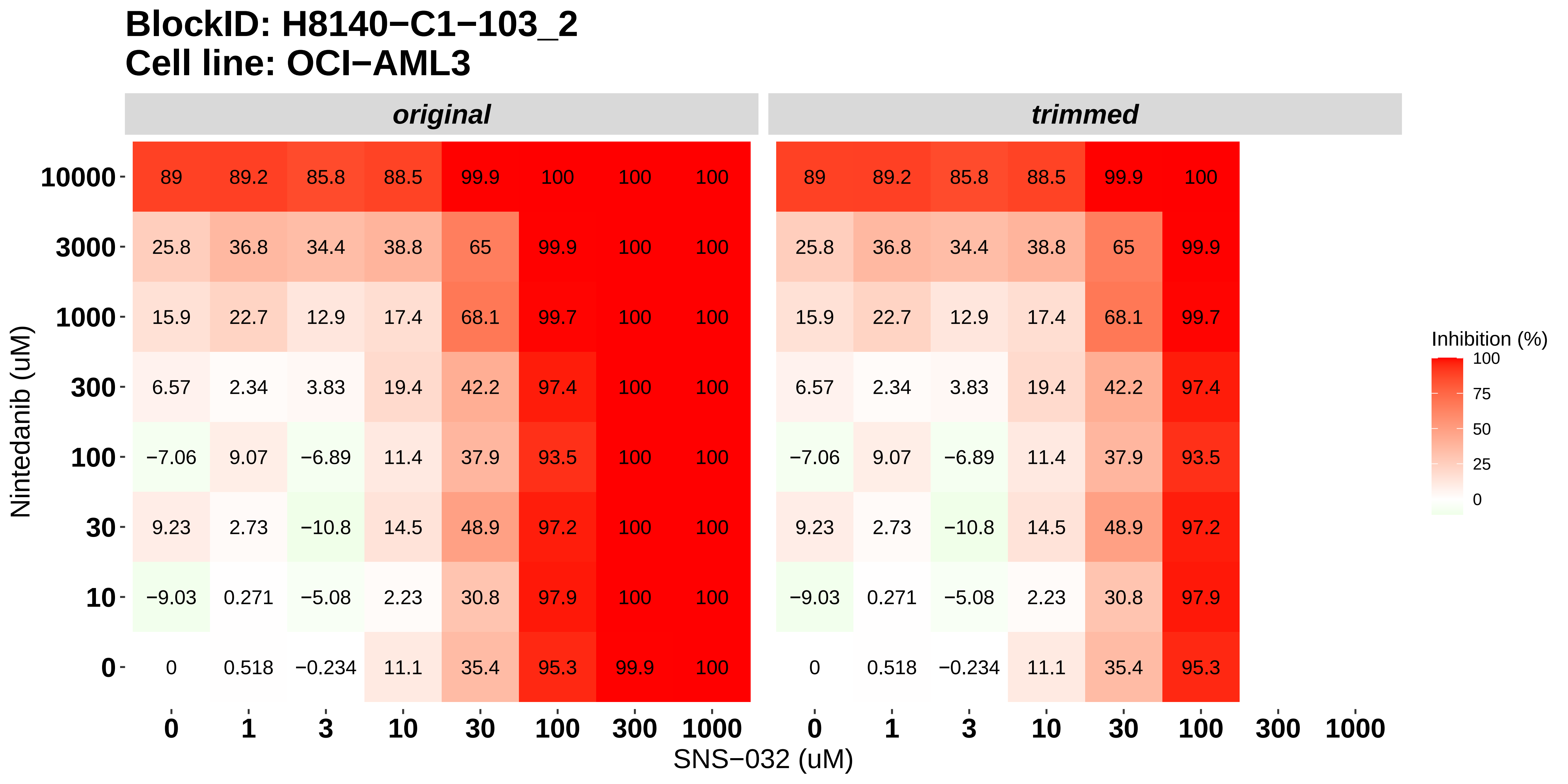

### H8140-C1-103_3_Doramapimod_SNS-032_OCI-AML3.pdf

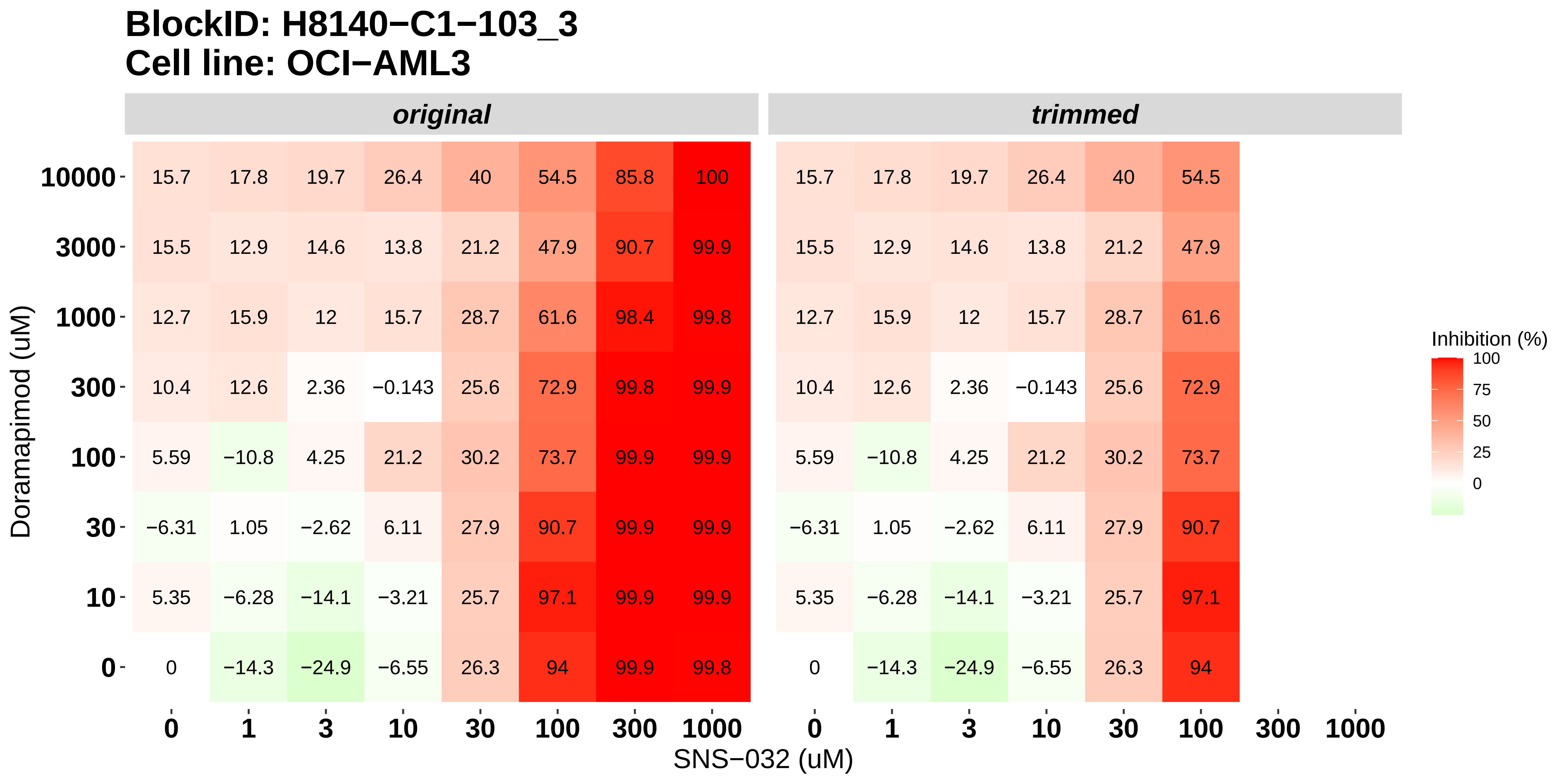

### H8140-C1-103_4_KI20227_SNS-032_OCI-AML3.pdf

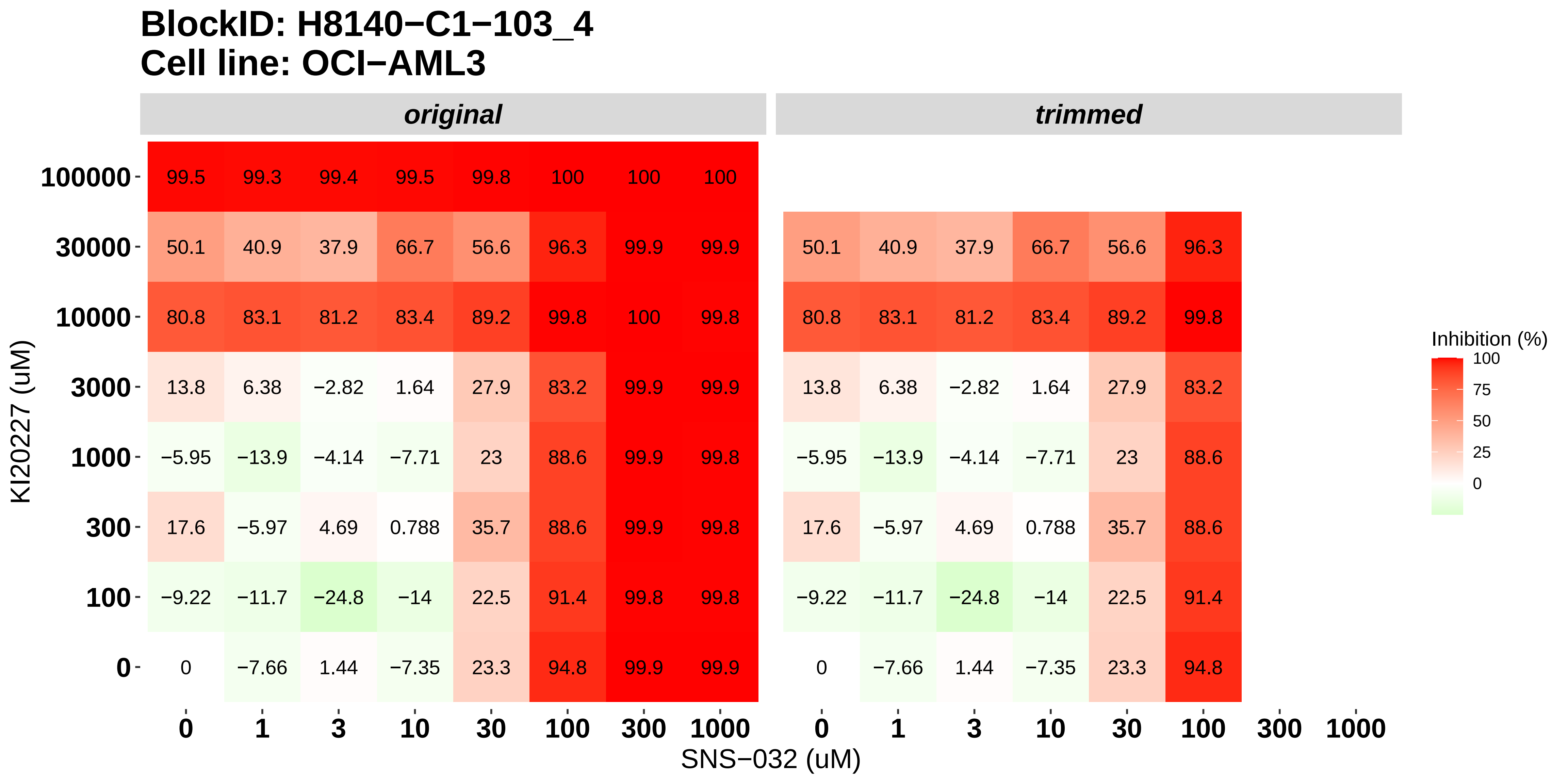

### H8140-C1-103_5_Cabozantinib_SNS-032_OCI-AML3.pdf

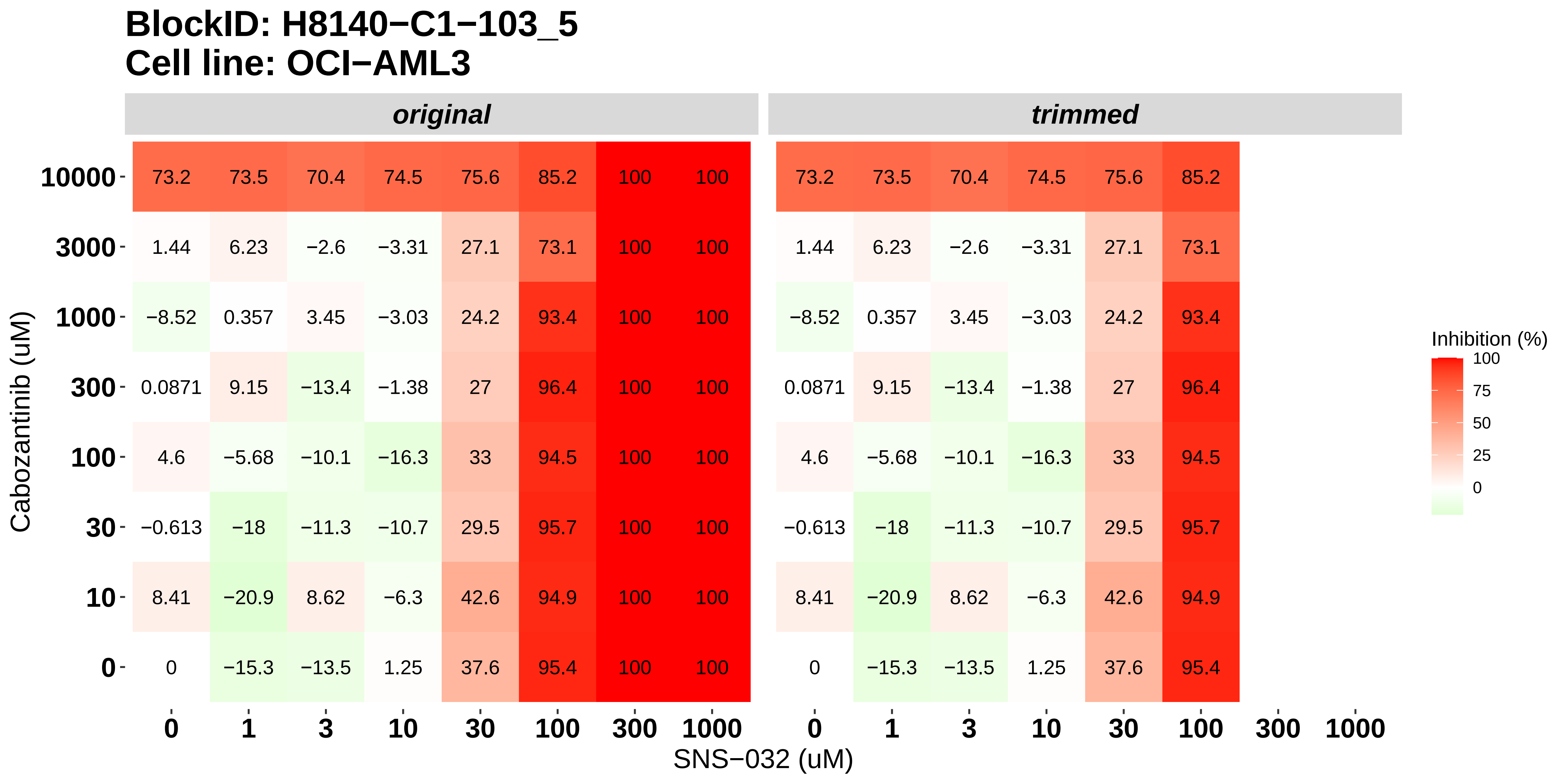

### H8140-C1-103_6_Dovitinib_Alvocidib_OCI-AML3.pdf

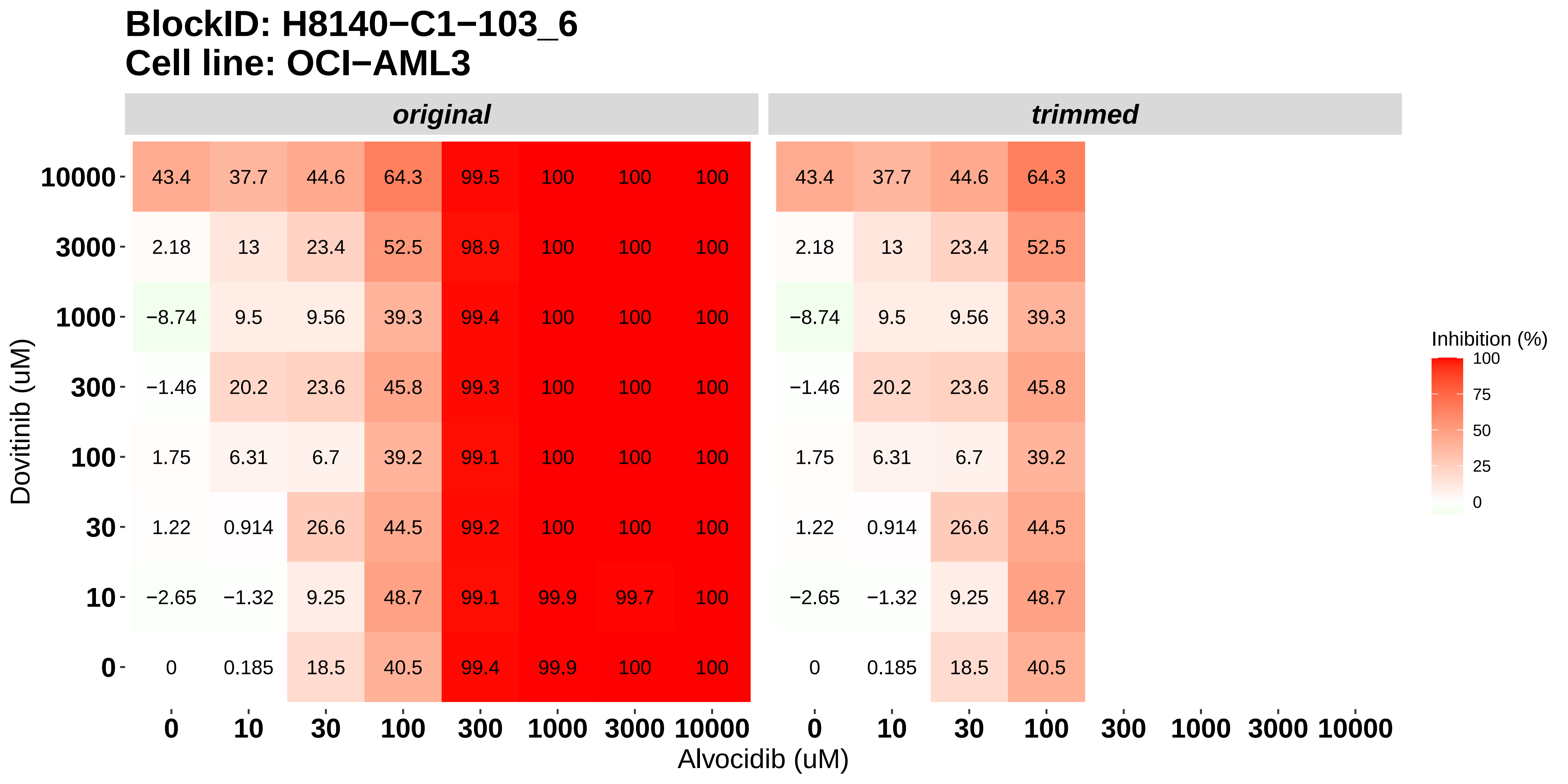

### H8140-C1-201_1_Nintedanib_Alvocidib_MOLM-16.pdf

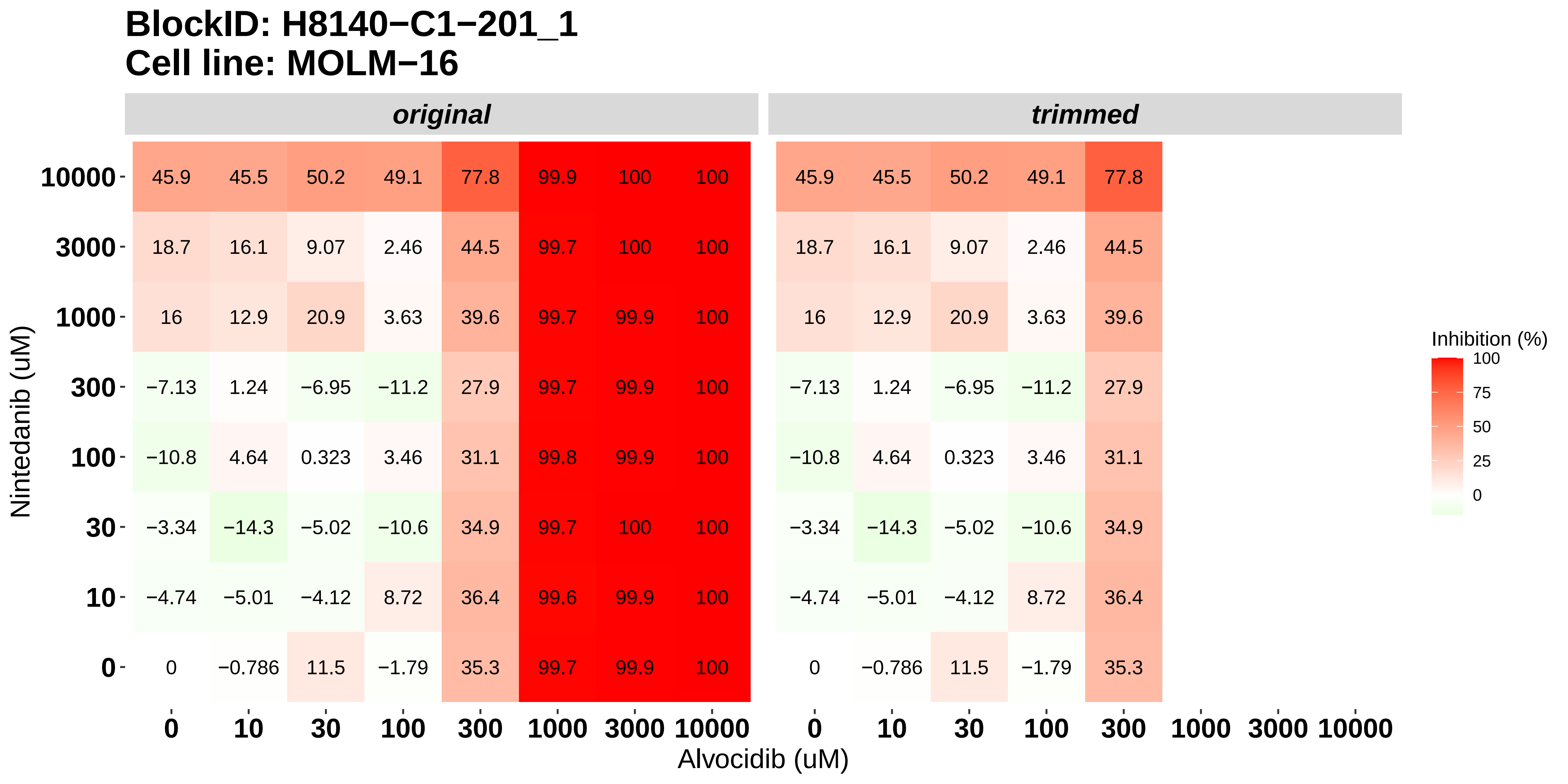

### H8140-C1-201_2_Doramapimod_Alvocidib_MOLM-16.pdf

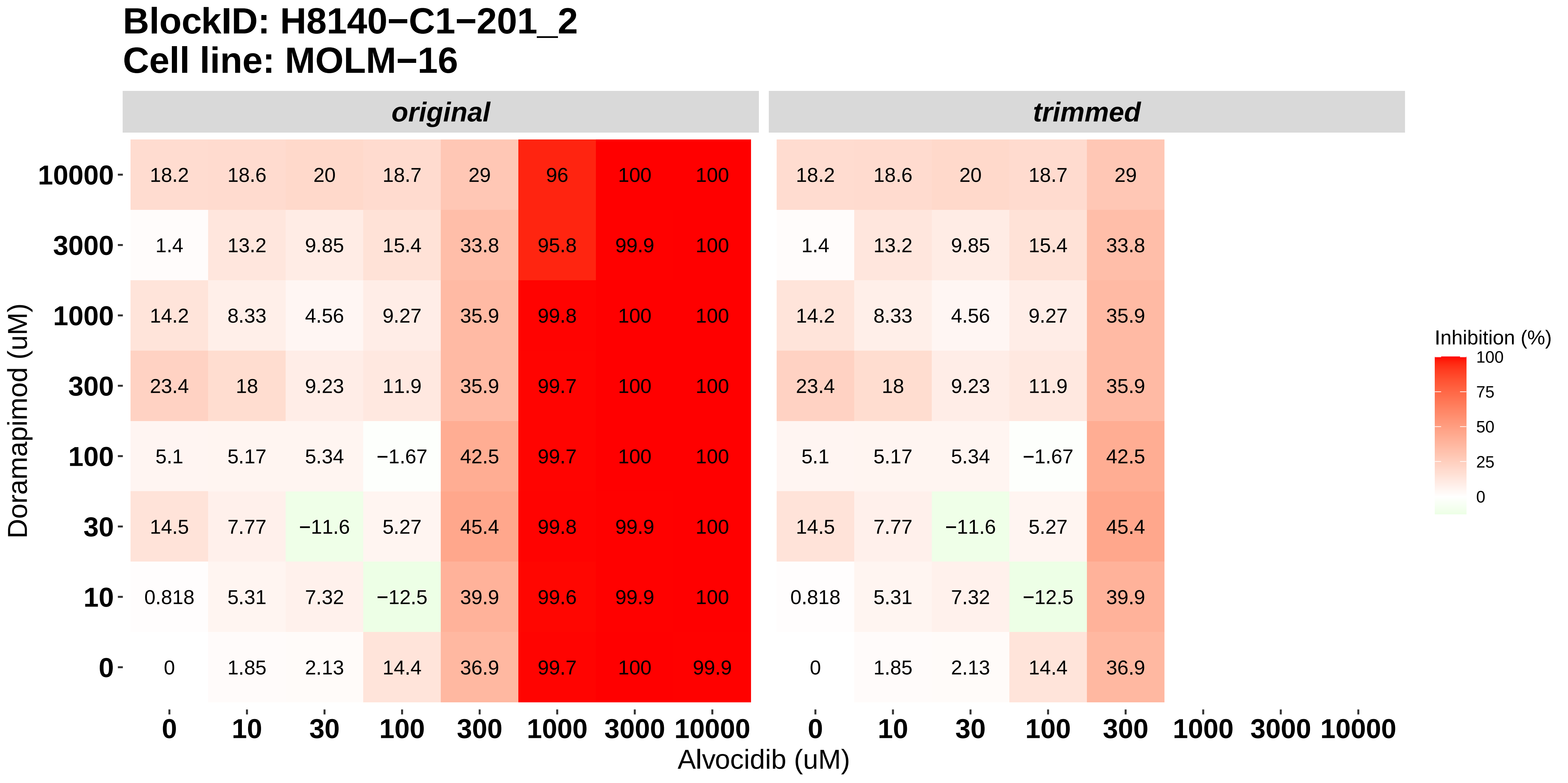

### H8140-C1-201_3_KI20227_Alvocidib_MOLM-16.pdf

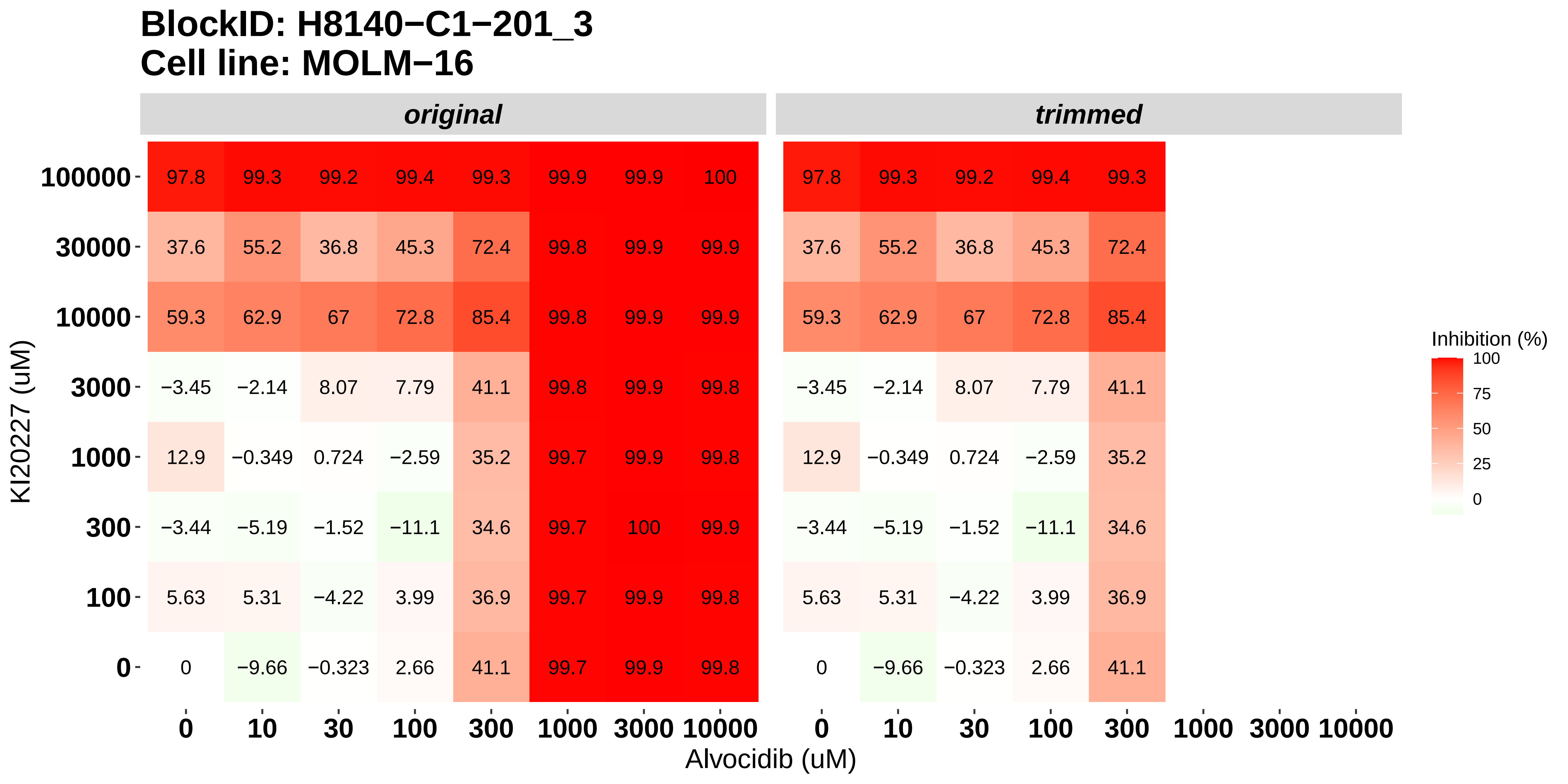

### H8140-C1-201_4_Cabozantinib_Alvocidib_MOLM-16.pdf

#### BlockID: H8140−C1−201_4
#### Cell line: MOLM−16

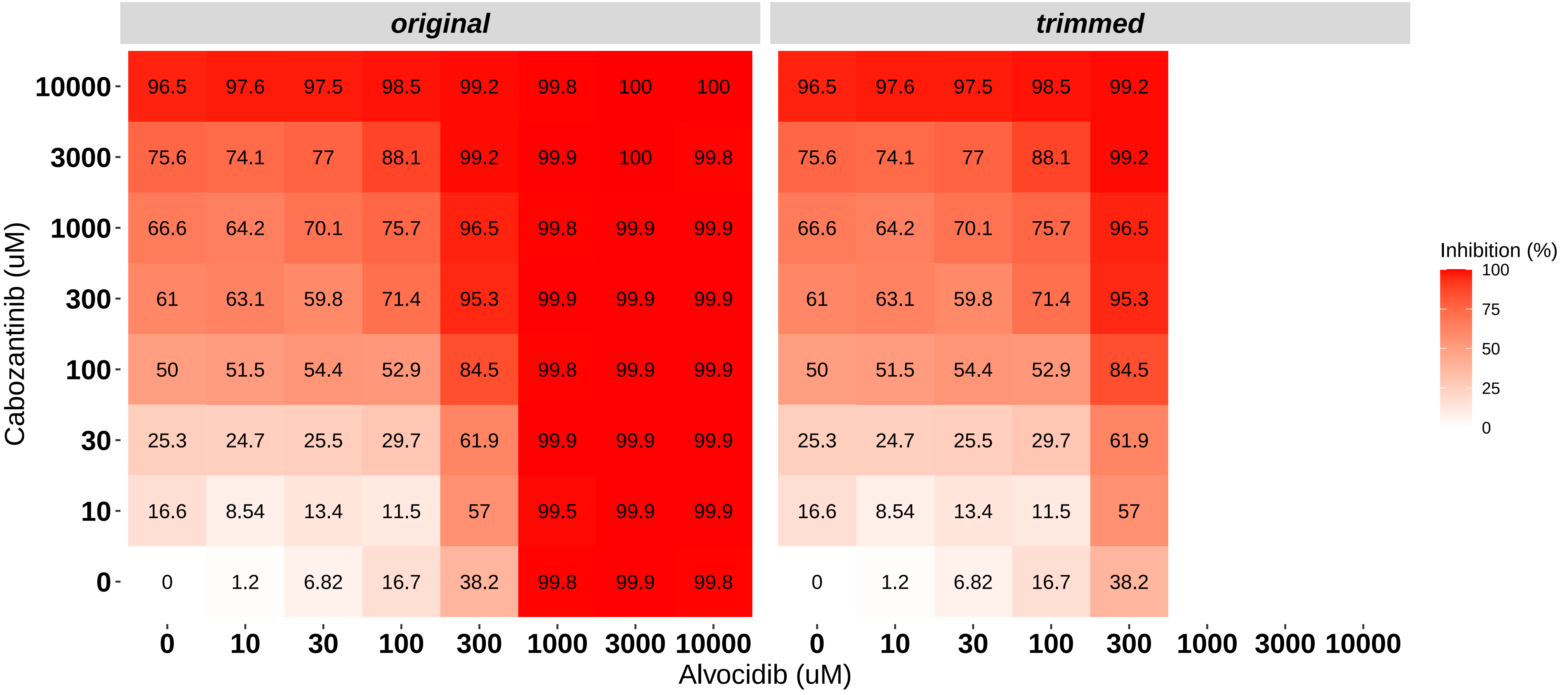

### H8140-C1-201_5_Dovitinib_Panobinostat_MOLM-16.pdf

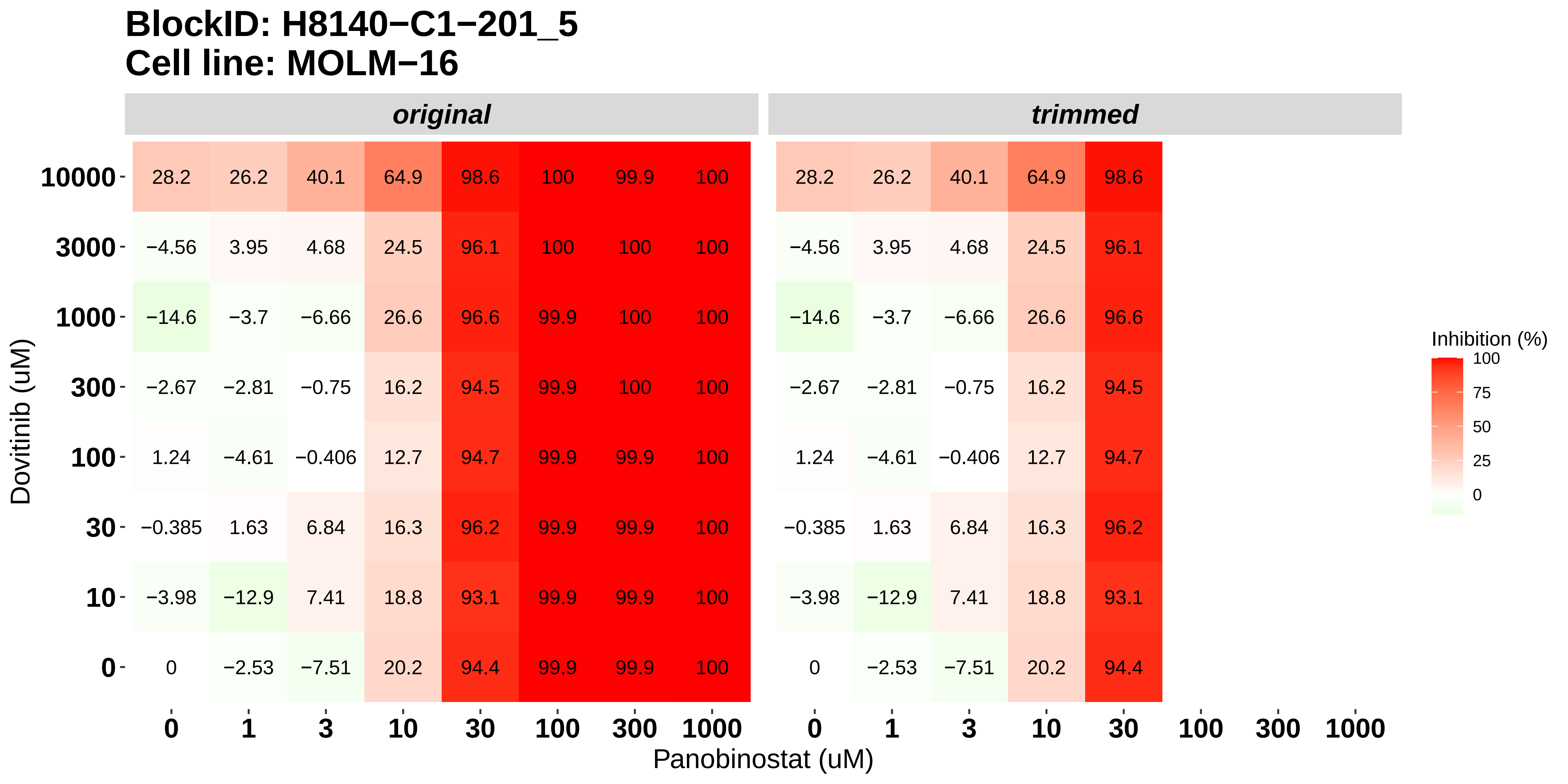

### H8140-C1-201_6_Nintedanib_Panobinostat_MOLM-16.pdf

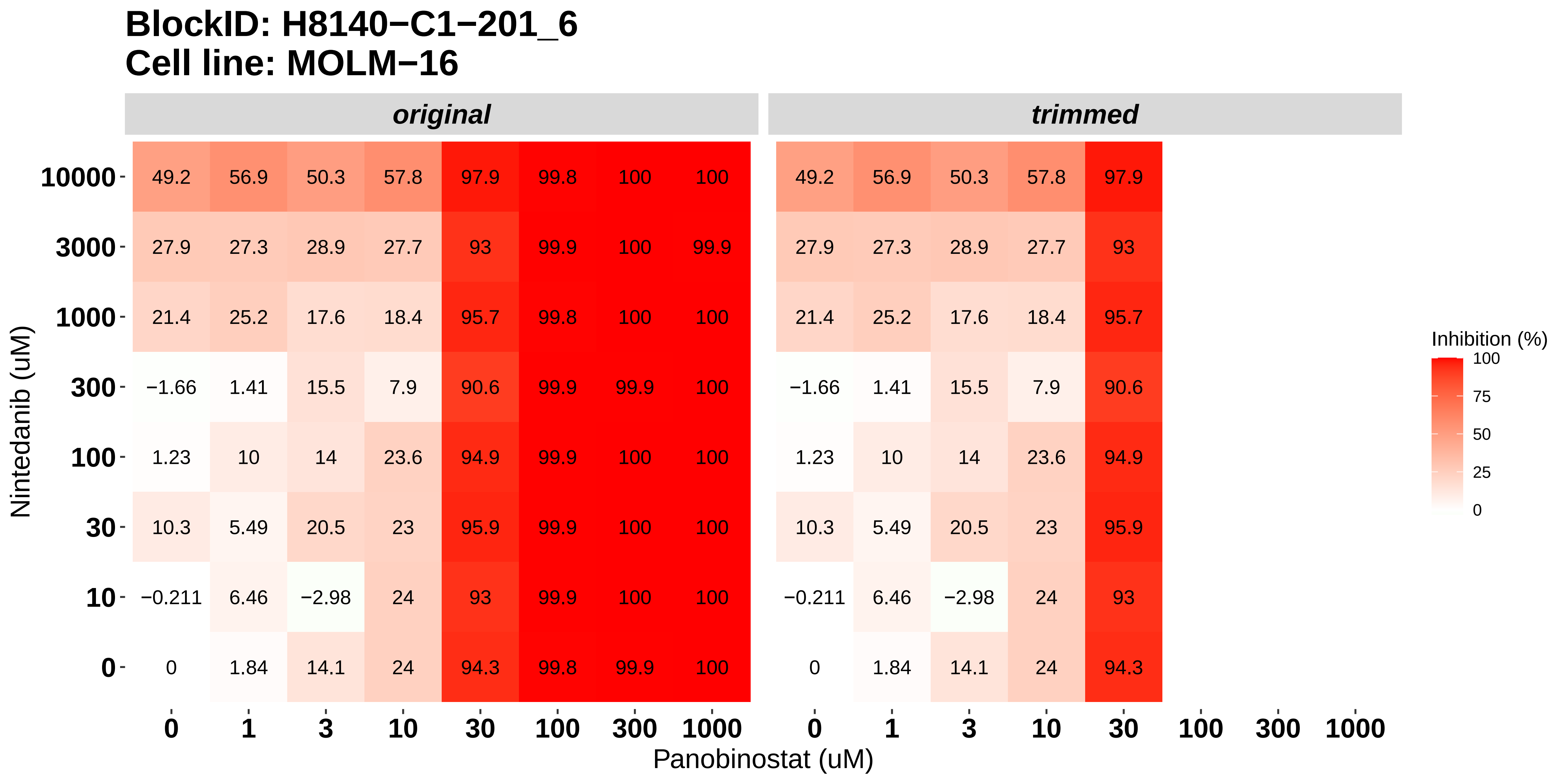

### H8140-C1-202_1_Nintedanib_Alvocidib_NOMO-1.pdf

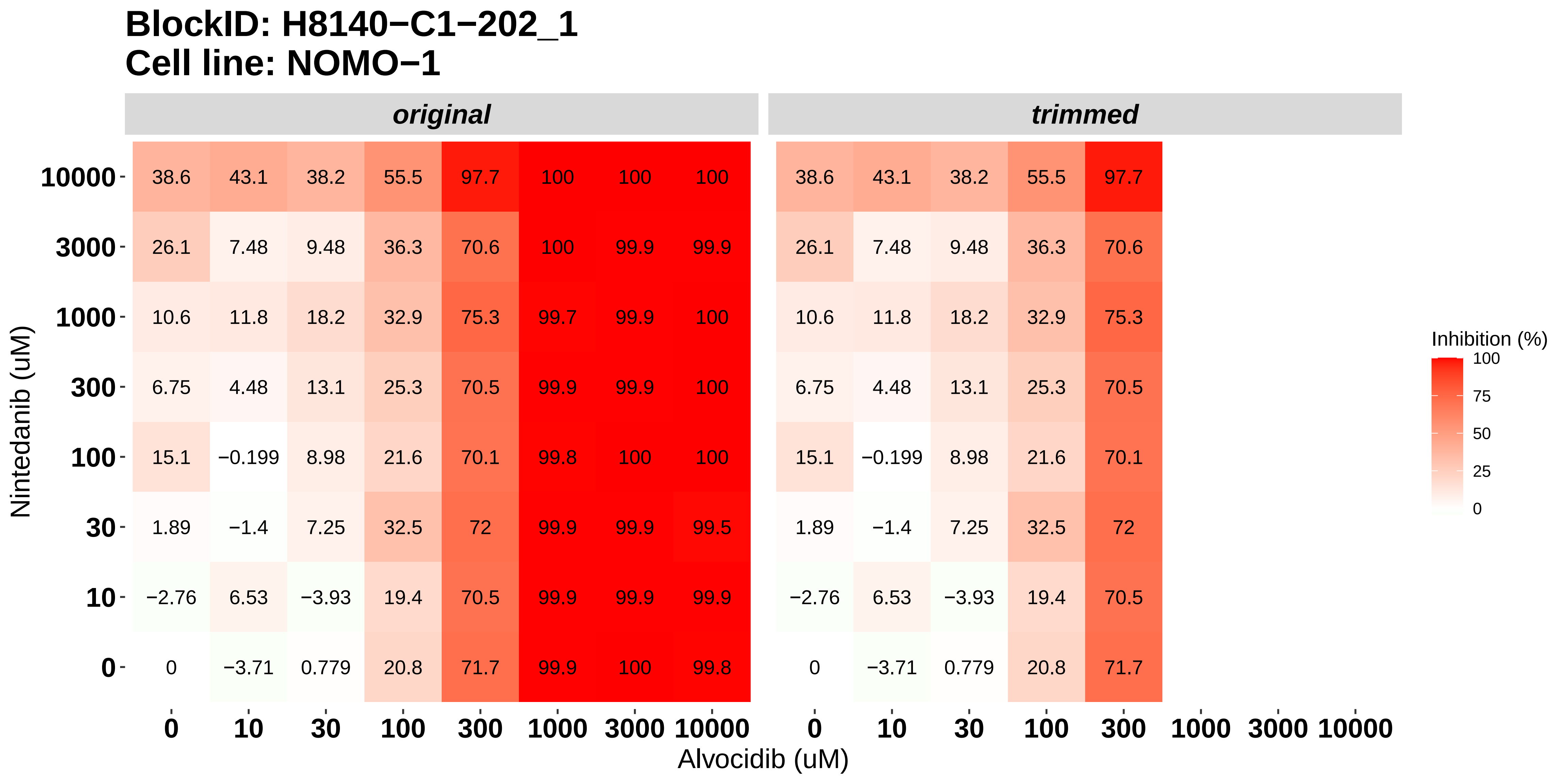

### H8140-C1-202_2_Doramapimod_Alvocidib_NOMO-1.pdf

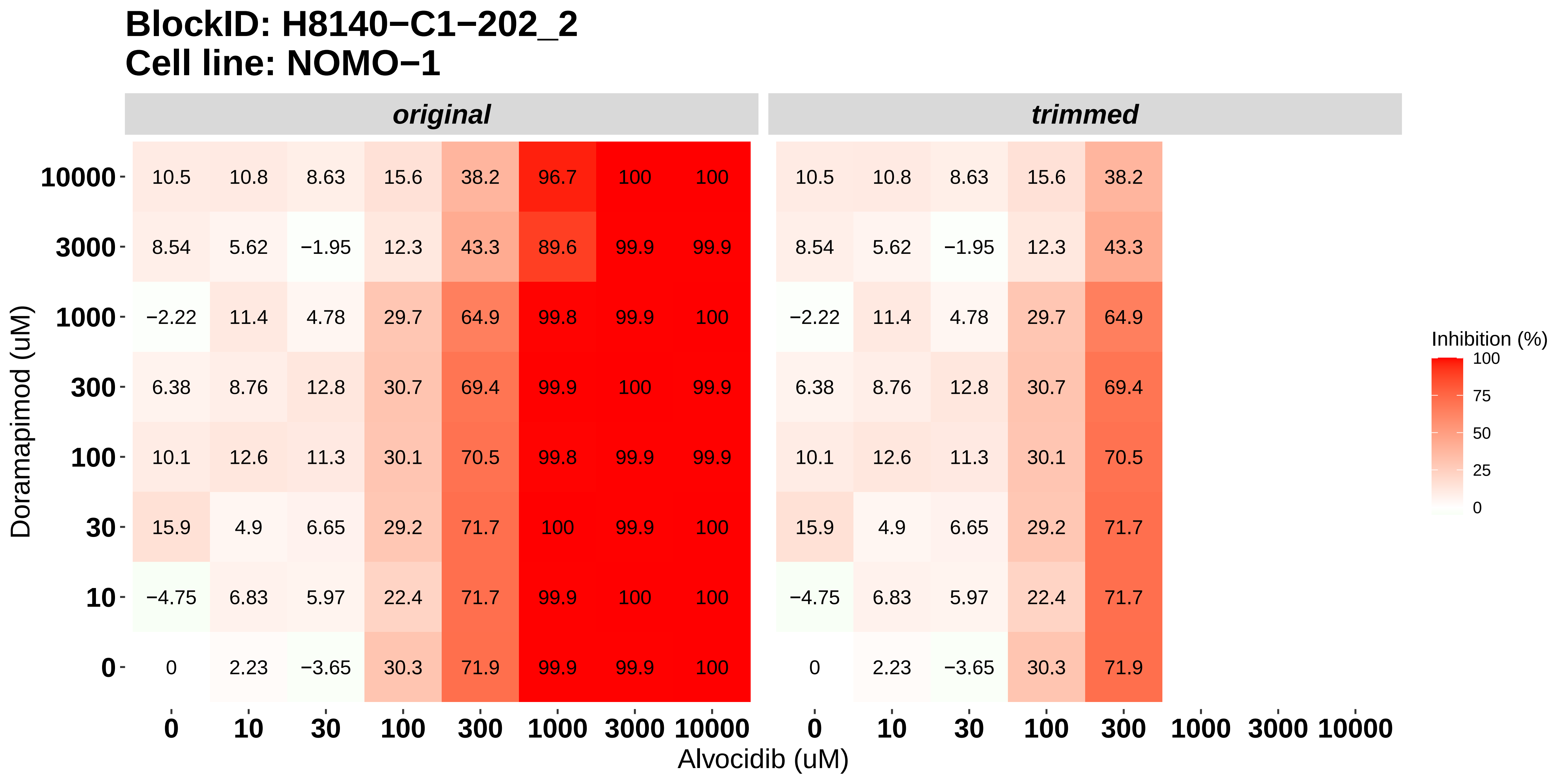

### H8140-C1-202_3_KI20227_Alvocidib_NOMO-1.pdf

#### BlockID: H8140−C1−202_3
#### Cell line: NOMO−1

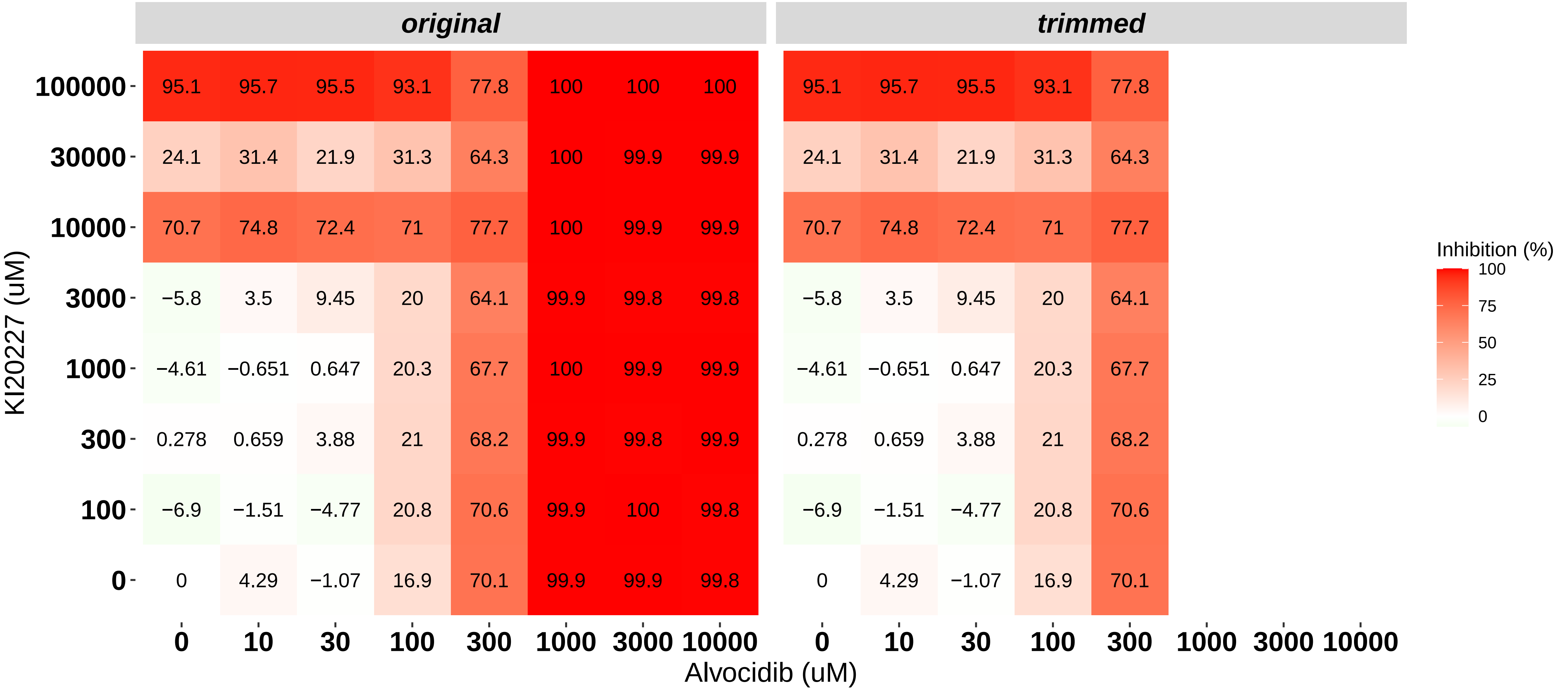

### H8140-C1-202_4_Cabozantinib_Alvocidib_NOMO-1.pdf

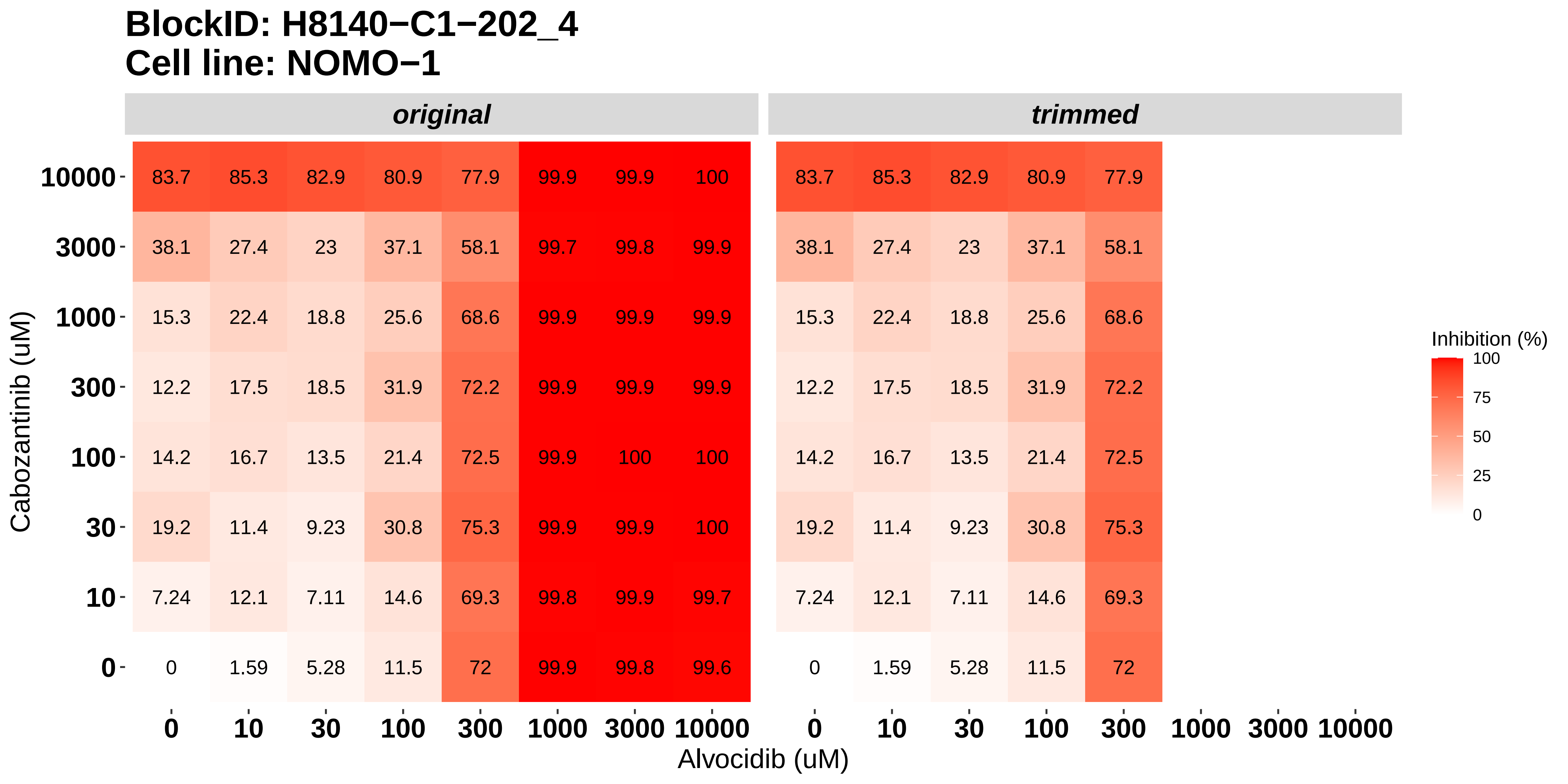

### H8140-C1-202_5_Dovitinib_Panobinostat_NOMO-1.pdf

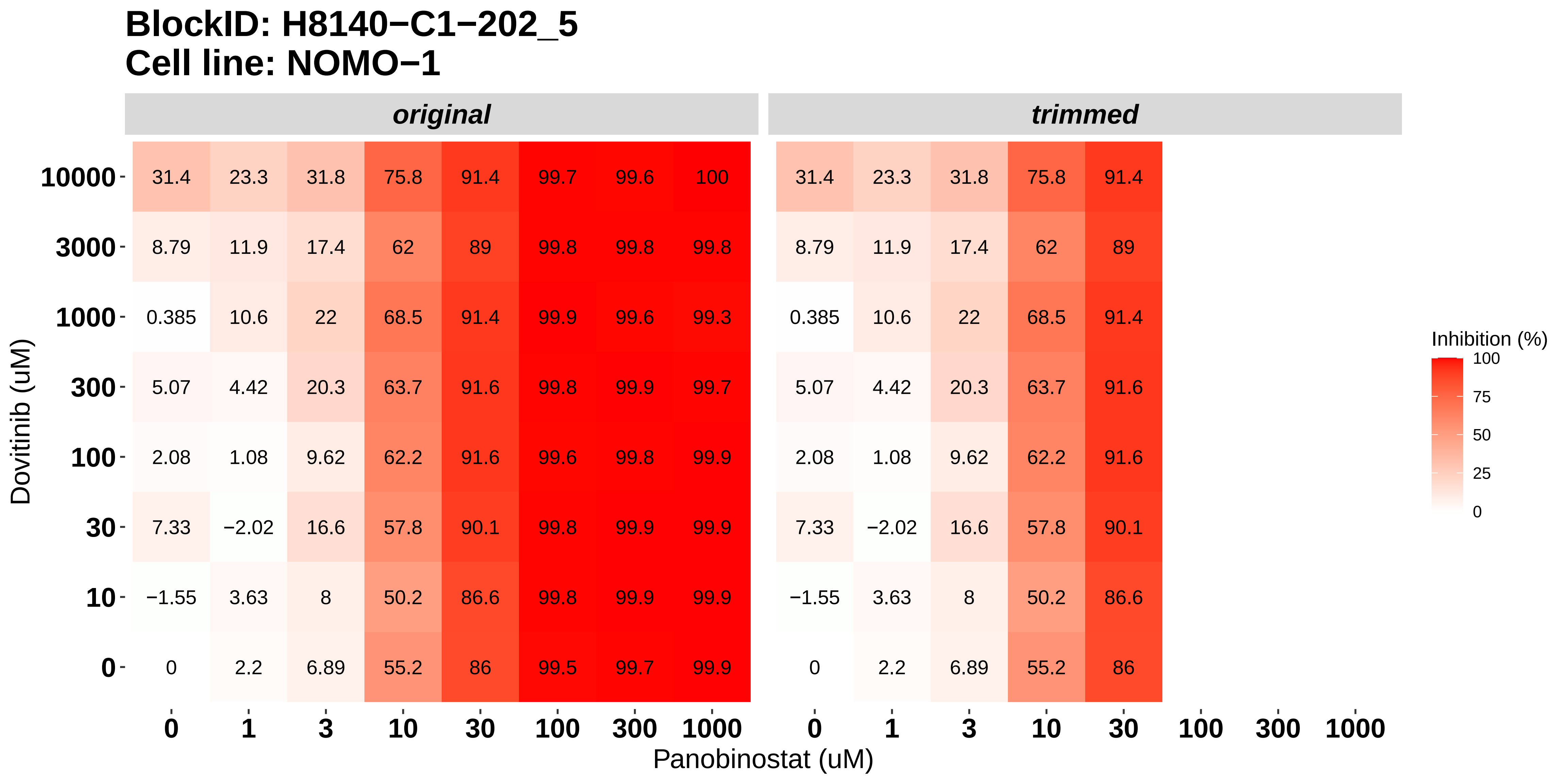

### H8140-C1-202_6_Nintedanib_Panobinostat_NOMO-1.pdf

#### BlockID: H8140−C1−202_6
#### Cell line: NOMO−1

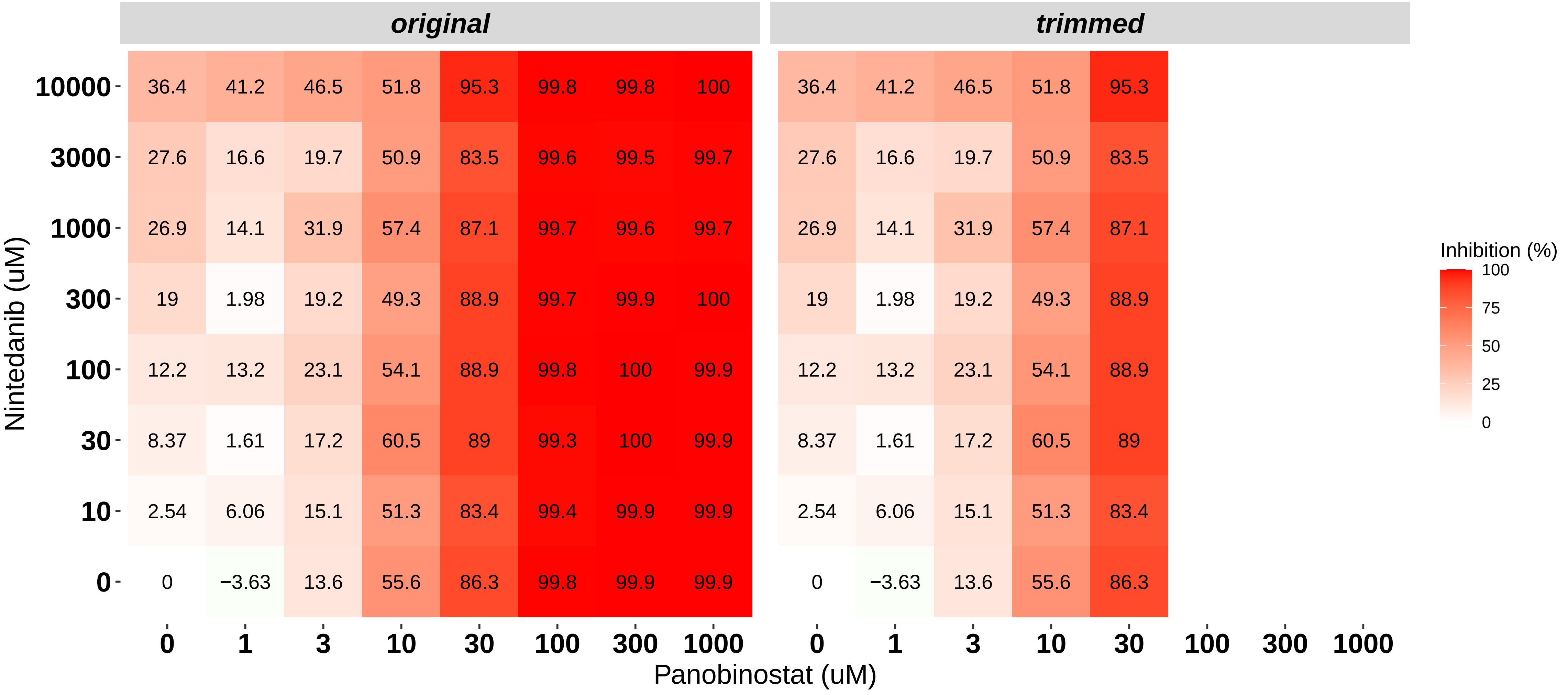

### H8140-C1-203_1_Nintedanib_Alvocidib_OCI-AML3.pdf

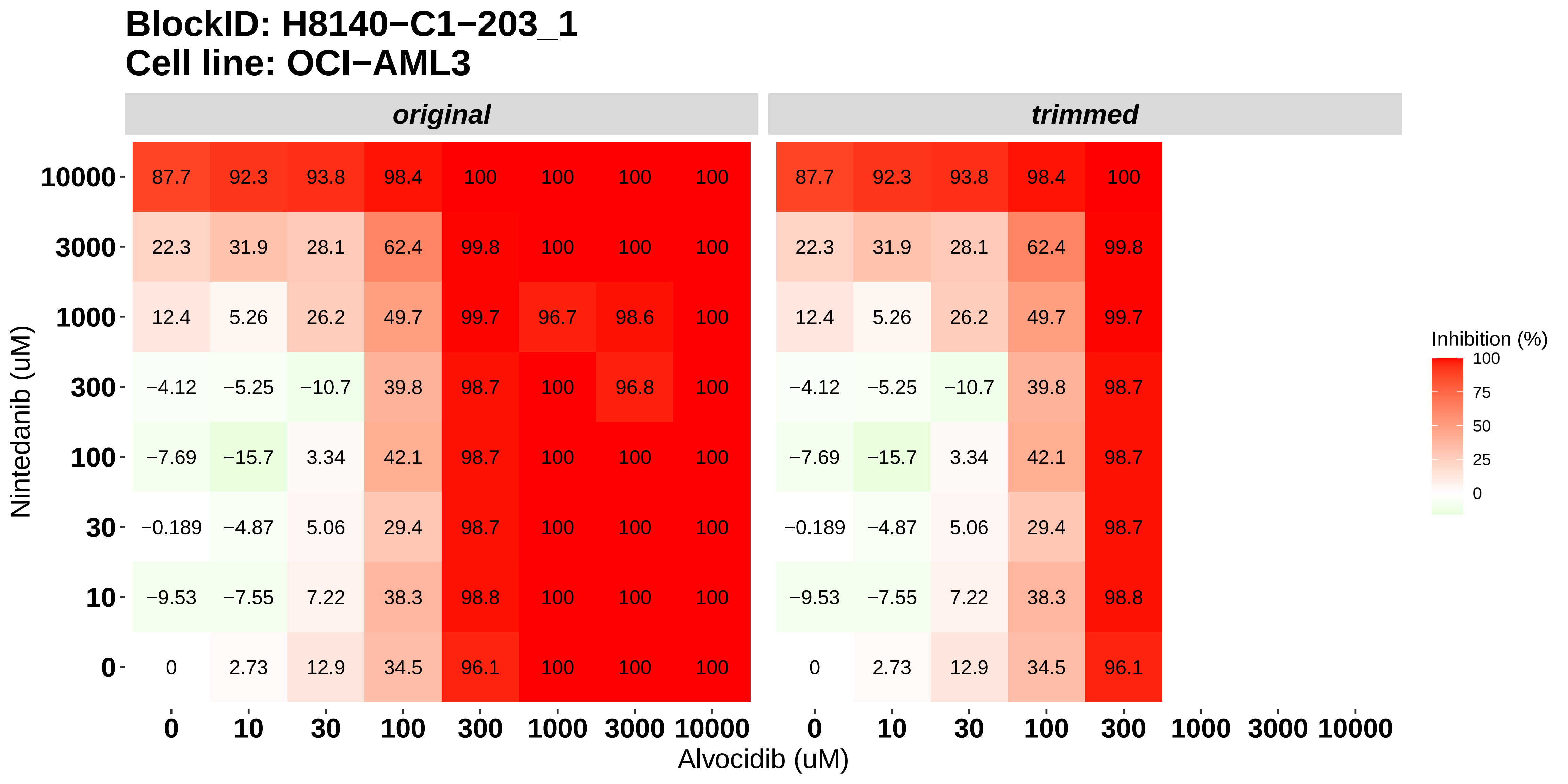

### H8140-C1-203_2_Doramapimod_Alvocidib_OCI-AML3.pdf

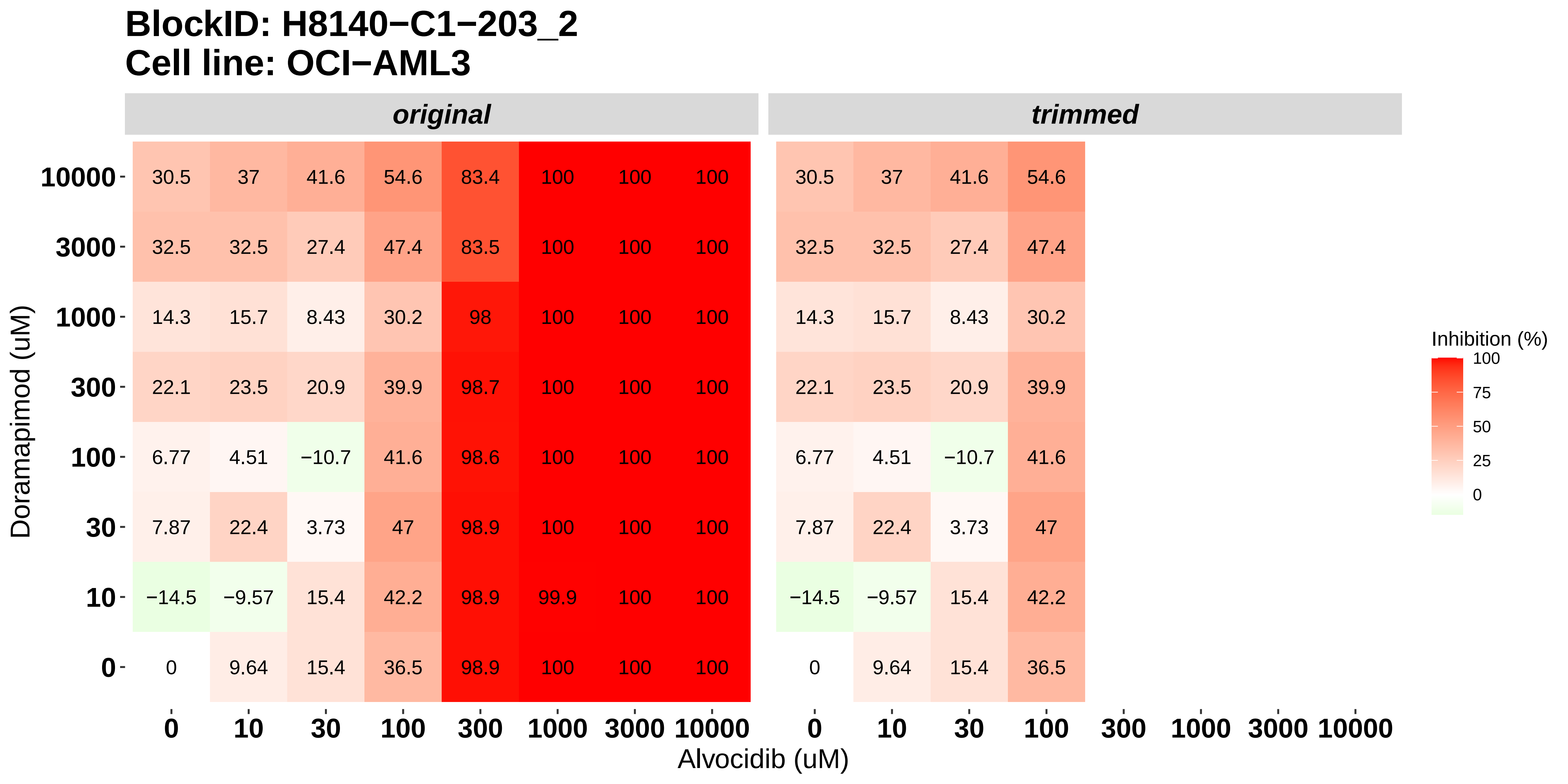

### H8140-C1-203_3_KI20227_Alvocidib_OCI-AML3.pdf

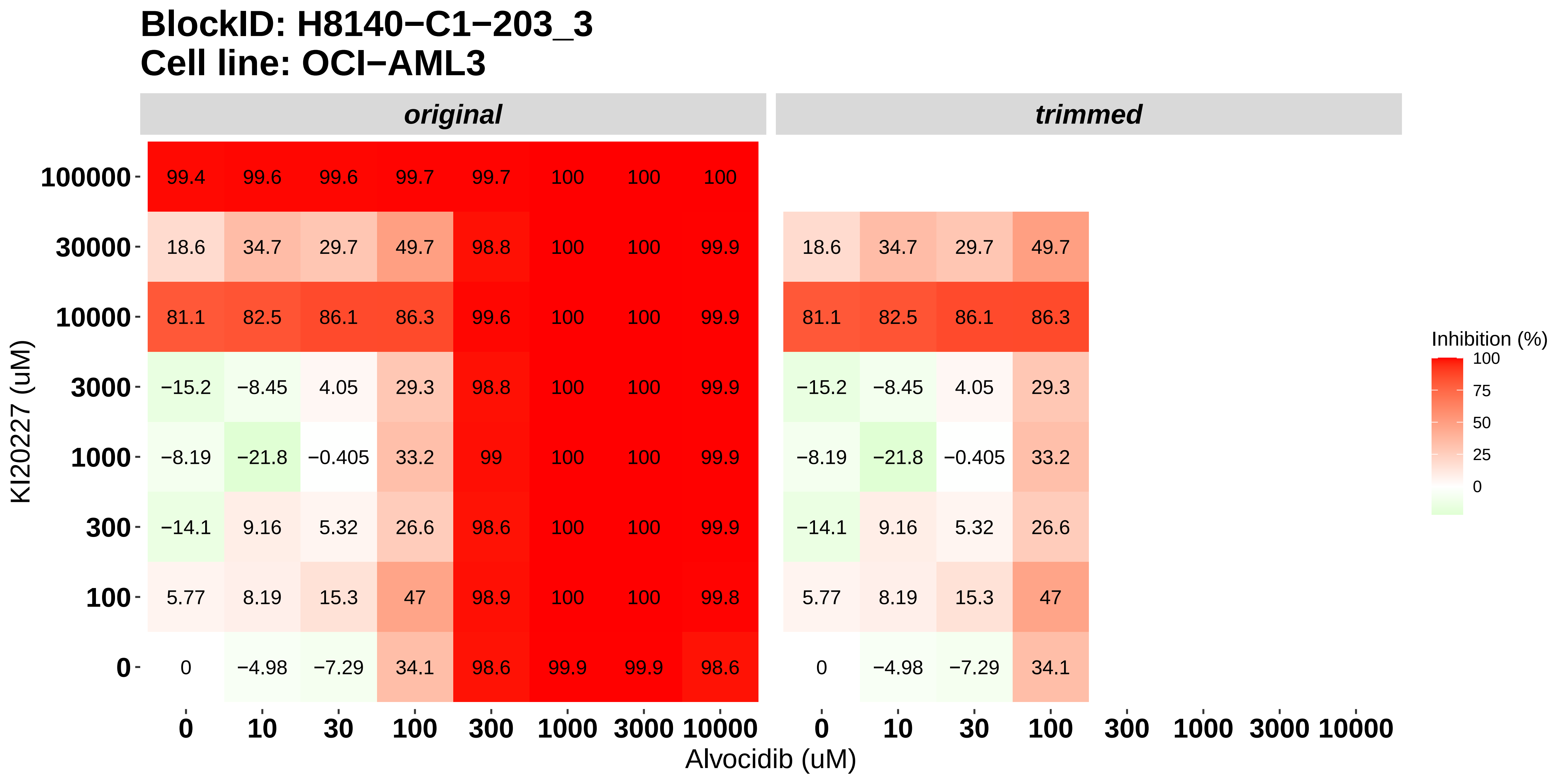

### H8140-C1-203_4_Cabozantinib_Alvocidib_OCI-AML3.pdf

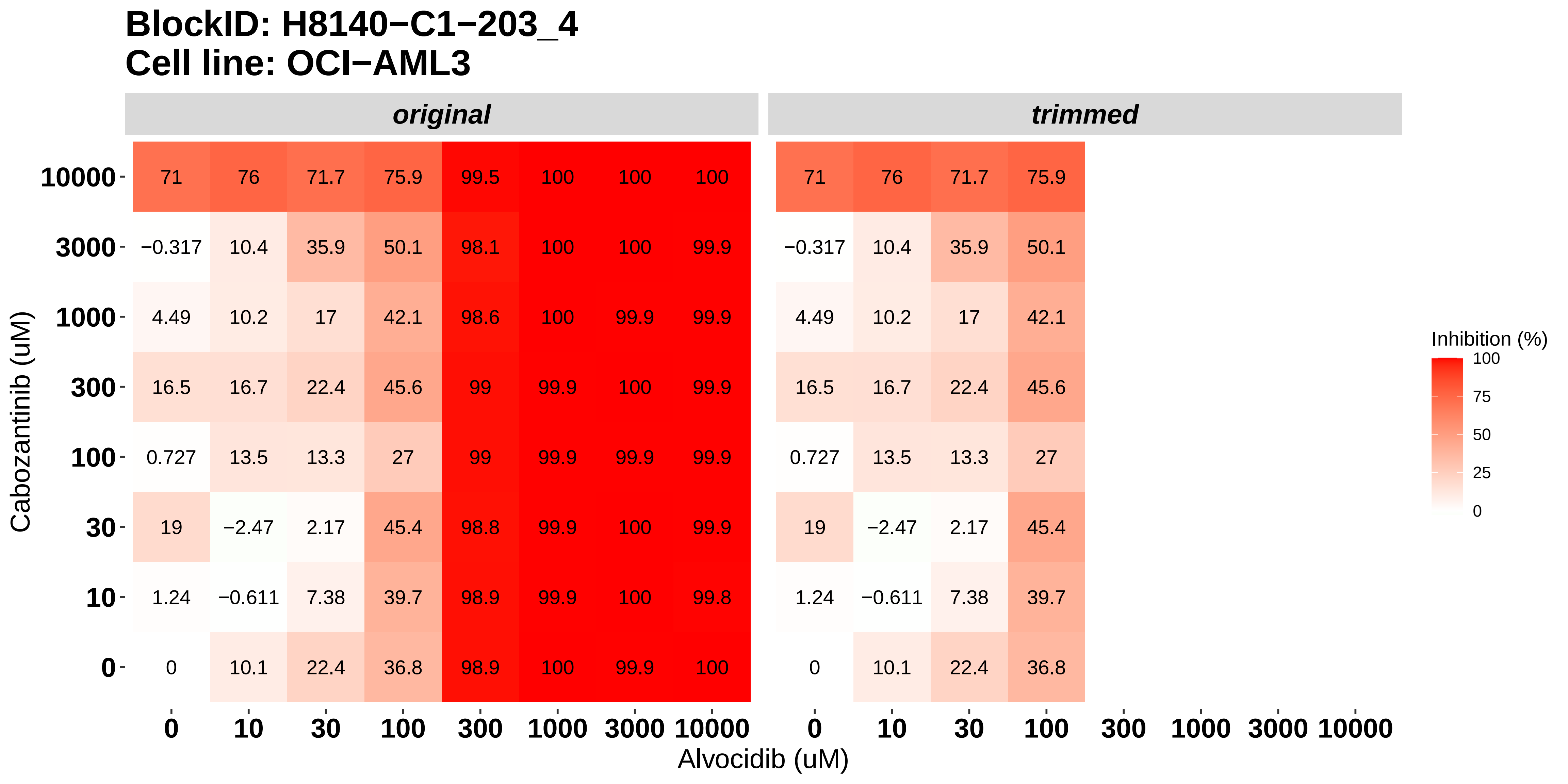

### H8140-C1-203_5_Dovitinib_Panobinostat_OCI-AML3.pdf

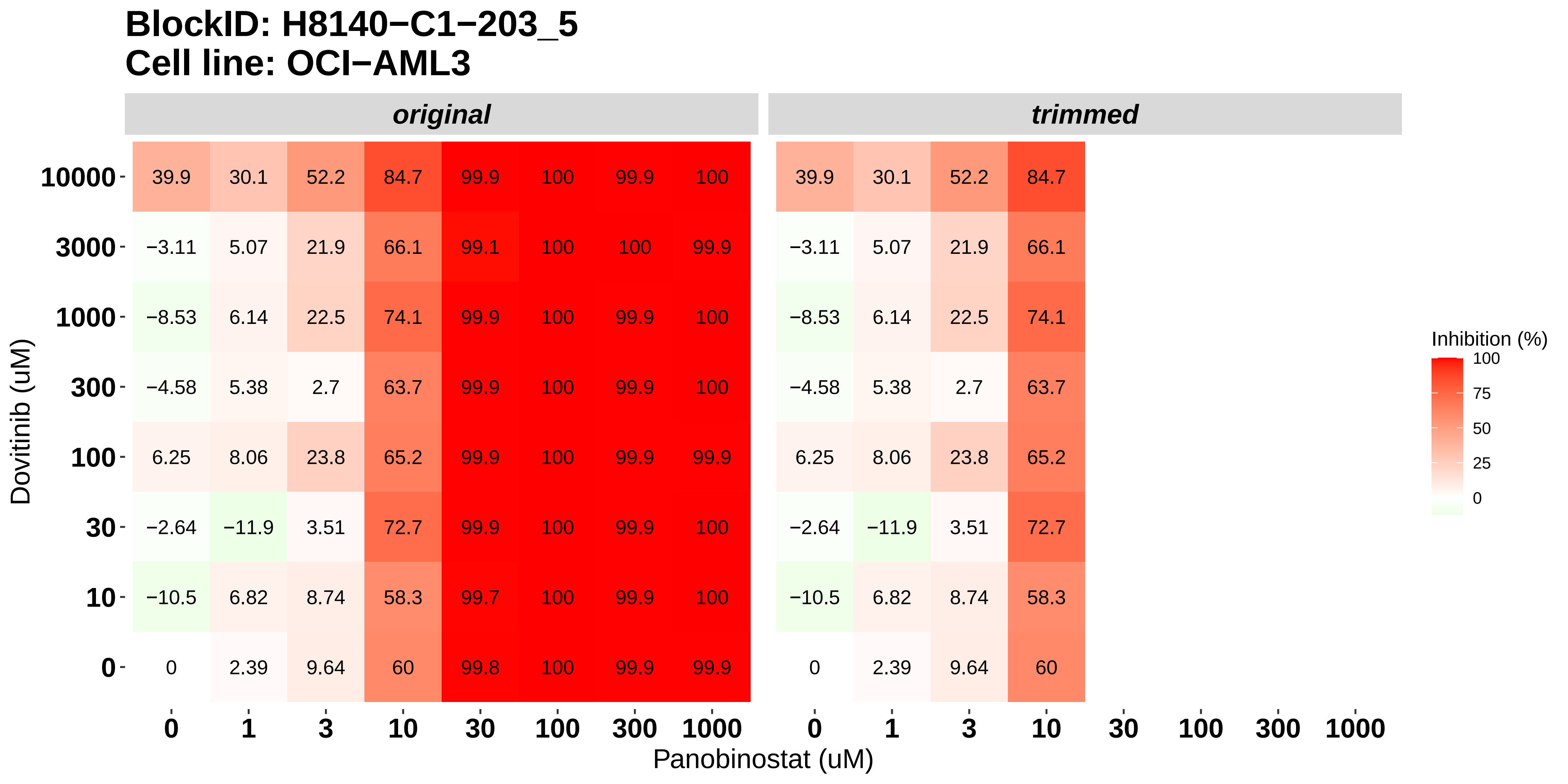

### H8140-C1-203_6_Nintedanib_Panobinostat_OCI-AML3.pdf

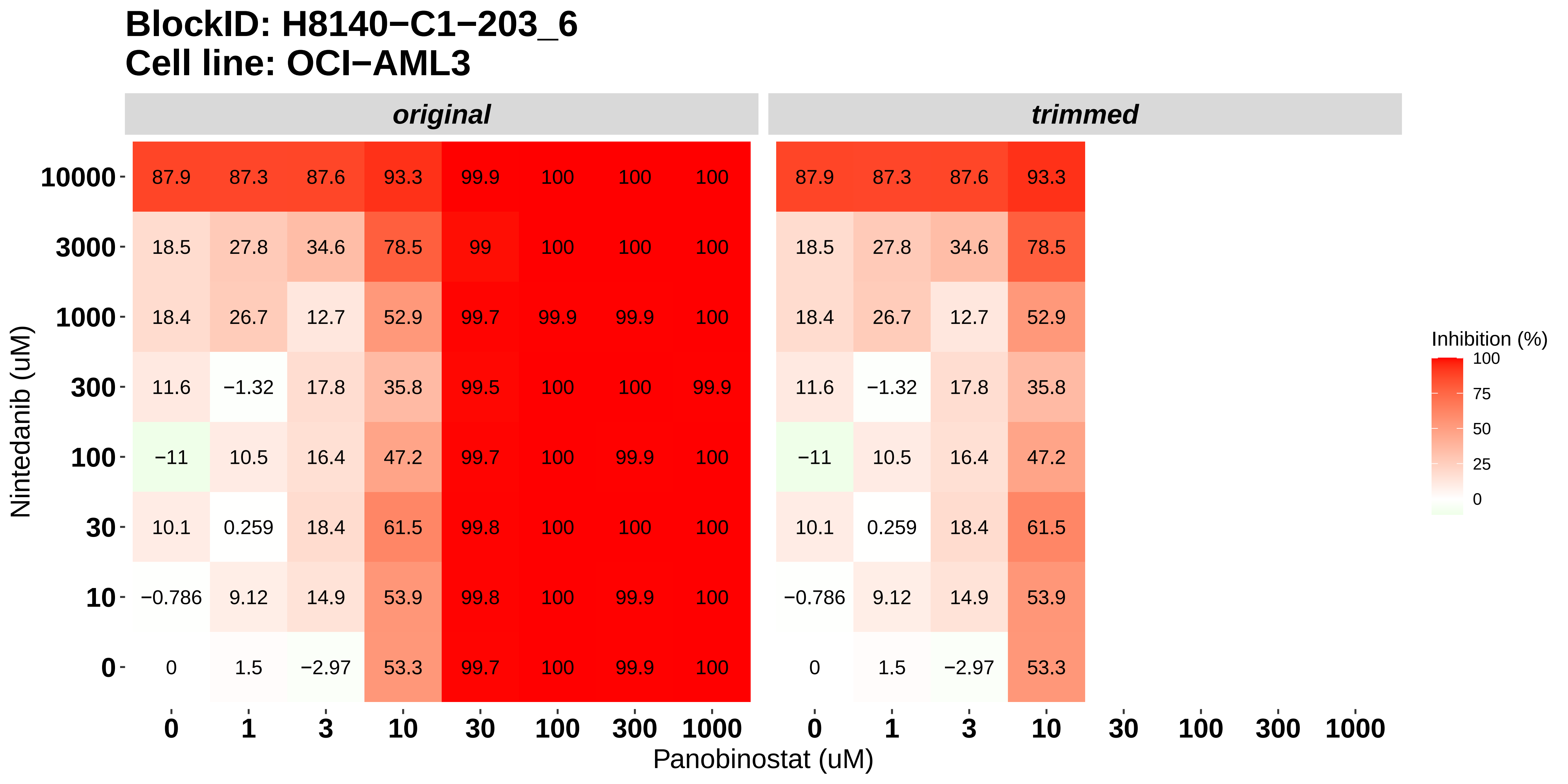
